# Local adaptation fuels cryptic speciation in terrestrial annelids

**DOI:** 10.1101/872309

**Authors:** Daniel Fernández Marchán, Marta Novo, Nuria Sánchez, Jorge Domínguez, Darío J. Díaz Cosín, Rosa Fernández

## Abstract

Uncovering the genetic and evolutionary basis of cryptic speciation is a major focus of evolutionary biology. Next Generation Sequencing (NGS) allows the identification of genome-wide local adaptation signatures, but has rarely been applied to cryptic complexes - particularly in the soil milieu - as is the case with integrative taxonomy. The earthworm genus *Carpetania*, comprising six previously suggested putative cryptic lineages, is a promising model to study the evolutionary phenomena shaping cryptic speciation in soil-dwelling lineages. Genotyping-By-Sequencing (GBS) was used to provide genome-wide information about genetic variability between seventeen populations, and geometric morphometrics analyses of genital chaetae were performed to investigate unexplored cryptic morphological evolution. Genomic analyses revealed the existence of three cryptic species, with half of the previously-identified potential cryptic lineages clustering within them. Local adaptation was detected in more than 800 genes putatively involved in a plethora of biological functions (most notably reproduction, metabolism, immunological response and morphogenesis). Several genes with selection signatures showed shared mutations for each of the cryptic species, and genes under selection were enriched in functions related to regulation of transcription, including SNPs located in UTR regions. Finally, geometric morphometrics approaches partially confirmed the phylogenetic signal of relevant morphological characters such as genital chaetae. Our study therefore unveils that local adaptation and regulatory divergence are key evolutionary forces orchestrating genome evolution in soil fauna.

## Introduction

Cryptic species are biological entities that cannot readily be distinguished morphologically - as defined principally through humans’ visual perception of morphology - yet evidence indicates that they followed different evolutionary trajectories (Bickford et al., 2007). The evolutionary processes underlying the origins of cryptic species remain largely unknown, therefore hampering our understanding on how they adapt to new environments and how frequently they exchange genes with each other, and consequently having an effect on biodiversity assessment. For instance, while in species with high morphological disparity (such as butterflies or fish) divergent selection often leads to sister species with markedly different body shapes or colours (Jiggins et al., 2001; Langerhans et al., 2007), in other species it might affect traits causing reproductive isolation with no clear morphological basis, such as behaviours (Janzen et al., 2009; Jones et al., 2016).

Identifying and defining cryptic species is even more challenging in the case of soil fauna due to their marked degree of morphological stasis, where diagnostic morphological characters are often scarce. This lack of morphological diversification could result from a series of factors, including low standing genetic variation and/or developmental constraints on the morphospace (Bickford et al., 2007; Appeltans et al., 2012), and a relatively constant through time environment coupled to strong stabilizing selection leading to retention of a common, shared morphology (reviewed in Marchán et al. (2018a) and Struck et al. (2018). The answer to this may rely on understanding how adaptation to different environments orchestrates speciation and to what extent it depends on morphological change. Another aspect that should not be disregarded is the existence of differences in unexplored (cryptic) morphological characters, as in the case of pseudocryptic or pseudo-sibling species (Knowlton, 1993), particularly in groups were taxonomy is daunting.

Local adaptation occurs when a population of organisms have higher average fitness in their local environment compared to individuals elsewhere (Kaweki and Ebert 2004). As environments vary across space and time, local conditions determine which traits are favoured by natural selection. Next Generation Sequencing (NGS) techniques have definitely facilitated the identification of some of the loci responsible for adaptive differences among populations. Two basic approaches for identifying putatively locally adaptive loci are mainly used: the identification of loci with unusually high genetic differentiation among populations (differentiation outlier methods), and the search for correlations between local population allele frequencies and local environments (genetic-environment association methods) (Hoban et al., 2016), the latter definitively more challenging to measure in soil environments or ecosystems. As natural selection acts on phenotypic traits, detecting the loci that accumulate high genetic differentiation would elucidate which loci are eventually responsible for such phenotypes. Such an approach has been largely unexplored in cryptic taxa. In a similar and complementary way, integrative taxonomy has yet to be widely applied to cryptic taxonomic complexes, especially through the application of state-of-the-art methodologies such as high-throughput sequencing or geometric morphometrics. To date, approaches aiming at exploring all these different levels of variation in cryptic lineages - particularly in soil fauna - are limited.

Cryptic diversity is widespread in annelids (e.g., Struck et al., 2017), particularly amongst earthworms (King et al., 2008; Novo et al., 2010; Shekhovtsov et al., 2013; Taheri et al., 2018). One of the most studied cryptic species groups among these animals is the former *Hormogaster elisae* Álvarez, 1977 (Annelida, Oligochaeta, Hormogastridae), recognized as the genus *Carpetania* after Marchán et al. (2018b). Six highly divergent cryptic lineages were identified using a set of mitochondrial and nuclear markers (Marchán et al., 2017), but their validation as species and consequent description has been hindered by the absence of clear-cut limits between the putative species. This species complex shows high mitochondrial divergence between lineages (11-17% cytochrome C oxidase subunit I average uncorrected pairwise distance) confirmed by segregation of nuclear haplotypes, as well as an ancient estimated divergence age (60-35 mya) (Marchán et al. 2017). Those lineages display strong spatial isolation with non-overlapping ranges separated by narrow borders and clear hints of reproductive isolation shown by cross-breeding experiments (Marchán et al. 2017). In addition, slight variation in some cryptic morphological characters (genital chaetae, relative position of septa and spermathecae) was found promising but was not fully tested in a comparative framework (Marchán et al., 2017, 2018b). These characteristics make the *Carpetania* complex a promising model to further explore the underlying evolutionary phenomena shaping cryptic speciation in soil-dwelling lineages.

Herein, we test the hypothesis that genome-wide and unnoticed morphological differences may have fueled cryptic speciation in soil fauna by exploring the terrestrial annelid *Carpetania* cryptic complex through an integrative taxonomic approach informed by state-of-the-art methodologies including Genotyping-by-Sequencing (GBS) and geometric morphometrics. Furthermore, we identify and characterize which loci may be putatively involved in local adaptation, which functions they may fulfil in the biology of this species complex and how they may have triggered cryptic speciation.

## Materials and methods

### 1. Specimen sampling

Different *Carpetania* populations were sampled from 2007 to 2015, usually in November (autumn) or February-April (spring) sampling seasons. Information about their localities was published in (Novo et al., 2009, 2010) and Marchán et al. (2017). In all cases, specimens were collected by hand and fixed in the field in ca. 96% EtOH, with subsequent ethanol changes and finally preserved at −20 °C.

Seventeen of those populations (Suppl. Table 1, Figure 1) representing all the lineages and internal clades from Marchán et al. (2017) were chosen for the GBS analysis, with five individuals from each population totaling 85 individuals. When possible, populations at both sides of a border between range of the previously-identified cryptic lineages were chosen.

**Figure 1.**
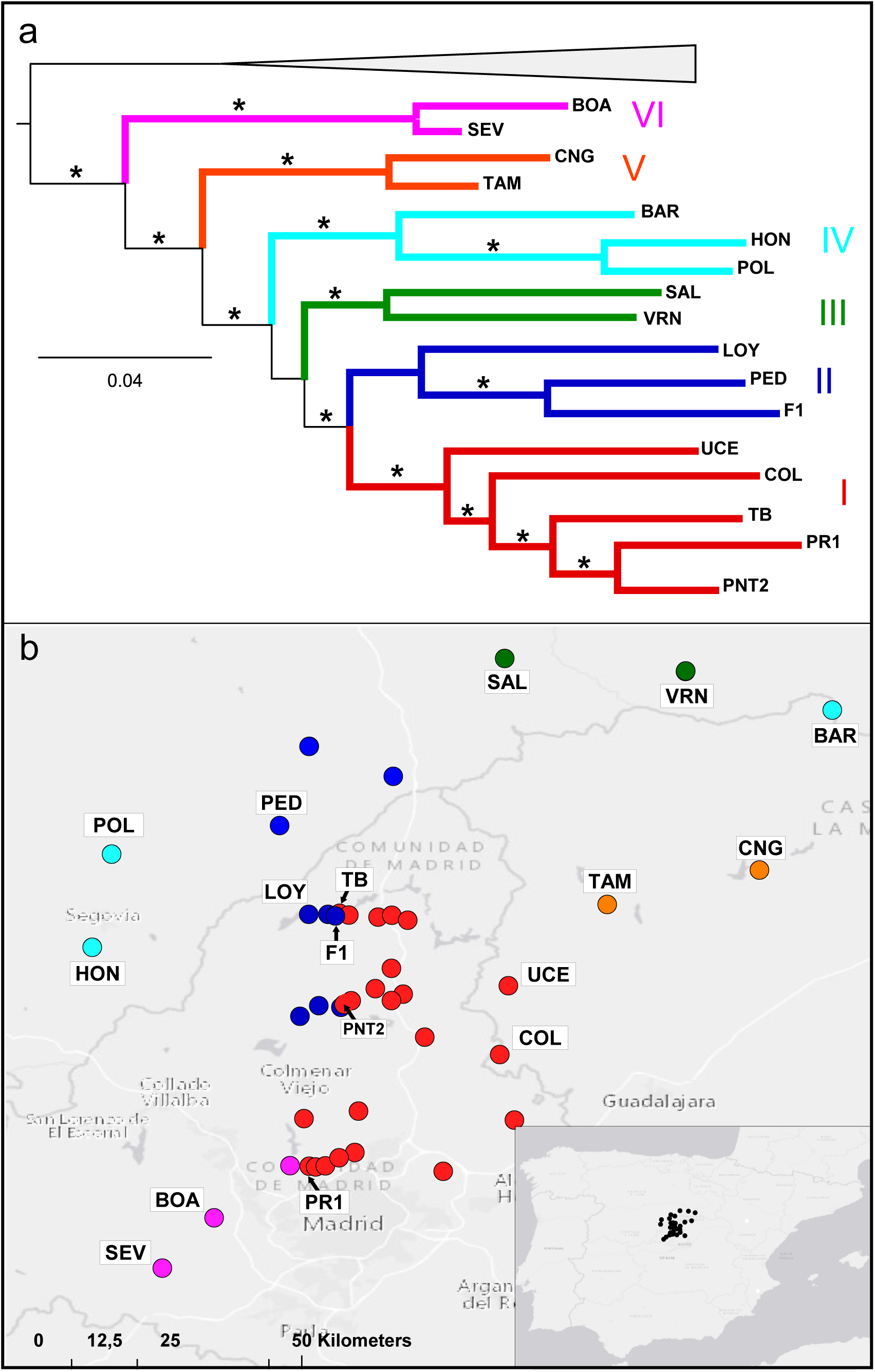
Modified from(Marchán et al., 2017). a) Populations of the *Carpetania elisae* complex included in this work shown in a phylogenetic tree obtained by bayesian inference from a COI, 16S-tRNAs, 28S, H3 concatenated dataset. Populations not included in this work were collapsed. Asterisks show maximum support (99-100%) b) Populations of the *Carpetania elisae* complex included in this work shown in a geographical context. Previously defined cryptic lineages are represented by a colour and roman number. Color codes and nomenclature are kept throughout the manuscript.

### 2. GBS library preparation and sequencing

DNA extraction from gizzard muscle tissue was performed at the Biotechnology Resource Center (BRC) at Cornell University. DNA concentrations were measured and normalized so as all samples had the same concentration when pooling, and a GBS library was prepared using the restriction enzyme *PstI* following the GBS protocol from Elshire et al. (2011). Sequencing was carried out in a NextSeq500 Illumina platform with 85 multiplexed samples on a single lane, with a read length of 75 bp reads.

### 2. SNP calling and filtering

SNP calling was performed using the software STACKS2 v 2.3e (Catchen et al., 2013). Raw reads were quality-filtered and demultiplexed according to individual barcodes using the script *process_radtags.pl* as implemented in STACKS (Catchen et al., 2013). GBS loci were further assembled, and SNPs were called using the *denovo_map.pl* pipeline also implemented in STACKS2. A first dataset (‘de novo-all SNPs’ hereafter) used for subsequent phylogenetic reconstruction was built using a minimum coverage to identify a stack of 3X (−m 3), a maximum number of differences between two stacks in a locus in each sample of five (−M 3), and a maximum number of differences among loci to be considered as orthologous across multiple samples of five (−n 5). The script *export_sql.p*l in the STACKS2 package was used to extract locus information from the catalogue, filtering for a maximum number of missing samples per locus of 50%. The function *populations* in the STACKS2 package was used to export a dataset of full sequences and a dataset of SNPs of the filtered loci in VCF and Phylip formats. A second dataset (‘de novo-one SNP’ hereafter) was inferred following the same pipeline but just selecting a random SNP per locus in order to leverage further phylogenetic analysis (see below). In addition, we constructed a third dataset (‘reference-one SNP’) in a similar way as described above but mapping against a reference transcriptome of *Carpetania elisae* lineage 1 (collected in El Molar, Spain), therefore calling only SNPs from protein coding genes (including both protein coding regions - CDS - and untranslated region - UTR, the regions of an mRNA directly upstream (5′-UTR) or downstream (3′-UTR) from the initiation codon). Transcriptome reads for the reference transcriptome assembly were retrieved from Novo et al. (2013) (NCBI Short Read Archive project: PRJNA196484) and the assembly assembled using Trinity v. r2013-08-14 (Haas et al., 2013) was provided by the authors and has been deposited in the Harvard Dataverse repository associated to this study (https://doi.org/10.7910/DVN/RVMQND).

These three datasets were used for phylogenetic reconstruction and further analyses of selection.

Genomic diversity and population genomics statistics (number of private alleles, number of polymorphic nucleotide sites across the dataset, percentage of polymorphic loci, average observed and expected heterozygosity per locus, average nucleotide diversity π , average pairwise fixation index (F_ST_) and average inbreeding coefficient F_IS_) were obtained from the output of the *populations* function in STACKS2.

### 3. Phylogenetic analyses

Phylogenetic relationships were inferred from the concatenated sequences of the three SNP datasets using a maximum likelihood (ML) approach as implemented in RAXML-HPC v8 <u>(Stamatakis, 2014)</u> in Cipres Science Gateway (https://www.phylo.org/) with default parameters (GTRCAT model, ascertainment bias correction(Lewis, 2001), 1000 rapid bootstrap inferences).

The multispecies coalescent model was implemented through SVDQuartets (Chifman and Kubatko, 2014) to infer the unrooted species tree. This method has the additional advantage of providing indirect evidence of the presence (or absence) of processes such as introgression or incomplete lineage sorting, displayed by the proportion of quartets (four-taxon subtrees) compatible with the obtained species tree. The three SNP datasets were analyzed in PAUP* 4.0a (Cummings, 2004) with the following settings: evaluate all possible quartets, handling of ambiguities: distribute and 100 bootstrap replicates. Two types of analysis were run: using a taxon partition to assign individual sequences to the six lineages from Marchán et al. (2017) and without taxon grouping.

### 4. Genetic structure

In order to visualize major trends of genetic structure on the studied individuals, a principal component analysis (PCA) was performed using the function *glPCA* in the R package adegenet v2.0.1(Jombart, 2008; Jombart and Ahmed, 2011). Unlinked SNP datasets were used as input.

Bayesian clustering was performed in STRUCTURE 2.3.4 (Pritchard et al., 2000). STRUCTURE is frequently chosen to study population structure within species, but an increasing number of works have shown its potential to study closely related species: this approach detects the uppermost level of genetic structure, which can correspond to species-level genetic differentiation (Garg et al., 2016). After exploratory runs, the number of genetic clusters K was limited to the range 1-7, and 10 separate runs for each K (1-7) were performed, each consisting on 100,000 generations of burn-in and 100,000 generations of MCMC sampling. Structure Harvester (Earl and vonHoldt, 2012) was used to identify the optimal number of clusters through the Evanno’s method (ΔK criterion) (Evanno et al., 2005). The output of the different iterations was summarized and visualized in CLUMPAK (Kopelman et al., 2015).

### 5. Genital chaetae extraction and imaging

Thirteen populations were chosen from amongst the previous nineteen to undergo an in-depth morphological analysis of their genital chaetae. Seven of them -BAR, UCE, LOY, SAL, SEV, TAM and TB (see Fig. 1)-were newly studied for this work. The obtained data was combined with the information from the six populations included in Marchán et al. (2016), providing a good representation of the different lineages and their internal clades.

Genital chaetae were extracted from three different adult specimens from each of the seven populations. The chaetae were cleaned from remaining tissues by being treated with hydrogen peroxide. Chaetae from each specimen were pooled together and glued on aluminum stubs using double-sided carbon tape, air-dried and sputter-coated 90 seconds with gold. Scanning electron micrographs (see Fig. 2) were taken on a JEOL JSM-6335F field emission scanning electron microscope.

**Figure 2.**
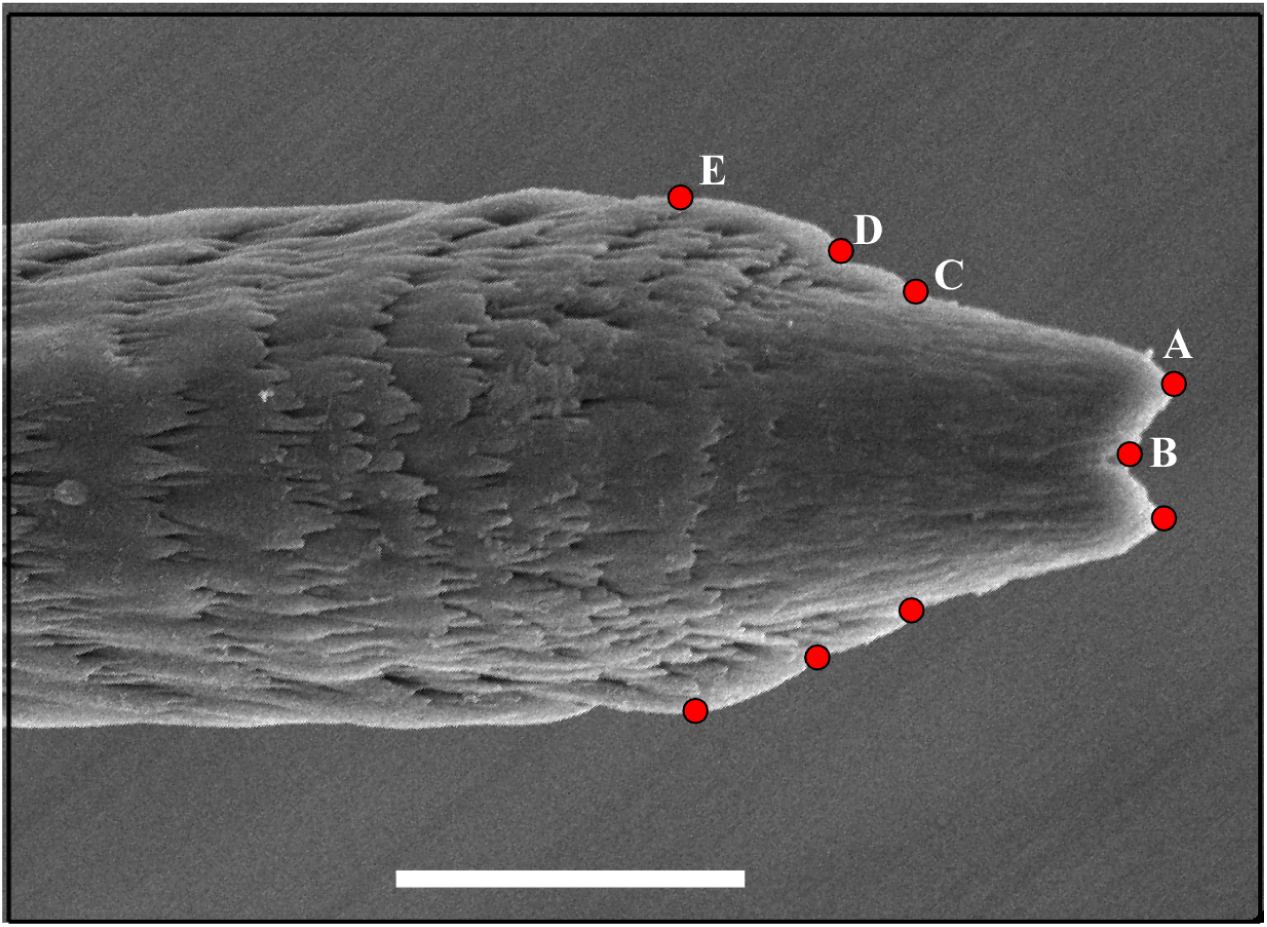
Example of scanning electron micrograph of the distal tip of the genital chaetae of *Carpetania elisae*, showing morphological landmarks chosen for the geometric morphometrics analyses: A) tip of distal denticles, B) mid-point between the distal denticles, C) anterior end of the lateral ridges, D) lateral limits of the first clearly developed ring, E) points of maximum width between the lateral ridges.

### 6. Geometric morphometrics

Geometric morphometrics analyses of the genital chaetae were performed using the software tpsUtil and tpsDig2 for the acquisition of landmarks, and MorphoJ for the Canonical Variate Analysis (CVA) and Discriminant Function Analysis (DFA). Landmarks established in Marchán et al. (2016) (tip of distal denticles, mid-point between the distal denticles, anterior end of the lateral ridges, lateral limits of the first clearly developed ring, points of maximum width between the lateral ridges -see Fig. 2) were chosen for the analysis, obtained from dorsal views of the distal part of genital chaetae. Genital chaetae were grouped by population and populations were subsequently grouped according to the main hypotheses recovered by the phylogenetic reconstruction, PCA and Bayesian clustering analysis.

### 7. Detection of selection and functional analyses

To detect selection signatures, the ‘reference-one SNP’ dataset was analyzed with PCAdapt, which jointly determines population structure and outlier loci (Duforet-Frebourg et al., 2014) and is, therefore, independent from any *a priori* assumption on population structure. Fsthet (Flanagan and Jones, 2017) was chosen as a second method relying on a different approach to outlier identification. Fsthet calculates smoothed quantiles from a SNP dataset, identifying SNPs as candidates for selection when their FST values relative to expected heterozigosity values fall outside such quantiles. In both cases, detected outlier SNPs were parsed to the corresponding contig using the *Carpetania elisae* transcriptome, and the surrounding regions (i.e., the coding sequence - CDS - region where each SNP lies) were analyzed to identify their most likely biological function as described below.

Untranslated regions (both 3’ and 5’-UTR) and CDS within transcripts (i.e., defined by opening reading frames, ORFs) were identified with Trinity and TransDecoder respectively (Haas et al., 2013) both for the reference transcriptome (*reference* hereafter) and the subset of contigs containing the SNPs with selection signatures (*under selection* hereafter). The location of SNPs under selection detected by PCAdapt and fsthet in both UTR and ORF regions was annotated. *Reference* and *under selection* datasets were annotated with eggNOG-mapper (Huerta-Cepas et al., 2017), which uses precomputed eggNOG-based orthology assignments for fast functional annotation.

A Gene Ontology (GO) enrichment analysis was performed with fatiGO (Al-Shahrour et al., 2007) to test if certain biological functions were more represented in one of the datasets. ‘Enrichment’ refers to genes (or their putative functions) that are over-represented in a list of genes (for instance, the list of genes where SNPs under selection lie) compared to the whole set of genes (i.e., the full transcriptome). Enrichment detection relies on a statistical comparison of the annotations of both sets of genes considering a p-value of 0.05. Two analyses were performed: (a) enrichment of *under selection* genes vs all the transcriptome -*reference*, and (b) enrichment of *under selection* genes vs non-selected genes from the transcriptome (to enhance detection). For a more clear visualization, reduction of redundancy of GO terms (for the *under selection* dataset) and visualization were performed using REVIGO (http://revigo.irb.hr/) (Supek et al., 2011). Default parameters were used, with an ‘allowed similarity’ threshold of 0.5. Genes annotated with GO terms associated with particularly relevant biological processes (see below) were parsed from the *reference* dataset. The peptide sequences were further annotated manually with BLASTp (https://blast.ncbi.nlm.nih.gov/) and Uniprot (https://www.uniprot.org/).

The concatenated sequences of SNPs with selection signatures were used as the input for a ML phylogenetic analysis (see section 3 above) in order to check if selection was limited to population-level local adaptation (i.e., a star-shaped phylogenetic tree would be recovered) or selected mutations were shared at higher phylogenetic levels (phylogenetic relationships between populations would reflect the species tree).

## Results

### 1. Genotyping-by-sequencing and SNP discovery

Illumina sequencing of GBS libraries for 95 individuals resulted in a total of 610,184,897 reads. Numbers of reads per individual ranged from 505,635 to 7,110,219, with a mean of 2,418,275. One individual (SEV5) was removed due to the low number of reads (2,948).

The different datasets contained the following number of SNPs: ‘de novo-all SNPs’ - 26,240, ‘de novo-one SNP’ −4,767, ‘reference-one SNP’ −3,181.

As genomic diversity and population genomics statistics were similar across the datasets, only those obtained from the ‘de novo-all SNPs’ are reported in Table 1. Most populations showed similar variability, with five of them (COL, PNT2, PR1, UCE and TAM) showing noticeably higher values for most of the parameters. Observed heterozigosity was higher than expected for all populations. Six populations (BAR, BOA, F1, LOY, PR1, TAM) showed negative FIS values, indicating individuals are less related than expected under a model of random rating, while the other eleven populations showed positive values, indicative of more closely related individuals than expected.

**Table 1.** Genomic diversity statistics obtained from the denovo_all dataset. Private: number of private alleles. Sites: number of variant nucleotide positions across the dataset. % Poly: percentage of polymorphic loci within the population. H_obs_: average observed heterozigosity. H_exp_ : average expected heterozigosity :average nucleotidic diversity. F_IS_: average inbreeding coefficient.

| Population | Sites | %Poly | Private | H<sub>obs</sub> | H<sub>exp</sub> | □ | F<sub>IS</sub> |
| --- | --- | --- | --- | --- | --- | --- | --- |
| BAR | 17224 | 0.74301 | 912 | 0.0192 | 0.0170 | 0.0189 | -0.0003 |
| BOA | 13621 | 0.51916 | 751 | 0.0131 | 0.0115 | 0.0128 | -0.0004 |
| CNG | 16036 | 0.9129 | 799 | 0.0199 | 0.0184 | 0.0204 | 0.0012 |
| COL | 18531 | 1.41761 | 1337 | 0.0355 | 0.0324 | 0.0360 | 0.0011 |
| F1 | 18849 | 0.41221 | 703 | 0.0103 | 0.0091 | 0.0101 | -0.0005 |
| HON | 17647 | 0.75366 | 794 | 0.0185 | 0.0166 | 0.0185 | 0.0003 |
| LOY | 19128 | 0.4398 | 1051 | 0.0137 | 0.0110 | 0.0122 | -0.0029 |
| PED | 16785 | 0.96378 | 722 | 0.0211 | 0.0191 | 0.0212 | 0.0005 |
| PNT2 | 18268 | 1.72951 | 1339 | 0.0412 | 0.0398 | 0.0443 | 0.0067 |
| POL | 13904 | 1.39305 | 727 | 0.0289 | 0.0272 | 0.0303 | 0.0029 |
| PR1 | 18003 | 1.32603 | 896 | 0.0330 | 0.0281 | 0.0312 | -0.0035 |
| SAL | 15772 | 0.98848 | 1362 | 0.0239 | 0.0218 | 0.0243 | 0.0014 |
| SEV | 13602 | 0.79526 | 722 | 0.0190 | 0.0175 | 0.0200 | 0.0021 |
| TAM | 15961 | 1.55836 | 1454 | 0.0571 | 0.0389 | 0.0432 | -0.0264 |
| TB | 18073 | 0.66944 | 943 | 0.0170 | 0.0157 | 0.0174 | 0.0011 |
| UCE | 18007 | 1.29482 | 1561 | 0.0297 | 0.0286 | 0.0318 | 0.0051 |
| **VRN** | 16362 | 0.99629 | 1509 | 0.0259 | 0.0236 | 0.0262 | 0.0005 |

Pairwise F_ST_ values (Suppl. Table 2) were high (mean= 0.64), with FST values between populations included in different clusters (see *Phylogenetic analysis*) (mean=0.67) being higher than FST values within clusters (mean=0.56).

### 2. Phylogenetic analysis

Phylogenetic relationships recovered from the datasets ‘de novo-all SNPs’ (Fig. 3), ‘de novo-one SNP’ and ‘reference-one SNP’ were mostly congruent. Main clades mimicked the deep lineages found in Marchán et al. (2017), separated in three main groups congruent with the population structure analysis (see below): A) lineage I B) lineages II and IV and C) lineages III, V and VI. Relationships within these groups were not clearly defined, as lineage II was recovered as paraphyletic with LOY branching either basally or closer to lineage IV populations, and lineages III and V were recovered in a clade with a very short branch separating them from lineage VI.

**Figure 3.**
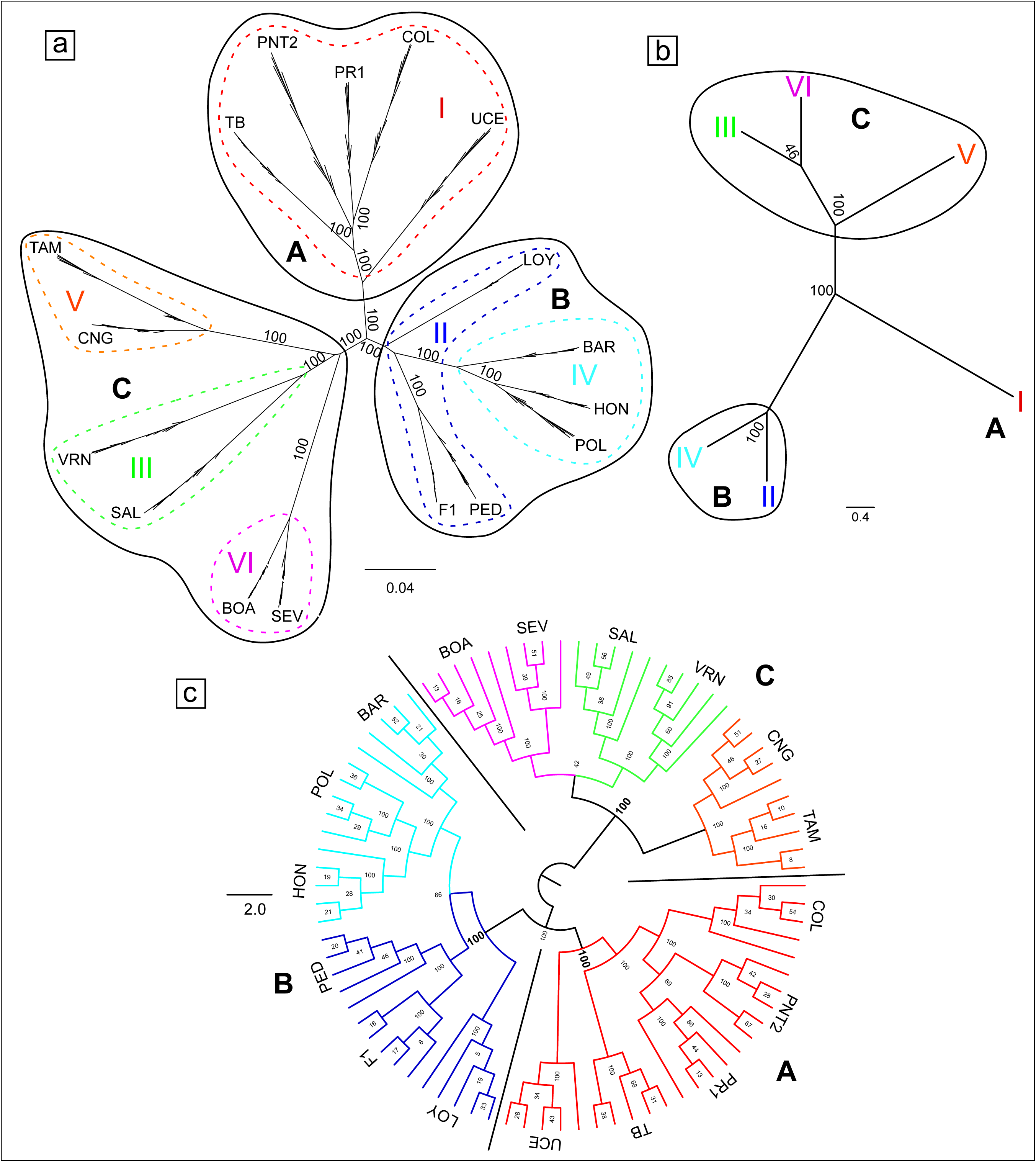
a) Maximum likelihood inference of the phylogenetic relationships of the studied populations of *Carpetania elisae*, based on ‘de novo-all SNPs’ dataset. Species trees obtained with SVDQuartets from the ‘de novo-all SNPs’ dataset b) grouping individuals by lineages from Marchán et al. (2017) and c) with ungrouped individuals. Lineages from Marchán et al. (2017) are shown with the same colors and roman numbers. The three main recovered clusters (A, B, and C) are indicated by a black outline.

The species trees obtained with SVDQuartets are shown in Figure 3b, 3c. Topologies obtained from the three SNP datasets were congruent. The total weight of compatible quartets for analyses grouping sequences by lineages from Marchán et al. (2017) was 87.65%, and 96.90% for analyses with ungrouped individuals. These high values indicate an almost complete absence of introgression and incomplete lineage sorting. In both cases, the three main clades shown by the maximum likelihood phylogenetic analyses were recovered with high bootstrap support.

### 3. Genetic structure

First five principal components obtained by the Principal Component Analysis (PCA) explained 13.6% / 11.6% / 9.1% / 8.0 % / 7.4% of variance. Representation of first two PCs (Fig. 4a) showed three main genetic clusters, which corresponded with the three main groups of populations recovered in the phylogenetic trees (A, B and C). Populations within lineage I clustered tightly with UCE, being the most divergent within that cluster. Populations from lineages II and IV showed no clear separation within cluster B, while in cluster C lineage V diverged the most from the rest of the populations.

**Figure 4.**
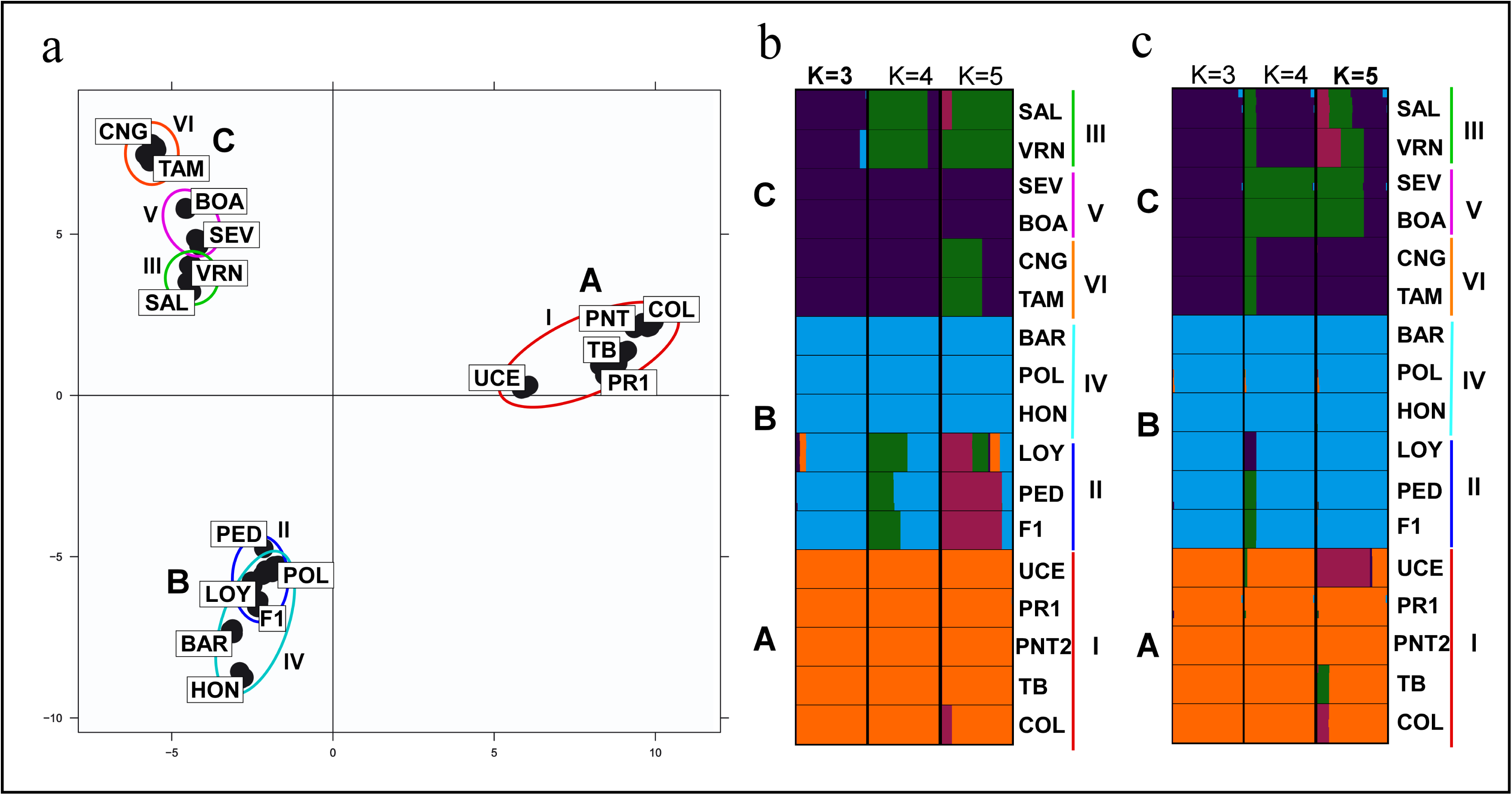
a) Principal Component Analysis (PCA) of ‘de novo-one SNP’ dataset. First two Principal Components are shown. Barplot of STRUCTURE analyses of b) ‘de novo-one SNP’ dataset and c) ‘reference-one SNP’ dataset, major modes for each k are summarized. Each color shows percentage of assignment to a cluster or ancestral population. b) Optimal K=3; c) Optimal K=5; K=3, 4 and 5 are shown for comparison. Lineages from Marchán et al. (2017) are shown with the same colors and roman numbers.

Statistically significant correlation was found between percentage of missingness and main principal components (r= −0.5158, p= 0.0000; r= 0,3875, p= 0.0003).

STRUCTURE analysis estimated the optimal K for ‘de novo-one SNP’ dataset to be 3, while the optimal K for ‘reference-one SNP’ dataset was 5. Genetic clusters at K=3 included the same individuals and populations for both datasets (Fig. 4b), which matched with the previously identified groups (A: lineage I, B: lineages II + IV and C: lineages III, V and VI). K=5 identified further subdivision within these clusters, which differed between the datasets. In the case of ‘de novo-one SNP’, the individuals corresponding to lineage II formed a separate cluster from lineage IV individuals, while they were all recovered in a homogeneous cluster in the ‘reference-one SNP’ dataset analysis. In ‘de novo-one SNP’ analysis individuals from lineage III and VI were clearly separated in two clusters, while individuals from lineage V showed a strong admixture of both; in ‘reference-one SNP’ analysis lineage V and VI were assigned to different clusters while individuals from lineage III showed admixture between the former clusters and a third one corresponding to the ancestral population dominating UCE individuals (belonging to lineage I). The distinctness of UCE individuals was not recovered in the ‘de novo-one SNP’ analysis, in which lineage I individuals formed a very homogeneous cluster.

### 4. Geometric morphometric analyses of genital chaetae

Canonical variate analysis of the shape of the genital chaetae showed differences when the input groupings represented the K=3 STRUCTURE clusters and the cryptic lineages from Marchán et al. (2017) (Fig. 5). For K=3, genital chaetae from specimens assigned to the cluster A (Lineage I) were clearly separated from chaetae from clusters B (lineages II and IV) and C (lineages III, V and VI), though the latter showed a moderate overlap in their confidence ellipses. The analysis based on the grouping by lineages showed the same separation of lineage I, and also a clear separation of lineage II, with lineage IV overlapping with the rest of the lineages.

**Figure 5.**
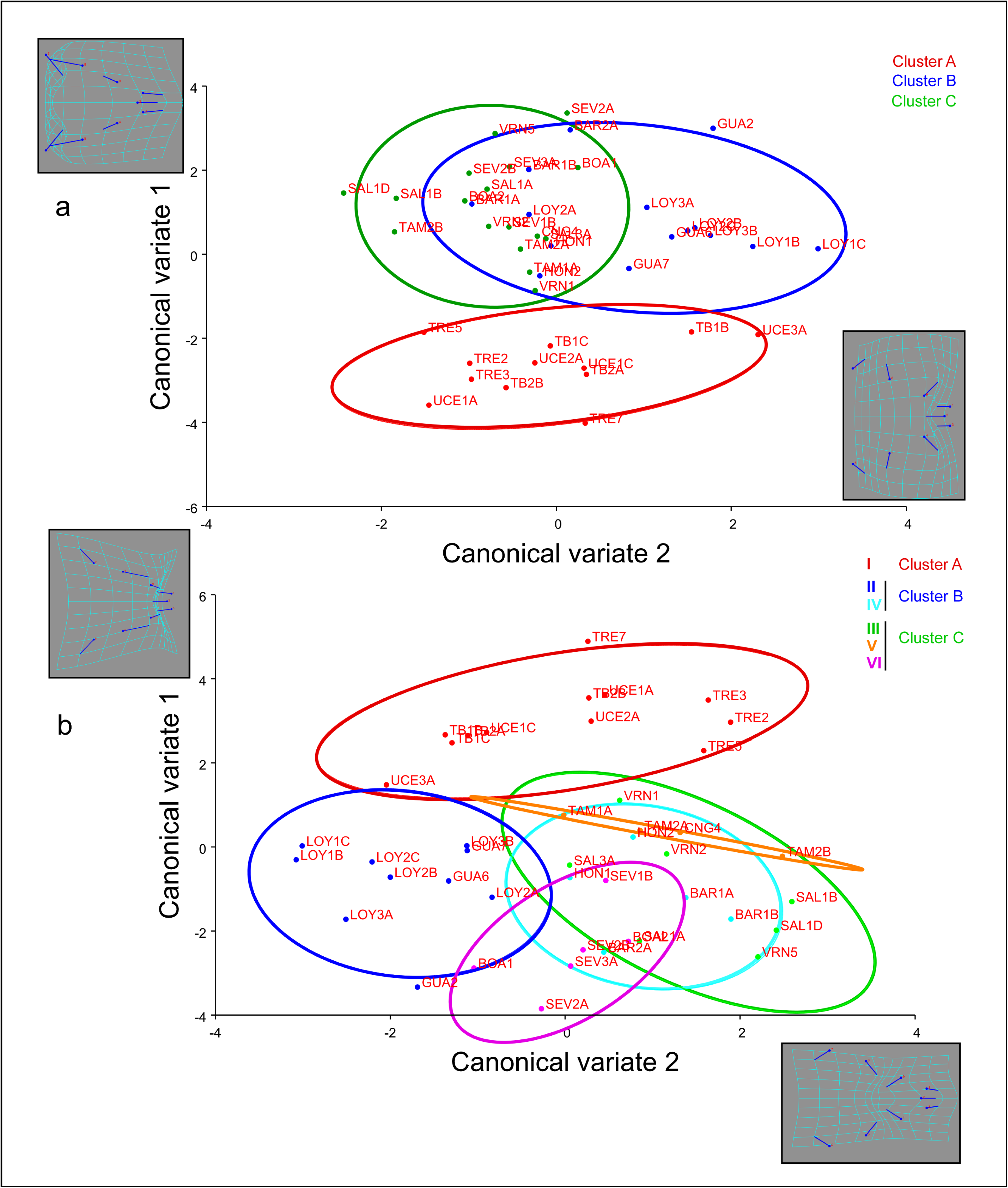
Geometric morphometrics analysis of distal tip of genital chaetae of *Carpetania* populations. A) Canonical variate analysis grouping observations by STRUCTURE K=3 clusters. B) Canonical variate analysis grouping observations by cryptic lineages defined in Marchán et al. (2017). Equal frequency ellipses (probability = 0.9) are displayed for observations belonging to the different groups. Deformation grids display morphological landmarks and shape change represented by each axis.

Results of the Discriminant Function Analysis using the K=3 grouping are shown in Suppl. table 3. Assignments and cross-validation tests were highly accurate except for Cluster B-Cluster C comparisons, were 40% and 29,5% of chaetae were missasigned in cross-validation. Nonetheless, all pairwise comparisons were statistically significant (p-value <0.05 for 1,000 permutation runs).

### 5. Putative loci under selection

PCAdapt found 867 outlier SNPs. In the *reference* dataset, 51,720 Opening Reading Frames (ORFs), a proxy for peptides, were identified, and 20,817 ORFs were successfully annotated with eggNOG-mapper (40.25% of the reference transcriptome). In the *under selection* dataset 1,406 ORFs were identified, and 557 ORFs were annotated with 4,871 GOs (39.615% of the contigs at protein level). These GO terms are summarized in a treemap graph obtained in REVIGO (Fig. 6a, Suppl. Fig. 1). The whole annotated dataset is shown in Suppl. Table 4. Some examples of putative proteins with local adaptation signatures involve functions related to metabolism, reproduction, reception of stimuli, development, among others, and their inferred biological function are shown in detail in Suppl. Table 5. Several protein-coding genes with selection signatures showed shared alleles within each of the three main clusters. Putative proteins and their most likely biological function are shown in Table 2.

**Figure 6.**
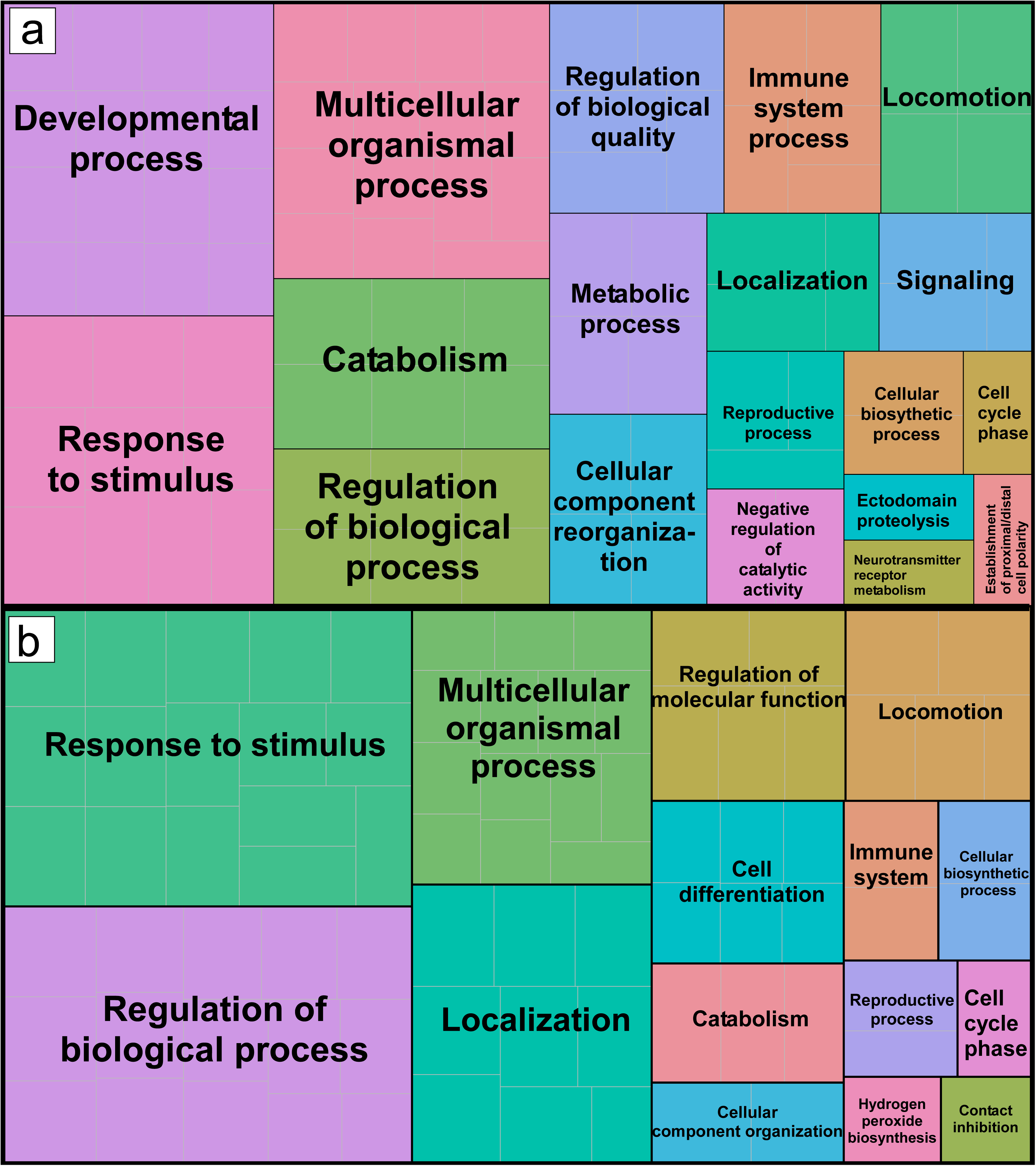
Treemap graph summarizing the GO terms (biological process level) of the genes containing SNPs under selection. High rank GO terms are shown (see also Suppl. Fig. 1 for full terms). The size of each main square is related to the frequency and the hierarchical relationship between GOs.

**Table 2.** Putative proteins with shared alleles (corresponding to SNPs with selection signatures detected by PCAdapt and fsthet -indicated by an asterisk) between populations assigned to clusters A-B-C

|  |  |  |
| --- | --- | --- |
| Cluster A | ***Nuclear hormone receptor HR96*** | Controls tryglycerid and cholesterol homeostasis, affecting response to starvation |
|  | ***Rabenosyn-5*** | Role in endosomal and lysosomal transport. Involved in the blood coagulation pathway |
|  | ***NADP-dependent oxidoreductase domain-containing protein 1*** | Probable oxidoreductase |
|  | ***JNK-interacting protein 1*** | Involved in axonal transport of mitochondrion and synaptic vesicle transport |
|  | ***ATP-dependent zinc metalloprotease YME1L1*** | Ensures cell proliferation, maintains normal cristae morphology and complex I respiration activity, promotes antiapoptotic activity and protects mitochondria from oxidatively damaged membrane proteins |
|  | ***Cleavage stimulation factor subunit 2*** | mRNA processing |
|  | ***Histone-lysine N-methyltransferase 2A*** | Essential role in early development, hematopoiesis and control of circadian gene expression |
|  | ***Nucleoporin p54*** | Component of the nuclear pore complex, |
|  | ***Collagen alpha-1(II) chain*** | In vertebrates, specific for cartilaginous tissues |
|  | ***Ral GTPase-activating protein subunit beta*** | GTPase activator |
|  | ***Protein crumbs homolog 2*** | Apical polarity protein that plays a central role during the epithelial-to-mesenchymal transition at gastrulation |
|  | ***snRNA-activating protein complex subunit 4*** | Required for the transcription of both RNA polymerase II and III small-nuclear RNA genes |
|  | ***Pericentrin*** | Integral component of the filamentous matrix of the centrosome involved in the initial establishment of organized microtubule arrays in both mitosis and meiosis |
|  | ***Collagen-like protein\**** |  |
|  | ***Histone-lysine N-methyltransferase, H3 lysine-79 specific\**** | Histone methyltransferase. Required for Polycomb Group and trithorax Group maintenance of expression in *Drosophila*. Also involved in telomeric silencing |
| Cluster B | ***JNK-interacting protein 1*** | May function as a regulator of synaptic vesicle transport. In *C. elegans* it has been linked to locomotion, ovoposition and defecation |
|  | ***Vinexin*** | Involved in smooth muscle contraction |
|  | ***Ankyrin repeat domain-containing protein 26\**** | Acts as a regulator of adipogenesis. Involved in the regulation of the feeding behavior. |
| Cluster C | ***RNA helicase aquarius*** | Involved in pre-mRNA splicing as component of the spliceosome |
|  | ***Probable sodium/potassium/calcium exchanger CG1090*** | Possible function in the removal and maintenance of calcium homeostasis |
|  | ***Cytochrome P450 2U1*** | Catalyzes the hydroxylation of long chain fatty acids |
|  | ***Zinc transporter 2*** | Zinc ion transmembrane transport |
|  | ***Host cell factor 1*** | Involved in control of the cell cycle |
|  | ***Protein eva-1-like\**** | Acts as a receptor for slt-1 . Required for the guidance of the AVM pioneer axon to the ventral nerve cord. |
|  | ***Rho GTPase-activating protein 32\**** | GTPase-activating protein promoting GTP hydrolysis. May be involved in the differentiation of neuronal cells during the formation of neurite extensions. |

fsthet detected 269 outlier SNPs, for which the F_ST_/heterozigosity ratio was significantly higher or lower than expected (Fig. 7). Fourteen outlier loci matched with the ones found by PCAdapt, and 73 were located in the same contigs as the PCAdapt SNPs with selection signatures. 489 ORFs were identified, and 129 ORFs were annotated. GO terms are summarized in a treemap graph obtained in REVIGO (Fig. 6b). A few protein-coding genes with selection signatures showed shared alleles within each of the three main clusters. Putative proteins and their most likely biological function are shown in Table 2.

**Figure 7.**
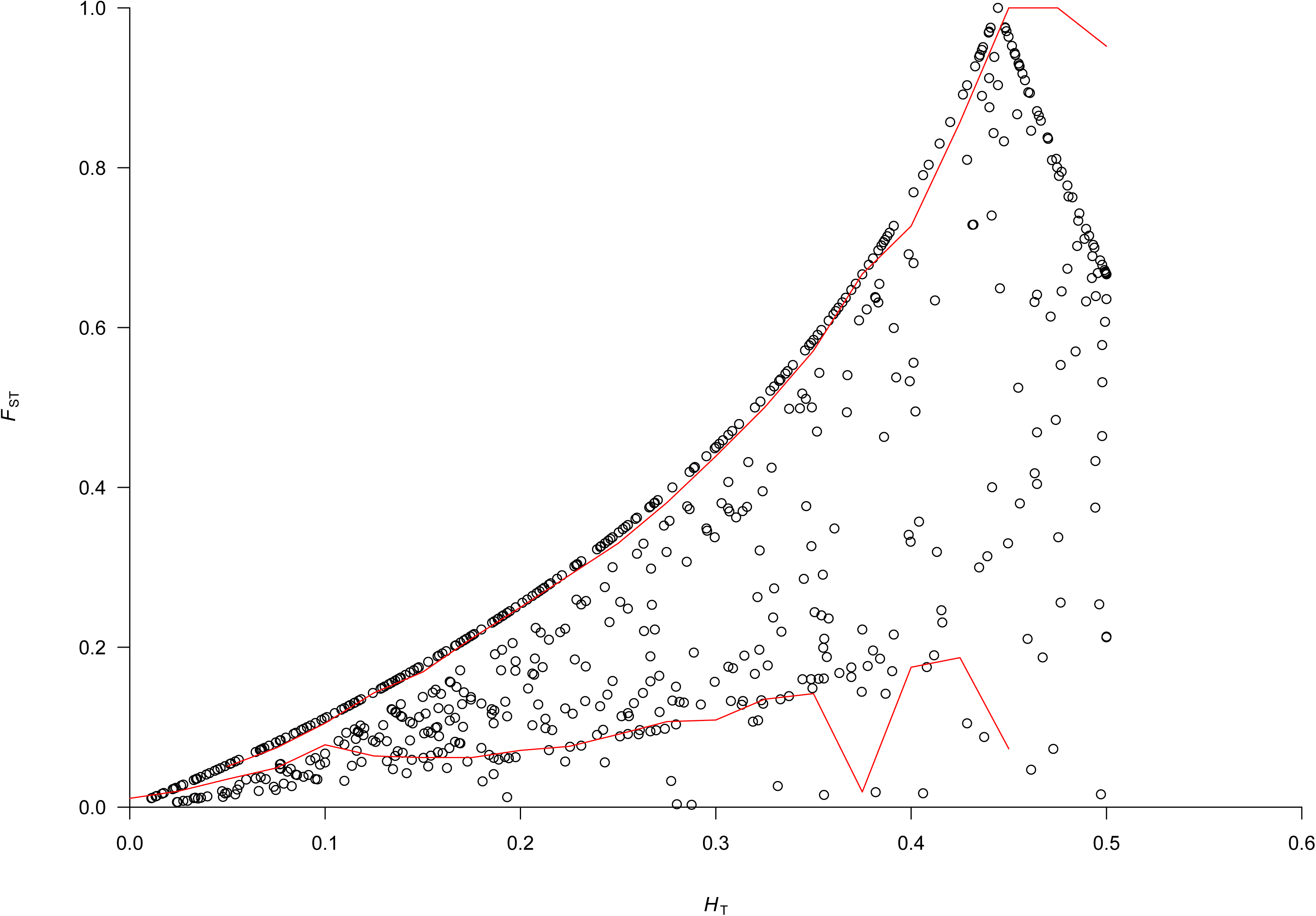
FST/heterozigosity plot obtained with fsthet, showing the smoothed quantiles as a red line. Outlier loci are the ones outside of the smoothed quantiles.

Maximum likelihood inference based on the concatenated sequence of the 867 outlier SNPs recovered well-resolved phylogenetic relationships between the populations (Suppl. Fig. 2), lineages and higher level groups, suggesting selection signature was pervasive through the different taxonomic levels.

GO enrichment analysis showed that genes with SNPs under selection (detected by PCAdapt) were statistically enriched at the biological function level (p-value < 0.05) in regulation of transcription regulation, as well as signal transduction, chromosome organization, development and mitotic anaphase, among others (Fig. 8a, Suppl. Fig. 3), as well as RNA binding, nucleoside triphosphatase activity and chromatin binding at the molecular function level (Fig. 8b, Suppl. Fig. 4) and complexes, cell, and cell part at the cellular component level (Fig. 8c, Suppl. Fig 5). Remarkably, several SNPs under selection were found in UTR regions placed in 83 different contigs (Suppl. Table. 6). Genes including outlier SNPs identified by fsthet showed enrichment for no biological function, molecular function or cellular component.

**Figure 8.**
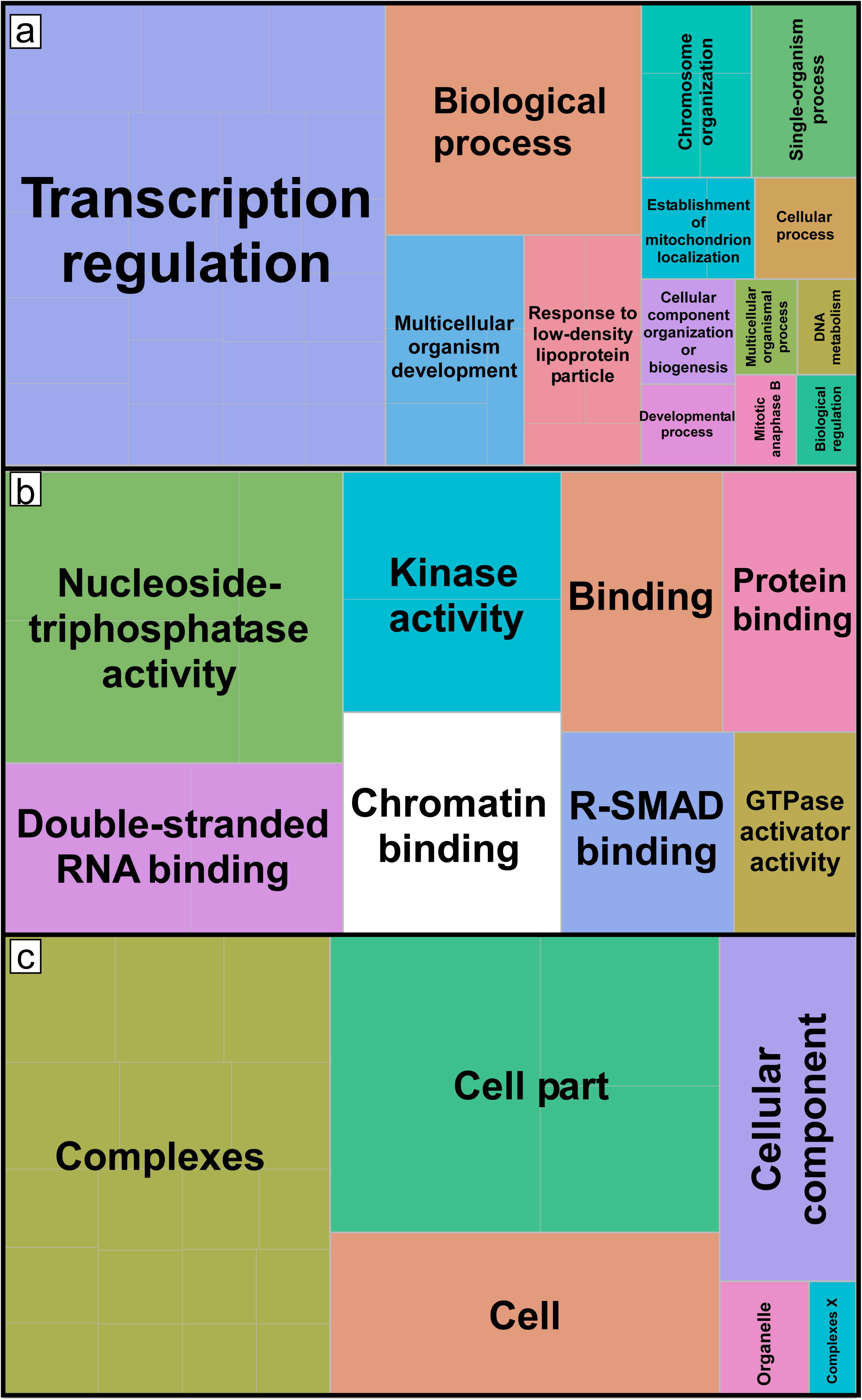
Treemaps showing GO enrichment functions enriched in the genes where SNPs under selection are placed. High rank GO terms are shown. (a) Biological process level. (b) Molecular function level. c) Cellular component. The size of each main square is related to the p-value of the enrichment analysis (see also Suppl. Figures. 2, 3 and 4).

## Discussion

### 1. Integrative taxonomy identified three putative cryptic species within the Carpetania species complex

The different analyses applied to the genome-wide genetic information provided by GBS evidenced two clear patterns. First, the uppermost level of genetic structure within the studied populations of *Carpetania* identified three main and distinctive clusters, each of them including one or several of the six cryptic lineages previously identified in(Marchán et al., 2017). Second, one of those clusters (termed C through this work) showed a higher level of internal genetic divergence, with presence of clear substructure in three lineages but hints of admixture between them, that is, presence of DNA from a distantly-related population or species as a result of interbreeding between genetically differentiated populations.

The information provided by the geometric morphometrics analyses of the genital chaetae of *Carpetania* supported to a significant extent the separation of the genetic clusters, albeit with a high degree of overlap as revealed by the canonical variate analysis. The shape of the genital chaetae of clusters B and C showed overlap according to the canonical variate analysis, explained by chaetae from lineage IV - comprised in cluster B as revealed by the phylogenetic analyses (Fig. 5, b) - being similar to chaetae from cluster C. Convergent evolution in the shape of chaetae of lineage IV and cluster C could explain the observed pattern. Morphological differentiation of these clusters, even if not absolutely clear-cut, suggest they should be considered pseudocryptic (as in above the resolution of morphological analysis) instead of cryptic taxonomical entities.

Several studies have assimilated this kind of above-population level clusters inferred from genome-wide diversity to species level (Brunet et al., 2017; Pinto et al., 2019). This has been done even in the absence of morphological evidence (Warner et al., 2015; Garg et al., 2016; Dincă et al., 2019), but has also been confirmed by geometric morphometrics and other cryptic characters (Alter et al., 2017). One criterium proposed for robust species delimitation in cryptic taxa is genetic distinctiveness in sympatry (Mallet, 1995). However, Marchán et al. (2017) found no sympatry of the different lineages in small scale transects separated just by a few hundred meters. On the other hand, genetic cohesiveness across allopatric populations (Good and Wake, 1992; Mallet, 1995) is clearly fulfilled: BAR and HON populations, for example, are separated 100 km but show very little differentiation.

Following these results, the most robust systematic proposal for the cryptic (or pseudo-cryptic) species within the genus *Carpetania* would therefore consist of three species. One of them (cluster C) may contain enough genetic diversity to consider the future possibility of assigning subspecific status to its genetic lineages in order to recognize their distinctness and improve biodiversity conservation efforts. Further research within this clade will help elucidate this taxonomic necessity. Formal description of the putative three cryptic species can be found in Marchán et al. (2019).

### 2. Local adaptation fuels cryptic speciation in the Carpetania species complex

The finding of selection signatures in genes across the genome provided a highly valuable insight into the evolutionary processes governing the *Carpetania* cryptic species complex. Several of the loci under selection respond to a pattern of local adaptation, as selected mutations appeared in single populations. This suggests that isolated populations of this cryptic complex evolved independently as a response to local environmental conditions, which could have fueled cryptic speciation in the long term. Little is known about the genomic basis of local adaptation in cryptic lineages in nonmodel organisms. Boissin et al. (2011) found compelling correlation between adaptation to local conditions (in their study correlated to oligotrophy), evolution of reproductive traits and cryptic speciation on the ophiuroid *Ophioderma longicauda.* Interestingly, locally-adapted lineages showed reduced dispersal ability when compared to other lineages. This is consistent with postulates from Bickford et al. (2007), that suggested that directional selection on traits with no apparent morphological correlates could drive cryptic diversification.

Moreover, the distinctiveness of the shared adaptive mutations across cryptic (or pseudocryptic) species confirmed their relevance as biological entities: the absence or scarcity of morphological differences does not reflect their genetic, physiological or metabolic diversity. As mentioned above, these cryptic features have been identified by different approaches in independent cryptic complexes (reviewed in Marchán et al., 2018a), but have rarely been studied by genome-wide analyses (Anderson et al., 2017; Shekhovtsov et al., 2019). We emphasize that integrative-centered studies guided by state-of-the-art methodologies are therefore most needed to further understand genome evolution and adaptation in cryptic non-model organisms.

### 3. Regulation of gene expression may drive cryptic speciation in the Carpetania species complex

Once considered as useless or junk mRNA, it is now well-known that UTR regions are involved in many aspects of regulation of gene expression. The 3 ′ UTR region has been shown to play a key role in translation termination as well as post-transcriptional gene expression (Matoulkova et al., 2012; Young and Wek, 2016; Leppek et al. 2017; Ren et al. 2017; Mayr 2018). Although a few case reports have shown that mutations and variants in these regions can have important genomic consequences, such as disease, genetic sequencing approaches typically focus on protein-coding regions and ignore these variants, particularly in studies dealing with non-model organisms, where this is a virtually unexplored field. In this work, a remarkable proportion (9.6%) of the SNPs under selection were located in UTR regions. It has been shown that a few nucleotide substitutions in UTR regions can significantly alter protein expression, measured as protein abundance (Dvir et al., 2013); that study, demonstrated the powerful consequences of sequence manipulations of even 1-10 nucleotides immediately upstream of the start codon, which resulted in significantly altered abundance of expressed proteins in yeast.

Likewise, GO enrichment analysis identified that genes were SNPs under selection concentrate (i.e., the *under selection* dataset as described above) are enriched in transcription regulation activity compared to all other genes in the transcriptome (as shown in Fig. 7a). The hypothesis that differences in gene regulation play an important role in speciation and adaptation is not new (as reviewed in Romero et al., 2012). Changes in gene regulation (i.e., regulatory divergence) have been shown to play a major role both in intrinsic pre- and/or post-zygotic isolation and in establishing other reproductive barriers as a byproduct of adaptive divergence, as in the case of ecological speciation (Pavey et al., 2010; Xu et al., 2016 Mack and Nachman, 2017; Deng et al., 2018). Gene expression might promote ecological speciation in two ways: indirectly by promoting population persistence (as suggested by studies of plasticity in morphological and behavioural traits related to fitness and population persistence, and studies of gene expression responses during ecological shifts, particularly those resulting in exposure to ecological stress), or more directly by affecting adaptive genetic divergence in traits causing reproductive isolation (Pavey et al., 2010). Indeed, as described below, we found genes with SNPs under selection related to reproduction, particularly hormonal pathways. Altogether, our results provide preliminary evidence showing that local adaptation may be reshaping regulation in gene expression in the *Carpetania* species complex, and it opens the door to further empirical testing of the hypothesis that regulatory divergence is indeed a major driver of cryptic speciation in soil fauna.

### 4. Loci under selection in the different cryptic species are putatively related to a plethora of biological functions related to reproduction and interactions with the environment

Even though selection signatures were detected in genes with putative biological functions related to a diversity of biological processes, some of those were especially suggestive from the point of view of the divergence and radiation of the cryptic complex. Several of the proteins studied in detail (featured in table 1) are expected to be involved in the interaction of *Carpetania* with their environment, with those related to metabolism, immune system and response to environmental stress being the most relevant, such as *14-3-3 protein zeta-like* or *Serine/threonine-protein kinase* TBK1. Local adaptation in such proteins could lead to the evolution of differential ecological preferences as seen in other cryptic species complexes, in diatoms, wildflowers and earthworms (Vanelslander et al., 2009; Yost et al., 2012; Spurgeon et al., 2016).

Considering the particularities of soil as a habitat and the biology of earthworms, proteins related to hypoxia (as *Manganese superoxide dismutase*, *Homeodomain-interacting protein kinase 2* or *Hypoxia-inducible factor 1-alpha isoform X1*) appear as a potentially-relevant target for adaptation. Soil flooding or soil compaction can result in deficient soil aeration and reduced available oxygen, representing a limiting factor for soil fauna. An increase in the ability to cope with these environmental pressures could lead to an increase in population persistence or the colonization of new niches. MAP3K12, with its role in UV-induced DNA damage and osmolarity changes regulation, could also be a relevant protein in local adaptation to environment by providing enhanced resistance to the harmful effect of exposure to solar radiation and changes in water availability.

It should be noted that some genes with putative sensory or behaviour-related functions showed signals of local adaptation as well. For example, annetocin receptor was among these genes: the closely related arginine vasopressin (AVP) receptors in the brain of rodents have an important effect on sexual behaviour and mate choice (Horth, 2007). Other examples are the *Coiled-coil and C2 domain-containing protein 1-like protein* and *Sal-like protein 1*, involved in sensory organ development (Klein, 2003; Celis et al., 2009) or *Putative transcription factor capicua*, involved in central nervous system development (Lu et al., 2017). These mutations under selection could affect intraspecific mate recognition, promoting the development of pre-zygotic reproductive barriers and isolation between the evolutionary diverging cryptic species. A precopulatory sexual selection behaviour has been previously observed for cryptic earthworm species (Jones et al., 2016).

Among the reproduction-related genes found to possess selection signatures, the annetocin receptor (Kawada et al., 2004) is noteworthy. Annetocin, a neuropeptide related to oxytocin, has been found to elicit egg-laying behaviour in *Eisenia fetida* (Oumi et al., 1996). Beyond this well-known function, this signaling pathway could be involved in other reproductive behaviours, as is the case with oxytocin - male copulative behaviour in snail *Lymnaea stagnalis*, coordination of reproductive behaviour in roundworm *C. elegans* (Gruber, 2014) - thus having a potential effect on reproductive isolation and differential reproductive success. Although putative, these functions may provide hints about how local adaptation might be reshaping the genome of the *Carpetania* cryptic species complex. More conclusive functional experiments will help to validate these findings.

## 5. Conclusions

In this piece of work, we show the potential to characterize and delimit robust species within cryptic complexes in soil milieu through the study of genome-wide genetic variability in the terrestrial annelid *Carpetania*, together with the exploration of inconspicuous morphological variability. The pervasive presence of local adaptation signatures in functionally diverse loci across the genome of these pseudocryptic species does not only confirm their biological relevance as distinct entities beyond their apparent homogeneity, but it also sheds light into the underlying genetic basis for cryptic speciation. These selection signatures also confirms the potentially important role of regulation of gene expression in adaptation and speciation within species complexes. Finally, the detection of putative genes subjected to local adaptation allowed identifying biologically relevant proteins with a potential role in the interaction of soil fauna with their environment, as well as proteins that could be involved in the reproductive isolation of cryptic species. Altogether, our results indicate that local adaptation and regulatory divergence provided an arena of genetic novelty to reshape the genome of three cryptic species of terrestrial annelids, possibly fueling ecological speciation. Future genomic studies will help elucidate with more precision the genetic underpinnings of these evolutionary processes. We emphasize that integrative taxonomic-centered studies are therefore most needed to further understand genome evolution in nonmodel soil organisms.

## Acknowledgments

This work was supported by Universidad Complutense de Madrid and Santander Group -Proyecto de Investigación Santander/Complutense PR41/17-21027, Systematics Research Fund (SRF) and Xunta de Galicia. Consellería de Cultura, Educación e Ordenación Universitaria. Secretaria Xeral de Universidades under grant ED431B 2019/038.

RF was funded by a Marie Skłodowska-Curie Fellowship (747607). DF was funded by a Juan de La Cierva-Formación grant (FJCI-2017-32895) from the Spanish Ministry of Sciences, Innovation and Universities. MN was funded by a Postdoctoral Fellowship UCM.

## Data Accessibility Statement

Scanning electron micrographs, landmark acquisition, GBS raw reads, SNP datasets and transcriptome assembly are available at Harvard Dataverse: FERNANDEZ MARCHAN, DANIEL, 2019, “Local adaptation and cryptic speciation in terrestrial annelids: GBS and geometric morphometrics data”, https://doi.org/10.7910/DVN/RVMQND, Harvard Dataverse, V1

## Author Contributions

DFM, MN, DJDC and RF designed the research. DFM and NS obtained and analyzed the morphological data. DFM, MN and RF analyzed the molecular data. DFM and RF wrote the first version of the manuscript, and MN, JD and DJDC made significant contributions to the final version of the manuscript.

## Supplementary Materials

**Suppl. Table 1.** Populations of *Carpetania* included in this work, geographical coordinates of the collection localities, sampling season and year of collection.

**Suppl. Table 2.** Pairwise FST values between the studied populations of *Carpetania* obtained from the ‘de novo-all SNPs’.

**Supp. Table 3.** Classification/misclassification tables from the Discriminant Function Analysis and cross-validation analysis with chaetae grouped by K=3 STRUCTURE clusters. ClA, ClB, ClC: Clusters A, B and C.

**Suppl. Table 4.** *Under selection* dataset (contigs containing SNPs with selection signatures) annotated Open Reading Frames (ORFs).

**Suppl. Table 5.** Most relevant proteins under selection with relevant putative functions in the biology of the Carpetania cryptic complex.

**Suppl. Table 6.** SNPs with selection signatures located in UTR regions.

**Suppl. Figure 1.** Treemap graph summarizing the GO terms (biological process level) of the genes containing SNPs under selection. a) PCAdapt, b) fsthet. The size of each main square is related to the frequency and the hierarchical relationship between GOs.

**Suppl. Figure 2.** Maximum likelihood inference based on the concatenated sequence of the 867 outlier SNPs detected by PCAdapt. Lineages from Marchán et al. (2017) are shown with the same colors and roman numbers. The three main recovered clusters (A, B, and C) are indicated by a black outline.

**Suppl. Figure 3.** Treemap showing GO enrichment functions enriched in the genes where SNPs under selection are placed. Biological process level. The size of each main square is related to the p-value of the enrichment analysis

**Suppl. Figure 4.** Treemap showing GO enrichment functions enriched in the genes where SNPs under selection are placed. Molecular function level. The size of each main square is related to the p-value of the enrichment analysis

**Suppl. Figure 5.** Treemap showing GO enrichment functions enriched in the genes where SNPs under selection are placed. Cellular component level. The size of each main square is related to the p-value of the enrichment analysis

## Notes

https://doi.org/10.7910/DVN/RVMQND

