## Supplementary figures and images for "Local adaptation fuels cryptic speciation in terrestrial annelids"

### Suppl. Figures

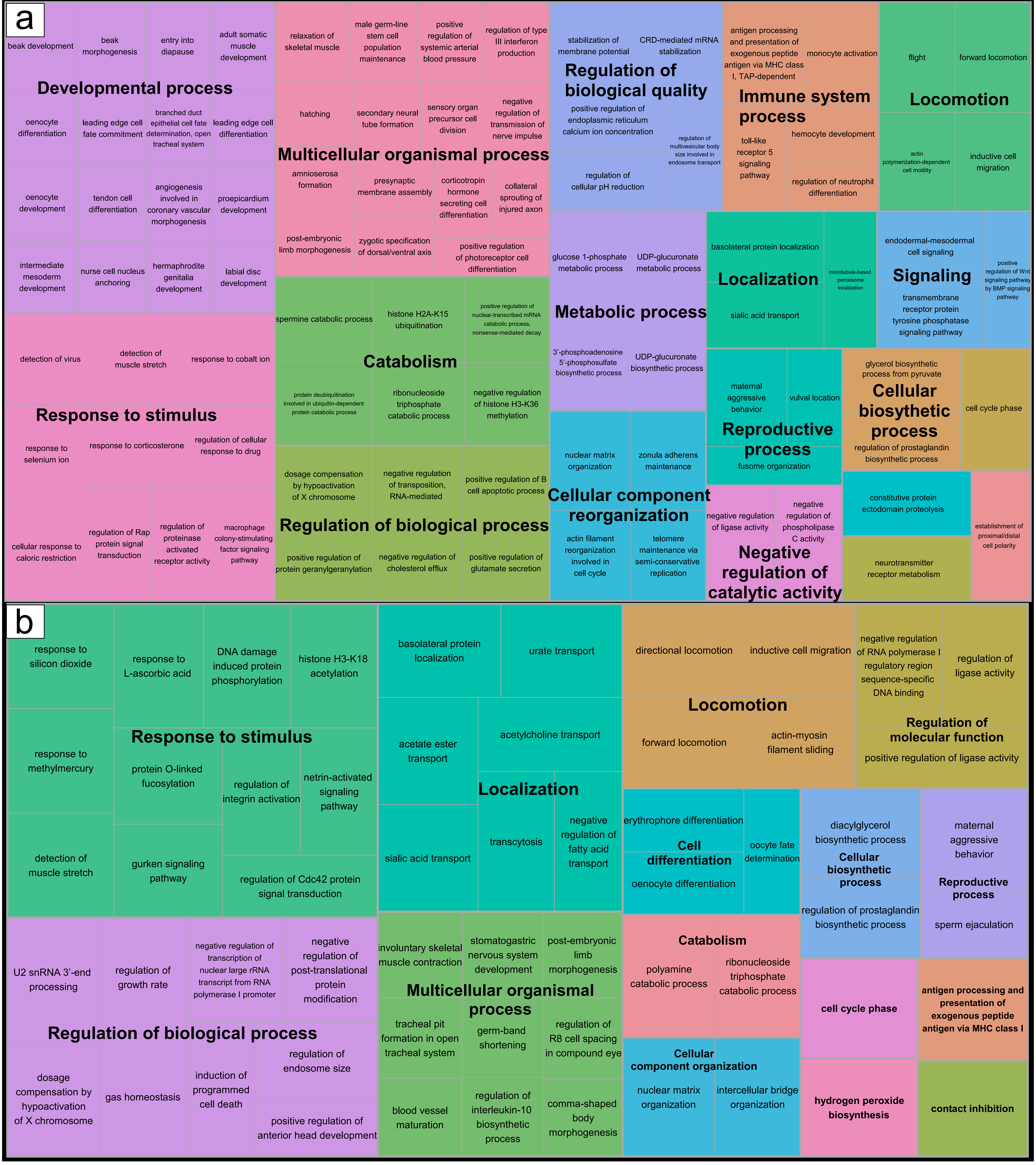

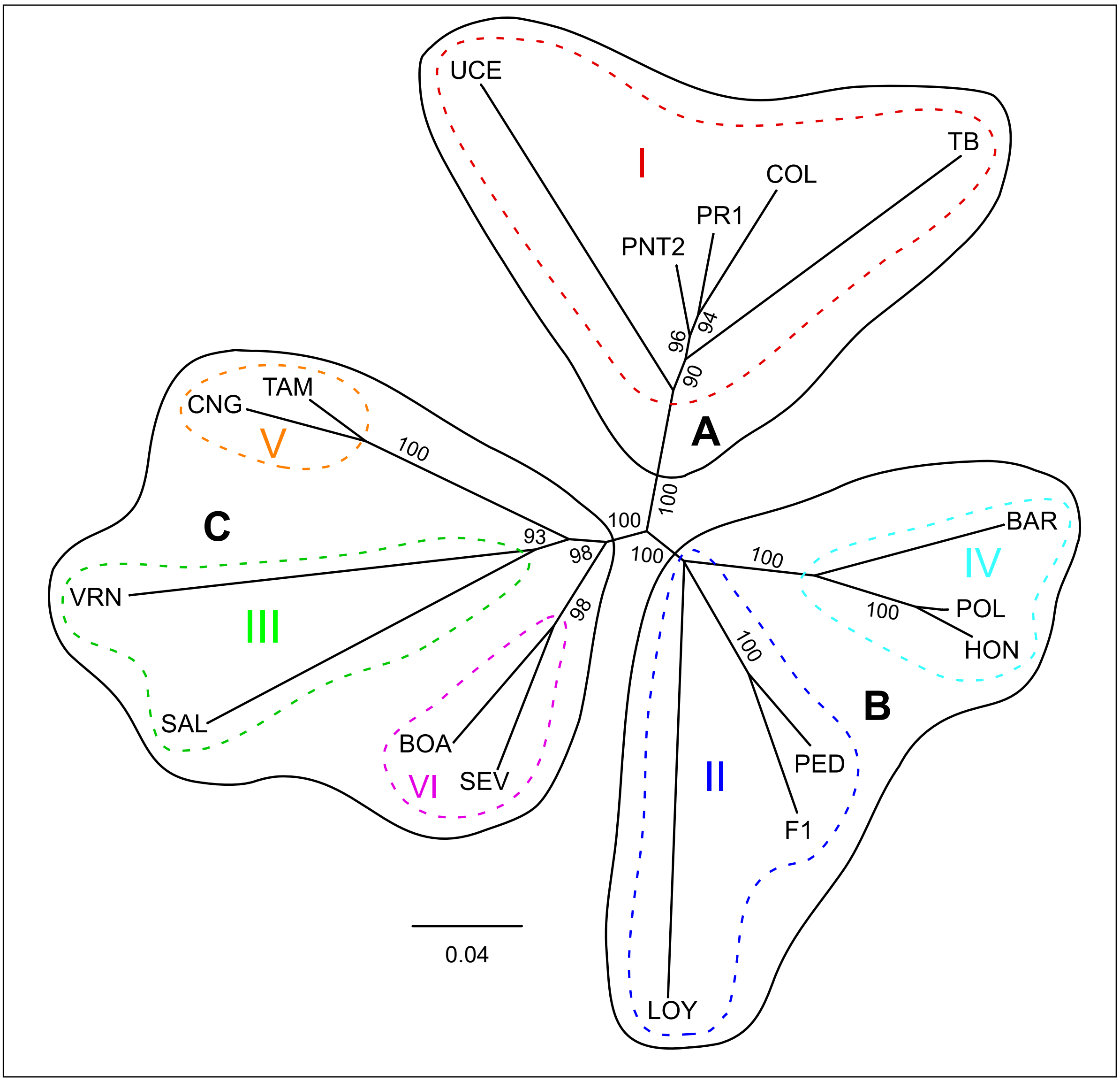

REVIGO Gene Ontology treemap

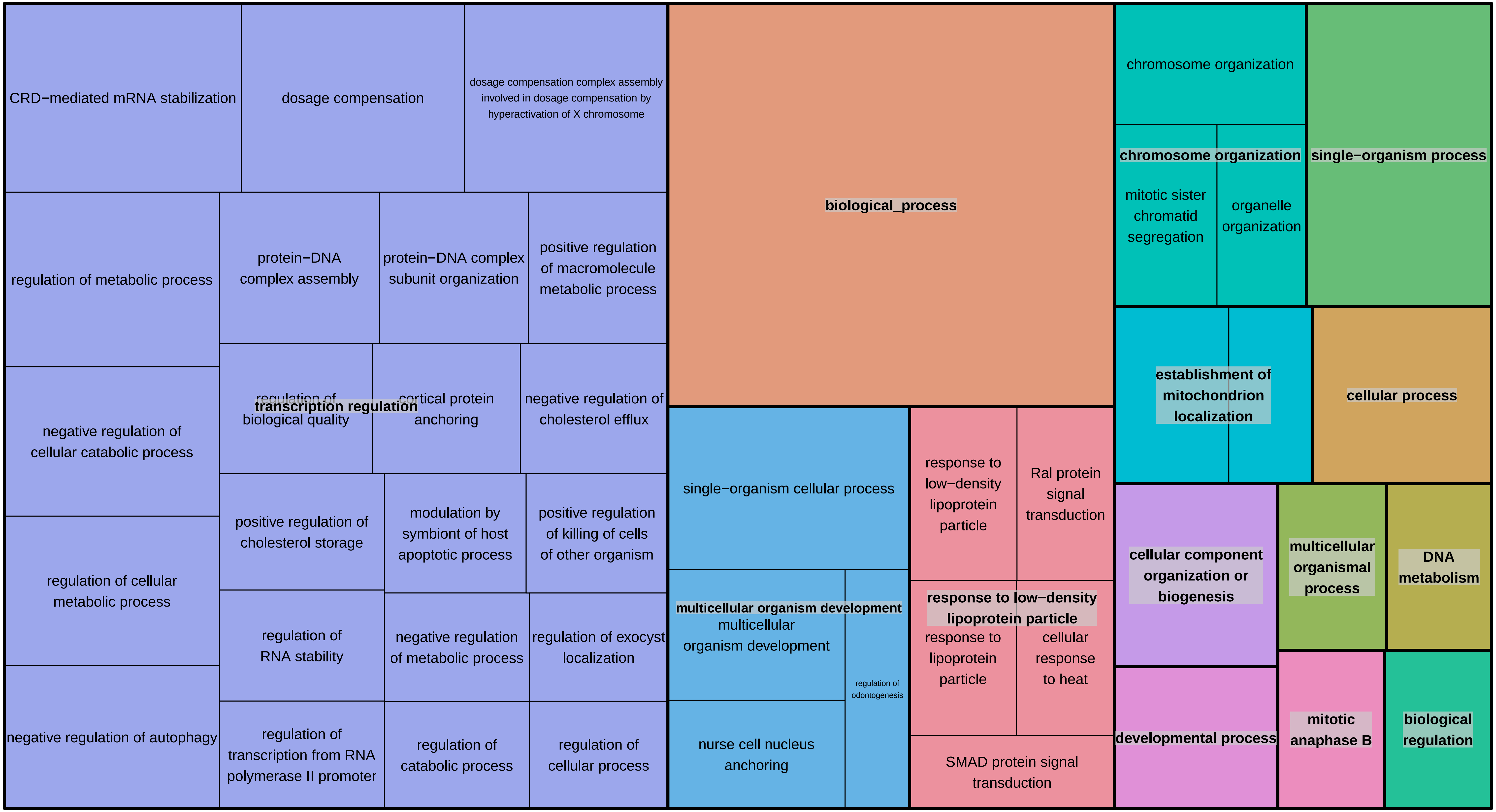

REVIGO Gene Ontology treemap

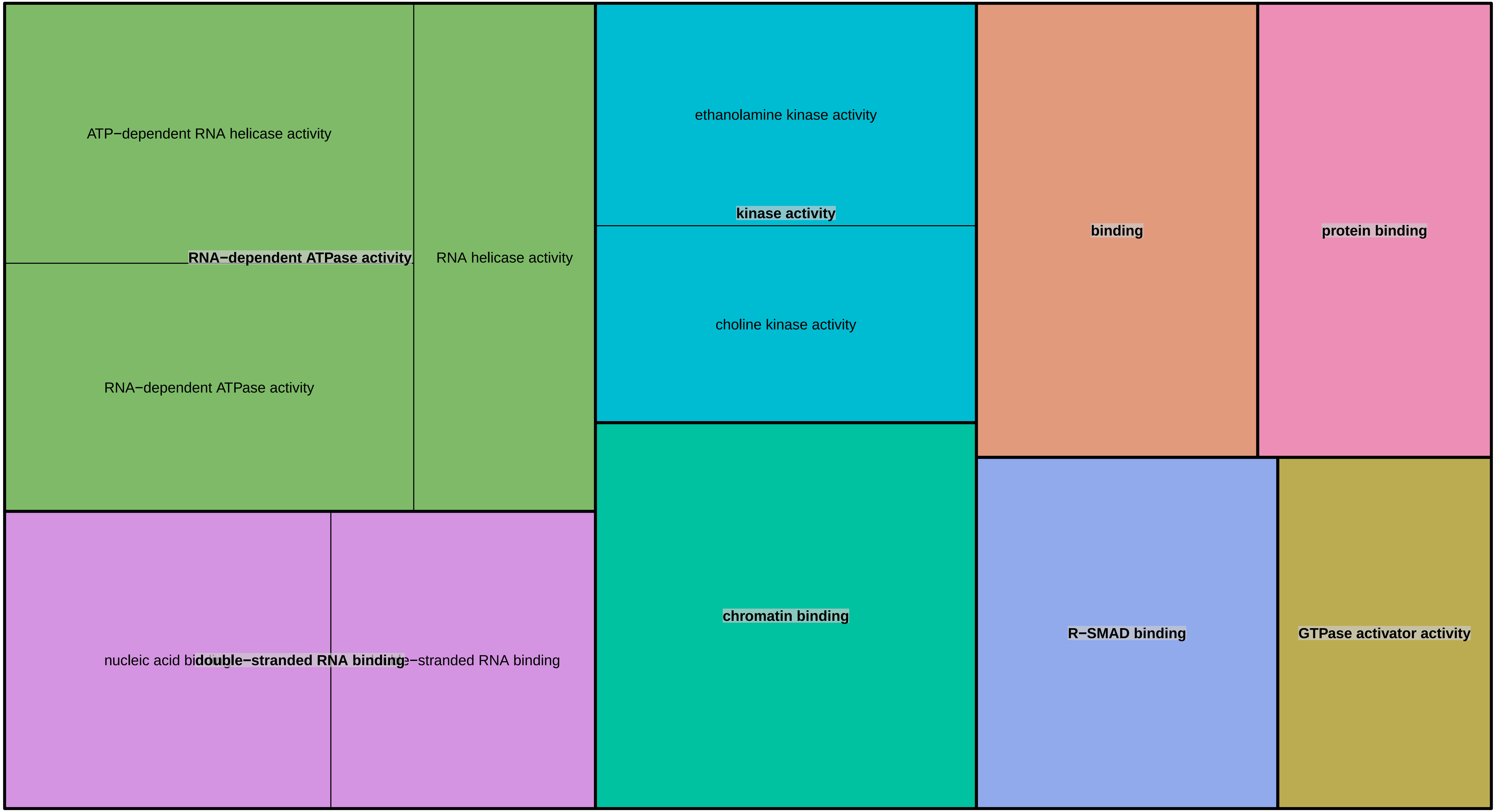

REVIGO Gene Ontology treemap

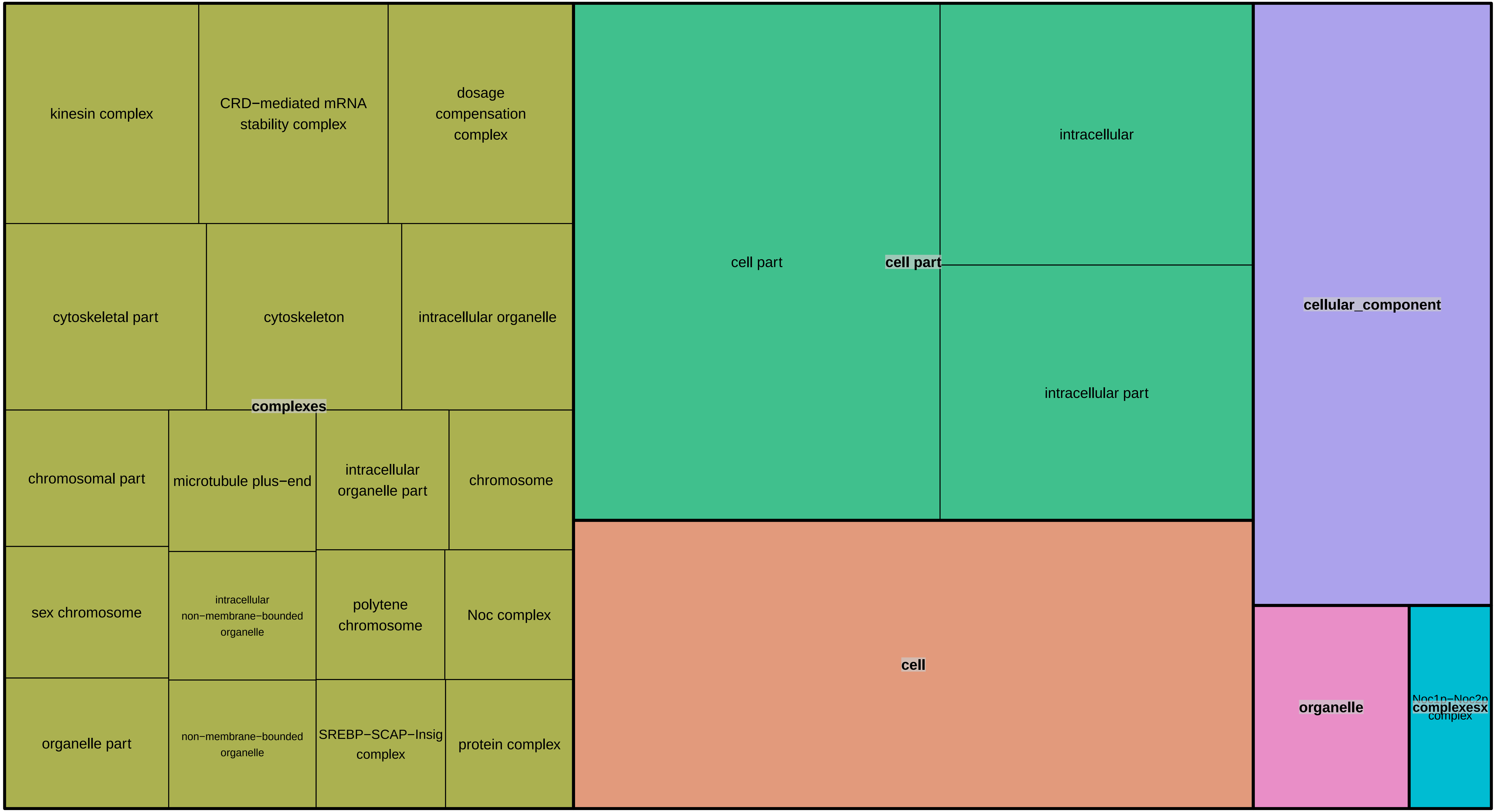
