## Supplementary material for "Local adaptation fuels cryptic speciation in terrestrial annelids": Suppl. tables

| Code | Population | Longitude | Latitude | Season | Year |
|---|---|---|---|---|---|
| BAR | Barcones | -2.8683 | 41.2983 | Autumn | 2013 |
| BOA | Boadilla del Monte | -3.925 | 40.4306 | Spring | 2008 |
| CNG | Congostrina | -2.9928 | 41.025 | Autumn | 2013 |
| COL | Las Colinas | -3.4367 | 40.7097 | Autumn | 2013 |
| F1 | Frontier 1 | -3.7175 | 40.9465 | Autumn | 2015 |
| HON | Hontoria | -4.1331 | 40.8928 | Autumn | 2012 |
| LOY | Lozoya | -3.7631 | 40.9492 | Spring | 2013 |
| PED | Pedraza | -3.8128 | 41.1008 | Autumn | 2012 |
| PNT2 | Punto 2 | -3.7028 | 40.7953 | Spring | 2011 |
| POL | Cabañas de Polendos | -4.1 | 41.0522 | Autumn | 2012 |
| PR1 | Pardo1 | -3.7622 | 40.5189 | Spring | 2013 |
| SAL | Saldaña de Ayllón | -3.4283 | 41.3864 | Spring | 2009 |
| SEV | Sevilla la Nueva | -4.0133 | 40.3447 | Spring | 2007 |
| TAM | Tamajón | -3.2531 | 40.9661 | Autumn | 2013 |
| TB | Tomillar B | -3.7103 | 40.9507 | Autumn | 2015 |
| UCE | Cubillo de Uceda | -3.4219 | 40.8272 | Spring | 2008 |
| VRN | Valderromán | -3.1192 | 41.3644 | Autumn | 2013 |

| | BAR | BOA | CNG | COL | F1 | HON | LOY | PED |
|---|---|---|---|---|---|---|---|---|
| BAR | 0 | 0.78 | 0.71 | 0.65 | 0.72 | 0.55 | 0.7 | 0.59 |
| BOA | 0.78 | 0 | 0.73 | 0.69 | 0.84 | 0.8 | 0.84 | 0.77 |
| CNG | 0.71 | 0.73 | 0 | 0.65 | 0.77 | 0.74 | 0.76 | 0.7 |
| COL | 0.65 | 0.69 | 0.65 | 0 | 0.68 | 0.66 | 0.66 | 0.59 |
| F1 | 0.72 | 0.84 | 0.77 | 0.68 | 0 | 0.73 | 0.76 | 0.42 |
| HON | 0.55 | 0.8 | 0.74 | 0.66 | 0.73 | 0 | 0.7 | 0.62 |
| LOY | 0.7 | 0.84 | 0.76 | 0.66 | 0.76 | 0.7 | 0 | 0.66 |
| PED | 0.59 | 0.77 | 0.7 | 0.59 | 0.42 | 0.62 | 0.66 | 0 |
| PNT2 | 0.6 | 0.66 | 0.62 | 0.34 | 0.62 | 0.6 | 0.62 | 0.55 |
| POL | 0.41 | 0.7 | 0.64 | 0.56 | 0.6 | 0.31 | 0.57 | 0.52 |
| PR1 | 0.62 | 0.72 | 0.67 | 0.35 | 0.64 | 0.63 | 0.67 | 0.58 |
| SAL | 0.7 | 0.74 | 0.66 | 0.64 | 0.77 | 0.74 | 0.76 | 0.7 |
| SEV | 0.75 | 0.44 | 0.7 | 0.67 | 0.78 | 0.74 | 0.79 | 0.7 |
| TAM | 0.6 | 0.63 | 0.3 | 0.56 | 0.62 | 0.63 | 0.64 | 0.59 |
| TB | 0.75 | 0.82 | 0.76 | 0.53 | 0.81 | 0.75 | 0.79 | 0.71 |
| UCE | 0.64 | 0.72 | 0.69 | 0.51 | 0.65 | 0.64 | 0.68 | 0.59 |
| VRN | 0.71 | 0.75 | 0.66 | 0.63 | 0.75 | 0.73 | 0.75 | 0.67 |

| PNT2 | POL | PR1 | SAL | SEV | TAM | TB | UCE | VRN |
|---|---|---|---|---|---|---|---|---|
| 0.6 | 0.41 | 0.62 | 0.7 | 0.75 | 0.6 | 0.75 | 0.64 | 0.71 |
| 0.66 | 0.7 | 0.72 | 0.74 | 0.44 | 0.63 | 0.82 | 0.72 | 0.75 |
| 0.62 | 0.64 | 0.67 | 0.66 | 0.7 | 0.3 | 0.76 | 0.69 | 0.66 |
| 0.34 | 0.56 | 0.35 | 0.64 | 0.67 | 0.56 | 0.53 | 0.51 | 0.63 |
| 0.62 | 0.6 | 0.64 | 0.77 | 0.78 | 0.62 | 0.81 | 0.65 | 0.75 |
| 0.6 | 0.31 | 0.63 | 0.74 | 0.74 | 0.63 | 0.75 | 0.64 | 0.73 |
| 0.62 | 0.57 | 0.67 | 0.76 | 0.79 | 0.64 | 0.79 | 0.68 | 0.75 |
| 0.55 | 0.52 | 0.58 | 0.7 | 0.7 | 0.59 | 0.71 | 0.59 | 0.67 |
| 0 | 0.51 | 0.3 | 0.61 | 0.62 | 0.55 | 0.45 | 0.48 | 0.62 |
| 0.51 | 0 | 0.52 | 0.62 | 0.66 | 0.55 | 0.64 | 0.55 | 0.62 |
| 0.3 | 0.52 | 0 | 0.67 | 0.66 | 0.55 | 0.51 | 0.49 | 0.66 |
| 0.61 | 0.62 | 0.67 | 0 | 0.7 | 0.59 | 0.75 | 0.66 | 0.6 |
| 0.62 | 0.66 | 0.66 | 0.7 | 0 | 0.62 | 0.78 | 0.67 | 0.7 |
| 0.55 | 0.55 | 0.55 | 0.59 | 0.62 | 0 | 0.66 | 0.59 | 0.57 |
| 0.45 | 0.64 | 0.51 | 0.75 | 0.78 | 0.66 | 0 | 0.6 | 0.74 |
| 0.48 | 0.55 | 0.49 | 0.66 | 0.67 | 0.59 | 0.6 | 0 | 0.65 |
| 0.62 | 0.62 | 0.66 | 0.6 | 0.7 | 0.57 | 0.74 | 0.65 | 0 |

| Discriminant function | | | Cross-validation | | |
|---|---|---|---|---|---|
| True | Allocated to | | Allocated to | | |
| | 1 | 2 | 1 | 2 | Total |
| 1 | 12 | 0 | 11 | 1 | 12 |
| 2 | 0 | 15 | 0 | 15 | 15 |
| | 1 | 3 | 1 | 3 | |
| 1 | 12 | 0 | 12 | 0 | 12 |
| 3 | 1 | 16 | 1 | 16 | 17 |
| | 2 | 3 | 2 | 3 | |
| 2 | 9 | 6 | 9 | 6 | 15 |
| 3 | 2 | 15 | 5 | 12 | 17 |
| Discriminant function | | | Cross-validation | | |
| True | Allocated to | | Allocated to | | |
| | 1 | 2 | 1 | 2 | Total |
| 1 | 12 | 0 | 10 | 2 | 12 |
| 2 | 0 | 10 | 0 | 10 | 10 |
| | 1 | 3 | 1 | 3 | |
| 1 | 12 | 0 | 12 | 0 | 12 |
| 3 | 1 | 6 | 1 | 6 | 7 |
| | 1 | 4 | 1 | 4 | |
| 1 | 12 | 0 | 11 | 1 | 12 |
| 4 | 0 | 5 | 0 | 5 | 5 |
| | 1 | 5 | 1 | 5 | |
| 1 | 12 | 0 | 11 | 1 | 12 |
| 5 | 0 | 4 | 0 | 4 | 4 |
| | 1 | 6 | 1 | 6 | |
| 1 | 12 | 0 | 11 | 1 | 12 |
| 6 | 0 | 6 | 1 | 5 | 6 |
| | 2 | 3 | 2 | 3 | |
| 2 | 10 | 0 | 6 | 4 | 10 |
| 3 | 0 | 7 | 1 | 6 | 7 |
| | 2 | 4 | 2 | 4 | |
| 2 | 10 | 0 | 6 | 4 | 10 |
| 4 | 0 | 5 | 1 | 4 | 5 |
| | 2 | 5 | 2 | 5 | |
| 2 | 10 | 0 | 8 | 2 | 10 |
| 5 | 0 | 4 | 1 | 3 | 4 |
| | 2 | 6 | 2 | 6 | |
| 2 | 10 | 0 | 8 | 2 | 10 |
| 6 | 0 | 6 | 0 | 6 | 6 |
| | 3 | 4 | 3 | 4 | |
| 3 | 7 | 0 | 5 | 2 | 7 |
| 4 | 0 | 5 | 2 | 3 | 5 |
| | 3 | 5 | 3 | 5 | |
| 3 | 6 | 1 | 4 | 3 | 7 |
| 5 | 1 | 3 | 3 | 1 | 4 |
| | 3 | 6 | 3 | 6 | |
| 3 | 7 | 0 | 5 | 2 | 7 |
| 6 | 0 | 6 | 2 | 4 | 6 |
| | 4 | 5 | 4 | 5 | |

| | | | | | |
|---|---|---|---|---|---|
| 4 | 5 | 0 | 4 | 1 | 5 |
| 5 | 0 | 4 | 2 | 2 | 4 |
| | 4 | 6 | 4 | 6 | |
| 4 | 5 | 0 | 4 | 1 | 5 |
| 6 | 0 | 6 | 2 | 4 | 6 |
| | 5 | 6 | 5 | 6 | |
| 5 | 4 | 0 | 3 | 1 | 4 |
| 6 | 0 | 6 | 2 | 4 | 6 |

| Contig | GO terms | Putative protein |
| --- | --- | --- |
| comp217417_c2_seq8.p1 | GO:0000122,GO:0000578,GO:0000902,GO:0000904,GO:0000981,GO:0000982,GO:0001071,GO:0001078,GO:0001227,GO:0001667,GO:0001709,GO:0003002,GO:0003674,GO:0003700,GO:0004871,GO:0004872,GO:0004879,GO:0005488,GO:0005515,GO:0005575,GO:0005622,GO:0005623,GO:0005634,GO:0006139,GO:0006351,GO:0006355,GO:0006357,GO:0006366,GO:0006725,GO:0006807,GO:0006928,GO:0007088,GO:0007154,GO:0007165,GO:0007275,GO:0007346,GO:0007350,GO:0007351,GO:0007354,GO:0007389,GO:0007399,GO:0007417,GO:0007424,GO:0007427,GO:0007517,GO:0008134,GO:0008150,GO:0008152,GO:0008544,GO:0008595,GO:0009058,GO:0009059,GO:0009653,GO:0009790,GO:0009798,GO:0009880,GO:0009888,GO:0009889,GO:0009890,GO:0009892,GO:0009948,GO:0009952,GO:0009987,GO:0010467,GO:0010468,GO:0010556,GO:0010558,GO:0010564,GO:0010605,GO:0010629,GO:0010631,GO:0016043,GO:0016070,GO:0016358,GO:0016477,GO:0018130,GO:0019219,GO:0019222,GO:0019438,GO:0022008,GO:0023052,GO:0030030,GO:0030154,GO:0030182,GO:0030522,GO:0031175,GO:0031323,GO:0031324,GO:0031326,GO:0031327,GO:0032501,GO:0032502,GO:0032774,GO:0032989,GO:0032990,GO:0033043,GO:0034641,GO:0034645,GO:0034654,GO:0035150,GO:0035151,GO:0035152,GO:0035282,GO:0035290,GO:0038023,GO:0040011,GO:0043170,GO:0043226,GO:0043227,GO:0043229,GO:0043231,GO:0043588,GO:0044237,GO:0044238,GO:0044249,GO:0044260,GO:0044271,GO:0044424,GO:0044464,GO:0044699,GO:0044700,GO:0044707,GO:0044763,GO:0044767,GO:0045165,GO:0045892,GO:0045934,GO:0046483,GO:0046845,GO:0048468,GO:0048513,GO:0048519,GO:0048523,GO:0048666,GO:0048667,GO:0048699,GO:0048731,GO:0048812,GO:0048813,GO:0048856,GO:0048858,GO:0048869,GO:0048870,GO:0050789,GO:0050794,GO:0050896,GO:0051128,GO:0051171,GO:0051172,GO:0051179,GO:0051252,GO:0051253,GO:0051674,GO:0051716,GO:0051726,GO:0051783,GO:0060089,GO:0060255,GO:0060541,GO:0061061,GO:0065007,GO:0065008,GO:0070491,GO:0071704,GO:0071840,GO:0080090,GO:0090066,GO:0090130,GO:0090132,GO:0090304,GO:1901360,GO:1901362,GO:1901576,GO:2000112,GO:2000113,GO:2001141 | branched duct epithelial cell fate determination, open tracheal system |

| | | |
| --- | --- | --- |
| comp220340_c0_seq3.p1 | GO:0000003,GO:0000070,GO:0000090,GO:0000212,GO:0000226,GO:0000228,GO:0000278,GO:0000279,GO:0000280,GO:0000793,GO:0000794,GO:0000819,GO:0000910,GO:0001775,GO:0003006,GO:0003674,GO:0003824,GO:0005488,GO:0005515,GO:0005575,GO:0005622,GO:0005623,GO:0005634,GO:0005694,GO:0005737,GO:0005813,GO:0005815,GO:0005819,GO:0005856,GO:0005938,GO:0006810,GO:0006887,GO:0006897,GO:0006909,GO:0006911,GO:0006915,GO:0006996,GO:0007010,GO:0007017,GO:0007049,GO:0007051,GO:0007059,GO:0007063,GO:0007067,GO:0007088,GO:0007097,GO:0007126,GO:0007127,GO:0007143,GO:0007275,GO:0007276,GO:0007292,GO:0007338,GO:0007343,GO:0007346,GO:0007399,GO:0007548,GO:0008104,GO:0008150,GO:0008152,GO:0008219,GO:0008233,GO:0008234,GO:0009566,GO:0009653,GO:0009790,GO:0009792,GO:0009987,GO:0010324,GO:0010564,GO:0010638,GO:0010639,GO:0010948,GO:0012501,GO:0015630,GO:0016023,GO:0016043,GO:0016044,GO:0016192,GO:0016265,GO:0016787,GO:0017156,GO:0019953,GO:0022008,GO:0022402,GO:0022403,GO:0022414,GO:0030071,GO:0030141,GO:0030154,GO:0030496,GO:0030703,GO:0031410,GO:0031503,GO:0031974,GO:0031981,GO:0031982,GO:0031988,GO:0032153,GO:0032154,GO:0032155,GO:0032501,GO:0032502,GO:0032504,GO:0032940,GO:0032991,GO:0033036,GO:0033043,GO:0033044,GO:0033206,GO:0035046,GO:0040001,GO:0040012,GO:0040023,GO:0040035,GO:0040038,GO:0043226,GO:0043227,GO:0043228,GO:0043229,GO:0043231,GO:0043232,GO:0043233,GO:0043234,GO:0043277,GO:0043652,GO:0043933,GO:0044422,GO:0044424,GO:0044428,GO:0044430,GO:0044444,GO:0044446,GO:0044464,GO:0044699,GO:0044702,GO:0044707,GO:0044763,GO:0044765,GO:0044767,GO:0045132,GO:0045143,GO:0045787,GO:0045840,GO:0045842,GO:0045875,GO:0046903,GO:0048285,GO:0048477,GO:0048513,GO:0048518,GO:0048519,GO:0048522,GO:0048523,GO:0048580,GO:0048608,GO:0048609,GO:0048610,GO:0048646,GO:0048731,GO:0048806,GO:0048856,GO:0048869,GO:0050789,GO:0050793,GO:0050794,GO:0051128,GO:0051129,GO:0051130,GO:0051179,GO:0051234,GO:0051239,GO:0051276,GO:0051293,GO:0051301,GO:0051304,GO:0051306,GO:0051321,GO:0051322,GO:0051640,GO:0051641,GO:0051647,GO:0051649,GO:0051653,GO:0051656,GO:0051726,GO:0051783,GO:0051785,GO:0051983,GO:0060467,GO:0060468,GO:0060471,GO:0060473,GO:0061024,GO:0061062,GO:0061458,GO:0065007,GO:0070011,GO:0070013,GO:0070090,GO:0071822,GO:0071840,GO:0071944,GO:0072686,GO:0072687,GO:0090068,GO:1901987,GO:1901989,GO:1901990,GO:1901992,GO:1902099,GO:1902101,GO:2000026,GO:2000241,GO:2000242,GO:2001251 | extra spindle pole bodies homolog 1 (S. cerevisiae) |
| comp210439_c0_seq1.p1 | GO:0001756,GO:0001775,GO:0001776,GO:0001782,GO:0002253,GO:0002260,GO:0002376,GO:0002429,GO:0002520,GO:0002521,GO:0002682,GO:0002684,GO:0002757,GO:0002764,GO:0002768,GO:0003002,GO:0003674,GO:0004857,GO:0004866,GO:0004869,GO:0005575,GO:0005622,GO:0005623,GO:0005737,GO:0005739,GO:0006915,GO:0007154,GO:0007165,GO:0007166,GO:0007275,GO:0007369,GO:0007389,GO:0008150,GO:0008219,GO:0009653,GO:0009790,GO:0009792,GO:0009952,GO:0009987,GO:0010466,GO:0010522,GO:0010941,GO:0010951,GO:0010959,GO:0012501,GO:0016265,GO:0019222,GO:0023052,GO:0030097,GO:0030098,GO:0030154,GO:0030217,GO:0030234,GO:0030414,GO:0032386,GO:0032501,GO:0032502,GO:0032844,GO:0032879,GO:0035282,GO:0042110,GO:0042592,GO:0042981,GO:0043009,GO:0043027,GO:0043028,GO:0043066,GO:0043067,GO:0043069,GO:0043086,GO:0043154,GO:0043226,GO:0043227,GO:0043229,GO:0043231,GO:0043269,GO:0043281,GO:0044092,GO:0044424,GO:0044444,GO:0044464,GO:0044699,GO:0044700,GO:0044707,GO:0044763,GO:0044767,GO:0045321,GO:0046649,GO:0048513,GO:0048518,GO:0048519,GO:0048523,GO:0048534,GO:0048583,GO:0048584,GO:0048598,GO:0048646,GO:0048731,GO:0048856,GO:0048869,GO:0048872,GO:0050776,GO:0050778,GO:0050789,GO:0050790,GO:0050794,GO:0050851,GO:0050852,GO:0050896,GO:0051049,GO:0051279,GO:0051336,GO:0051346,GO:0051716,GO:0051924,GO:0052547,GO:0052548,GO:0060341,GO:0060548,GO:0061053,GO:0061134,GO:0061135,GO:0065007,GO:0065008,GO:0065009,GO:2000021,GO:2000116,GO:2000117 | Bcl-2-related protein A1 |
| comp213097_c0_seq8.p1 | | PTC7 protein phosphatase homolog (S. cerevisiae) |

| | | |
|---|---|---|
| comp219622_c0_seq5.p1 | GO:0003674,GO:0003824,GO:0004842,GO:0005575,GO:0005622,GO:0005623,GO:0005634,GO:0005737,GO:0006464,GO:0006508,GO:0006511,GO:0008150,GO:0008152,GO:0009056,GO:0009057,GO:0009987,GO:0016567,GO:0016874,GO:0016879,GO:0016881,GO:0019538,GO:0019787,GO:0019941,GO:0030163,GO:0032446,GO:0036211,GO:0042787,GO:0043170,GO:0043226,GO:0043227,GO:0043229,GO:0043231,GO:0043412,GO:0043632,GO:0044237,GO:0044238,GO:0044248,GO:0044257,GO:0044260,GO:0044265,GO:0044267,GO:0044424,GO:0044464,GO:0051603,GO:0070647,GO:0071704,GO:1901575 | HECT domain containing E3 ubiquitin protein ligase 1 |
| comp213188_c0_seq4.p1 | | coiled-coil domain containing 60 |
| comp219530_c3_seq2.p1 | GO:0000375,GO:0000377,GO:0000398,GO:0005575,GO:0005622,GO:0005623,GO:0005634,GO:0005654,GO:0006139,GO:0006396,GO:0006397,GO:0006725,GO:0006807,GO:0008150,GO:0008152,GO:0008380,GO:0009987,GO:0010467,GO:0016070,GO:0016071,GO:0031974,GO:0031981,GO:0034641,GO:0043170,GO:0043226,GO:0043227,GO:0043229,GO:0043231,GO:0043233,GO:0044237,GO:0044238,GO:0044260,GO:0044422,GO:0044424,GO:0044428,GO:0044446,GO:0044464,GO:0046483,GO:0070013,GO:0071704,GO:0090304,GO:1901360 | SURP and G patch domain containing 1 |
| comp209633_c0_seq1.p1 | | Coiled-coil domain containing |
| comp204426_c0_seq2.p1 | GO:0005575,GO:0005622,GO:0005623,GO:0005737,GO:0044424,GO:0044464 | Ubiquitin associated protein 1 |
| comp219472_c0_seq6.p1 | | Inherit from biNOG: Family with sequence similarity 193, member A |
| comp211313_c0_seq1.p1 | GO:0000070,GO:0000278,GO:0000280,GO:0000819,GO:0005575,GO:0005622,GO:0005623,GO:0005856,GO:0005871,GO:0005875,GO:0006996,GO:0007049,GO:0007059,GO:0007067,GO:0008150,GO:0009892,GO:0009894,GO:0009895,GO:0009987,GO:0010506,GO:0010507,GO:0015630,GO:0016043,GO:0019222,GO:0022402,GO:0031323,GO:0031324,GO:0031329,GO:0031330,GO:0032991,GO:0043226,GO:0043228,GO:0043229,GO:0043232,GO:0043234,GO:0044422,GO:0044424,GO:0044430,GO:0044446,GO:0044464,GO:0044699,GO:0044763,GO:0048285,GO:0048519,GO:0048523,GO:0050789,GO:0050794,GO:0051276,GO:0065007,GO:0071840 | kinesin family member 25 |
| comp211313_c0_seq1.p2 | GO:0000070,GO:0000278,GO:0000280,GO:0000819,GO:0005575,GO:0005622,GO:0005623,GO:0005856,GO:0005871,GO:0005875,GO:0006996,GO:0007049,GO:0007059,GO:0007067,GO:0008150,GO:0009892,GO:0009894,GO:0009895,GO:0009987,GO:0010506,GO:0010507,GO:0015630,GO:0016043,GO:0019222,GO:0022402,GO:0031323,GO:0031324,GO:0031329,GO:0031330,GO:0032991,GO:0043226,GO:0043228,GO:0043229,GO:0043232,GO:0043234,GO:0044422,GO:0044424,GO:0044430,GO:0044446,GO:0044464,GO:0044699,GO:0044763,GO:0048285,GO:0048519,GO:0048523,GO:0050789,GO:0050794,GO:0051276,GO:0065007,GO:0071840 | kinesin family member 25 |
| comp211313_c0_seq1.p3 | GO:0000070,GO:0000278,GO:0000280,GO:0000819,GO:0005575,GO:0005622,GO:0005623,GO:0005856,GO:0005871,GO:0005875,GO:0006996,GO:0007049,GO:0007059,GO:0007067,GO:0008150,GO:0009892,GO:0009894,GO:0009895,GO:0009987,GO:0010506,GO:0010507,GO:0015630,GO:0016043,GO:0019222,GO:0022402,GO:0031323,GO:0031324,GO:0031329,GO:0031330,GO:0032991,GO:0043226,GO:0043228,GO:0043229,GO:0043232,GO:0043234,GO:0044422,GO:0044424,GO:0044430,GO:0044446,GO:0044464,GO:0044699,GO:0044763,GO:0048285,GO:0048519,GO:0048523,GO:0050789,GO:0050794,GO:0051276,GO:0065007,GO:0071840 | kinesin family member 25 |
| comp203597_c2_seq5.p1 | GO:0001525,GO:0001568,GO:0001944,GO:0007275,GO:0008150,GO:0009653,GO:0032501,GO:0032502,GO:0044699,GO:0044707,GO:0044767,GO:0048514,GO:0048646,GO:0048731,GO:0048856,GO:0072358,GO:0072359 | hexosaminidase B (beta polypeptide) |

| | | |
|---|---|---|
| comp203597_c2_seq5.p2 | GO:0001525,GO:0001568,GO:0001944,GO:0007275,GO:0008150,GO:0009653,GO:0032501,GO:0032502,GO:0044699,GO:0044707,GO:0044767,GO:0048514,GO:0048646,GO:0048731,GO:0048856,GO:0072358,GO:0072359 | hexosaminidase B (beta polypeptide) |
| comp209470_c0_seq4.p1 | | serine threonine-protein phosphatase 6 regulatory ankyrin repeat subunit |
| comp200606_c0_seq2.p1 | GO:0000988,GO:0000989,GO:0003674,GO:0003712,GO:0003713,GO:0005575,GO:0005622,GO:0005623,GO:0005634,GO:0005737,GO:0005793,GO:0005801,GO:0005856,GO:0006355,GO:0006357,GO:0006917,GO:0006928,GO:0006950,GO:0006996,GO:0007010,GO:0007015,GO:0007049,GO:0007050,GO:0008064,GO:0008150,GO:0009889,GO:0009987,GO:0010468,GO:0010556,GO:0010638,GO:0010941,GO:0010942,GO:0012502,GO:0012506,GO:0016020,GO:0016023,GO:0016043,GO:0019219,GO:0019222,GO:0022402,GO:0030029,GO:0030036,GO:0030659,GO:0030832,GO:0030833,GO:0030838,GO:0031090,GO:0031252,GO:0031323,GO:0031326,GO:0031334,GO:0031410,GO:0031982,GO:0031988,GO:0032271,GO:0032273,GO:0032535,GO:0032956,GO:0032970,GO:0033043,GO:0033554,GO:0034314,GO:0040011,GO:0042981,GO:0043065,GO:0043067,GO:0043068,GO:0043226,GO:0043227,GO:0043228,GO:0043229,GO:0043231,GO:0043232,GO:0043254,GO:0043620,GO:0043933,GO:0044087,GO:0044093,GO:0044422,GO:0044424,GO:0044433,GO:0044444,GO:0044446,GO:0044464,GO:0044699,GO:0044763,GO:0045010,GO:0045786,GO:0048518,GO:0048519,GO:0048522,GO:0048523,GO:0048870,GO:0050789,GO:0050794,GO:0050896,GO:0051090,GO:0051091,GO:0051128,GO:0051130,GO:0051171,GO:0051179,GO:0051252,GO:0051493,GO:0051495,GO:0051674,GO:0051716,GO:0051726,GO:0060255,GO:0065007,GO:0065008,GO:0065009,GO:0070060,GO:0070358,GO:0071822,GO:0071840,GO:0080090,GO:0090066,GO:2000112,GO:2001141 | WAS protein homolog associated with actin, golgi membranes and microtubules |
| comp220940_c0_seq10.p1 | GO:0003674,GO:0003824,GO:0004842,GO:0006464,GO:0008150,GO:0008152,GO:0009987,GO:0016567,GO:0016874,GO:0016879,GO:0016881,GO:0019538,GO:0019787,GO:0032446,GO:0036211,GO:0043170,GO:0043412,GO:0044237,GO:0044238,GO:0044260,GO:0044267,GO:0051865,GO:0070647,GO:0071704 | listerin E3 ubiquitin protein ligase 1 |
| comp220940_c0_seq10.p2 | | Reeler domain |
| comp215017_c3_seq2.p1 | | Ricin-type beta-trefoil lectin domain |

| | | |
|---|---|---|
| comp213838_c0_seq1.p1 | GO:0003674,GO:0005488,GO:0005575,GO:0005622,GO:0005623,GO:0005634,GO:0005737,GO:0005829,GO:0008150,GO:0032886,GO:0033218,GO:0042277,GO:0043226,GO:0043227,GO:0043229,GO:0043231,GO:0044424,GO:0044444,GO:0044464,GO:0050789,GO:0050794,GO:0065007 | May control cell migration by relaying extracellular chemotactic signals to the microtubule cytoskeleton. Mediator of ERBB2 signaling. The MEMO1-RHOA-DIAPH1 signaling pathway plays an important role in ERBB2-dependent stabilization of microtubules at the cell cortex. It controls the localization of APC and CLASP2 to the cell membrane, via the regulation of GSK3B activity. In turn, membrane-bound APC allows the localization of the MACF1 to the cell membrane, which is required for microtubule capture and stabilization (By similarity) |
| comp221241_c0_seq6.p1 | | Inherit from KOG: protein tyrosine phosphatase receptor type |
| comp221241_c0_seq6.p2 | | Kinesin family member |
| comp206443_c0_seq3.p1 | GO:0005575,GO:0005622,GO:0005623,GO:0005737,GO:0044424,GO:0044464 | Tetratricopeptide repeat domain 25 |

| | | |
|---|---|---|
| comp208585_c0_seq1.p1 | GO:0000075,GO:0000082,GO:0000084,GO:0000122,GO:0000226,GO:0000278,GO:0000302,GO:0000910,GO:0001666,GO:0002039,GO:0003008,GO:0003674,GO:0003824,GO:0004672,GO:0004674,GO:0004871,GO:0005488,GO:0005515,GO:0005575,GO:0005622,GO:0005623,GO:0005634,GO:0005730,GO:0005737,GO:0005794,GO:0005795,GO:0005813,GO:0005814,GO:0005815,GO:0005856,GO:0006355,GO:0006357,GO:0006464,GO:0006468,GO:0006793,GO:0006796,GO:0006915,GO:0006950,GO:0006970,GO:0006974,GO:0006979,GO:0006996,GO:0007010,GO:0007017,GO:0007030,GO:0007049,GO:0007051,GO:0007052,GO:0007093,GO:0007113,GO:0007154,GO:0007165,GO:0007243,GO:0007264,GO:0007265,GO:0007346,GO:0007610,GO:0007611,GO:0007613,GO:0008150,GO:0008152,GO:0008219,GO:0009314,GO:0009628,GO:0009889,GO:0009890,GO:0009892,GO:0009893,GO:0009894,GO:0009896,GO:0009966,GO:0009967,GO:0009987,GO:0010468,GO:0010556,GO:0010558,GO:0010564,GO:0010604,GO:0010605,GO:0010627,GO:0010629,GO:0010646,GO:0010647,GO:0010648,GO:0010740,GO:0010824,GO:0010941,GO:0010948,GO:0012501,GO:0015630,GO:0016043,GO:0016265,GO:0016301,GO:0016310,GO:0016740,GO:0016772,GO:0016773,GO:0019219,GO:0019222,GO:0019538,GO:0022402,GO:0022403,GO:0022411,GO:0023051,GO:0023052,GO:0023056,GO:0023057,GO:0030162,GO:0030425,GO:0031323,GO:0031324,GO:0031325,GO:0031326,GO:0031327,GO:0031329,GO:0031331,GO:0031644,GO:0031645,GO:0031646,GO:0031974,GO:0031981,GO:0032268,GO:0032270,GO:0032434,GO:0032436,GO:0032486,GO:0032501,GO:0032886,GO:0033043,GO:0033554,GO:0035556,GO:0036211,GO:0036293,GO:0036294,GO:0042176,GO:0042221,GO:0042981,GO:0042995,GO:0043005,GO:0043025,GO:0043066,GO:0043067,GO:0043069,GO:0043122,GO:0043123,GO:0043170,GO:0043226,GO:0043227,GO:0043228,GO:0043229,GO:0043231,GO:0043232,GO:0043233,GO:0043412,GO:0043491,GO:0043933,GO:0044057,GO:0044237,GO:0044238,GO:0044260,GO:0044267,GO:0044297,GO:0044422,GO:0044424,GO:0044428,GO:0044430,GO:0044431,GO:0044444,GO:0044446,GO:0044450,GO:0044464,GO:0044699,GO:0044700,GO:0044707,GO:0044708,GO:0044763,GO:0044770,GO:0044772,GO:0045732,GO:0045862,GO:0045892,GO:0045934,GO:0046599,GO:0046605,GO:0048167,GO:0048168,GO:0048169,GO:0048308,GO:0048313,GO:0048518,GO:0048519,GO:0048522,GO:0048523,GO:0048583,GO:0048584,GO:0050789,GO:0050794,GO:0050804,GO:0050805,GO:0050806,GO:0050877,GO:0050890,GO:0050896,GO:0051128,GO:0051171,GO:0051172,GO:0051239,GO:0051240,GO:0051241,GO:0051246,GO:0051247,GO:0051252,GO:0051253,GO:0051301,GO:0051320,GO:0051325,GO:0051329,GO:0051493,GO:0051716,GO:0051726,GO:0051969,GO:0051970,GO:0051971,GO:0060089,GO:0060255,GO:0060291,GO:0060292,GO:0060548,GO:0061136,GO:0065007,GO:0065008,GO:0070013,GO:0070482,GO:0070507,GO:0070887,GO:0071453,GO:0071456,GO:0071704,GO:0071822,GO:0071840,GO:0080090,GO:0090166,GO:0097458,GO:1901700,GO:1901800,GO:1901987,GO:1901988,GO:1901990,GO:1901991,GO:2000112,GO:2000113,GO:2000777,GO:2001141 | Polo-like kinase |

| | | |
|---|---|---|
| comp210368_c0_seq4.p1 | GO:0000166,GO:0003008,GO:0003674,GO:0003824,GO:0003924,GO:0005488,GO:0005515,GO:0005575,GO:0005622,GO:0005623,GO:0005737,GO:0005783,GO:0005789,GO:0005794,GO:0006139,GO:0006152,GO:0006163,GO:0006184,GO:0006195,GO:0006725,GO:0006753,GO:0006793,GO:0006796,GO:0006807,GO:0006944,GO:0006996,GO:0007154,GO:0007267,GO:0007268,GO:0007275,GO:0007517,GO:0007519,GO:0007528,GO:0008150,GO:0008152,GO:0008582,GO:0009056,GO:0009116,GO:0009117,GO:0009119,GO:0009141,GO:0009143,GO:0009144,GO:0009146,GO:0009150,GO:0009154,GO:0009164,GO:0009166,GO:0009199,GO:0009203,GO:0009205,GO:0009207,GO:0009259,GO:0009261,GO:0009888,GO:0009987,GO:0012505,GO:0014706,GO:0016020,GO:0016043,GO:0016044,GO:0016202,GO:0016320,GO:0016462,GO:0016787,GO:0016817,GO:0016818,GO:0017076,GO:0017111,GO:0019001,GO:0019226,GO:0019439,GO:0019637,GO:0019693,GO:0023052,GO:0030154,GO:0031090,GO:0031110,GO:0031114,GO:0031224,GO:0031227,GO:0031300,GO:0032501,GO:0032502,GO:0032553,GO:0032555,GO:0032561,GO:0032886,GO:0033043,GO:0034641,GO:0034655,GO:0035637,GO:0036094,GO:0040008,GO:0042175,GO:0042278,GO:0042454,GO:0042692,GO:0042802,GO:0042803,GO:0043226,GO:0043227,GO:0043229,GO:0043231,GO:0043244,GO:0044087,GO:0044237,GO:0044238,GO:0044248,GO:0044270,GO:0044281,GO:0044422,GO:0044424,GO:0044425,GO:0044432,GO:0044444,GO:0044446,GO:0044464,GO:0044699,GO:0044700,GO:0044707,GO:0044710,GO:0044763,GO:0044767,GO:0045595,GO:0046039,GO:0046128,GO:0046130,GO:0046434,GO:0046483,GO:0046700,GO:0046983,GO:0048284,GO:0048468,GO:0048513,GO:0048634,GO:0048638,GO:0048641,GO:0048731,GO:0048741,GO:0048742,GO:0048747,GO:0048856,GO:0048869,GO:0050789,GO:0050793,GO:0050794,GO:0050803,GO:0050807,GO:0050808,GO:0050877,GO:0051128,GO:0051146,GO:0051147,GO:0051153,GO:0051239,GO:0051493,GO:0051960,GO:0051963,GO:0055001,GO:0055002,GO:0055086,GO:0060284,GO:0060537,GO:0060538,GO:0061024,GO:0061025,GO:0061061,GO:0065007,GO:0065008,GO:0070507,GO:0071704,GO:0071840,GO:0072521,GO:0072523,GO:0090174,GO:0097159,GO:1901068,GO:1901069,GO:1901135,GO:1901136,GO:1901265,GO:1901292,GO:1901360,GO:1901361,GO:1901363,GO:1901564,GO:1901565,GO:1901575,GO:1901657,GO:1901658,GO:1901861,GO:1901879,GO:2000026 | Guanylate-binding protein, N-terminal domain |
| comp213562_c0_seq5.p1 | | Gamma-glutamyltranspeptidase |
| comp209455_c0_seq6.p1 | GO:0003674,GO:0003824,GO:0008150,GO:0008152,GO:0016491,GO:0016614,GO:0044710,GO:0055114 | dehydrogenase |
| comp211928_c0_seq7.p1 | | nucleoporin 214kDa |
| comp219252_c2_seq2.p1 | | mediator complex subunit 23 |
| comp187914_c1_seq1.p1 | GO:0003674,GO:0004872,GO:0005575,GO:0005622,GO:0005623,GO:0005737,GO:0005783,GO:0005789,GO:0005811,GO:0006412,GO:0006605,GO:0006612,GO:0006613,GO:0006614,GO:0006810,GO:0006886,GO:0006897,GO:0006909,GO:0008104,GO:0008150,GO:0008152,GO:0009058,GO:0009059,GO:0009987,GO:0010467,GO:0010876,GO:0012505,GO:0015031,GO:0016020,GO:0016021,GO:0016192,GO:0016482,GO:0019538,GO:0019915,GO:0031090,GO:0031224,GO:0032501,GO:0032963,GO:0032964,GO:0033036,GO:0033365,GO:0034613,GO:0034645,GO:0042175,GO:0043170,GO:0043226,GO:0043227,GO:0043229,GO:0043231,GO:0044236,GO:0044237,GO:0044238,GO:0044249,GO:0044259,GO:0044260,GO:0044267,GO:0044422,GO:0044424,GO:0044425,GO:0044432,GO:0044444,GO:0044446,GO:0044464,GO:0044699,GO:0044707,GO:0044710,GO:0044763,GO:0044765,GO:0045047,GO:0045184,GO:0046907,GO:0051179,GO:0051234,GO:0051641,GO:0051649,GO:0070727,GO:0070972,GO:0071702,GO:0071704,GO:0072594,GO:0072599,GO:1901576 | translocation associated membrane protein |

| | | |
|---|---|---|
| comp220560_c1_seq1.p1 | GO:0000122,GO:0003674,GO:0003824,GO:0004842,GO:0005488,GO:0005515,GO:0005575,GO:0005622,GO:0005623,GO:0005634,GO:0005654,GO:0006139,GO:0006351,GO:0006352,GO:0006355,GO:0006357,GO:0006366,GO:0006367,GO:0006464,GO:0006725,GO:0006807,GO:0007154,GO:0007165,GO:0007166,GO:0007167,GO:0007178,GO:0007179,GO:0008150,GO:0008152,GO:0008270,GO:0009058,GO:0009059,GO:0009719,GO:0009889,GO:0009890,GO:0009892,GO:0009966,GO:0009968,GO:0009987,GO:0010033,GO:0010467,GO:0010468,GO:0010556,GO:0010558,GO:0010605,GO:0010629,GO:0010646,GO:0010648,GO:0016070,GO:0016567,GO:0016874,GO:0016879,GO:0016881,GO:0017015,GO:0018130,GO:0019219,GO:0019222,GO:0019438,GO:0019538,GO:0019787,GO:0023051,GO:0023052,GO:0023057,GO:0030510,GO:0030514,GO:0031323,GO:0031324,GO:0031326,GO:0031327,GO:0031974,GO:0031981,GO:0032446,GO:0032774,GO:0034641,GO:0034645,GO:0034654,GO:0036211,GO:0042221,GO:0043167,GO:0043169,GO:0043170,GO:0043226,GO:0043227,GO:0043229,GO:0043231,GO:0043233,GO:0043412,GO:0044237,GO:0044238,GO:0044249,GO:0044260,GO:0044267,GO:0044271,GO:0044422,GO:0044424,GO:0044428,GO:0044446,GO:0044464,GO:0044699,GO:0044700,GO:0044763,GO:0045892,GO:0045934,GO:0046332,GO:0046483,GO:0046872,GO:0046914,GO:0048519,GO:0048523,GO:0048583,GO:0048585,GO:0050789,GO:0050794,GO:0050896,GO:0051171,GO:0051172,GO:0051252,GO:0051253,GO:0051716,GO:0060255,GO:0065007,GO:0070013,GO:0070410,GO:0070412,GO:0070647,GO:0070848,GO:0070887,GO:0071310,GO:0071363,GO:0071495,GO:0071559,GO:0071560,GO:0071704,GO:0080090,GO:0090092,GO:0090101,GO:0090287,GO:0090304,GO:1901360,GO:1901362,GO:1901576,GO:2000112,GO:2000113,GO:2001141 | BBC |
| comp219942_c0_seq9.p1 | | |
| comp217552_c1_seq2.p1 | | cAMP-specific 3,5-cyclic phosphodiesterase |
| comp215497_c0_seq2.p1 | GO:0005575,GO:0005622,GO:0005623,GO:0005737,GO:0030016,GO:0030017,GO:0030018,GO:0031430,GO:0031672,GO:0031674,GO:0043226,GO:0043228,GO:0043229,GO:0043232,GO:0043292,GO:0044422,GO:0044424,GO:0044444,GO:0044449,GO:0044464,GO:0055120 | Heat shock protein |
| comp211047_c0_seq1.p1 | GO:0000302,GO:0005575,GO:0005622,GO:0005623,GO:0005737,GO:0005739,GO:0005740,GO:0005743,GO:0005753,GO:0006873,GO:0006874,GO:0006875,GO:0006950,GO:0006979,GO:0006996,GO:0007005,GO:0007006,GO:0007007,GO:0007275,GO:0007399,GO:0008150,GO:0009653,GO:0009987,GO:0016020,GO:0016043,GO:0016044,GO:0016469,GO:0019725,GO:0019866,GO:0022008,GO:0030003,GO:0030154,GO:0031090,GO:0031966,GO:0031967,GO:0031975,GO:0032501,GO:0032502,GO:0032989,GO:0032990,GO:0032991,GO:0042221,GO:0042407,GO:0042592,GO:0043226,GO:0043227,GO:0043229,GO:0043231,GO:0043234,GO:0044422,GO:0044424,GO:0044425,GO:0044429,GO:0044444,GO:0044446,GO:0044455,GO:0044464,GO:0044699,GO:0044707,GO:0044763,GO:0044767,GO:0045259,GO:0048731,GO:0048856,GO:0048869,GO:0048878,GO:0050801,GO:0050896,GO:0051560,GO:0055065,GO:0055074,GO:0055080,GO:0055082,GO:0061024,GO:0065007,GO:0065008,GO:0070584,GO:0071840,GO:0072503,GO:0072507,GO:1901700 | inner membrane protein, mitochondrial |

| | | |
|---|---|---|
| comp213330_c0_seq4.p1 | GO:0000003,GO:0000226,GO:0000278,GO:0000280,GO:0000281,GO:0000902,GO:0000904,GO:0000910,GO:0000912,GO:0000915,GO:0000920,GO:0001654,GO:0001669,GO:0001709,GO:0001745,GO:0002009,GO:0002376,GO:0002478,GO:0002495,GO:0002504,GO:0003006,GO:0003674,GO:0004871,GO:0005083,GO:0005096,GO:0005099,GO:0005100,GO:0005488,GO:0005515,GO:0005575,GO:0005622,GO:0005623,GO:0005634,GO:0005737,GO:0005819,GO:0005829,GO:0005856,GO:0006140,GO:0006461,GO:0006810,GO:0006811,GO:0006820,GO:0006897,GO:0006898,GO:0006915,GO:0006928,GO:0006950,GO:0006996,GO:0007010,GO:0007017,GO:0007018,GO:0007049,GO:0007051,GO:0007052,GO:0007067,GO:0007098,GO:0007108,GO:0007109,GO:0007126,GO:0007143,GO:0007154,GO:0007165,GO:0007264,GO:0007265,GO:0007266,GO:0007275,GO:0007276,GO:0007283,GO:0007399,GO:0007405,GO:0007423,GO:0007525,GO:0007548,GO:0007596,GO:0007599,GO:0008017,GO:0008047,GO:0008092,GO:0008104,GO:0008150,GO:0008219,GO:0008272,GO:0008283,GO:0008289,GO:0009118,GO:0009611,GO:0009653,GO:0009790,GO:0009792,GO:0009887,GO:0009888,GO:0009894,GO:0009966,GO:0009968,GO:0009987,GO:0010171,GO:0010646,GO:0010648,GO:0012501,GO:0015630,GO:0015631,GO:0015698,GO:0016023,GO:0016043,GO:0016192,GO:0016265,GO:0016358,GO:0019219,GO:0019220,GO:0019222,GO:0019882,GO:0019884,GO:0019886,GO:0019953,GO:0019992,GO:0022008,GO:0022402,GO:0022414,GO:0022607,GO:0023051,GO:0023052,GO:0023057,GO:0030029,GO:0030030,GO:0030036,GO:0030111,GO:0030141,GO:0030154,GO:0030178,GO:0030182,GO:0030234,GO:0030496,GO:0030695,GO:0030811,GO:0031023,GO:0031032,GO:0031175,GO:0031323,GO:0031329,GO:0031410,GO:0031982,GO:0031988,GO:0032153,GO:0032154,GO:0032155,GO:0032318,GO:0032319,GO:0032320,GO:0032321,GO:0032489,GO:0032501,GO:0032502,GO:0032504,GO:0032506,GO:0032507,GO:0032855,GO:0032879,GO:0032880,GO:0032886,GO:0032989,GO:0032990,GO:0032991,GO:0033036,GO:0033043,GO:0033121,GO:0033124,GO:0033206,GO:0034613,GO:0034622,GO:0035023,GO:0035556,GO:0040011,GO:0040035,GO:0040038,GO:0042060,GO:0043014,GO:0043015,GO:0043085,GO:0043087,GO:0043088,GO:0043089,GO:0043226,GO:0043227,GO:0043228,GO:0043229,GO:0043231,GO:0043232,GO:0043234,GO:0043547,GO:0043623,GO:0043933,GO:0044085,GO:0044093,GO:0044422,GO:0044424,GO:0044430,GO:0044444,GO:0044446,GO:0044464,GO:0044699,GO:0044700,GO:0044702,GO:0044707,GO:0044763,GO:0044765,GO:0044767,GO:0045165,GO:0045185,GO:0046578,GO:0048002,GO:0048232,GO:0048285,GO:0048468,GO:0048471,GO:0048487,GO:0048513,GO:0048519,GO:0048523,GO:0048583,GO:0048585,GO:0048592,GO:0048608,GO:0048609,GO:0048610,GO:0048666,GO:0048667,GO:0048699,GO:0048729,GO:0048731,GO:0048749,GO:0048806,GO:0048812,GO:0048813,GO:0048856,GO:0048858,GO:0048869,GO:0050789,GO:0050790,GO:0050794,GO:0050817,GO:0050878,GO:0050896,GO:0051056,GO:0051128,GO:0051171,GO:0051174,GO:0051179,GO:0051225,GO:0051233,GO:0051234,GO:0051235,GO:0051255,GO:0051256,GO:0051297,GO:0051298,GO:0051301,GO:0051321,GO:0051336,GO:0051345,GO:0051493,GO:0051641,GO:0051651,GO:0051716,GO:0051983,GO:0051988,GO:0060089,GO:0060341,GO:0060429,GO:0060589,GO:0061061,GO:0061351,GO:0061458,GO:0065003,GO:0065007,GO:0065008,GO:0065009,GO:0070271,GO:0070507,GO:0070727,GO:0070925,GO:0071822,GO:0071840,GO:0072089,GO:0072348,GO:0072686,GO:0080090,GO:0090307,GO:0097149,GO:0097223,GO:1900542 | Rac GTPase activating protein 1 |
| comp217871_c0_seq5.p1 | | Ankyrin Repeat Protein |

| | | |
|---|---|---|
| comp215128_c1_seq4.p1 | GO:0000151,GO:0000209,GO:0001501,GO:0002253,GO:0002376,GO:0002429,GO:0002682,GO:0002684,GO:0002757,GO:0002764,GO:0002768,GO:0003674,GO:0003676,GO:0003677,GO:0003690,GO:0003824,GO:0004842,GO:0005488,GO:0005515,GO:0005575,GO:0005622,GO:0005623,GO:0006355,GO:0006357,GO:0006464,GO:0006508,GO:0006511,GO:0007154,GO:0007165,GO:0007166,GO:0007275,GO:0008150,GO:0008152,GO:0009056,GO:0009057,GO:0009653,GO:0009790,GO:0009792,GO:0009887,GO:0009889,GO:0009891,GO:0009893,GO:0009966,GO:0009967,GO:0009987,GO:0010468,GO:0010498,GO:0010556,GO:0010557,GO:0010604,GO:0010627,GO:0010628,GO:0010646,GO:0010647,GO:0010740,GO:0010941,GO:0010942,GO:0016567,GO:0016874,GO:0016879,GO:0016881,GO:0019219,GO:0019222,GO:0019538,GO:0019787,GO:0019941,GO:0023051,GO:0023052,GO:0023056,GO:0030163,GO:0031323,GO:0031325,GO:0031326,GO:0031328,GO:0032088,GO:0032182,GO:0032386,GO:0032388,GO:0032446,GO:0032501,GO:0032502,GO:0032879,GO:0032880,GO:0032991,GO:0033157,GO:0036211,GO:0042306,GO:0042307,GO:0042345,GO:0042346,GO:0042981,GO:0042990,GO:0042993,GO:0043009,GO:0043065,GO:0043067,GO:0043068,GO:0043122,GO:0043123,GO:0043130,GO:0043161,GO:0043170,GO:0043234,GO:0043412,GO:0043433,GO:0043566,GO:0043632,GO:0044092,GO:0044093,GO:0044237,GO:0044238,GO:0044248,GO:0044257,GO:0044260,GO:0044265,GO:0044267,GO:0044424,GO:0044464,GO:0044699,GO:0044700,GO:0044707,GO:0044763,GO:0044767,GO:0045893,GO:0045935,GO:0045944,GO:0046822,GO:0046824,GO:0048513,GO:0048518,GO:0048522,GO:0048562,GO:0048568,GO:0048583,GO:0048584,GO:0048598,GO:0048701,GO:0048704,GO:0048705,GO:0048706,GO:0048731,GO:0048856,GO:0050776,GO:0050778,GO:0050789,GO:0050794,GO:0050851,GO:0050852,GO:0050896,GO:0051049,GO:0051050,GO:0051090,GO:0051091,GO:0051092,GO:0051171,GO:0051173,GO:0051222,GO:0051223,GO:0051252,GO:0051254,GO:0051603,GO:0051716,GO:0060255,GO:0060341,GO:0065007,GO:0065009,GO:0070201,GO:0070647,GO:0071704,GO:0071797,GO:0080090,GO:0090316,GO:0097039,GO:0097159,GO:1900180,GO:1901363,GO:1901575,GO:2000112,GO:2001141 | RanBP-type and C3HC4-type zinc finger containing 1 |
| comp215128_c1_seq4.p2 | GO:0001501,GO:0007275,GO:0008150,GO:0009653,GO:0009790,GO:0009792,GO:0009887,GO:0032501,GO:0032502,GO:0043009,GO:0044699,GO:0044707,GO:0044767,GO:0048513,GO:0048562,GO:0048568,GO:0048598,GO:0048701,GO:0048704,GO:0048705,GO:0048706,GO:0048731,GO:0048856 | RanBP-type and C3HC4-type zinc finger containing 1 |
| comp209705_c0_seq2.p1 | | Glucosaminyl (N-acetyl) transferase |
| comp221240_c1_seq6.p1 | GO:0000381,GO:0003674,GO:0003676,GO:0003723,GO:0003729,GO:0005488,GO:0005575,GO:0005622,GO:0005623,GO:0005634,GO:0005654,GO:0005737,GO:0006915,GO:0008150,GO:0008219,GO:0009987,GO:0010468,GO:0012501,GO:0016265,GO:0016604,GO:0016607,GO:0019219,GO:0019222,GO:0031323,GO:0031974,GO:0031981,GO:0043226,GO:0043227,GO:0043229,GO:0043231,GO:0043233,GO:0043484,GO:0044422,GO:0044424,GO:0044428,GO:0044446,GO:0044451,GO:0044464,GO:0044699,GO:0044763,GO:0048024,GO:0050684,GO:0050789,GO:0050794,GO:0051171,GO:0051252,GO:0060255,GO:0065007,GO:0070013,GO:0080090,GO:0097159,GO:1901363 | RNA binding motif protein 25 |
| comp221240_c1_seq6.p2 | GO:0003674,GO:0005488,GO:0005515,GO:0005575,GO:0005622,GO:0005623,GO:0005634,GO:0043226,GO:0043227,GO:0043229,GO:0043231,GO:0044424,GO:0044464 | HOX |

| | | |
|---|---|---|
| comp189162_c0_seq1.p1 | GO:0001071,GO:0001525,GO:0001568,GO:0001944,GO:0003674,GO:0003676,GO:0003677,GO:0003700,GO:0005488,GO:0005575,GO:0005622,GO:0005623,GO:0005634,GO:0005654,GO:0005737,GO:0005794,GO:0006139,GO:0006351,GO:0006355,GO:0006366,GO:0006725,GO:0006807,GO:0007275,GO:0007517,GO:0007519,GO:0008150,GO:0008152,GO:0008284,GO:0009058,GO:0009059,GO:0009653,GO:0009719,GO:0009888,GO:0009889,GO:0009891,GO:0009893,GO:0009987,GO:0010033,GO:0010243,GO:0010467,GO:0010468,GO:0010556,GO:0010557,GO:0010604,GO:0010628,GO:0014070,GO:0014706,GO:0016043,GO:0016070,GO:0018130,GO:0019219,GO:0019222,GO:0019438,GO:0022607,GO:0030030,GO:0030031,GO:0030033,GO:0030154,GO:0031323,GO:0031325,GO:0031326,GO:0031328,GO:0031974,GO:0031981,GO:0032501,GO:0032502,GO:0032528,GO:0032774,GO:0034641,GO:0034645,GO:0034654,GO:0035914,GO:0042127,GO:0042221,GO:0042692,GO:0043170,GO:0043226,GO:0043227,GO:0043229,GO:0043231,GO:0043233,GO:0044085,GO:0044237,GO:0044238,GO:0044249,GO:0044260,GO:0044271,GO:0044422,GO:0044424,GO:0044428,GO:0044444,GO:0044446,GO:0044464,GO:0044699,GO:0044707,GO:0044763,GO:0044767,GO:0045893,GO:0045935,GO:0046483,GO:0048513,GO:0048514,GO:0048518,GO:0048522,GO:0048646,GO:0048731,GO:0048856,GO:0048869,GO:0050789,GO:0050794,GO:0050896,GO:0051146,GO:0051171,GO:0051173,GO:0051252,GO:0051254,GO:0051716,GO:0060255,GO:0060537,GO:0060538,GO:0061061,GO:0065007,GO:0070013,GO:0070887,GO:0071310,GO:0071407,GO:0071417,GO:0071495,GO:0071704,GO:0071840,GO:0072358,GO:0072359,GO:0080090,GO:0090304,GO:0097159,GO:1901360,GO:1901362,GO:1901363,GO:1901576,GO:1901652,GO:1901653,GO:1901698,GO:1901699,GO:1901700,GO:1901701,GO:2000112,GO:2001141 | Kruppel-like factor 5 (intestinal) |

| | | |
|---|---|---|
| comp209552_c0_seq16.p1 | GO:0000003,GO:0000375,GO:0000377,GO:0000398,GO:0000785,GO:0000803,GO:0000805,GO:0000902,GO:0000904,GO:0003006,GO:0003674,GO:0003676,GO:0003678,GO:0003682,GO:0003723,GO:0003724,GO:0003725,GO:0003824,GO:0004003,GO:0004004,GO:0004386,GO:0005488,GO:0005575,GO:0005622,GO:0005623,GO:0005634,GO:0005654,GO:0005694,GO:0005700,GO:0005730,GO:0005737,GO:0005813,GO:0005815,GO:0005856,GO:0006139,GO:0006152,GO:0006163,GO:0006195,GO:0006200,GO:0006259,GO:0006355,GO:0006357,GO:0006396,GO:0006397,GO:0006725,GO:0006753,GO:0006793,GO:0006796,GO:0006807,GO:0006950,GO:0007275,GO:0007276,GO:0007281,GO:0007399,GO:0007409,GO:0007549,GO:0007568,GO:0007610,GO:0007617,GO:0007618,GO:0007619,GO:0007623,GO:0008026,GO:0008049,GO:0008094,GO:0008150,GO:0008152,GO:0008186,GO:0008340,GO:0008380,GO:0009047,GO:0009056,GO:0009116,GO:0009117,GO:0009119,GO:0009123,GO:0009125,GO:0009126,GO:0009128,GO:0009141,GO:0009143,GO:0009144,GO:0009146,GO:0009150,GO:0009154,GO:0009158,GO:0009161,GO:0009164,GO:0009166,GO:0009167,GO:0009169,GO:0009199,GO:0009203,GO:0009205,GO:0009207,GO:0009259,GO:0009261,GO:0009266,GO:0009408,GO:0009628,GO:0009653,GO:0009790,GO:0009792,GO:0009889,GO:0009891,GO:0009892,GO:0009893,GO:0009987,GO:0010259,GO:0010467,GO:0010468,GO:0010556,GO:0010557,GO:0010604,GO:0010605,GO:0010608,GO:0010628,GO:0010629,GO:0015630,GO:0016043,GO:0016049,GO:0016070,GO:0016071,GO:0016456,GO:0016457,GO:0016462,GO:0016545,GO:0016787,GO:0016817,GO:0016818,GO:0016887,GO:0017111,GO:0019098,GO:0019219,GO:0019222,GO:0019439,GO:0019637,GO:0019693,GO:0019953,GO:0022008,GO:0022412,GO:0022414,GO:0022607,GO:0030030,GO:0030154,GO:0030182,GO:0030529,GO:0031175,GO:0031323,GO:0031325,GO:0031326,GO:0031328,GO:0031974,GO:0031981,GO:0032392,GO:0032501,GO:0032502,GO:0032504,GO:0032508,GO:0032989,GO:0032990,GO:0032991,GO:0033057,GO:0033554,GO:0034605,GO:0034622,GO:0034641,GO:0034655,GO:0040007,GO:0040029,GO:0042278,GO:0042454,GO:0042623,GO:0042714,GO:0043170,GO:0043226,GO:0043227,GO:0043228,GO:0043229,GO:0043231,GO:0043232,GO:0043233,GO:0043234,GO:0043487,GO:0043488,GO:0043489,GO:0043933,GO:0044085,GO:0044237,GO:0044238,GO:0044248,GO:0044260,GO:0044270,GO:0044281,GO:0044422,GO:0044424,GO:0044427,GO:0044428,GO:0044430,GO:0044444,GO:0044446,GO:0044464,GO:0044699,GO:0044702,GO:0044703,GO:0044706,GO:0044707,GO:0044710,GO:0044763,GO:0044767,GO:0045433,GO:0045814,GO:0045893,GO:0045935,GO:0045944,GO:0046034,GO:0046128,GO:0046130,GO:0046434,GO:0046483,GO:0046536,GO:0046700,GO:0048065,GO:0048255,GO:0048468,GO:0048511,GO:0048518,GO:0048519,GO:0048522,GO:0048588,GO:0048589,GO:0048609,GO:0048610,GO:0048666,GO:0048667,GO:0048675,GO:0048699,GO:0048731,GO:0048812,GO:0048856,GO:0048858,GO:0048869,GO:0050789,GO:0050794,GO:0050896,GO:0051171,GO:0051173,GO:0051252,GO:0051254,GO:0051704,GO:0051705,GO:0051716,GO:0055086,GO:0060179,GO:0060255,GO:0060560,GO:0061564,GO:0065003,GO:0065004,GO:0065007,GO:0065008,GO:0070013,GO:0070035,GO:0070934,GO:0070937,GO:0071103,GO:0071704,GO:0071824,GO:0071840,GO:0072521,GO:0072523,GO:0080090,GO:0090304,GO:0097159,GO:1901135,GO:1901136,GO:1901292,GO:1901360,GO:1901361,GO:1901363,GO:1901564,GO:1901565,GO:1901575,GO:1901657,GO:1901658,GO:1990138,GO:2000112,GO:2001141 | ATPdependent RNA helicase |

| | | |
|---|---|---|
| comp209552_c0_seq16.p2 | GO:0000003,GO:0000375,GO:0000377,GO:0000398,GO:0000785,GO:0000803,GO:0000805,GO:0000902,GO:0000904,GO:0003006,GO:0003674,GO:0003676,GO:0003678,GO:0003682,GO:0003723,GO:0003724,GO:0003725,GO:0003824,GO:0004003,GO:0004004,GO:0004386,GO:0005488,GO:0005575,GO:0005622,GO:0005623,GO:0005634,GO:0005654,GO:0005694,GO:0005700,GO:0005730,GO:0005737,GO:0005813,GO:0005815,GO:0005856,GO:0006139,GO:0006152,GO:0006163,GO:0006195,GO:0006200,GO:0006259,GO:0006355,GO:0006357,GO:0006396,GO:0006397,GO:0006725,GO:0006753,GO:0006793,GO:0006796,GO:0006807,GO:0006950,GO:0007275,GO:0007276,GO:0007281,GO:0007399,GO:0007409,GO:0007549,GO:0007568,GO:0007610,GO:0007617,GO:0007618,GO:0007619,GO:0007623,GO:0008026,GO:0008049,GO:0008094,GO:0008150,GO:0008152,GO:0008186,GO:0008340,GO:0008380,GO:0009047,GO:0009056,GO:0009116,GO:0009117,GO:0009119,GO:0009123,GO:0009125,GO:0009126,GO:0009128,GO:0009141,GO:0009143,GO:0009144,GO:0009146,GO:0009150,GO:0009154,GO:0009158,GO:0009161,GO:0009164,GO:0009166,GO:0009167,GO:0009169,GO:0009199,GO:0009203,GO:0009205,GO:0009207,GO:0009259,GO:0009261,GO:0009266,GO:0009408,GO:0009628,GO:0009653,GO:0009790,GO:0009792,GO:0009889,GO:0009891,GO:0009892,GO:0009893,GO:0009987,GO:0010259,GO:0010467,GO:0010468,GO:0010556,GO:0010557,GO:0010604,GO:0010605,GO:0010608,GO:0010628,GO:0010629,GO:0015630,GO:0016043,GO:0016049,GO:0016070,GO:0016071,GO:0016456,GO:0016457,GO:0016462,GO:0016545,GO:0016787,GO:0016817,GO:0016818,GO:0016887,GO:0017111,GO:0019098,GO:0019219,GO:0019222,GO:0019439,GO:0019637,GO:0019693,GO:0019953,GO:0022008,GO:0022412,GO:0022414,GO:0022607,GO:0030030,GO:0030154,GO:0030182,GO:0030529,GO:0031175,GO:0031323,GO:0031325,GO:0031326,GO:0031328,GO:0031974,GO:0031981,GO:0032392,GO:0032501,GO:0032502,GO:0032504,GO:0032508,GO:0032989,GO:0032990,GO:0032991,GO:0033057,GO:0033554,GO:0034605,GO:0034622,GO:0034641,GO:0034655,GO:0040007,GO:0040029,GO:0042278,GO:0042454,GO:0042623,GO:0042714,GO:0043170,GO:0043226,GO:0043227,GO:0043228,GO:0043229,GO:0043231,GO:0043232,GO:0043233,GO:0043234,GO:0043487,GO:0043488,GO:0043489,GO:0043933,GO:0044085,GO:0044237,GO:0044238,GO:0044248,GO:0044260,GO:0044270,GO:0044281,GO:0044422,GO:0044424,GO:0044427,GO:0044428,GO:0044430,GO:0044444,GO:0044446,GO:0044464,GO:0044699,GO:0044702,GO:0044703,GO:0044706,GO:0044707,GO:0044710,GO:0044763,GO:0044767,GO:0045433,GO:0045814,GO:0045893,GO:0045935,GO:0045944,GO:0046034,GO:0046128,GO:0046130,GO:0046434,GO:0046483,GO:0046536,GO:0046700,GO:0048065,GO:0048255,GO:0048468,GO:0048511,GO:0048518,GO:0048519,GO:0048522,GO:0048588,GO:0048589,GO:0048609,GO:0048610,GO:0048666,GO:0048667,GO:0048675,GO:0048699,GO:0048731,GO:0048812,GO:0048856,GO:0048858,GO:0048869,GO:0050789,GO:0050794,GO:0050896,GO:0051171,GO:0051173,GO:0051252,GO:0051254,GO:0051704,GO:0051705,GO:0051716,GO:0055086,GO:0060179,GO:0060255,GO:0060560,GO:0061564,GO:0065003,GO:0065004,GO:0065007,GO:0065008,GO:0070013,GO:0070035,GO:0070934,GO:0070937,GO:0071103,GO:0071704,GO:0071824,GO:0071840,GO:0072521,GO:0072523,GO:0080090,GO:0090304,GO:0097159,GO:1901135,GO:1901136,GO:1901292,GO:1901360,GO:1901361,GO:1901363,GO:1901564,GO:1901565,GO:1901575,GO:1901657,GO:1901658,GO:1990138,GO:2000112,GO:2001141 | DEAH (Asp-Glu-Ala-His) box polypeptide 9 |

| | | |
|---|---|---|
| comp209552_c0_seq16.p3 | GO:0000003,GO:0000375,GO:0000377,GO:0000398,GO:0000785,GO:0000803,GO:0000805,GO:0000902,GO:0000904,GO:0003006,GO:0003674,GO:0003676,GO:0003678,GO:0003682,GO:0003723,GO:0003724,GO:0003725,GO:0003824,GO:0004003,GO:0004004,GO:0004386,GO:0005488,GO:0005575,GO:0005622,GO:0005623,GO:0005634,GO:0005654,GO:0005694,GO:0005700,GO:0005730,GO:0005737,GO:0005813,GO:0005815,GO:0005856,GO:0006139,GO:0006152,GO:0006163,GO:0006195,GO:0006200,GO:0006259,GO:0006355,GO:0006357,GO:0006396,GO:0006397,GO:0006725,GO:0006753,GO:0006793,GO:0006796,GO:0006807,GO:0006950,GO:0007275,GO:0007276,GO:0007281,GO:0007399,GO:0007409,GO:0007549,GO:0007568,GO:0007610,GO:0007617,GO:0007618,GO:0007619,GO:0007623,GO:0008026,GO:0008049,GO:0008094,GO:0008150,GO:0008152,GO:0008186,GO:0008340,GO:0008380,GO:0009047,GO:0009056,GO:0009116,GO:0009117,GO:0009119,GO:0009123,GO:0009125,GO:0009126,GO:0009128,GO:0009141,GO:0009143,GO:0009144,GO:0009146,GO:0009150,GO:0009154,GO:0009158,GO:0009161,GO:0009164,GO:0009166,GO:0009167,GO:0009169,GO:0009199,GO:0009203,GO:0009205,GO:0009207,GO:0009259,GO:0009261,GO:0009266,GO:0009408,GO:0009628,GO:0009653,GO:0009790,GO:0009792,GO:0009889,GO:0009891,GO:0009892,GO:0009893,GO:0009987,GO:0010259,GO:0010467,GO:0010468,GO:0010556,GO:0010557,GO:0010604,GO:0010605,GO:0010608,GO:0010628,GO:0010629,GO:0015630,GO:0016043,GO:0016049,GO:0016070,GO:0016071,GO:0016456,GO:0016457,GO:0016462,GO:0016545,GO:0016787,GO:0016817,GO:0016818,GO:0016887,GO:0017111,GO:0019098,GO:0019219,GO:0019222,GO:0019439,GO:0019637,GO:0019693,GO:0019953,GO:0022008,GO:0022412,GO:0022414,GO:0022607,GO:0030030,GO:0030154,GO:0030182,GO:0030529,GO:0031175,GO:0031323,GO:0031325,GO:0031326,GO:0031328,GO:0031974,GO:0031981,GO:0032392,GO:0032501,GO:0032502,GO:0032504,GO:0032508,GO:0032989,GO:0032990,GO:0032991,GO:0033057,GO:0033554,GO:0034605,GO:0034622,GO:0034641,GO:0034655,GO:0040007,GO:0040029,GO:0042278,GO:0042454,GO:0042623,GO:0042714,GO:0043170,GO:0043226,GO:0043227,GO:0043228,GO:0043229,GO:0043231,GO:0043232,GO:0043233,GO:0043234,GO:0043487,GO:0043488,GO:0043489,GO:0043933,GO:0044085,GO:0044237,GO:0044238,GO:0044248,GO:0044260,GO:0044270,GO:0044281,GO:0044422,GO:0044424,GO:0044427,GO:0044428,GO:0044430,GO:0044444,GO:0044446,GO:0044464,GO:0044699,GO:0044702,GO:0044703,GO:0044706,GO:0044707,GO:0044710,GO:0044763,GO:0044767,GO:0045433,GO:0045814,GO:0045893,GO:0045935,GO:0045944,GO:0046034,GO:0046128,GO:0046130,GO:0046434,GO:0046483,GO:0046536,GO:0046700,GO:0048065,GO:0048255,GO:0048468,GO:0048511,GO:0048518,GO:0048519,GO:0048522,GO:0048588,GO:0048589,GO:0048609,GO:0048610,GO:0048666,GO:0048667,GO:0048675,GO:0048699,GO:0048731,GO:0048812,GO:0048856,GO:0048858,GO:0048869,GO:0050789,GO:0050794,GO:0050896,GO:0051171,GO:0051173,GO:0051252,GO:0051254,GO:0051704,GO:0051705,GO:0051716,GO:0055086,GO:0060179,GO:0060255,GO:0060560,GO:0061564,GO:0065003,GO:0065004,GO:0065007,GO:0065008,GO:0070013,GO:0070035,GO:0070934,GO:0070937,GO:0071103,GO:0071704,GO:0071824,GO:0071840,GO:0072521,GO:0072523,GO:0080090,GO:0090304,GO:0097159,GO:1901135,GO:1901136,GO:1901292,GO:1901360,GO:1901361,GO:1901363,GO:1901564,GO:1901565,GO:1901575,GO:1901657,GO:1901658,GO:1990138,GO:2000112,GO:2001141 | DEAH (Asp-Glu-Ala-His) box polypeptide 9 |
| comp220198_c4_seq14.p1 | | Pfam:DUF23 |
| comp220198_c4_seq14.p2 | | Reverse transcriptase (RNA-dependent DNA polymerase) |
| comp219827_c0_seq2.p1 | GO:0005575,GO:0005622,GO:0005623,GO:0005737,GO:0005794,GO:0043226,GO:0043227,GO:0043229,GO:0043231,GO:0044424,GO:0044444,GO:0044464 | janus kinase and microtubule interacting protein 3 |

| | | |
|---|---|---|
| comp218338_c1_seq6.p1 | GO:0000003,GO:0003006,GO:0003674,GO:0003676,GO:0003677,GO:0003682,GO:0005488,GO:0005575,GO:0005622,GO:0005623,GO:0005634,GO:0006139,GO:0006351,GO:0006725,GO:0006807,GO:0006996,GO:0007275,GO:0007276,GO:0007281,GO:0007292,GO:0007293,GO:0007548,GO:0008150,GO:0008152,GO:0008406,GO:0008584,GO:0009058,GO:0009059,GO:0009653,GO:0009987,GO:0010467,GO:0010468,GO:0016043,GO:0016070,GO:0018130,GO:0019222,GO:0019438,GO:0019827,GO:0019953,GO:0022412,GO:0022414,GO:0030154,GO:0030707,GO:0030718,GO:0030723,GO:0031490,GO:0032501,GO:0032502,GO:0032504,GO:0032774,GO:0034641,GO:0034645,GO:0034654,GO:0036098,GO:0043170,GO:0043226,GO:0043227,GO:0043229,GO:0043231,GO:0043566,GO:0044237,GO:0044238,GO:0044249,GO:0044260,GO:0044271,GO:0044424,GO:0044464,GO:0044699,GO:0044702,GO:0044707,GO:0044763,GO:0044767,GO:0045137,GO:0045478,GO:0045595,GO:0045596,GO:0046483,GO:0046546,GO:0046661,GO:0048468,GO:0048477,GO:0048513,GO:0048519,GO:0048523,GO:0048608,GO:0048609,GO:0048610,GO:0048646,GO:0048731,GO:0048856,GO:0048863,GO:0048864,GO:0048869,GO:0050789,GO:0050793,GO:0050794,GO:0051093,GO:0060255,GO:0061458,GO:0065007,GO:0071704,GO:0071840,GO:0090304,GO:0097159,GO:1901360,GO:1901362,GO:1901363,GO:1901576 | PWP1 homolog (S. cerevisiae) |
| comp218338_c1_seq6.p2 | | KR domain |
| comp213767_c1_seq2.p1 | GO:0000166,GO:0001882,GO:0001883,GO:0003674,GO:0003824,GO:0004103,GO:0004104,GO:0004305,GO:0005488,GO:0005524,GO:0005575,GO:0005622,GO:0005623,GO:0005737,GO:0006066,GO:0006576,GO:0006629,GO:0006644,GO:0006646,GO:0006650,GO:0006656,GO:0006657,GO:0006793,GO:0006796,GO:0006807,GO:0008144,GO:0008150,GO:0008152,GO:0008610,GO:0008654,GO:0009058,GO:0009308,GO:0009987,GO:0016301,GO:0016310,GO:0016740,GO:0016772,GO:0016773,GO:0016787,GO:0016788,GO:0017076,GO:0019637,GO:0019695,GO:0030554,GO:0032549,GO:0032550,GO:0032553,GO:0032555,GO:0032559,GO:0033265,GO:0034641,GO:0035639,GO:0036094,GO:0042439,GO:0043167,GO:0043168,GO:0043169,GO:0043178,GO:0044106,GO:0044237,GO:0044238,GO:0044249,GO:0044255,GO:0044281,GO:0044283,GO:0044424,GO:0044464,GO:0044710,GO:0044711,GO:0045017,GO:0046165,GO:0046337,GO:0046470,GO:0046474,GO:0046486,GO:0050997,GO:0052689,GO:0071704,GO:0090407,GO:0097159,GO:1901265,GO:1901363,GO:1901564,GO:1901566,GO:1901576,GO:1901615,GO:1901617 | choline kinase alpha |
| comp213767_c1_seq2.p2 | GO:0000166,GO:0001882,GO:0001883,GO:0003674,GO:0003824,GO:0004103,GO:0004104,GO:0004305,GO:0005488,GO:0005524,GO:0005575,GO:0005622,GO:0005623,GO:0005737,GO:0006066,GO:0006576,GO:0006629,GO:0006644,GO:0006646,GO:0006650,GO:0006656,GO:0006657,GO:0006793,GO:0006796,GO:0006807,GO:0008144,GO:0008150,GO:0008152,GO:0008610,GO:0008654,GO:0009058,GO:0009308,GO:0009987,GO:0016301,GO:0016310,GO:0016740,GO:0016772,GO:0016773,GO:0016787,GO:0016788,GO:0017076,GO:0019637,GO:0019695,GO:0030554,GO:0032549,GO:0032550,GO:0032553,GO:0032555,GO:0032559,GO:0033265,GO:0034641,GO:0035639,GO:0036094,GO:0042439,GO:0043167,GO:0043168,GO:0043169,GO:0043178,GO:0044106,GO:0044237,GO:0044238,GO:0044249,GO:0044255,GO:0044281,GO:0044283,GO:0044424,GO:0044464,GO:0044710,GO:0044711,GO:0045017,GO:0046165,GO:0046337,GO:0046470,GO:0046474,GO:0046486,GO:0050997,GO:0052689,GO:0071704,GO:0090407,GO:0097159,GO:1901265,GO:1901363,GO:1901564,GO:1901566,GO:1901576,GO:1901615,GO:1901617 | choline kinase alpha |
| comp218568_c1_seq17.p1 | GO:0003674,GO:0003824,GO:0006793,GO:0006796,GO:0008150,GO:0008152,GO:0009987,GO:0016301,GO:0016310,GO:0016740,GO:0016772,GO:0016773,GO:0044237,GO:0050201 | fucokinase |
| comp218243_c1_seq17.p1 | | membrane-associated ring finger (C3HC4) 7, E3 ubiquitin protein ligase |
| comp207312_c1_seq13.p1 | | LIM domain |

| | | |
|---|---|---|
| comp219607_c0_seq1.p1 | GO:0003008,GO:0003674,GO:0005070,GO:0005488,GO:0005515,GO:0005575,GO:0005622,GO:0005623,GO:0005737,GO:0005768,GO:0007154,GO:0007267,GO:0007268,GO:0007274,GO:0008064,GO:0008150,GO:0008582,GO:0009966,GO:0009967,GO:0009987,GO:0010646,GO:0010647,GO:0016043,GO:0016202,GO:0019226,GO:0023051,GO:0023052,GO:0023056,GO:0030674,GO:0030832,GO:0030833,GO:0032271,GO:0032501,GO:0032535,GO:0032956,GO:0032970,GO:0033043,GO:0035591,GO:0035637,GO:0040008,GO:0043226,GO:0043227,GO:0043229,GO:0043231,GO:0043254,GO:0044087,GO:0044424,GO:0044444,GO:0044464,GO:0044699,GO:0044700,GO:0044707,GO:0044763,GO:0045595,GO:0045886,GO:0045926,GO:0048518,GO:0048519,GO:0048522,GO:0048523,GO:0048583,GO:0048584,GO:0048634,GO:0048638,GO:0048640,GO:0048641,GO:0048742,GO:0050789,GO:0050793,GO:0050794,GO:0050803,GO:0050807,GO:0050877,GO:0051093,GO:0051128,GO:0051129,GO:0051147,GO:0051153,GO:0051239,GO:0051493,GO:0051960,GO:0051961,GO:0051963,GO:0051964,GO:0055037,GO:0060090,GO:0060284,GO:0065007,GO:0065008,GO:0071840,GO:0090066,GO:1901861,GO:2000026 | FCH and double SH3 domains |
| comp211796_c0_seq2.p1 | | LATS, large tumor suppressor, homolog 2 (Drosophila) |
| comp216002_c0_seq2.p1 | | Zinc finger protein |
| comp209845_c0_seq2.p1 | GO:0000122,GO:0005575,GO:0005622,GO:0005623,GO:0005634,GO:0005737,GO:0006355,GO:0006357,GO:0008150,GO:0009889,GO:0009890,GO:0009891,GO:0009892,GO:0009893,GO:0010468,GO:0010556,GO:0010557,GO:0010558,GO:0010604,GO:0010605,GO:0010628,GO:0010629,GO:0016020,GO:0019219,GO:0019222,GO:0031323,GO:0031324,GO:0031325,GO:0031326,GO:0031327,GO:0031328,GO:0043226,GO:0043227,GO:0043229,GO:0043231,GO:0044424,GO:0044444,GO:0044464,GO:0045892,GO:0045893,GO:0045934,GO:0045935,GO:0045944,GO:0048471,GO:0048518,GO:0048519,GO:0048522,GO:0048523,GO:0050789,GO:0050794,GO:0051171,GO:0051172,GO:0051173,GO:0051252,GO:0051253,GO:0051254,GO:0060255,GO:0065007,GO:0080090,GO:2000112,GO:2000113,GO:2001141 | motile sperm domain containing 1 |
| comp220104_c0_seq9.p1 | GO:0000003,GO:0008150 | Pfam:efhand_like |
| comp218926_c0_seq3.p1 | GO:0001700,GO:0002009,GO:0003674,GO:0003824,GO:0005488,GO:0005515,GO:0005575,GO:0005622,GO:0005623,GO:0005856,GO:0006139,GO:0006152,GO:0006163,GO:0006195,GO:0006200,GO:0006725,GO:0006753,GO:0006793,GO:0006796,GO:0006807,GO:0006810,GO:0006886,GO:0007275,GO:0007391,GO:0008104,GO:0008150,GO:0008152,GO:0009056,GO:0009116,GO:0009117,GO:0009119,GO:0009123,GO:0009125,GO:0009126,GO:0009128,GO:0009141,GO:0009143,GO:0009144,GO:0009146,GO:0009150,GO:0009154,GO:0009158,GO:0009161,GO:0009164,GO:0009166,GO:0009167,GO:0009169,GO:0009199,GO:0009203,GO:0009205,GO:0009207,GO:0009259,GO:0009261,GO:0009653,GO:0009790,GO:0009792,GO:0009888,GO:0009987,GO:0015031,GO:0015629,GO:0016043,GO:0016331,GO:0016459,GO:0016461,GO:0016462,GO:0016787,GO:0016817,GO:0016818,GO:0016887,GO:0017111,GO:0019439,GO:0019637,GO:0019693,GO:0022607,GO:0030030,GO:0030031,GO:0030175,GO:0032501,GO:0032502,GO:0032991,GO:0033036,GO:0034613,GO:0034641,GO:0034655,GO:0042278,GO:0042454,GO:0042623,GO:0042995,GO:0043025,GO:0043226,GO:0043228,GO:0043229,GO:0043232,GO:0043234,GO:0044085,GO:0044237,GO:0044238,GO:0044248,GO:0044270,GO:0044281,GO:0044297,GO:0044422,GO:0044424,GO:0044430,GO:0044446,GO:0044464,GO:0044699,GO:0044707,GO:0044710,GO:0044763,GO:0044767,GO:0045184,GO:0046034,GO:0046128,GO:0046130,GO:0046434,GO:0046483,GO:0046700,GO:0046847,GO:0046907,GO:0048598,GO:0048729,GO:0048856,GO:0051179,GO:0051234,GO:0051641,GO:0051649,GO:0055086,GO:0060429,GO:0070727,GO:0071702,GO:0071704,GO:0071840,GO:0072521,GO:0072523,GO:0097458,GO:1901135,GO:1901136,GO:1901292,GO:1901360,GO:1901361,GO:1901564,GO:1901565,GO:1901575,GO:1901657,GO:1901658 | myosin xv |

| | | |
|---|---|---|
| comp220750_c2_seq1.p1 | GO:0001520,GO:0005575,GO:0005622,GO:0005623,GO:0005737,GO:0005930,GO:0006996,GO:0007010,GO:0008150,GO:0009987,GO:0016043,GO:0042995,GO:0043226,GO:0043227,GO:0043229,GO:0043231,GO:0044422,GO:0044424,GO:0044447,GO:0044463,GO:0044464,GO:0044699,GO:0044763,GO:0071840 | polyamine modulated factor 1 binding protein 1 |
| comp220750_c2_seq1.p2 | | |
| comp213467_c0_seq1.p1 | | guanylate cyclase |
| comp213467_c0_seq1.p2 | GO:0003674,GO:0003824,GO:0005488,GO:0005515,GO:0005575,GO:0005622,GO:0005623,GO:0005737,GO:0005739,GO:0005811,GO:0005856,GO:0005875,GO:0006139,GO:0006152,GO:0006163,GO:0006195,GO:0006200,GO:0006457,GO:0006725,GO:0006753,GO:0006793,GO:0006796,GO:0006807,GO:0006950,GO:0008150,GO:0008152,GO:0009056,GO:0009116,GO:0009117,GO:0009119,GO:0009123,GO:0009125,GO:0009126,GO:0009128,GO:0009141,GO:0009143,GO:0009144,GO:0009146,GO:0009150,GO:0009154,GO:0009158,GO:0009161,GO:0009164,GO:0009166,GO:0009167,GO:0009169,GO:0009199,GO:0009203,GO:0009205,GO:0009207,GO:0009259,GO:0009261,GO:0009266,GO:0009408,GO:0009628,GO:0009987,GO:0015630,GO:0016462,GO:0016787,GO:0016817,GO:0016818,GO:0016887,GO:0017111,GO:0019439,GO:0019538,GO:0019637,GO:0019693,GO:0032991,GO:0034641,GO:0034655,GO:0042278,GO:0042454,GO:0043170,GO:0043226,GO:0043227,GO:0043228,GO:0043229,GO:0043231,GO:0043232,GO:0043234,GO:0044237,GO:0044238,GO:0044248,GO:0044260,GO:0044267,GO:0044270,GO:0044281,GO:0044422,GO:0044424,GO:0044430,GO:0044444,GO:0044446,GO:0044464,GO:0044710,GO:0046034,GO:0046128,GO:0046130,GO:0046434,GO:0046483,GO:0046700,GO:0050896,GO:0051082,GO:0055086,GO:0071704,GO:0072521,GO:0072523,GO:1901135,GO:1901136,GO:1901292,GO:1901360,GO:1901361,GO:1901564,GO:1901565,GO:1901575,GO:1901657,GO:1901658 | heat shock 70 kDa protein cognate |
| comp216653_c0_seq4.p1 | GO:0003674,GO:0005488,GO:0005515,GO:0005575,GO:0005622,GO:0005623,GO:0005634,GO:0005681,GO:0005737,GO:0007275,GO:0007399,GO:0008150,GO:0009987,GO:0010941,GO:0017134,GO:0019838,GO:0022008,GO:0030154,GO:0030529,GO:0032501,GO:0032502,GO:0032991,GO:0042981,GO:0043066,GO:0043067,GO:0043069,GO:0043226,GO:0043227,GO:0043229,GO:0043231,GO:0044422,GO:0044424,GO:0044428,GO:0044446,GO:0044464,GO:0044699,GO:0044707,GO:0044763,GO:0048519,GO:0048523,GO:0048731,GO:0048856,GO:0048869,GO:0050789,GO:0050794,GO:0060548,GO:0065007 | Apoptosis inhibitor 5 |
| comp216221_c0_seq7.p1 | GO:0000375,GO:0000377,GO:0000398,GO:0003674,GO:0003676,GO:0003723,GO:0003729,GO:0005488,GO:0005575,GO:0005622,GO:0005623,GO:0005634,GO:0005654,GO:0005847,GO:0005848,GO:0005849,GO:0006139,GO:0006351,GO:0006353,GO:0006366,GO:0006369,GO:0006378,GO:0006379,GO:0006396,GO:0006397,GO:0006725,GO:0006807,GO:0007275,GO:0007399,GO:0008150,GO:0008152,GO:0008380,GO:0009058,GO:0009059,GO:0009987,GO:0010467,GO:0016070,GO:0016071,GO:0016604,GO:0018130,GO:0019438,GO:0022008,GO:0030154,GO:0031123,GO:0031124,GO:0031974,GO:0031981,GO:0032501,GO:0032502,GO:0032774,GO:0032991,GO:0034641,GO:0034645,GO:0034654,GO:0043170,GO:0043226,GO:0043227,GO:0043229,GO:0043231,GO:0043233,GO:0043234,GO:0043631,GO:0044237,GO:0044238,GO:0044249,GO:0044260,GO:0044271,GO:0044422,GO:0044424,GO:0044428,GO:0044446,GO:0044451,GO:0044464,GO:0044699,GO:0044707,GO:0044763,GO:0046483,GO:0048731,GO:0048856,GO:0048869,GO:0070013,GO:0071704,GO:0071920,GO:0090304,GO:0090305,GO:0097159,GO:1901360,GO:1901362,GO:1901363,GO:1901576 | Cleavage stimulation factor, 3' pre-RNA subunit 2 |

| | | |
|---|---|---|
| comp210207_c2_seq7.p1 | GO:0000123,GO:0000988,GO:0000989,GO:0001071,GO:0003674,GO:0003682,GO:0003700,GO:0003712,GO:0003713,GO:0005488,GO:0005575,GO:0005622,GO:0005623,GO:0005634,GO:0005654,GO:0006325,GO:0006338,GO:0006355,GO:0006464,GO:0006473,GO:0006475,GO:0006479,GO:0006996,GO:0008150,GO:0008152,GO:0008213,GO:0009889,GO:0009891,GO:0009893,GO:0009987,GO:0010468,GO:0010556,GO:0010557,GO:0010604,GO:0010628,GO:0016043,GO:0016568,GO:0016569,GO:0016570,GO:0016571,GO:0016573,GO:0018193,GO:0018205,GO:0018393,GO:0018394,GO:0019219,GO:0019222,GO:0019538,GO:0031323,GO:0031325,GO:0031326,GO:0031328,GO:0031974,GO:0031981,GO:0032259,GO:0032991,GO:0034708,GO:0034968,GO:0035097,GO:0036211,GO:0040008,GO:0043170,GO:0043226,GO:0043227,GO:0043229,GO:0043231,GO:0043233,GO:0043234,GO:0043412,GO:0043414,GO:0043543,GO:0043966,GO:0043967,GO:0044237,GO:0044238,GO:0044260,GO:0044267,GO:0044422,GO:0044424,GO:0044428,GO:0044446,GO:0044451,GO:0044464,GO:0044665,GO:0044666,GO:0044699,GO:0044763,GO:0045893,GO:0045927,GO:0045935,GO:0048188,GO:0048518,GO:0048522,GO:0050789,GO:0050794,GO:0051171,GO:0051173,GO:0051252,GO:0051254,GO:0051276,GO:0051568,GO:0060255,GO:0065007,GO:0070013,GO:0071704,GO:0071840,GO:0080090,GO:2000112,GO:2001141 | Kelch motif |
| comp210207_c2_seq7.p2 | GO:0005575,GO:0032991,GO:0043234 | GPN-loop GTPase 3 |
| comp213423_c0_seq1.p1 | GO:0001071,GO:0001501,GO:0003674,GO:0003676,GO:0003677,GO:0003700,GO:0005488,GO:0005575,GO:0005622,GO:0005623,GO:0005634,GO:0006355,GO:0007275,GO:0008150,GO:0009889,GO:0010468,GO:0010556,GO:0019219,GO:0019222,GO:0031323,GO:0031326,GO:0032501,GO:0032502,GO:0043226,GO:0043227,GO:0043229,GO:0043231,GO:0044424,GO:0044464,GO:0044699,GO:0044707,GO:0044767,GO:0048731,GO:0048856,GO:0050789,GO:0050794,GO:0051171,GO:0051252,GO:0060255,GO:0060348,GO:0065007,GO:0080090,GO:0097159,GO:1901363,GO:2000112,GO:2001141 | Bobby sox homolog (Drosophila) |
| comp215015_c1_seq4.p1 | GO:0000003,GO:0002064,GO:0003006,GO:0003674,GO:0005488,GO:0005496,GO:0005575,GO:0005622,GO:0005623,GO:0005737,GO:0005783,GO:0005794,GO:0006810,GO:0006886,GO:0006996,GO:0007030,GO:0007275,GO:0007276,GO:0007281,GO:0007283,GO:0007286,GO:0007291,GO:0007349,GO:0008104,GO:0008142,GO:0008150,GO:0008289,GO:0009653,GO:0009888,GO:0009987,GO:0015031,GO:0016043,GO:0019953,GO:0022412,GO:0022414,GO:0030154,GO:0030855,GO:0032501,GO:0032502,GO:0032504,GO:0032934,GO:0033036,GO:0034613,GO:0036094,GO:0043178,GO:0043226,GO:0043227,GO:0043229,GO:0043231,GO:0044424,GO:0044444,GO:0044464,GO:0044699,GO:0044702,GO:0044707,GO:0044763,GO:0044767,GO:0045184,GO:0046907,GO:0048232,GO:0048468,GO:0048515,GO:0048609,GO:0048610,GO:0048646,GO:0048856,GO:0048869,GO:0051179,GO:0051234,GO:0051641,GO:0051649,GO:0060429,GO:0070727,GO:0071702,GO:0071840,GO:0097159 | Oxysterol-binding protein |
| comp185773_c1_seq1.p1 | | Ankyrin repeat |
| comp201923_c0_seq1.p1 | | transporter |
| comp214186_c0_seq2.p1 | GO:0005575,GO:0016020 | Transmembrane protein 179B |
| comp212635_c0_seq5.p1 | | musashi homolog 1 (Drosophila) |

| | | |
|---|---|---|
| comp214040_c0_seq2.p1 | GO:0000785,GO:0002039,GO:0002119,GO:0002164,GO:0003674,GO:0003824,GO:0004175,GO:0004197,GO:0004221,GO:0004843,GO:0005488,GO:0005515,GO:0005575,GO:0005622,GO:0005623,GO:0005634,GO:0005654,GO:0005694,GO:0005700,GO:0005737,GO:0005829,GO:0006139,GO:0006259,GO:0006281,GO:0006283,GO:0006289,GO:0006304,GO:0006305,GO:0006306,GO:0006325,GO:0006342,GO:0006355,GO:0006464,GO:0006725,GO:0006807,GO:0006917,GO:0006950,GO:0006974,GO:0006996,GO:0007044,GO:0007275,GO:0008150,GO:0008152,GO:0008233,GO:0008234,GO:0009791,GO:0009889,GO:0009890,GO:0009892,GO:0009987,GO:0010216,GO:0010468,GO:0010556,GO:0010558,GO:0010605,GO:0010608,GO:0010629,GO:0010941,GO:0010942,GO:0012502,GO:0016043,GO:0016458,GO:0016568,GO:0016569,GO:0016570,GO:0016578,GO:0016579,GO:0016604,GO:0016605,GO:0016787,GO:0016788,GO:0016790,GO:0019219,GO:0019222,GO:0019538,GO:0019783,GO:0022607,GO:0031323,GO:0031324,GO:0031326,GO:0031327,GO:0031581,GO:0031647,GO:0031974,GO:0031981,GO:0032088,GO:0032259,GO:0032501,GO:0032502,GO:0032991,GO:0033554,GO:0034329,GO:0034330,GO:0034641,GO:0035328,GO:0036211,GO:0040008,GO:0040009,GO:0040010,GO:0040014,GO:0040018,GO:0040029,GO:0042802,GO:0042803,GO:0042981,GO:0043065,GO:0043067,GO:0043068,GO:0043170,GO:0043226,GO:0043227,GO:0043228,GO:0043229,GO:0043231,GO:0043232,GO:0043233,GO:0043234,GO:0043412,GO:0043414,GO:0043433,GO:0044085,GO:0044092,GO:0044237,GO:0044238,GO:0044260,GO:0044267,GO:0044422,GO:0044424,GO:0044427,GO:0044428,GO:0044444,GO:0044446,GO:0044451,GO:0044464,GO:0044699,GO:0044707,GO:0044728,GO:0044763,GO:0045814,GO:0045892,GO:0045927,GO:0045934,GO:0046483,GO:0046983,GO:0048518,GO:0048519,GO:0048522,GO:0048523,GO:0050789,GO:0050794,GO:0050896,GO:0051090,GO:0051171,GO:0051172,GO:0051239,GO:0051240,GO:0051252,GO:0051253,GO:0051276,GO:0051716,GO:0060255,GO:0065007,GO:0065008,GO:0065009,GO:0070011,GO:0070013,GO:0070646,GO:0070647,GO:0071704,GO:0071840,GO:0080090,GO:0090304,GO:1901360,GO:2000112,GO:2000113,GO:2001141 | ubiquitin specific peptidase 7 (herpes virus-associated) |
| comp220752_c1_seq8.p1 | GO:0001071,GO:0003674,GO:0003700,GO:0005575,GO:0005622,GO:0005623,GO:0005634,GO:0006355,GO:0008150,GO:0009889,GO:0009890,GO:0009892,GO:0010468,GO:0010556,GO:0010558,GO:0010605,GO:0010629,GO:0019219,GO:0019222,GO:0031323,GO:0031324,GO:0031326,GO:0031327,GO:0043226,GO:0043227,GO:0043229,GO:0043231,GO:0044424,GO:0044464,GO:0045892,GO:0045934,GO:0048519,GO:0048523,GO:0050789,GO:0050794,GO:0051171,GO:0051172,GO:0051252,GO:0051253,GO:0060255,GO:0065007,GO:0080090,GO:2000112,GO:2000113,GO:2001141 | BTAF1 RNA polymerase II, B-TFIID transcription factor-associated, 170kDa (Mot1 homolog, S. cerevisiae) |
| comp205625_c0_seq1.p1 | | Inherit from NOG: pathogenesis |
| comp205625_c0_seq1.p2 | GO:0005575,GO:0005618,GO:0005623,GO:0030312,GO:0044464,GO:0071944 | Inherit from bactNOG: (LipO)protein |
| comp205625_c0_seq1.p3 | | |
| comp205625_c0_seq1.p5 | GO:0000147,GO:0003674,GO:0003779,GO:0005488,GO:0005515,GO:0005575,GO:0005622,GO:0005623,GO:0005737,GO:0005856,GO:0005937,GO:0005938,GO:0006996,GO:0007010,GO:0007163,GO:0008092,GO:0008104,GO:0008150,GO:0009987,GO:0015629,GO:0016043,GO:0022607,GO:0030010,GO:0030029,GO:0030036,GO:0030427,GO:0030479,GO:0030863,GO:0030864,GO:0030865,GO:0030866,GO:0032153,GO:0033036,GO:0042995,GO:0043226,GO:0043228,GO:0043229,GO:0043232,GO:0043332,GO:0044085,GO:0044422,GO:0044424,GO:0044430,GO:0044444,GO:0044446,GO:0044448,GO:0044463,GO:0044464,GO:0044699,GO:0044763,GO:0051015,GO:0051179,GO:0051286,GO:0060187,GO:0071840,GO:0071944 | actin binding protein |
| comp202572_c0_seq1.p1 | | ChaC-like protein |

| | | |
|---|---|---|
| comp213459_c0_seq8.p1 | GO:0000902,GO:0000904,GO:0001763,GO:0002009,GO:0002165,GO:0005575,GO:0005623,GO:0005886,GO:0006928,GO:0006935,GO:0007154,GO:0007155,GO:0007156,GO:0007267,GO:0007275,GO:0007399,GO:0007409,GO:0007411,GO:0007424,GO:0007444,GO:0007472,GO:0007476,GO:0007552,GO:0007560,GO:0008045,GO:0008150,GO:0009605,GO:0009653,GO:0009791,GO:0009886,GO:0009887,GO:0009888,GO:0009987,GO:0016020,GO:0016043,GO:0016337,GO:0016477,GO:0022008,GO:0022610,GO:0023052,GO:0030030,GO:0030154,GO:0030182,GO:0031175,GO:0032501,GO:0032502,GO:0032989,GO:0032990,GO:0035107,GO:0035114,GO:0035120,GO:0035146,GO:0035147,GO:0035220,GO:0035239,GO:0035295,GO:0040011,GO:0042221,GO:0042330,GO:0042995,GO:0044464,GO:0044699,GO:0044700,GO:0044707,GO:0044763,GO:0044767,GO:0045165,GO:0045168,GO:0046331,GO:0048468,GO:0048513,GO:0048563,GO:0048569,GO:0048666,GO:0048667,GO:0048699,GO:0048707,GO:0048729,GO:0048731,GO:0048736,GO:0048737,GO:0048754,GO:0048812,GO:0048856,GO:0048858,GO:0048869,GO:0048870,GO:0050896,GO:0051179,GO:0051674,GO:0060429,GO:0060446,GO:0060541,GO:0060562,GO:0061138,GO:0061564,GO:0071840,GO:0071944,GO:0072499,GO:0097485 | LRRCT |
| comp195514_c1_seq1.p1 | GO:0000165,GO:0000902,GO:0001654,GO:0001700,GO:0001708,GO:0001745,GO:0001763,GO:0002009,GO:0003674,GO:0003824,GO:0004385,GO:0005488,GO:0005515,GO:0005575,GO:0005623,GO:0005886,GO:0005911,GO:0005912,GO:0005913,GO:0005917,GO:0006139,GO:0006725,GO:0006753,GO:0006793,GO:0006796,GO:0006807,GO:0006950,GO:0007043,GO:0007154,GO:0007155,GO:0007163,GO:0007165,GO:0007243,GO:0007254,GO:0007267,GO:0007275,GO:0007391,GO:0007399,GO:0007423,GO:0007424,GO:0007444,GO:0008150,GO:0008152,GO:0009117,GO:0009653,GO:0009790,GO:0009792,GO:0009887,GO:0009888,GO:0009966,GO:0009987,GO:0010627,GO:0010646,GO:0016020,GO:0016043,GO:0016301,GO:0016310,GO:0016323,GO:0016331,GO:0016337,GO:0016740,GO:0016772,GO:0016776,GO:0019201,GO:0019205,GO:0019637,GO:0022008,GO:0022416,GO:0022607,GO:0022610,GO:0023051,GO:0023052,GO:0030030,GO:0030054,GO:0030154,GO:0030155,GO:0030182,GO:0030424,GO:0031098,GO:0031175,GO:0032501,GO:0032502,GO:0032872,GO:0032989,GO:0032990,GO:0033554,GO:0034329,GO:0034330,GO:0034332,GO:0034333,GO:0034641,GO:0035146,GO:0035147,GO:0035153,GO:0035156,GO:0035220,GO:0035239,GO:0035295,GO:0035556,GO:0036056,GO:0036058,GO:0036059,GO:0042995,GO:0043005,GO:0043408,GO:0044085,GO:0044237,GO:0044238,GO:0044281,GO:0044425,GO:0044459,GO:0044464,GO:0044699,GO:0044700,GO:0044707,GO:0044710,GO:0044763,GO:0044767,GO:0045165,GO:0045168,GO:0045177,GO:0045216,GO:0046328,GO:0046331,GO:0046483,GO:0046939,GO:0048468,GO:0048513,GO:0048583,GO:0048592,GO:0048598,GO:0048666,GO:0048699,GO:0048729,GO:0048731,GO:0048749,GO:0048754,GO:0048812,GO:0048856,GO:0048858,GO:0048869,GO:0050789,GO:0050794,GO:0050839,GO:0050896,GO:0051403,GO:0051716,GO:0055086,GO:0060429,GO:0060438,GO:0060439,GO:0060446,GO:0060541,GO:0060562,GO:0061138,GO:0065007,GO:0070161,GO:0070302,GO:0071704,GO:0071840,GO:0071944,GO:0080134,GO:0080135,GO:0097458,GO:1901360 | GuKc |
| comp220770_c0_seq3.p1 | GO:0005575,GO:0005622,GO:0005623,GO:0005634,GO:0005737,GO:0005929,GO:0031514,GO:0036126,GO:0042995,GO:0043226,GO:0043227,GO:0043229,GO:0043231,GO:0044424,GO:0044464,GO:0097223 | sperm associated antigen 17 |
| comp218895_c1_seq4.p1 | | Type III restriction enzyme, res subunit |
| comp211388_c0_seq1.p1 | GO:0000003,GO:0007276,GO:0007283,GO:0008150,GO:0019953,GO:0022414,GO:0032501,GO:0032504,GO:0044699,GO:0044702,GO:0048232,GO:0048609 | Spermatogenesis associated 7 |
| comp220378_c0_seq2.p1 | | Kinesin family member |

| | | |
|---|---|---|
| comp220414_c0_seq2.p1 | GO:0000003,GO:0000123,GO:0000228,GO:0000785,GO:0000786,GO:0000790,GO:0000902,GO:0000904,GO:0000988,GO:0000989,GO:0001501,GO:0001502,GO:0001568,GO:0001708,GO:0001837,GO:0001944,GO:0002376,GO:0002520,GO:0003002,GO:0003006,GO:0003007,GO:0003674,GO:0003676,GO:0003677,GO:0003682,GO:0003712,GO:0003713,GO:0003824,GO:0004402,GO:0005488,GO:0005515,GO:0005575,GO:0005622,GO:0005623,GO:0005634,GO:0005654,GO:0005694,GO:0006139,GO:0006259,GO:0006323,GO:0006325,GO:0006333,GO:0006334,GO:0006355,GO:0006464,GO:0006473,GO:0006475,GO:0006725,GO:0006807,GO:0006810,GO:0006897,GO:0006898,GO:0006928,GO:0006935,GO:0006996,GO:0007155,GO:0007275,GO:0007389,GO:0007399,GO:0007405,GO:0007409,GO:0007411,GO:0007417,GO:0007420,GO:0007507,GO:0007548,GO:0008080,GO:0008134,GO:0008150,GO:0008152,GO:0008270,GO:0008283,GO:0009605,GO:0009653,GO:0009790,GO:0009792,GO:0009880,GO:0009887,GO:0009888,GO:0009889,GO:0009890,GO:0009891,GO:0009892,GO:0009893,GO:0009952,GO:0009966,GO:0009987,GO:0010171,GO:0010172,GO:0010468,GO:0010556,GO:0010557,GO:0010558,GO:0010604,GO:0010605,GO:0010627,GO:0010628,GO:0010629,GO:0010646,GO:0014029,GO:0014031,GO:0014033,GO:0014034,GO:0014036,GO:0016043,GO:0016192,GO:0016319,GO:0016337,GO:0016407,GO:0016410,GO:0016568,GO:0016569,GO:0016570,GO:0016573,GO:0016740,GO:0016746,GO:0016747,GO:0018193,GO:0018205,GO:0018393,GO:0018394,GO:0019219,GO:0019222,GO:0019538,GO:0019827,GO:0022008,GO:0022414,GO:0022607,GO:0022610,GO:0023051,GO:0030030,GO:0030097,GO:0030099,GO:0030154,GO:0030182,GO:0031175,GO:0031323,GO:0031324,GO:0031325,GO:0031326,GO:0031327,GO:0031328,GO:0031497,GO:0031974,GO:0031981,GO:0032501,GO:0032502,GO:0032989,GO:0032990,GO:0032991,GO:0032993,GO:0033563,GO:0034622,GO:0034641,GO:0034728,GO:0035019,GO:0035162,GO:0035904,GO:0035909,GO:0036211,GO:0040008,GO:0040011,GO:0040014,GO:0040018,GO:0040026,GO:0040028,GO:0040035,GO:0042221,GO:0042330,GO:0043009,GO:0043167,GO:0043169,GO:0043170,GO:0043226,GO:0043227,GO:0043228,GO:0043229,GO:0043231,GO:0043232,GO:0043233,GO:0043234,GO:0043412,GO:0043433,GO:0043543,GO:0043933,GO:0043966,GO:0044085,GO:0044092,GO:0044237,GO:0044238,GO:0044260,GO:0044267,GO:0044422,GO:0044424,GO:0044427,GO:0044428,GO:0044446,GO:0044451,GO:0044454,GO:0044464,GO:0044699,GO:0044707,GO:0044763,GO:0044767,GO:0045165,GO:0045595,GO:0045596,GO:0045892,GO:0045893,GO:0045927,GO:0045934,GO:0045935,GO:0046425,GO:0046483,GO:0046872,GO:0046914,GO:0048468,GO:0048513,GO:0048514,GO:0048518,GO:0048519,GO:0048522,GO:0048523,GO:0048534,GO:0048562,GO:0048568,GO:0048580,GO:0048583,GO:0048598,GO:0048608,GO:0048666,GO:0048667,GO:0048699,GO:0048701,GO:0048703,GO:0048704,GO:0048705,GO:0048706,GO:0048731,GO:0048762,GO:0048806,GO:0048812,GO:0048844,GO:0048856,GO:0048858,GO:0048863,GO:0048864,GO:0048869,GO:0050789,GO:0050793,GO:0050794,GO:0050896,GO:0051090,GO:0051093,GO:0051094,GO:0051171,GO:0051172,GO:0051173,GO:0051179,GO:0051216,GO:0051234,GO:0051239,GO:0051240,GO:0051252,GO:0051253,GO:0051254,GO:0051276,GO:0060255,GO:0060322,GO:0060323,GO:0060324,GO:0060325,GO:0060485,GO:0060840,GO:0061062,GO:0061351,GO:0061448,GO:0061458,GO:0061564,GO:0065003,GO:0065004,GO:0065007,GO:0065009,GO:0070013,GO:0070775,GO:0070776,GO:0071103,GO:0071704,GO:0071824,GO:0071840,GO:0072089,GO:0072358,GO:0072359,GO:0080090,GO:0090304,GO:0097159,GO:0097485,GO:1901360,GO:1901363,GO:2000026,GO:2000112,GO:2000113,GO:2001141 | K(lysine) acetyltransferase |
| comp213150_c0_seq5.p1 | | RING |
| comp221016_c0_seq11.p1 | | GAS2 |

| | | |
|---|---|---|
| comp219559_c2_seq7.p1 | GO:0000003,GO:0000226,GO:0000578,GO:0001664,GO:0001709,GO:0003002,GO:0003006,GO:0003008,GO:0003674,GO:0005102,GO:0005198,GO:0005200,GO:0005488,GO:0005515,GO:0005575,GO:0005576,GO:0005622,GO:0005623,GO:0005737,GO:0005794,GO:0005802,GO:0005829,GO:0005856,GO:0005886,GO:0006139,GO:0006396,GO:0006403,GO:0006725,GO:0006807,GO:0006810,GO:0006897,GO:0006928,GO:0006996,GO:0007010,GO:0007017,GO:0007018,GO:0007097,GO:0007154,GO:0007163,GO:0007267,GO:0007268,GO:0007275,GO:0007276,GO:0007281,GO:0007292,GO:0007293,GO:0007294,GO:0007308,GO:0007309,GO:0007310,GO:0007312,GO:0007314,GO:0007350,GO:0007351,GO:0007389,GO:0007399,GO:0007548,GO:0008092,GO:0008093,GO:0008103,GO:0008104,GO:0008150,GO:0008152,GO:0008277,GO:0008298,GO:0008358,GO:0008595,GO:0009653,GO:0009790,GO:0009792,GO:0009798,GO:0009880,GO:0009948,GO:0009950,GO:0009952,GO:0009953,GO:0009966,GO:0009968,GO:0009987,GO:0009994,GO:0010467,GO:0010469,GO:0010517,GO:0010519,GO:0010646,GO:0010648,GO:0010970,GO:0015931,GO:0016020,GO:0016032,GO:0016043,GO:0016070,GO:0016192,GO:0016325,GO:0016358,GO:0017016,GO:0017137,GO:0017157,GO:0018995,GO:0019222,GO:0019226,GO:0019899,GO:0019953,GO:0021700,GO:0022008,GO:0022412,GO:0022414,GO:0022607,GO:0022613,GO:0022618,GO:0023051,GO:0023052,GO:0023057,GO:0030030,GO:0030100,GO:0030154,GO:0030182,GO:0030276,GO:0030430,GO:0030674,GO:0030705,GO:0030706,GO:0030716,GO:0030722,GO:0030951,GO:0030952,GO:0031175,GO:0031267,GO:0031410,GO:0031644,GO:0031871,GO:0031982,GO:0032050,GO:0032501,GO:0032502,GO:0032504,GO:0032507,GO:0032879,GO:0033036,GO:0033365,GO:0033643,GO:0033646,GO:0033647,GO:0033648,GO:0033655,GO:0034063,GO:0034452,GO:0034453,GO:0034613,GO:0034622,GO:0034641,GO:0035282,GO:0035637,GO:0040008,GO:0040011,GO:0040014,GO:0040018,GO:0040023,GO:0040035,GO:0043086,GO:0043170,GO:0043226,GO:0043227,GO:0043228,GO:0043229,GO:0043231,GO:0043232,GO:0043245,GO:0043656,GO:0043657,GO:0043933,GO:0044057,GO:0044085,GO:0044092,GO:0044215,GO:0044216,GO:0044217,GO:0044237,GO:0044238,GO:0044260,GO:0044403,GO:0044419,GO:0044421,GO:0044424,GO:0044444,GO:0044464,GO:0044699,GO:0044700,GO:0044702,GO:0044707,GO:0044763,GO:0044764,GO:0044765,GO:0044767,GO:0045165,GO:0045185,GO:0045502,GO:0045744,GO:0045807,GO:0045921,GO:0045927,GO:0046483,GO:0046907,GO:0046928,GO:0048259,GO:0048260,GO:0048468,GO:0048469,GO:0048471,GO:0048477,GO:0048488,GO:0048489,GO:0048513,GO:0048518,GO:0048519,GO:0048522,GO:0048523,GO:0048583,GO:0048585,GO:0048599,GO:0048608,GO:0048609,GO:0048610,GO:0048646,GO:0048666,GO:0048699,GO:0048731,GO:0048806,GO:0048856,GO:0048869,GO:0050657,GO:0050658,GO:0050789,GO:0050790,GO:0050794,GO:0050804,GO:0050877,GO:0051020,GO:0051028,GO:0051046,GO:0051047,GO:0051049,GO:0051050,GO:0051128,GO:0051130,GO:0051179,GO:0051234,GO:0051235,GO:0051236,GO:0051239,GO:0051240,GO:0051336,GO:0051346,GO:0051588,GO:0051640,GO:0051641,GO:0051647,GO:0051648,GO:0051649,GO:0051650,GO:0051651,GO:0051656,GO:0051663,GO:0051704,GO:0051969,GO:0060090,GO:0060191,GO:0060192,GO:0060341,GO:0060627,GO:0061458,GO:0065003,GO:0065007,GO:0065008,GO:0065009,GO:0070727,GO:0070925,GO:0071702,GO:0071704,GO:0071705,GO:0071826,GO:0071840,GO:0071944,GO:0072384,GO:0072385,GO:0072393,GO:0072517,GO:0090304,GO:0097479,GO:0097480,GO:1900274,GO:1900275,GO:1900276,GO:1900736,GO:1900737,GO:1901360,GO:2000300,GO:2000302,GO:2000369,GO:2000370 | bicaudal D homolog |
| comp216121_c0_seq1.p1 | GO:0003674,GO:0003824,GO:0008150,GO:0008152,GO:0008194,GO:0008378,GO:0008499,GO:0016740,GO:0016757,GO:0016758,GO:0035250,GO:0048531 | Galactosyltransferase |
| comp214011_c0_seq2.p1 | | CHY zinc finger |

| | | |
|---|---|---|
| comp215010_c0_seq2.p1 | GO:0001525,GO:0001568,GO:0001838,GO:0001841,GO:0001906,GO:0001907,GO:0001944,GO:0002009,GO:0002218,GO:0002221,GO:0002237,GO:0002253,GO:0002376,GO:0002520,GO:0002521,GO:0002573,GO:0002682,GO:0002684,GO:0002753,GO:0002757,GO:0002758,GO:0002761,GO:0002763,GO:0002764,GO:0003674,GO:0003824,GO:0004175,GO:0004197,GO:0005575,GO:0005622,GO:0005623,GO:0005634,GO:0005737,GO:0005739,GO:0005740,GO:0005741,GO:0005813,GO:0005815,GO:0005829,GO:0005856,GO:0005886,GO:0006461,GO:0006508,GO:0006915,GO:0006917,GO:0006919,GO:0006921,GO:0006926,GO:0006950,GO:0006952,GO:0006955,GO:0007154,GO:0007165,GO:0007166,GO:0007275,GO:0007399,GO:0007507,GO:0008150,GO:0008152,GO:0008219,GO:0008233,GO:0008234,GO:0009056,GO:0009057,GO:0009266,GO:0009409,GO:0009605,GO:0009607,GO:0009612,GO:0009617,GO:0009628,GO:0009636,GO:0009653,GO:0009719,GO:0009725,GO:0009790,GO:0009792,GO:0009888,GO:0009966,GO:0009967,GO:0009968,GO:0009987,GO:0010033,GO:0010035,GO:0010038,GO:0010627,GO:0010646,GO:0010647,GO:0010648,GO:0010740,GO:0010741,GO:0010941,GO:0010942,GO:0010950,GO:0010952,GO:0012501,GO:0012502,GO:0014070,GO:0015630,GO:0016020,GO:0016032,GO:0016043,GO:0016265,GO:0016331,GO:0016787,GO:0019048,GO:0019051,GO:0019054,GO:0019222,GO:0019538,GO:0019867,GO:0021915,GO:0022411,GO:0022607,GO:0023051,GO:0023052,GO:0023056,GO:0023057,GO:0030097,GO:0030099,GO:0030154,GO:0030163,GO:0030225,GO:0030522,GO:0030529,GO:0030684,GO:0030686,GO:0030687,GO:0030689,GO:0030690,GO:0031090,GO:0031264,GO:0031265,GO:0031341,GO:0031343,GO:0031347,GO:0031349,GO:0031640,GO:0031966,GO:0031967,GO:0031968,GO:0031975,GO:0032025,GO:0032355,GO:0032496,GO:0032501,GO:0032502,GO:0032991,GO:0033993,GO:0034097,GO:0034612,GO:0035148,GO:0035239,GO:0035295,GO:0035556,GO:0035821,GO:0035872,GO:0039526,GO:0042221,GO:0042981,GO:0042995,GO:0043005,GO:0043009,GO:0043065,GO:0043067,GO:0043068,GO:0043085,GO:0043122,GO:0043123,GO:0043124,GO:0043170,GO:0043226,GO:0043227,GO:0043228,GO:0043229,GO:0043231,GO:0043232,GO:0043234,GO:0043280,GO:0043281,GO:0043627,GO:0043900,GO:0043902,GO:0043903,GO:0043933,GO:0044003,GO:0044004,GO:0044068,GO:0044085,GO:0044093,GO:0044237,GO:0044238,GO:0044248,GO:0044257,GO:0044260,GO:0044265,GO:0044267,GO:0044297,GO:0044364,GO:0044403,GO:0044419,GO:0044422,GO:0044424,GO:0044425,GO:0044428,GO:0044429,GO:0044430,GO:0044444,GO:0044446,GO:0044459,GO:0044464,GO:0044531,GO:0044532,GO:0044533,GO:0044699,GO:0044700,GO:0044707,GO:0044763,GO:0044764,GO:0044767,GO:0045087,GO:0045088,GO:0045089,GO:0045121,GO:0045471,GO:0045595,GO:0045597,GO:0045637,GO:0045639,GO:0045649,GO:0045651,GO:0046677,GO:0048513,GO:0048514,GO:0048518,GO:0048519,GO:0048522,GO:0048523,GO:0048534,GO:0048545,GO:0048583,GO:0048584,GO:0048585,GO:0048598,GO:0048646,GO:0048729,GO:0048731,GO:0048856,GO:0048869,GO:0050776,GO:0050778,GO:0050789,GO:0050790,GO:0050793,GO:0050794,GO:0050896,GO:0051094,GO:0051239,GO:0051259,GO:0051291,GO:0051336,GO:0051345,GO:0051603,GO:0051701,GO:0051704,GO:0051707,GO:0051709,GO:0051712,GO:0051716,GO:0051817,GO:0051818,GO:0051883,GO:0052040,GO:0052042,GO:0052150,GO:0052151,GO:0052248,GO:0052330,GO:0052433,GO:0052501,GO:0052547,GO:0052548,GO:0060139,GO:0060429,GO:0060562,GO:0065003,GO:0065007,GO:0065008,GO:0065009,GO:0070011,GO:0070271,GO:0070423,GO:0070887,GO:0071214,GO:0071260,GO:0071310,GO:0071407,GO:0071496,GO:0071704,GO:0071822,GO:0071840,GO:0071944,GO:0072175,GO:0072358,GO:0072359,GO:0080134,GO:0097153,GO:0097190,GO:0097191,GO:0097194,GO:0097202,GO:0097285,GO:0097305,GO:0097458,GO:1901575,GO:1901700,GO:1902105,GO:1902107,GO:2000026,GO:2000116,GO:2001056 | apoptosis-related cysteine peptidase |

| | | |
|---|---|---|
| comp215010_c0_seq2.p2 | GO:0001525,GO:0001568,GO:0001838,GO:0001841,GO:0001906,GO:0001907,GO:0001944,GO:0002009,GO:0002218,GO:0002221,GO:0002237,GO:0002253,GO:0002376,GO:0002520,GO:0002521,GO:0002573,GO:0002682,GO:0002684,GO:0002753,GO:0002757,GO:0002758,GO:0002761,GO:0002763,GO:0002764,GO:0003674,GO:0003824,GO:0004175,GO:0004197,GO:0005575,GO:0005622,GO:0005623,GO:0005634,GO:0005737,GO:0005739,GO:0005740,GO:0005741,GO:0005813,GO:0005815,GO:0005829,GO:0005856,GO:0005886,GO:0006461,GO:0006508,GO:0006915,GO:0006917,GO:0006919,GO:0006921,GO:0006926,GO:0006950,GO:0006952,GO:0006955,GO:0007154,GO:0007165,GO:0007166,GO:0007275,GO:0007399,GO:0007507,GO:0008150,GO:0008152,GO:0008219,GO:0008233,GO:0008234,GO:0009056,GO:0009057,GO:0009266,GO:0009409,GO:0009605,GO:0009607,GO:0009612,GO:0009617,GO:0009628,GO:0009636,GO:0009653,GO:0009719,GO:0009725,GO:0009790,GO:0009792,GO:0009888,GO:0009966,GO:0009967,GO:0009968,GO:0009987,GO:0010033,GO:0010035,GO:0010038,GO:0010627,GO:0010646,GO:0010647,GO:0010648,GO:0010740,GO:0010741,GO:0010941,GO:0010942,GO:0010950,GO:0010952,GO:0012501,GO:0012502,GO:0014070,GO:0015630,GO:0016020,GO:0016032,GO:0016043,GO:0016265,GO:0016331,GO:0016787,GO:0019048,GO:0019051,GO:0019054,GO:0019222,GO:0019538,GO:0019867,GO:0021915,GO:0022411,GO:0022607,GO:0023051,GO:0023052,GO:0023056,GO:0023057,GO:0030097,GO:0030099,GO:0030154,GO:0030163,GO:0030225,GO:0030522,GO:0030529,GO:0030684,GO:0030686,GO:0030687,GO:0030689,GO:0030690,GO:0031090,GO:0031264,GO:0031265,GO:0031341,GO:0031343,GO:0031347,GO:0031349,GO:0031640,GO:0031966,GO:0031967,GO:0031968,GO:0031975,GO:0032025,GO:0032355,GO:0032496,GO:0032501,GO:0032502,GO:0032991,GO:0033993,GO:0034097,GO:0034612,GO:0035148,GO:0035239,GO:0035295,GO:0035556,GO:0035821,GO:0035872,GO:0039526,GO:0042221,GO:0042981,GO:0042995,GO:0043005,GO:0043009,GO:0043065,GO:0043067,GO:0043068,GO:0043085,GO:0043122,GO:0043123,GO:0043124,GO:0043170,GO:0043226,GO:0043227,GO:0043228,GO:0043229,GO:0043231,GO:0043232,GO:0043234,GO:0043280,GO:0043281,GO:0043627,GO:0043900,GO:0043902,GO:0043903,GO:0043933,GO:0044003,GO:0044004,GO:0044068,GO:0044085,GO:0044093,GO:0044237,GO:0044238,GO:0044248,GO:0044257,GO:0044260,GO:0044265,GO:0044267,GO:0044297,GO:0044364,GO:0044403,GO:0044419,GO:0044422,GO:0044424,GO:0044425,GO:0044428,GO:0044429,GO:0044430,GO:0044444,GO:0044446,GO:0044459,GO:0044464,GO:0044531,GO:0044532,GO:0044533,GO:0044699,GO:0044700,GO:0044707,GO:0044763,GO:0044764,GO:0044767,GO:0045087,GO:0045088,GO:0045089,GO:0045121,GO:0045471,GO:0045595,GO:0045597,GO:0045637,GO:0045639,GO:0045649,GO:0045651,GO:0046677,GO:0048513,GO:0048514,GO:0048518,GO:0048519,GO:0048522,GO:0048523,GO:0048534,GO:0048545,GO:0048583,GO:0048584,GO:0048585,GO:0048598,GO:0048646,GO:0048729,GO:0048731,GO:0048856,GO:0048869,GO:0050776,GO:0050778,GO:0050789,GO:0050790,GO:0050793,GO:0050794,GO:0050896,GO:0051094,GO:0051239,GO:0051259,GO:0051291,GO:0051336,GO:0051345,GO:0051603,GO:0051701,GO:0051704,GO:0051707,GO:0051709,GO:0051712,GO:0051716,GO:0051817,GO:0051818,GO:0051883,GO:0052040,GO:0052042,GO:0052150,GO:0052151,GO:0052248,GO:0052330,GO:0052433,GO:0052501,GO:0052547,GO:0052548,GO:0060139,GO:0060429,GO:0060562,GO:0065003,GO:0065007,GO:0065008,GO:0065009,GO:0070011,GO:0070271,GO:0070423,GO:0070887,GO:0071214,GO:0071260,GO:0071310,GO:0071407,GO:0071496,GO:0071704,GO:0071822,GO:0071840,GO:0071944,GO:0072175,GO:0072358,GO:0072359,GO:0080134,GO:0097153,GO:0097190,GO:0097191,GO:0097194,GO:0097202,GO:0097285,GO:0097305,GO:0097458,GO:1901575,GO:1901700,GO:1902105,GO:1902107,GO:2000026,GO:2000116,GO:2001056 | apoptosis-related cysteine peptidase |
| comp221244_c0_seq7.p1 | GO:0000003,GO:0000139,GO:0003006,GO:0005575,GO:0005622,GO:0005623,GO:0005737,GO:0005794,GO:0006810,GO:0006892,GO:0006895,GO:0006996,GO:0007029,GO:0007275,GO:0007548,GO:0008150,GO:0009790,GO:0009792,GO:0009987,GO:0012505,GO:0016020,GO:0016043,GO:0016192,GO:0016197,GO:0016482,GO:0022414,GO:0031090,GO:0032501,GO:0032502,GO:0040008,GO:0040009,GO:0040010,GO:0040035,GO:0043226,GO:0043227,GO:0043229,GO:0043231,GO:0044422,GO:0044424,GO:0044431,GO:0044444,GO:0044446,GO:0044464,GO:0044699,GO:0044707,GO:0044763,GO:0044765,GO:0044767,GO:0045927,GO:0046907,GO:0048193,GO:0048513,GO:0048518,GO:0048608,GO:0048731,GO:0048806,GO:0048856,GO:0050789,GO:0051179,GO:0051234,GO:0051641,GO:0051649,GO:0061458,GO:0065007,GO:0071840 | Dopey family member |

| | | |
|---|---|---|
| comp212879_c0_seq5.p1 | | transformer 2 beta homolog (Drosophila) |
| comp217337_c1_seq3.p1 | GO:0000226,GO:0003674,GO:0005198,GO:0005488,GO:0005515,GO:0005575,GO:0005622,GO:0005623,GO:0005737,GO:0005813,GO:0005814,GO:0005815,GO:0005856,GO:0005886,GO:0005929,GO:0006996,GO:0007010,GO:0007017,GO:0008092,GO:0008104,GO:0008150,GO:0009987,GO:0015629,GO:0015630,GO:0016020,GO:0016043,GO:0019894,GO:0031023,GO:0031514,GO:0033036,GO:0035253,GO:0042995,GO:0043226,GO:0043227,GO:0043228,GO:0043229,GO:0043231,GO:0043232,GO:0044422,GO:0044424,GO:0044430,GO:0044441,GO:0044444,GO:0044446,GO:0044450,GO:0044463,GO:0044464,GO:0044699,GO:0044763,GO:0051179,GO:0051297,GO:0071840,GO:0071944 | Ciliary rootlet coiled-coil, rootletin |
| comp205932_c1_seq2.p1 | GO:0000003,GO:0002119,GO:0002164,GO:0003674,GO:0005488,GO:0005515,GO:0005575,GO:0005622,GO:0005623,GO:0005634,GO:0005737,GO:0006810,GO:0006897,GO:0006898,GO:0007275,GO:0007568,GO:0007610,GO:0008150,GO:0008340,GO:0009790,GO:0009791,GO:0009792,GO:0010259,GO:0016192,GO:0018991,GO:0019098,GO:0022414,GO:0022611,GO:0032501,GO:0032502,GO:0032504,GO:0033057,GO:0035188,GO:0040007,GO:0040008,GO:0040009,GO:0040010,GO:0040024,GO:0043053,GO:0043226,GO:0043227,GO:0043229,GO:0043231,GO:0044424,GO:0044464,GO:0044699,GO:0044706,GO:0044707,GO:0044767,GO:0045927,GO:0048518,GO:0048609,GO:0048856,GO:0050789,GO:0050896,GO:0051179,GO:0051234,GO:0051704,GO:0055115,GO:0065007,GO:0071684 | 14_3_3 |
| comp217837_c0_seq5.p1 | | Zinc finger protein |
| comp216281_c0_seq13.p1 | GO:0000003,GO:0003674,GO:0005215,GO:0005216,GO:0005261,GO:0005262,GO:0005488,GO:0005515,GO:0005575,GO:0005622,GO:0005623,GO:0005929,GO:0006810,GO:0006811,GO:0006812,GO:0006816,GO:0007610,GO:0007617,GO:0007618,GO:0008150,GO:0008324,GO:0009581,GO:0009582,GO:0009605,GO:0009612,GO:0009628,GO:0009987,GO:0015075,GO:0015085,GO:0015267,GO:0016020,GO:0019098,GO:0019904,GO:0022414,GO:0022803,GO:0022838,GO:0022857,GO:0022890,GO:0022891,GO:0022892,GO:0030001,GO:0031513,GO:0032501,GO:0032504,GO:0033057,GO:0034220,GO:0034606,GO:0034608,GO:0040008,GO:0040009,GO:0040010,GO:0042995,GO:0043025,GO:0043226,GO:0043227,GO:0043229,GO:0043231,GO:0044297,GO:0044424,GO:0044464,GO:0044699,GO:0044703,GO:0044706,GO:0044763,GO:0044765,GO:0045927,GO:0046873,GO:0048518,GO:0048609,GO:0050789,GO:0050896,GO:0050982,GO:0051179,GO:0051234,GO:0051606,GO:0051704,GO:0051705,GO:0055085,GO:0060179,GO:0065007,GO:0070588,GO:0070838,GO:0072372,GO:0072509,GO:0072511,GO:0097458 | polycystic kidney disease 1-like 2 |
| comp208251_c1_seq2.p1 | | Src homology 2 domain containing F |
| comp207861_c1_seq3.p1 | GO:0000003,GO:0001709,GO:0003006,GO:0003674,GO:0005488,GO:0005515,GO:0005575,GO:0006139,GO:0006259,GO:0006281,GO:0006302,GO:0006457,GO:0006725,GO:0006807,GO:0006950,GO:0006974,GO:0007276,GO:0007292,GO:0007293,GO:0007294,GO:0008150,GO:0008152,GO:0009653,GO:0009987,GO:0009994,GO:0010564,GO:0019538,GO:0019953,GO:0022412,GO:0022414,GO:0030154,GO:0030706,GO:0030716,GO:0032501,GO:0032502,GO:0032504,GO:0032991,GO:0033554,GO:0034641,GO:0040020,GO:0043170,GO:0043234,GO:0044237,GO:0044238,GO:0044260,GO:0044267,GO:0044699,GO:0044702,GO:0044763,GO:0044767,GO:0045165,GO:0046483,GO:0048477,GO:0048609,GO:0048610,GO:0048646,GO:0048856,GO:0048869,GO:0050789,GO:0050794,GO:0050896,GO:0051082,GO:0051445,GO:0051716,GO:0051726,GO:0065007,GO:0071704,GO:0090304,GO:1901360 | missing oocyte, meiosis regulator, homolog (Drosophila) |

| | | |
|---|---|---|
| comp214593_c0_seq1.p1 | GO:0001786,GO:0001948,GO:0003002,GO:0003674,GO:0003779,GO:0005488,GO:0005509,GO:0005515,GO:0005539,GO:0005543,GO:0005544,GO:0005545,GO:0005575,GO:0005622,GO:0005623,GO:0005634,GO:0005635,GO:0005737,GO:0005783,GO:0005789,GO:0005829,GO:0005886,GO:0005938,GO:0006139,GO:0006351,GO:0006355,GO:0006357,GO:0006725,GO:0006807,GO:0006810,GO:0006873,GO:0006874,GO:0006875,GO:0006884,GO:0006897,GO:0006909,GO:0006911,GO:0006944,GO:0006950,GO:0006970,GO:0007155,GO:0007163,GO:0007275,GO:0007389,GO:0007444,GO:0007447,GO:0007450,GO:0007599,GO:0007610,GO:0008092,GO:0008104,GO:0008105,GO:0008150,GO:0008152,GO:0008201,GO:0008283,GO:0008289,GO:0008360,GO:0008429,GO:0009058,GO:0009059,GO:0009628,GO:0009651,GO:0009790,GO:0009792,GO:0009889,GO:0009953,GO:0009987,GO:0009992,GO:0010033,GO:0010035,GO:0010038,GO:0010324,GO:0010467,GO:0010468,GO:0010556,GO:0010796,GO:0010797,GO:0012505,GO:0014070,GO:0016020,GO:0016043,GO:0016044,GO:0016070,GO:0016192,GO:0016197,GO:0016597,GO:0018130,GO:0019219,GO:0019222,GO:0019438,GO:0019725,GO:0022603,GO:0022604,GO:0022610,GO:0030003,GO:0030011,GO:0030104,GO:0030507,GO:0031090,GO:0031323,GO:0031326,GO:0031406,GO:0031967,GO:0031975,GO:0031982,GO:0032501,GO:0032502,GO:0032509,GO:0032535,GO:0032774,GO:0033036,GO:0034641,GO:0034645,GO:0034654,GO:0035091,GO:0035176,GO:0035220,GO:0035222,GO:0042175,GO:0042221,GO:0042592,GO:0043167,GO:0043168,GO:0043169,GO:0043170,GO:0043226,GO:0043227,GO:0043229,GO:0043231,GO:0043277,GO:0043394,GO:0043395,GO:0043652,GO:0044237,GO:0044238,GO:0044249,GO:0044260,GO:0044271,GO:0044422,GO:0044424,GO:0044425,GO:0044428,GO:0044432,GO:0044444,GO:0044446,GO:0044464,GO:0044699,GO:0044707,GO:0044763,GO:0044765,GO:0044767,GO:0046483,GO:0046872,GO:0046907,GO:0048190,GO:0048513,GO:0048731,GO:0048856,GO:0048878,GO:0050789,GO:0050793,GO:0050794,GO:0050801,GO:0050878,GO:0050896,GO:0051036,GO:0051128,GO:0051171,GO:0051179,GO:0051234,GO:0051252,GO:0051592,GO:0051641,GO:0051649,GO:0051703,GO:0051704,GO:0051705,GO:0055065,GO:0055074,GO:0055080,GO:0055082,GO:0060255,GO:0061024,GO:0061025,GO:0061467,GO:0065007,GO:0065008,GO:0071704,GO:0071840,GO:0071944,GO:0072341,GO:0072503,GO:0072507,GO:0080090,GO:0090066,GO:0090304,GO:0097367,GO:1901360,GO:1901362,GO:1901576,GO:1901681,GO:2000112,GO:2001141 | annexin A11 |
| comp215854_c0_seq3.p1 | GO:0003674,GO:0005488,GO:0005515,GO:0005575,GO:0005622,GO:0005623,GO:0005634,GO:0005737,GO:0005739,GO:0006457,GO:0008150,GO:0008152,GO:0009987,GO:0019538,GO:0030424,GO:0030425,GO:0042995,GO:0043005,GO:0043170,GO:0043226,GO:0043227,GO:0043229,GO:0043231,GO:0044087,GO:0044237,GO:0044238,GO:0044260,GO:0044267,GO:0044424,GO:0044444,GO:0044464,GO:0048519,GO:0050789,GO:0051087,GO:0051128,GO:0051129,GO:0065007,GO:0070852,GO:0071704,GO:0090083,GO:0090084,GO:0097458 | Spastic ataxia of Charlevoix-Saguenay (Sacsin) |
| comp211515_c2_seq5.p1 | | ZIP Zinc transporter |
| comp200867_c0_seq1.p1 | | PDZ domain containing 2 |
| comp214066_c0_seq8.p1 | GO:0000902,GO:0000904,GO:0001709,GO:0001764,GO:0005575,GO:0005622,GO:0005623,GO:0005634,GO:0005730,GO:0005737,GO:0005938,GO:0006928,GO:0006935,GO:0007275,GO:0007399,GO:0007405,GO:0007409,GO:0007411,GO:0008045,GO:0008104,GO:0008105,GO:0008150,GO:0008283,GO:0009605,GO:0009653,GO:0009987,GO:0016043,GO:0016477,GO:0017145,GO:0022008,GO:0030030,GO:0030154,GO:0030182,GO:0031175,GO:0031974,GO:0031981,GO:0032501,GO:0032502,GO:0032989,GO:0032990,GO:0033036,GO:0040011,GO:0042221,GO:0042330,GO:0043226,GO:0043227,GO:0043228,GO:0043229,GO:0043231,GO:0043232,GO:0043233,GO:0044422,GO:0044424,GO:0044428,GO:0044444,GO:0044446,GO:0044464,GO:0044699,GO:0044707,GO:0044763,GO:0044767,GO:0045165,GO:0045167,GO:0048103,GO:0048468,GO:0048666,GO:0048667,GO:0048699,GO:0048731,GO:0048812,GO:0048856,GO:0048858,GO:0048869,GO:0048870,GO:0050896,GO:0051179,GO:0051301,GO:0051674,GO:0055057,GO:0055059,GO:0061351,GO:0061564,GO:0070013,GO:0071840,GO:0071944,GO:0072089,GO:0097485 | Storkhead box 1 |

| | | |
|---|---|---|
| comp217136_c0_seq2.p1 | GO:0005575,GO:0005622,GO:0005623,GO:0005634,GO:0043226,GO:0043227,GO:0043229,GO:0043231,GO:0044424,GO:0044464 | Inherit from biNOG: cut-like homeobox |
| comp219117_c1_seq2.p1 | GO:0003674,GO:0003824,GO:0004672,GO:0004674,GO:0005575,GO:0005622,GO:0005623,GO:0005634,GO:0005737,GO:0005794,GO:0006464,GO:0006468,GO:0006793,GO:0006796,GO:0008150,GO:0008152,GO:0009987,GO:0016301,GO:0016310,GO:0016740,GO:0016772,GO:0016773,GO:0019538,GO:0036211,GO:0043170,GO:0043226,GO:0043227,GO:0043229,GO:0043231,GO:0043412,GO:0044237,GO:0044238,GO:0044260,GO:0044267,GO:0044424,GO:0044444,GO:0044464,GO:0046777,GO:0048471,GO:0071704 | Serine threonine kinase 33 |
| comp194622_c1_seq3.p1 | GO:0003674,GO:0004889,GO:0005215,GO:0005216,GO:0005230,GO:0005231,GO:0005261,GO:0005575,GO:0005623,GO:0005886,GO:0005887,GO:0005892,GO:0006810,GO:0006811,GO:0006812,GO:0008150,GO:0008324,GO:0009987,GO:0015075,GO:0015267,GO:0015276,GO:0016020,GO:0016021,GO:0022803,GO:0022834,GO:0022836,GO:0022838,GO:0022839,GO:0022857,GO:0022891,GO:0022892,GO:0031224,GO:0031226,GO:0032991,GO:0034220,GO:0034702,GO:0043234,GO:0043235,GO:0044425,GO:0044456,GO:0044459,GO:0044464,GO:0044699,GO:0044763,GO:0044765,GO:0045202,GO:0045211,GO:0051179,GO:0051234,GO:0055085,GO:0071944,GO:0097060 | Neurotransmitter-gated ion-channel transmembrane region |
| comp220022_c0_seq1.p1 | | coiled-coil domain containing 18 |
| comp175699_c0_seq2.p1 | | conserved hypothetical protein |
| comp215078_c1_seq1.p1 | GO:0000132,GO:0000226,GO:0000278,GO:0001764,GO:0001838,GO:0001841,GO:0002009,GO:0005575,GO:0005622,GO:0005623,GO:0005737,GO:0005886,GO:0005911,GO:0005912,GO:0005913,GO:0006140,GO:0006915,GO:0006928,GO:0006935,GO:0006996,GO:0007010,GO:0007017,GO:0007049,GO:0007155,GO:0007163,GO:0007275,GO:0007346,GO:0007369,GO:0007399,GO:0008104,GO:0008150,GO:0008219,GO:0008283,GO:0009118,GO:0009605,GO:0009653,GO:0009790,GO:0009792,GO:0009888,GO:0009894,GO:0009966,GO:0009987,GO:0010623,GO:0010646,GO:0010941,GO:0010942,GO:0012501,GO:0014021,GO:0016020,GO:0016043,GO:0016265,GO:0016331,GO:0016337,GO:0016477,GO:0019219,GO:0019220,GO:0019222,GO:0021915,GO:0022008,GO:0022402,GO:0022610,GO:0023051,GO:0030010,GO:0030054,GO:0030154,GO:0030811,GO:0031323,GO:0031329,GO:0031503,GO:0032318,GO:0032319,GO:0032320,GO:0032321,GO:0032501,GO:0032502,GO:0032855,GO:0032856,GO:0032862,GO:0032863,GO:0033036,GO:0033121,GO:0033124,GO:0035023,GO:0035088,GO:0035089,GO:0035148,GO:0035239,GO:0035295,GO:0040001,GO:0040011,GO:0042221,GO:0042330,GO:0042981,GO:0043009,GO:0043065,GO:0043067,GO:0043068,GO:0043085,GO:0043087,GO:0043547,GO:0044093,GO:0044424,GO:0044464,GO:0044699,GO:0044707,GO:0044763,GO:0044767,GO:0045786,GO:0045930,GO:0046578,GO:0048468,GO:0048518,GO:0048519,GO:0048522,GO:0048523,GO:0048583,GO:0048598,GO:0048646,GO:0048699,GO:0048729,GO:0048731,GO:0048856,GO:0048869,GO:0048870,GO:0050789,GO:0050790,GO:0050794,GO:0050896,GO:0050918,GO:0051056,GO:0051171,GO:0051174,GO:0051179,GO:0051234,GO:0051293,GO:0051294,GO:0051336,GO:0051345,GO:0051640,GO:0051641,GO:0051649,GO:0051653,GO:0051656,GO:0051674,GO:0051726,GO:0060026,GO:0060027,GO:0060429,GO:0060561,GO:0060562,GO:0061162,GO:0061245,GO:0061339,GO:0065007,GO:0065009,GO:0070161,GO:0071840,GO:0071944,GO:0072175,GO:0080090,GO:1900542 | scribbled homolog (Drosophila) |

| | | |
|---|---|---|
| comp220769_c1_seq17.p1 | GO:0000003,GO:0000166,GO:0000578,GO:0001654,GO:0001745,GO:0001751,GO:0001754,GO:0001882,GO:0001883,GO:0002064,GO:0003002,GO:0003006,GO:0003674,GO:0003774,GO:0003779,GO:0003824,GO:0005488,GO:0005515,GO:0005516,GO:0005524,GO:0005575,GO:0005622,GO:0005623,GO:0005856,GO:0006139,GO:0006152,GO:0006163,GO:0006195,GO:0006200,GO:0006725,GO:0006753,GO:0006793,GO:0006796,GO:0006807,GO:0006810,GO:0006928,GO:0006996,GO:0007017,GO:0007018,GO:0007275,GO:0007276,GO:0007281,GO:0007283,GO:0007286,GO:0007291,GO:0007317,GO:0007349,GO:0007350,GO:0007351,GO:0007389,GO:0007399,GO:0007423,GO:0008092,GO:0008150,GO:0008152,GO:0008595,GO:0009056,GO:0009116,GO:0009117,GO:0009119,GO:0009123,GO:0009125,GO:0009126,GO:0009128,GO:0009141,GO:0009143,GO:0009144,GO:0009146,GO:0009150,GO:0009154,GO:0009158,GO:0009161,GO:0009164,GO:0009166,GO:0009167,GO:0009169,GO:0009199,GO:0009203,GO:0009205,GO:0009207,GO:0009259,GO:0009261,GO:0009653,GO:0009790,GO:0009798,GO:0009880,GO:0009887,GO:0009888,GO:0009948,GO:0009952,GO:0009987,GO:0010970,GO:0015629,GO:0016043,GO:0016459,GO:0016461,GO:0016462,GO:0016787,GO:0016817,GO:0016818,GO:0016887,GO:0017022,GO:0017076,GO:0017111,GO:0019439,GO:0019637,GO:0019693,GO:0019953,GO:0022008,GO:0022412,GO:0022414,GO:0030029,GO:0030048,GO:0030154,GO:0030182,GO:0030554,GO:0030705,GO:0030855,GO:0030898,GO:0031475,GO:0032027,GO:0032501,GO:0032502,GO:0032504,GO:0032549,GO:0032550,GO:0032553,GO:0032555,GO:0032559,GO:0032879,GO:0032991,GO:0034641,GO:0034643,GO:0034655,GO:0035282,GO:0035639,GO:0036094,GO:0042051,GO:0042052,GO:0042278,GO:0042454,GO:0042461,GO:0042462,GO:0042623,GO:0043167,GO:0043168,GO:0043226,GO:0043228,GO:0043229,GO:0043232,GO:0043234,GO:0044087,GO:0044237,GO:0044238,GO:0044248,GO:0044270,GO:0044281,GO:0044422,GO:0044424,GO:0044430,GO:0044446,GO:0044464,GO:0044699,GO:0044702,GO:0044707,GO:0044710,GO:0044763,GO:0044765,GO:0044767,GO:0045595,GO:0045856,GO:0045995,GO:0046034,GO:0046128,GO:0046130,GO:0046434,GO:0046483,GO:0046530,GO:0046700,GO:0046907,GO:0047497,GO:0048232,GO:0048468,GO:0048513,GO:0048515,GO:0048518,GO:0048522,GO:0048592,GO:0048609,GO:0048610,GO:0048646,GO:0048666,GO:0048699,GO:0048731,GO:0048749,GO:0048856,GO:0048869,GO:0050789,GO:0050793,GO:0050794,GO:0051128,GO:0051179,GO:0051234,GO:0051239,GO:0051640,GO:0051641,GO:0051646,GO:0051649,GO:0051654,GO:0051656,GO:0055086,GO:0060281,GO:0060284,GO:0060341,GO:0060429,GO:0065007,GO:0071704,GO:0071840,GO:0072384,GO:0072521,GO:0072523,GO:0097159,GO:1901135,GO:1901136,GO:1901265,GO:1901292,GO:1901360,GO:1901361,GO:1901363,GO:1901564,GO:1901565,GO:1901575,GO:1901657,GO:1901658,GO:2000026,GO:2000241,GO:2000243 | MYSc |

| | | |
|---|---|---|
| comp220769_c1_seq17.p2 | GO:0000146,GO:0001508,GO:0001676,GO:0001726,GO:0001750,GO:0001942,GO:0002065,GO:0002790,GO:0003001,GO:0003008,GO:0003674,GO:0003774,GO:0003779,GO:0003824,GO:0005488,GO:0005509,GO:0005515,GO:0005516,GO:0005575,GO:0005622,GO:0005623,GO:0005737,GO:0005794,GO:0005829,GO:0005856,GO:0005874,GO:0005882,GO:0005929,GO:0006082,GO:0006582,GO:0006629,GO:0006631,GO:0006633,GO:0006725,GO:0006810,GO:0006887,GO:0006928,GO:0007009,GO:0007154,GO:0007267,GO:0007268,GO:0007272,GO:0007275,GO:0007399,GO:0007600,GO:0007601,GO:0007610,GO:0007626,GO:0008092,GO:0008104,GO:0008150,GO:0008152,GO:0008366,GO:0008544,GO:0008610,GO:0009058,GO:0009653,GO:0009719,GO:0009725,GO:0009887,GO:0009888,GO:0009914,GO:0009966,GO:0009987,GO:0010033,GO:0010243,GO:0010469,GO:0010646,GO:0010817,GO:0010959,GO:0015629,GO:0015630,GO:0015833,GO:0016023,GO:0016043,GO:0016044,GO:0016053,GO:0016192,GO:0016459,GO:0016462,GO:0016787,GO:0016817,GO:0016818,GO:0017111,GO:0018958,GO:0019226,GO:0019228,GO:0019438,GO:0019748,GO:0019752,GO:0021700,GO:0022404,GO:0022405,GO:0022898,GO:0023051,GO:0023052,GO:0023061,GO:0030029,GO:0030048,GO:0030050,GO:0030072,GO:0030073,GO:0030141,GO:0030154,GO:0030318,GO:0030426,GO:0030427,GO:0030855,GO:0031252,GO:0031410,GO:0031513,GO:0031585,GO:0031982,GO:0031987,GO:0031988,GO:0032252,GO:0032400,GO:0032401,GO:0032402,GO:0032409,GO:0032412,GO:0032501,GO:0032502,GO:0032593,GO:0032787,GO:0032868,GO:0032869,GO:0032870,GO:0032879,GO:0032940,GO:0032991,GO:0033036,GO:0033059,GO:0034613,GO:0034762,GO:0034765,GO:0035371,GO:0035637,GO:0040007,GO:0040011,GO:0042221,GO:0042303,GO:0042391,GO:0042438,GO:0042440,GO:0042470,GO:0042476,GO:0042552,GO:0042633,GO:0042640,GO:0042641,GO:0042759,GO:0042886,GO:0042995,GO:0043005,GO:0043025,GO:0043167,GO:0043169,GO:0043226,GO:0043227,GO:0043228,GO:0043229,GO:0043231,GO:0043232,GO:0043234,GO:0043269,GO:0043434,GO:0043436,GO:0043473,GO:0043588,GO:0044237,GO:0044238,GO:0044249,GO:0044255,GO:0044281,GO:0044283,GO:0044297,GO:0044422,GO:0044424,GO:0044430,GO:0044444,GO:0044446,GO:0044463,GO:0044464,GO:0044550,GO:0044699,GO:0044700,GO:0044707,GO:0044708,GO:0044710,GO:0044711,GO:0044763,GO:0044765,GO:0044767,GO:0045111,GO:0046148,GO:0046189,GO:0046394,GO:0046872,GO:0046879,GO:0046903,GO:0048066,GO:0048513,GO:0048583,GO:0048589,GO:0048731,GO:0048770,GO:0048820,GO:0048856,GO:0048869,GO:0050789,GO:0050794,GO:0050808,GO:0050877,GO:0050896,GO:0050931,GO:0050953,GO:0051049,GO:0051179,GO:0051234,GO:0051640,GO:0051641,GO:0051643,GO:0051648,GO:0051649,GO:0051650,GO:0051656,GO:0051716,GO:0051875,GO:0051904,GO:0051905,GO:0051924,GO:0060429,GO:0060563,GO:0061024,GO:0065007,GO:0065008,GO:0065009,GO:0070727,GO:0070887,GO:0071310,GO:0071375,GO:0071417,GO:0071495,GO:0071695,GO:0071702,GO:0071704,GO:0071705,GO:0071840,GO:0072330,GO:0072372,GO:0072657,GO:0072659,GO:0097458,GO:1901019,GO:1901360,GO:1901362,GO:1901576,GO:1901615,GO:1901617,GO:1901652,GO:1901653,GO:1901698,GO:1901699,GO:1901700,GO:1901701,GO:2001257 | myosin VA (heavy chain 12, myoxin) |
| comp208980_c2_seq2.p1 | | Inherit from NOG: Chromosome 11 open reading frame 63 |
| comp217101_c1_seq1.p1 | GO:0001817,GO:0001818,GO:0002376,GO:0003674,GO:0005488,GO:0005515,GO:0005575,GO:0005622,GO:0005623,GO:0005737,GO:0005829,GO:0006355,GO:0006950,GO:0006952,GO:0006955,GO:0008150,GO:0009889,GO:0010468,GO:0010556,GO:0010941,GO:0019219,GO:0019222,GO:0019899,GO:0019900,GO:0031323,GO:0031326,GO:0032088,GO:0032182,GO:0032479,GO:0032480,GO:0042981,GO:0043066,GO:0043067,GO:0043069,GO:0043130,GO:0043433,GO:0044092,GO:0044424,GO:0044444,GO:0044464,GO:0045087,GO:0048519,GO:0048523,GO:0050789,GO:0050794,GO:0050896,GO:0051090,GO:0051171,GO:0051239,GO:0051241,GO:0051252,GO:0060255,GO:0060548,GO:0065007,GO:0065009,GO:0080090,GO:2000112,GO:2001141 | Tax1 (human T-cell leukemia virus type I) binding protein 1 |

| | | |
|---|---|---|
| comp201466_c0_seq1.p2 | GO:0005575,GO:0005622,GO:0005623,GO:0005634,GO:0005654,GO:0006325,GO:0006464,GO:0006479,GO:0006996,GO:0008150,GO:0008152,GO:0008213,GO:0009987,GO:0016043,GO:0016568,GO:0016569,GO:0016570,GO:0016571,GO:0019538,GO:0031974,GO:0031981,GO:0032259,GO:0032991,GO:0034708,GO:0034968,GO:0035097,GO:0036211,GO:0043170,GO:0043226,GO:0043227,GO:0043229,GO:0043231,GO:0043233,GO:0043234,GO:0043412,GO:0043414,GO:0044237,GO:0044238,GO:0044260,GO:0044267,GO:0044422,GO:0044424,GO:0044428,GO:0044446,GO:0044451,GO:0044464,GO:0044666,GO:0044699,GO:0044763,GO:0051276,GO:0051568,GO:0070013,GO:0071704,GO:0071840 | Inherit from biNOG: Nuclear receptor coactivator 6 |
| comp219873_c0_seq6.p1 | | Inherit from biNOG: SOGA family member |
| comp206293_c0_seq1.p1 | GO:0000323,GO:0002376,GO:0002520,GO:0003674,GO:0003824,GO:0004842,GO:0005575,GO:0005622,GO:0005623,GO:0005737,GO:0005764,GO:0005765,GO:0005768,GO:0005769,GO:0005773,GO:0005774,GO:0005783,GO:0005789,GO:0006464,GO:0007275,GO:0008150,GO:0008152,GO:0009790,GO:0009792,GO:0009987,GO:0010008,GO:0012505,GO:0012506,GO:0016020,GO:0016023,GO:0016567,GO:0016874,GO:0016879,GO:0016881,GO:0019538,GO:0019787,GO:0030097,GO:0030659,GO:0031090,GO:0031410,GO:0031901,GO:0031982,GO:0031988,GO:0032446,GO:0032501,GO:0032502,GO:0036211,GO:0042175,GO:0043170,GO:0043226,GO:0043227,GO:0043229,GO:0043231,GO:0043412,GO:0044237,GO:0044238,GO:0044260,GO:0044267,GO:0044422,GO:0044424,GO:0044425,GO:0044432,GO:0044433,GO:0044437,GO:0044440,GO:0044444,GO:0044446,GO:0044464,GO:0044699,GO:0044707,GO:0044767,GO:0048513,GO:0048534,GO:0048731,GO:0048856,GO:0070647,GO:0071704 | membrane-associated ring finger (C3HC4) |
| comp207317_c0_seq4.p1 | | peptide-methionine (R)-S-oxide reductase |
| comp213245_c0_seq1.p1 | | Patatin-like phospholipase |
| comp214535_c0_seq4.p1 | | RING |

| | | |
|---|---|---|
| comp213622_c0_seq1.p1 | GO:0000003,GO:0000075,GO:0000077,GO:0000122,GO:0000715,GO:0000781,GO:0000988,GO:0000989,GO:0001085,GO:0001654,GO:0001817,GO:0001818,GO:0001959,GO:0001960,GO:0002376,GO:0002520,GO:0002682,GO:0003006,GO:0003674,GO:0003676,GO:0003677,GO:0003712,GO:0003713,GO:0003714,GO:0003824,GO:0005102,GO:0005488,GO:0005515,GO:0005575,GO:0005622,GO:0005623,GO:0005634,GO:0005652,GO:0005654,GO:0005694,GO:0005700,GO:0005737,GO:0006139,GO:0006259,GO:0006281,GO:0006289,GO:0006323,GO:0006351,GO:0006355,GO:0006357,GO:0006464,GO:0006725,GO:0006807,GO:0006950,GO:0006952,GO:0006955,GO:0006974,GO:0006996,GO:0007049,GO:0007093,GO:0007095,GO:0007154,GO:0007165,GO:0007166,GO:0007243,GO:0007259,GO:0007275,GO:0007276,GO:0007283,GO:0007346,GO:0007399,GO:0007423,GO:0007444,GO:0007446,GO:0007548,GO:0008022,GO:0008134,GO:0008150,GO:0008152,GO:0008270,GO:0009058,GO:0009059,GO:0009653,GO:0009719,GO:0009725,GO:0009790,GO:0009792,GO:0009889,GO:0009890,GO:0009891,GO:0009892,GO:0009893,GO:0009894,GO:0009966,GO:0009968,GO:0009987,GO:0010033,GO:0010171,GO:0010467,GO:0010468,GO:0010556,GO:0010557,GO:0010558,GO:0010564,GO:0010604,GO:0010605,GO:0010627,GO:0010628,GO:0010629,GO:0010646,GO:0010648,GO:0010720,GO:0010741,GO:0010769,GO:0010941,GO:0010942,GO:0010948,GO:0010975,GO:0014070,GO:0015459,GO:0016043,GO:0016070,GO:0016247,GO:0016363,GO:0016604,GO:0016605,GO:0016607,GO:0016874,GO:0016879,GO:0016881,GO:0016925,GO:0017151,GO:0018130,GO:0019219,GO:0019221,GO:0019222,GO:0019438,GO:0019538,GO:0019787,GO:0019789,GO:0019899,GO:0019953,GO:0022008,GO:0022402,GO:0022414,GO:0022603,GO:0022604,GO:0022607,GO:0023051,GO:0023052,GO:0023057,GO:0030097,GO:0030154,GO:0030261,GO:0030278,GO:0030425,GO:0030518,GO:0030521,GO:0030522,GO:0031323,GO:0031324,GO:0031325,GO:0031326,GO:0031327,GO:0031328,GO:0031344,GO:0031346,GO:0031347,GO:0031399,GO:0031401,GO:0031570,GO:0031572,GO:0031974,GO:0031981,GO:0032088,GO:0032268,GO:0032270,GO:0032446,GO:0032479,GO:0032480,GO:0032501,GO:0032502,GO:0032504,GO:0032774,GO:0033143,GO:0033144,GO:0033233,GO:0033235,GO:0033554,GO:0033574,GO:0033993,GO:0034097,GO:0034341,GO:0034399,GO:0034622,GO:0034641,GO:0034645,GO:0034654,GO:0035012,GO:0035257,GO:0035258,GO:0035265,GO:0035556,GO:0036211,GO:0040007,GO:0040008,GO:0040009,GO:0040010,GO:0040014,GO:0040018,GO:0040035,GO:0042176,GO:0042221,GO:0042391,GO:0042981,GO:0042995,GO:0043005,GO:0043065,GO:0043067,GO:0043068,GO:0043167,GO:0043169,GO:0043170,GO:0043226,GO:0043227,GO:0043228,GO:0043229,GO:0043231,GO:0043232,GO:0043233,GO:0043412,GO:0043433,GO:0043933,GO:0044085,GO:0044092,GO:0044237,GO:0044238,GO:0044249,GO:0044260,GO:0044267,GO:0044271,GO:0044422,GO:0044424,GO:0044427,GO:0044428,GO:0044446,GO:0044451,GO:0044464,GO:0044699,GO:0044700,GO:0044702,GO:0044707,GO:0044763,GO:0044767,GO:0044773,GO:0044774,GO:0045087,GO:0045088,GO:0045202,GO:0045595,GO:0045597,GO:0045664,GO:0045667,GO:0045838,GO:0045892,GO:0045893,GO:0045927,GO:0045934,GO:0045935,GO:0045944,GO:0046425,GO:0046426,GO:0046483,GO:0046872,GO:0046914,GO:0047485,GO:0048232,GO:0048513,GO:0048518,GO:0048519,GO:0048522,GO:0048523,GO:0048534,GO:0048545,GO:0048583,GO:0048585,GO:0048589,GO:0048608,GO:0048609,GO:0048699,GO:0048731,GO:0048749,GO:0048806,GO:0048814,GO:0048856,GO:0048869,GO:0050681,GO:0050767,GO:0050769,GO:0050773,GO:0050775,GO:0050776,GO:0050789,GO:0050793,GO:0050794,GO:0050896,GO:0051090,GO:0051094,GO:0051128,GO:0051130,GO:0051147,GO:0051149,GO:0051150,GO:0051152,GO:0051171,GO:0051172,GO:0051173,GO:0051239,GO:0051240,GO:0051241,GO:0051246,GO:0051247,GO:0051252,GO:0051253,GO:0051254,GO:0051276,GO:0051427,GO:0051716,GO:0051726,GO:0051960,GO:0060255,GO:0060284,GO:0060330,GO:0060333,GO:0060334,GO:0060338,GO:0060339,GO:0060759,GO:0060761,GO:0060765,GO:0060766,GO:0061458,GO:0065003,GO:0065004,GO:0065007,GO:0065008,GO:0065009,GO:0070013,GO:0070647,GO:0070887,GO:0071103,GO:0071310,GO:0071345,GO:0071346,GO:0071704,GO:0071824,GO:0071840,GO:0080090,GO:0080134,GO:0090304,GO:0097159,GO:0097305,GO:0097458,GO:1901360,GO:1901362,GO:1901363,GO:1901576,GO:1901654,GO:1901700,GO:1901987,GO:1901988,GO:1901990,GO:1901991,GO:2000026,GO:2000112,GO:2000113,GO:2001141 | Protein inhibitor of activated STAT |

| | | |
|---|---|---|
| comp213622_c0_seq1.p2 | GO:0000139,GO:0005575,GO:0005622,GO:0005623,GO:0005737,GO:0005794,GO:0005798,GO:0005829,GO:0006810,GO:0006890,GO:0006891,GO:0006900,GO:0006901,GO:0006903,GO:0006996,GO:0008150,GO:0009987,GO:0012505,GO:0012506,GO:0016020,GO:0016023,GO:0016043,GO:0016044,GO:0016050,GO:0016192,GO:0030117,GO:0030120,GO:0030126,GO:0030135,GO:0030137,GO:0030659,GO:0030660,GO:0030662,GO:0030663,GO:0031090,GO:0031410,GO:0031982,GO:0031988,GO:0032991,GO:0035964,GO:0043226,GO:0043227,GO:0043229,GO:0043231,GO:0043234,GO:0044422,GO:0044424,GO:0044425,GO:0044431,GO:0044433,GO:0044444,GO:0044446,GO:0044464,GO:0044699,GO:0044763,GO:0044765,GO:0046907,GO:0048193,GO:0048194,GO:0048199,GO:0048200,GO:0048205,GO:0048475,GO:0051179,GO:0051234,GO:0051640,GO:0051641,GO:0051648,GO:0051649,GO:0051650,GO:0051656,GO:0061024,GO:0071840 | Coatomer protein complex, subunit epsilon |
| comp220930_c3_seq1.p1 | GO:0002376,GO:0002520,GO:0003674,GO:0003824,GO:0004842,GO:0006464,GO:0007275,GO:0008150,GO:0008152,GO:0009790,GO:0009987,GO:0016567,GO:0016874,GO:0016879,GO:0016881,GO:0019538,GO:0019787,GO:0030097,GO:0030099,GO:0030154,GO:0032446,GO:0032501,GO:0032502,GO:0035162,GO:0036211,GO:0043170,GO:0043412,GO:0044237,GO:0044238,GO:0044260,GO:0044267,GO:0044699,GO:0044707,GO:0044763,GO:0044767,GO:0048513,GO:0048534,GO:0048568,GO:0048731,GO:0048856,GO:0048869,GO:0060215,GO:0070647,GO:0071704 | tripartite motif containing |
| comp148580_c0_seq1.p1 | | coiled-coil domain containing 176 |
| comp207582_c0_seq2.p1 | GO:0000166,GO:0001775,GO:0001882,GO:0001883,GO:0002200,GO:0002204,GO:0002208,GO:0002250,GO:0002252,GO:0002263,GO:0002285,GO:0002312,GO:0002366,GO:0002376,GO:0002377,GO:0002381,GO:0002440,GO:0002443,GO:0002449,GO:0002460,GO:0002520,GO:0002562,GO:0003674,GO:0003676,GO:0003677,GO:0005488,GO:0005524,GO:0005575,GO:0005622,GO:0005623,GO:0005634,GO:0005737,GO:0005886,GO:0006139,GO:0006259,GO:0006310,GO:0006725,GO:0006807,GO:0006955,GO:0007088,GO:0007096,GO:0007275,GO:0007346,GO:0007399,GO:0008150,GO:0008152,GO:0009987,GO:0010564,GO:0010720,GO:0014013,GO:0014015,GO:0016020,GO:0016064,GO:0016444,GO:0016445,GO:0016447,GO:0017076,GO:0019724,GO:0022008,GO:0030027,GO:0030154,GO:0030554,GO:0031252,GO:0032501,GO:0032502,GO:0032549,GO:0032550,GO:0032553,GO:0032555,GO:0032559,GO:0033043,GO:0034641,GO:0035639,GO:0036094,GO:0042113,GO:0042995,GO:0043167,GO:0043168,GO:0043170,GO:0043226,GO:0043227,GO:0043229,GO:0043231,GO:0044237,GO:0044238,GO:0044260,GO:0044424,GO:0044464,GO:0044699,GO:0044707,GO:0044763,GO:0045190,GO:0045321,GO:0045595,GO:0045597,GO:0045685,GO:0045687,GO:0046483,GO:0046649,GO:0048518,GO:0048522,GO:0048699,GO:0048713,GO:0048714,GO:0048731,GO:0048856,GO:0048869,GO:0050767,GO:0050769,GO:0050789,GO:0050793,GO:0050794,GO:0050896,GO:0051094,GO:0051128,GO:0051239,GO:0051726,GO:0051783,GO:0051960,GO:0060284,GO:0065007,GO:0071704,GO:0071944,GO:0090304,GO:0097159,GO:1901265,GO:1901360,GO:1901363,GO:2000026 | SWAP switching B-cell complex 70kDa subunit |
| comp220682_c0_seq1.p1 | | ring finger protein 17 |
| comp212671_c1_seq2.p1 | GO:0000003,GO:0000323,GO:0003674,GO:0003824,GO:0004518,GO:0004519,GO:0004521,GO:0004540,GO:0005575,GO:0005576,GO:0005615,GO:0005622,GO:0005623,GO:0005737,GO:0005764,GO:0005773,GO:0005775,GO:0005783,GO:0005788,GO:0006139,GO:0006401,GO:0006725,GO:0006807,GO:0008150,GO:0008152,GO:0009056,GO:0009057,GO:0009987,GO:0016070,GO:0016072,GO:0016075,GO:0016787,GO:0016788,GO:0019439,GO:0031974,GO:0032501,GO:0032504,GO:0034641,GO:0034655,GO:0034660,GO:0034661,GO:0043170,GO:0043202,GO:0043226,GO:0043227,GO:0043229,GO:0043231,GO:0043233,GO:0044237,GO:0044238,GO:0044248,GO:0044260,GO:0044265,GO:0044270,GO:0044421,GO:0044422,GO:0044424,GO:0044432,GO:0044437,GO:0044444,GO:0044446,GO:0044464,GO:0046483,GO:0046700,GO:0070013,GO:0071704,GO:0090304,GO:0090305,GO:1901360,GO:1901361,GO:1901575 | ribonuclease T2 |
| comp215056_c0_seq5.p1 | | Neurotransmitter-gated ion-channel ligand binding domain |

| | | |
|---|---|---|
| comp211241_c0_seq12.p1 | | deafness, autosomal recessive |
| comp211241_c0_seq12.p2 | GO:0008150,GO:0008277,GO:0009966,GO:0010646,GO:0023051,GO:0048583,GO:0050789,GO:0050794,GO:0065007 | GGL domain |
| comp219343_c0_seq4.p1 | GO:0000003,GO:0000041,GO:0000220,GO:0000226,GO:0000278,GO:0001654,GO:0002119,GO:0002164,GO:0003674,GO:0003824,GO:0005215,GO:0005575,GO:0005622,GO:0005623,GO:0005737,GO:0005768,GO:0005773,GO:0005774,GO:0005886,GO:0005887,GO:0005889,GO:0006139,GO:0006152,GO:0006163,GO:0006195,GO:0006200,GO:0006725,GO:0006753,GO:0006793,GO:0006796,GO:0006807,GO:0006810,GO:0006811,GO:0006812,GO:0006818,GO:0006826,GO:0006873,GO:0006875,GO:0006879,GO:0006897,GO:0006898,GO:0006915,GO:0006996,GO:0007010,GO:0007017,GO:0007049,GO:0007051,GO:0007052,GO:0007154,GO:0007165,GO:0007166,GO:0007167,GO:0007169,GO:0007275,GO:0007423,GO:0008104,GO:0008150,GO:0008152,GO:0008219,GO:0008286,GO:0008324,GO:0008553,GO:0009056,GO:0009116,GO:0009117,GO:0009119,GO:0009123,GO:0009125,GO:0009126,GO:0009128,GO:0009141,GO:0009143,GO:0009144,GO:0009146,GO:0009150,GO:0009154,GO:0009158,GO:0009161,GO:0009164,GO:0009166,GO:0009167,GO:0009169,GO:0009199,GO:0009203,GO:0009205,GO:0009207,GO:0009259,GO:0009261,GO:0009719,GO:0009725,GO:0009790,GO:0009791,GO:0009792,GO:0009987,GO:0010008,GO:0010033,GO:0010243,GO:0012501,GO:0012506,GO:0015031,GO:0015075,GO:0015077,GO:0015078,GO:0015399,GO:0015405,GO:0015662,GO:0015672,GO:0015682,GO:0015988,GO:0015991,GO:0015992,GO:0016020,GO:0016021,GO:0016023,GO:0016043,GO:0016192,GO:0016265,GO:0016462,GO:0016469,GO:0016471,GO:0016787,GO:0016817,GO:0016818,GO:0016820,GO:0016887,GO:0017111,GO:0018996,GO:0019439,GO:0019637,GO:0019693,GO:0019725,GO:0019829,GO:0022402,GO:0022804,GO:0022857,GO:0022890,GO:0022891,GO:0022892,GO:0023052,GO:0030001,GO:0030003,GO:0030139,GO:0030659,GO:0030666,GO:0030670,GO:0031090,GO:0031224,GO:0031226,GO:0031410,GO:0031982,GO:0031988,GO:0032501,GO:0032502,GO:0032868,GO:0032869,GO:0032870,GO:0032991,GO:0033036,GO:0033176,GO:0033177,GO:0033179,GO:0033181,GO:0033572,GO:0034220,GO:0034641,GO:0034655,GO:0040011,GO:0042221,GO:0042278,GO:0042303,GO:0042454,GO:0042592,GO:0042623,GO:0042625,GO:0042626,GO:0043010,GO:0043226,GO:0043227,GO:0043229,GO:0043231,GO:0043234,GO:0043434,GO:0043473,GO:0043492,GO:0043933,GO:0044237,GO:0044238,GO:0044248,GO:0044270,GO:0044281,GO:0044403,GO:0044419,GO:0044422,GO:0044424,GO:0044425,GO:0044433,GO:0044437,GO:0044440,GO:0044444,GO:0044446,GO:0044459,GO:0044464,GO:0044699,GO:0044700,GO:0044707,GO:0044710,GO:0044763,GO:0044765,GO:0044767,GO:0045184,GO:0045335,GO:0046034,GO:0046128,GO:0046130,GO:0046434,GO:0046483,GO:0046700,GO:0046916,GO:0048066,GO:0048513,GO:0048731,GO:0048856,GO:0048878,GO:0050789,GO:0050794,GO:0050801,GO:0050896,GO:0051179,GO:0051234,GO:0051701,GO:0051704,GO:0051716,GO:0055065,GO:0055072,GO:0055076,GO:0055080,GO:0055082,GO:0055085,GO:0055086,GO:0060041,GO:0065007,GO:0065008,GO:0070887,GO:0071310,GO:0071375,GO:0071417,GO:0071495,GO:0071702,GO:0071704,GO:0071822,GO:0071840,GO:0071944,GO:0072512,GO:0072521,GO:0072523,GO:0090382,GO:0090533,GO:1901135,GO:1901136,GO:1901292,GO:1901360,GO:1901361,GO:1901564,GO:1901565,GO:1901575,GO:1901652,GO:1901653,GO:1901657,GO:1901658,GO:1901698,GO:1901699,GO:1901700,GO:1901701 | ATPase, H transporting, lysosomal 21kDa, V0 subunit b |
| comp210861_c0_seq1.p1 | | piggyBac transposable element derived 4 |

| | | |
|---|---|---|
| comp220533_c0_seq1.p1 | GO:0000003,GO:0000184,GO:0000785,GO:0000932,GO:0000956,GO:0001666,GO:0003006,GO:0003674,GO:0003676,GO:0003682,GO:0003723,GO:0003724,GO:0003824,GO:0004004,GO:0004386,GO:0005488,GO:0005515,GO:0005575,GO:0005622,GO:0005623,GO:0005634,GO:0005681,GO:0005694,GO:0005737,GO:0005829,GO:0006139,GO:0006152,GO:0006163,GO:0006195,GO:0006200,GO:0006259,GO:0006260,GO:0006281,GO:0006401,GO:0006402,GO:0006403,GO:0006405,GO:0006406,GO:0006417,GO:0006449,GO:0006725,GO:0006753,GO:0006793,GO:0006796,GO:0006807,GO:0006810,GO:0006913,GO:0006950,GO:0006974,GO:0007049,GO:0007275,GO:0007548,GO:0007549,GO:0008026,GO:0008150,GO:0008152,GO:0008186,GO:0008334,GO:0009048,GO:0009056,GO:0009057,GO:0009058,GO:0009059,GO:0009116,GO:0009117,GO:0009119,GO:0009123,GO:0009125,GO:0009126,GO:0009128,GO:0009141,GO:0009143,GO:0009144,GO:0009146,GO:0009150,GO:0009154,GO:0009158,GO:0009161,GO:0009164,GO:0009166,GO:0009167,GO:0009169,GO:0009199,GO:0009203,GO:0009205,GO:0009207,GO:0009259,GO:0009261,GO:0009628,GO:0009653,GO:0009790,GO:0009887,GO:0009889,GO:0009892,GO:0009893,GO:0009894,GO:0009896,GO:0009987,GO:0010467,GO:0010468,GO:0010556,GO:0010604,GO:0010605,GO:0010608,GO:0010629,GO:0015931,GO:0016070,GO:0016071,GO:0016246,GO:0016441,GO:0016458,GO:0016462,GO:0016482,GO:0016787,GO:0016817,GO:0016818,GO:0016887,GO:0017111,GO:0019219,GO:0019222,GO:0019439,GO:0019637,GO:0019693,GO:0019899,GO:0019902,GO:0019903,GO:0022414,GO:0030529,GO:0030538,GO:0031047,GO:0031323,GO:0031325,GO:0031326,GO:0031329,GO:0031331,GO:0032268,GO:0032501,GO:0032502,GO:0032991,GO:0033036,GO:0033554,GO:0034641,GO:0034645,GO:0034655,GO:0035112,GO:0035145,GO:0035194,GO:0035770,GO:0036293,GO:0036294,GO:0040029,GO:0040035,GO:0042221,GO:0042278,GO:0042454,GO:0042623,GO:0043170,GO:0043226,GO:0043227,GO:0043228,GO:0043229,GO:0043231,GO:0043232,GO:0043234,GO:0043244,GO:0044237,GO:0044238,GO:0044248,GO:0044249,GO:0044260,GO:0044265,GO:0044270,GO:0044281,GO:0044422,GO:0044424,GO:0044427,GO:0044428,GO:0044444,GO:0044446,GO:0044464,GO:0044530,GO:0044699,GO:0044707,GO:0044710,GO:0044763,GO:0044767,GO:0045935,GO:0046034,GO:0046128,GO:0046130,GO:0046434,GO:0046483,GO:0046700,GO:0046907,GO:0048513,GO:0048518,GO:0048519,GO:0048522,GO:0048562,GO:0048568,GO:0048598,GO:0048608,GO:0048731,GO:0048806,GO:0048856,GO:0050657,GO:0050658,GO:0050789,GO:0050794,GO:0050896,GO:0051028,GO:0051128,GO:0051168,GO:0051169,GO:0051171,GO:0051173,GO:0051179,GO:0051234,GO:0051236,GO:0051246,GO:0051252,GO:0051254,GO:0051641,GO:0051649,GO:0051716,GO:0051721,GO:0055086,GO:0060255,GO:0061013,GO:0061014,GO:0061458,GO:0065007,GO:0070035,GO:0070482,GO:0070887,GO:0071044,GO:0071453,GO:0071456,GO:0071702,GO:0071704,GO:0071705,GO:0072521,GO:0072523,GO:0080090,GO:0090304,GO:0097159,GO:1901135,GO:1901136,GO:1901292,GO:1901360,GO:1901361,GO:1901363,GO:1901564,GO:1901565,GO:1901575,GO:1901576,GO:1901657,GO:1901658,GO:2000112,GO:2000622,GO:2000624 | UPF1 regulator of nonsense transcripts homolog (Yeast) |

| | | |
|---|---|---|
| comp220533_c0_seq1.p2 | GO:0000166,GO:0000271,GO:0003674,GO:0003824,GO:0003983,GO:0005488,GO:0005536,GO:0005575,GO:0005622,GO:0005623,GO:0005737,GO:0005829,GO:0005975,GO:0005976,GO:0005977,GO:0005978,GO:0005996,GO:0006006,GO:0006011,GO:0006065,GO:0006073,GO:0006082,GO:0006091,GO:0006112,GO:0006139,GO:0006725,GO:0006753,GO:0006793,GO:0006796,GO:0006805,GO:0006807,GO:0008150,GO:0008152,GO:0009058,GO:0009059,GO:0009117,GO:0009165,GO:0009225,GO:0009226,GO:0009250,GO:0009410,GO:0009987,GO:0015980,GO:0016051,GO:0016053,GO:0016310,GO:0016740,GO:0016772,GO:0016779,GO:0018130,GO:0019103,GO:0019255,GO:0019318,GO:0019438,GO:0019637,GO:0019752,GO:0030246,GO:0032553,GO:0032557,GO:0033692,GO:0034637,GO:0034641,GO:0034645,GO:0034654,GO:0036094,GO:0042221,GO:0043170,GO:0043436,GO:0044042,GO:0044237,GO:0044238,GO:0044249,GO:0044260,GO:0044262,GO:0044264,GO:0044271,GO:0044281,GO:0044283,GO:0044424,GO:0044444,GO:0044464,GO:0044699,GO:0044710,GO:0044711,GO:0044723,GO:0044763,GO:0046394,GO:0046398,GO:0046483,GO:0048029,GO:0050896,GO:0051716,GO:0051748,GO:0055086,GO:0055114,GO:0070569,GO:0070887,GO:0071466,GO:0071704,GO:0090407,GO:0097159,GO:1901135,GO:1901137,GO:1901265,GO:1901293,GO:1901360,GO:1901362,GO:1901363,GO:1901576 | UDP-glucose pyrophosphorylase 2 |
| comp220533_c0_seq1.p7 | GO:0003008,GO:0003674,GO:0003824,GO:0004356,GO:0005575,GO:0005622,GO:0005623,GO:0005737,GO:0006082,GO:0006520,GO:0006536,GO:0006538,GO:0006541,GO:0006542,GO:0006807,GO:0007154,GO:0007267,GO:0007268,GO:0007275,GO:0007399,GO:0007416,GO:0008150,GO:0008152,GO:0008652,GO:0009056,GO:0009058,GO:0009063,GO:0009064,GO:0009065,GO:0009084,GO:0009987,GO:0016043,GO:0016053,GO:0016054,GO:0016211,GO:0016874,GO:0016879,GO:0016880,GO:0019226,GO:0019752,GO:0022607,GO:0023052,GO:0032501,GO:0032502,GO:0035637,GO:0043112,GO:0043170,GO:0043436,GO:0043648,GO:0043649,GO:0044085,GO:0044237,GO:0044238,GO:0044248,GO:0044249,GO:0044260,GO:0044281,GO:0044282,GO:0044283,GO:0044424,GO:0044464,GO:0044699,GO:0044700,GO:0044707,GO:0044710,GO:0044711,GO:0044712,GO:0044763,GO:0045213,GO:0046394,GO:0046395,GO:0048731,GO:0048856,GO:0050808,GO:0050877,GO:0071704,GO:0071840,GO:1901564,GO:1901565,GO:1901566,GO:1901575,GO:1901576,GO:1901605,GO:1901606,GO:1901607 | glutamine synthetase |
| comp218230_c0_seq2.p1 | GO:0001775,GO:0003674,GO:0003824,GO:0004143,GO:0005488,GO:0005515,GO:0005575,GO:0005622,GO:0005623,GO:0005634,GO:0005635,GO:0005737,GO:0005768,GO:0005886,GO:0006461,GO:0006629,GO:0006638,GO:0006639,GO:0006793,GO:0006796,GO:0006950,GO:0006979,GO:0007154,GO:0007165,GO:0007166,GO:0007167,GO:0007169,GO:0007173,GO:0007186,GO:0007205,GO:0007275,GO:0007596,GO:0007599,GO:0008150,GO:0008152,GO:0008289,GO:0009611,GO:0009987,GO:0010033,GO:0012505,GO:0016020,GO:0016023,GO:0016043,GO:0016049,GO:0016301,GO:0016310,GO:0016740,GO:0016772,GO:0016773,GO:0019932,GO:0019992,GO:0022607,GO:0023052,GO:0030168,GO:0031090,GO:0031410,GO:0031965,GO:0031967,GO:0031975,GO:0031982,GO:0031988,GO:0032501,GO:0032502,GO:0035556,GO:0038127,GO:0040007,GO:0042060,GO:0042221,GO:0042802,GO:0042803,GO:0043226,GO:0043227,GO:0043229,GO:0043231,GO:0043933,GO:0044085,GO:0044237,GO:0044238,GO:0044255,GO:0044422,GO:0044424,GO:0044428,GO:0044444,GO:0044446,GO:0044464,GO:0044699,GO:0044700,GO:0044707,GO:0044710,GO:0044763,GO:0046339,GO:0046486,GO:0046982,GO:0046983,GO:0050789,GO:0050794,GO:0050817,GO:0050878,GO:0050896,GO:0051259,GO:0051260,GO:0051716,GO:0065003,GO:0065007,GO:0065008,GO:0070271,GO:0071704,GO:0071822,GO:0071840,GO:0071944 | diacylglycerol kinase |
| comp219923_c1_seq2.p1 | | ZnF_C2H2 |
| comp221252_c1_seq1.p1 | | SCAN domain |
| comp221252_c1_seq1.p2 | | Inherit from opiNOG: to reverse transcriptase |

| | | |
|---|---|---|
| comp216702_c3_seq2.p1 | | Low-density lipoprotein receptor repeat class B |
| comp211445_c0_seq1.p1 | GO:0000139,GO:0005575,GO:0005622,GO:0005623,GO:0005634,GO:0005737,GO:0005768,GO:0005794,GO:0005795,GO:0005796,GO:0005801,GO:0010008,GO:0012505,GO:0016020,GO:0016021,GO:0016023,GO:0030139,GO:0031090,GO:0031224,GO:0031410,GO:0031974,GO:0031982,GO:0031984,GO:0031985,GO:0031988,GO:0032580,GO:0043226,GO:0043227,GO:0043229,GO:0043231,GO:0043233,GO:0044422,GO:0044424,GO:0044425,GO:0044431,GO:0044440,GO:0044444,GO:0044446,GO:0044464,GO:0070013 | golgi integral membrane protein 4 |
| comp220720_c0_seq1.p1 | | Tudor domain |
| comp196691_c1_seq2.p1 | GO:0005575,GO:0005623,GO:0005886,GO:0016020,GO:0016324,GO:0044425,GO:0044459,GO:0044464,GO:0045177,GO:0071944 | kiaa1919 |
| comp208523_c0_seq4.p1 | GO:0005575,GO:0016020 | Zinc finger, DHHC-type containing 14 |
| comp206442_c0_seq2.p1 | GO:0000003,GO:0000165,GO:0000186,GO:0000187,GO:0000902,GO:0001700,GO:0001932,GO:0001934,GO:0002009,GO:0002165,GO:0003006,GO:0003674,GO:0003824,GO:0004672,GO:0004674,GO:0004702,GO:0004706,GO:0004709,GO:0004712,GO:0004871,GO:0005057,GO:0005488,GO:0005515,GO:0006464,GO:0006468,GO:0006793,GO:0006796,GO:0006950,GO:0007154,GO:0007165,GO:0007243,GO:0007254,GO:0007256,GO:0007257,GO:0007275,GO:0007276,GO:0007292,GO:0007304,GO:0007306,GO:0007391,GO:0007392,GO:0007394,GO:0007444,GO:0007483,GO:0007484,GO:0007485,GO:0007486,GO:0007548,GO:0007552,GO:0007560,GO:0008150,GO:0008152,GO:0008360,GO:0009266,GO:0009408,GO:0009628,GO:0009653,GO:0009790,GO:0009791,GO:0009792,GO:0009826,GO:0009886,GO:0009887,GO:0009888,GO:0009893,GO:0009966,GO:0009967,GO:0009987,GO:0010033,GO:0010562,GO:0010604,GO:0010627,GO:0010646,GO:0010647,GO:0010740,GO:0010927,GO:0016043,GO:0016049,GO:0016301,GO:0016310,GO:0016331,GO:0016476,GO:0016740,GO:0016772,GO:0016773,GO:0019220,GO:0019222,GO:0019538,GO:0019953,GO:0022412,GO:0022414,GO:0022603,GO:0022604,GO:0023014,GO:0023051,GO:0023052,GO:0023056,GO:0030154,GO:0030539,GO:0030540,GO:0030703,GO:0030707,GO:0031098,GO:0031323,GO:0031325,GO:0031399,GO:0031401,GO:0032147,GO:0032268,GO:0032270,GO:0032501,GO:0032502,GO:0032504,GO:0032872,GO:0032874,GO:0032989,GO:0033554,GO:0033674,GO:0035112,GO:0035126,GO:0035215,GO:0035556,GO:0036211,GO:0040007,GO:0042221,GO:0042325,GO:0042327,GO:0042802,GO:0042803,GO:0043085,GO:0043170,GO:0043405,GO:0043406,GO:0043408,GO:0043410,GO:0043412,GO:0043506,GO:0043507,GO:0043549,GO:0044093,GO:0044237,GO:0044238,GO:0044260,GO:0044267,GO:0044699,GO:0044700,GO:0044702,GO:0044707,GO:0044763,GO:0044767,GO:0045137,GO:0045471,GO:0045859,GO:0045860,GO:0045937,GO:0045995,GO:0046328,GO:0046330,GO:0046528,GO:0046529,GO:0046545,GO:0046546,GO:0046660,GO:0046661,GO:0046777,GO:0046843,GO:0046983,GO:0048468,GO:0048477,GO:0048513,GO:0048518,GO:0048522,GO:0048563,GO:0048569,GO:0048583,GO:0048584,GO:0048589,GO:0048598,GO:0048608,GO:0048609,GO:0048610,GO:0048646,GO:0048707,GO:0048729,GO:0048731,GO:0048803,GO:0048804,GO:0048805,GO:0048806,GO:0048807,GO:0048808,GO:0048856,GO:0048869,GO:0050789,GO:0050790,GO:0050793,GO:0050794,GO:0050896,GO:0051128,GO:0051174,GO:0051239,GO:0051246,GO:0051247,GO:0051338,GO:0051347,GO:0051403,GO:0051716,GO:0060089,GO:0060255,GO:0060429,GO:0060560,GO:0061458,GO:0065007,GO:0065008,GO:0065009,GO:0070302,GO:0070304,GO:0070887,GO:0071310,GO:0071361,GO:0071704,GO:0071840,GO:0071900,GO:0071902,GO:0080090,GO:0080134,GO:0080135,GO:0097305,GO:0097306,GO:1901700,GO:1901701,GO:2000026 | Protein tyrosine kinase |
| comp210218_c0_seq2.p1 | | Inherit from biNOG: ATP-binding cassette sub-family G (WHITE) member 2 |

| | | |
|---|---|---|
| comp220139_c3_seq4.p1 | GO:0003674,GO:0005488,GO:0005515,GO:0008150,GO:0010921,GO:0010923,GO:0019220,GO:0019222,GO:0019899,GO:0019902,GO:0031323,GO:0035303,GO:0043086,GO:0044092,GO:0050789,GO:0050790,GO:0050794,GO:0051174,GO:0051336,GO:0051346,GO:0065007,GO:0065009 | Protein phosphatase 1 regulatory subunit 37 |
| comp218657_c0_seq5.p1 | GO:0003674,GO:0005215,GO:0005575,GO:0005623,GO:0005886,GO:0006810,GO:0006811,GO:0006820,GO:0006821,GO:0008150,GO:0008509,GO:0009987,GO:0015075,GO:0015103,GO:0015108,GO:0015291,GO:0015297,GO:0015301,GO:0015698,GO:0015701,GO:0015711,GO:0016020,GO:0016323,GO:0016324,GO:0022804,GO:0022857,GO:0022891,GO:0022892,GO:0034220,GO:0044425,GO:0044459,GO:0044464,GO:0044699,GO:0044763,GO:0044765,GO:0045177,GO:0051179,GO:0051234,GO:0055085,GO:0071702,GO:0071944 | Solute carrier family 4 (Anion exchanger), member |
| comp218657_c0_seq5.p2 | GO:0003674,GO:0005215,GO:0005452,GO:0005488,GO:0005515,GO:0005575,GO:0005622,GO:0005623,GO:0005737,GO:0005856,GO:0005886,GO:0005887,GO:0005938,GO:0006810,GO:0006811,GO:0006820,GO:0006821,GO:0006873,GO:0006885,GO:0008150,GO:0008152,GO:0008509,GO:0009987,GO:0015075,GO:0015103,GO:0015108,GO:0015291,GO:0015297,GO:0015301,GO:0015698,GO:0015701,GO:0015711,GO:0016020,GO:0016021,GO:0016323,GO:0016324,GO:0019725,GO:0022804,GO:0022857,GO:0022891,GO:0022892,GO:0030003,GO:0030004,GO:0030016,GO:0030017,GO:0030018,GO:0030641,GO:0030863,GO:0031224,GO:0031226,GO:0031674,GO:0034220,GO:0042592,GO:0043226,GO:0043228,GO:0043229,GO:0043232,GO:0043292,GO:0043495,GO:0044281,GO:0044422,GO:0044424,GO:0044425,GO:0044444,GO:0044448,GO:0044449,GO:0044459,GO:0044464,GO:0044699,GO:0044710,GO:0044763,GO:0044765,GO:0045177,GO:0048878,GO:0050801,GO:0051179,GO:0051234,GO:0051453,GO:0055067,GO:0055080,GO:0055082,GO:0055085,GO:0065007,GO:0065008,GO:0071702,GO:0071944 | solute carrier family 4 |
| comp46753_c0_seq1.p1 | GO:0003674,GO:0005083,GO:0005085,GO:0005088,GO:0005089,GO:0005575,GO:0005622,GO:0005623,GO:0007275,GO:0007399,GO:0007417,GO:0007420,GO:0008064,GO:0008150,GO:0009987,GO:0016043,GO:0016319,GO:0019222,GO:0030234,GO:0030695,GO:0030832,GO:0030833,GO:0031344,GO:0031346,GO:0032271,GO:0032501,GO:0032502,GO:0032535,GO:0032956,GO:0032970,GO:0033043,GO:0043254,GO:0044087,GO:0044464,GO:0044699,GO:0044707,GO:0044763,GO:0044767,GO:0048513,GO:0048518,GO:0048522,GO:0048731,GO:0048856,GO:0050789,GO:0050790,GO:0050794,GO:0051128,GO:0051130,GO:0051493,GO:0060491,GO:0060589,GO:0065007,GO:0065008,GO:0065009,GO:0071840,GO:0090066,GO:1900027,GO:1900029 | RhoGEF domain |

| | | |
|---|---|---|
| comp220746_c1_seq15.p1 | GO:0000003,GO:0000902,GO:0000904,GO:0001654,GO:0001655,GO:0001700,GO:0001738,GO:0001745,GO:0001751,GO:0001754,GO:0001822,GO:0001894,GO:0001895,GO:0002009,GO:0002119,GO:0002164,GO:0003008,GO:0003014,GO:0003094,GO:0003674,GO:0005080,GO:0005488,GO:0005515,GO:0005575,GO:0005576,GO:0005622,GO:0005623,GO:0005737,GO:0005886,GO:0005911,GO:0005912,GO:0005913,GO:0005918,GO:0006810,GO:0006996,GO:0007043,GO:0007163,GO:0007275,GO:0007276,GO:0007292,GO:0007391,GO:0007399,GO:0007417,GO:0007420,GO:0007423,GO:0007424,GO:0007431,GO:0007435,GO:0007442,GO:0007443,GO:0008092,GO:0008104,GO:0008105,GO:0008150,GO:0008284,GO:0008593,GO:0009653,GO:0009790,GO:0009791,GO:0009792,GO:0009887,GO:0009888,GO:0009893,GO:0009966,GO:0009968,GO:0009987,GO:0010562,GO:0010646,GO:0010648,GO:0010927,GO:0016020,GO:0016021,GO:0016028,GO:0016043,GO:0016044,GO:0016324,GO:0016327,GO:0016331,GO:0016332,GO:0016333,GO:0016334,GO:0019220,GO:0019222,GO:0019899,GO:0019900,GO:0019901,GO:0019953,GO:0022008,GO:0022414,GO:0022607,GO:0022612,GO:0023051,GO:0023057,GO:0030010,GO:0030011,GO:0030030,GO:0030031,GO:0030054,GO:0030154,GO:0030182,GO:0030507,GO:0030855,GO:0030859,GO:0031224,GO:0031323,GO:0031325,GO:0032501,GO:0032502,GO:0032504,GO:0032835,GO:0032989,GO:0032990,GO:0032991,GO:0033036,GO:0034329,GO:0034330,GO:0034331,GO:0034332,GO:0034333,GO:0035002,GO:0035003,GO:0035088,GO:0035089,GO:0035090,GO:0035239,GO:0035272,GO:0035295,GO:0035330,GO:0040007,GO:0040008,GO:0040011,GO:0042044,GO:0042045,GO:0042051,GO:0042052,GO:0042127,GO:0042325,GO:0042327,GO:0042384,GO:0042461,GO:0042462,GO:0042592,GO:0042995,GO:0043010,GO:0043234,GO:0043296,GO:0043297,GO:0043954,GO:0044085,GO:0044424,GO:0044425,GO:0044459,GO:0044464,GO:0044699,GO:0044702,GO:0044707,GO:0044763,GO:0044765,GO:0044767,GO:0044782,GO:0045176,GO:0045177,GO:0045186,GO:0045197,GO:0045198,GO:0045216,GO:0045217,GO:0045218,GO:0045494,GO:0045570,GO:0045746,GO:0045926,GO:0045937,GO:0046530,GO:0046620,GO:0046621,GO:0046664,GO:0048468,GO:0048477,GO:0048513,GO:0048518,GO:0048519,GO:0048522,GO:0048523,GO:0048546,GO:0048565,GO:0048583,GO:0048585,GO:0048592,GO:0048593,GO:0048598,GO:0048609,GO:0048619,GO:0048638,GO:0048646,GO:0048666,GO:0048699,GO:0048729,GO:0048731,GO:0048732,GO:0048749,GO:0048793,GO:0048856,GO:0048858,GO:0048869,GO:0048871,GO:0050789,GO:0050793,GO:0050794,GO:0051174,GO:0051179,GO:0051234,GO:0051239,GO:0051241,GO:0055123,GO:0060041,GO:0060042,GO:0060059,GO:0060249,GO:0060271,GO:0060429,GO:0060541,GO:0060562,GO:0061024,GO:0061162,GO:0061245,GO:0061326,GO:0061333,GO:0061336,GO:0061339,GO:0061525,GO:0065007,GO:0065008,GO:0070160,GO:0070161,GO:0070925,GO:0071840,GO:0071944,GO:0072001,GO:0072002,GO:0072006,GO:0090162,GO:0097205,GO:2000026 | crumbs homolog 2 (Drosophila) |
| comp219021_c0_seq21.p1 | | Calcium-binding EGF domain |
| comp219021_c0_seq21.p2 | | Polycystic kidney disease |
| comp219021_c0_seq21.p3 | GO:0000003,GO:0003674,GO:0005215,GO:0005216,GO:0005261,GO:0005262,GO:0005488,GO:0005515,GO:0005575,GO:0005622,GO:0005623,GO:0005929,GO:0006810,GO:0006811,GO:0006812,GO:0006816,GO:0007610,GO:0007617,GO:0007618,GO:0008150,GO:0008324,GO:0009581,GO:0009582,GO:0009605,GO:0009612,GO:0009628,GO:0009987,GO:0015075,GO:0015085,GO:0015267,GO:0016020,GO:0019098,GO:0019904,GO:0022414,GO:0022803,GO:0022838,GO:0022857,GO:0022890,GO:0022891,GO:0022892,GO:0030001,GO:0031513,GO:0032501,GO:0032504,GO:0033057,GO:0034220,GO:0034606,GO:0034608,GO:0040008,GO:0040009,GO:0040010,GO:0042995,GO:0043025,GO:0043226,GO:0043227,GO:0043229,GO:0043231,GO:0044297,GO:0044424,GO:0044464,GO:0044699,GO:0044703,GO:0044706,GO:0044763,GO:0044765,GO:0045927,GO:0046873,GO:0048518,GO:0048609,GO:0050789,GO:0050896,GO:0050982,GO:0051179,GO:0051234,GO:0051606,GO:0051704,GO:0051705,GO:0055085,GO:0060179,GO:0065007,GO:0070588,GO:0070838,GO:0072372,GO:0072509,GO:0072511,GO:0097458 | polycystic kidney disease 1-like 2 |

| | | |
|---|---|---|
| comp214083_c0_seq3.p1 | GO:0003674,GO:0005575,GO:0005622,GO:0005623,GO:0005737,GO:0005783,GO:0005789,GO:0006805,GO:0007275,GO:0007423,GO:0008150,GO:0008152,GO:0009055,GO:0009410,GO:0009987,GO:0012505,GO:0016020,GO:0031090,GO:0032501,GO:0032502,GO:0042175,GO:0042221,GO:0043226,GO:0043227,GO:0043229,GO:0043231,GO:0044237,GO:0044281,GO:0044422,GO:0044424,GO:0044425,GO:0044432,GO:0044444,GO:0044446,GO:0044464,GO:0044699,GO:0044707,GO:0044710,GO:0044763,GO:0044767,GO:0048513,GO:0048731,GO:0048856,GO:0050896,GO:0051716,GO:0070887,GO:0071466 | cytochrome P450 |
| comp207297_c0_seq1.p1 | GO:0005575,GO:0005622,GO:0005623,GO:0005634,GO:0043226,GO:0043227,GO:0043229,GO:0043231,GO:0044424,GO:0044464 | Zinc finger protein 596 |
| comp213020_c0_seq3.p1 | GO:0000003,GO:0000146,GO:0000166,GO:0000302,GO:0001667,GO:0001701,GO:0001725,GO:0001756,GO:0001757,GO:0001778,GO:0001882,GO:0001883,GO:0002026,GO:0002027,GO:0002119,GO:0002164,GO:0003002,GO:0003006,GO:0003007,GO:0003008,GO:0003009,GO:0003010,GO:0003012,GO:0003013,GO:0003015,GO:0003205,GO:0003206,GO:0003208,GO:0003209,GO:0003228,GO:0003229,GO:0003230,GO:0003231,GO:0003674,GO:0003774,GO:0003779,GO:0003824,GO:0005198,GO:0005488,GO:0005515,GO:0005516,GO:0005524,GO:0005575,GO:0005622,GO:0005623,GO:0005634,GO:0005694,GO:0005700,GO:0005703,GO:0005730,GO:0005737,GO:0005794,GO:0005826,GO:0005829,GO:0005856,GO:0005859,GO:0005863,GO:0005911,GO:0005912,GO:0005924,GO:0005925,GO:0005938,GO:0006139,GO:0006140,GO:0006152,GO:0006163,GO:0006195,GO:0006200,GO:0006355,GO:0006357,GO:0006461,GO:0006725,GO:0006753,GO:0006793,GO:0006796,GO:0006807,GO:0006810,GO:0006897,GO:0006898,GO:0006928,GO:0006936,GO:0006941,GO:0006950,GO:0006979,GO:0006996,GO:0007009,GO:0007010,GO:0007015,GO:0007154,GO:0007165,GO:0007166,GO:0007167,GO:0007178,GO:0007275,GO:0007276,GO:0007292,GO:0007297,GO:0007298,GO:0007379,GO:0007389,GO:0007399,GO:0007413,GO:0007424,GO:0007427,GO:0007507,GO:0007512,GO:0007517,GO:0007519,GO:0007522,GO:0007525,GO:0007527,GO:0007610,GO:0007626,GO:0007631,GO:0008015,GO:0008016,GO:0008037,GO:0008038,GO:0008092,GO:0008150,GO:0008152,GO:0008217,GO:0008307,GO:0009056,GO:0009116,GO:0009117,GO:0009118,GO:0009119,GO:0009123,GO:0009125,GO:0009126,GO:0009128,GO:0009141,GO:0009143,GO:0009144,GO:0009146,GO:0009150,GO:0009154,GO:0009158,GO:0009161,GO:0009164,GO:0009166,GO:0009167,GO:0009169,GO:0009199,GO:0009203,GO:0009205,GO:0009207,GO:0009259,GO:0009261,GO:0009611,GO:0009653,GO:0009790,GO:0009791,GO:0009792,GO:0009880,GO:0009887,GO:0009888,GO:0009889,GO:0009894,GO:0009952,GO:0009987,GO:0010171,GO:0010468,GO:0010556,GO:0010608,GO:0010631,GO:0010927,GO:0014704,GO:0014706,GO:0014721,GO:0014823,GO:0014866,GO:0015629,GO:0016020,GO:0016043,GO:0016044,GO:0016055,GO:0016192,GO:0016459,GO:0016460,GO:0016462,GO:0016477,GO:0016787,GO:0016817,GO:0016818,GO:0016887,GO:0017076,GO:0017111,GO:0018991,GO:0019098,GO:0019219,GO:0019220,GO:0019222,GO:0019439,GO:0019637,GO:0019693,GO:0019953,GO:0022008,GO:0022412,GO:0022414,GO:0022603,GO:0022607,GO:0023052,GO:0030016,GO:0030017,GO:0030018,GO:0030029,GO:0030030,GO:0030036,GO:0030048,GO:0030049,GO:0030054,GO:0030055,GO:0030154,GO:0030182,GO:0030239,GO:0030240,GO:0030241,GO:0030509,GO:0030554,GO:0030707,GO:0030811,GO:0030898,GO:0030899,GO:0031032,GO:0031033,GO:0031034,GO:0031175,GO:0031323,GO:0031326,GO:0031329,GO:0031444,GO:0031647,GO:0031672,GO:0031674,GO:0031974,GO:0031981,GO:0032153,GO:0032155,GO:0032432,GO:0032501,GO:0032502,GO:0032504,GO:0032549,GO:0032550,GO:0032553,GO:0032555,GO:0032559,GO:0032956,GO:0032970,GO:0032982,GO:0032989,GO:0032991,GO:0033043,GO:0033057,GO:0033121,GO:0033275,GO:0034622,GO:0034641,GO:0034655,GO:0035050,GO:0035051,GO:0035282,GO:0035295,GO:0035556,GO:0035639,GO:0036094,GO:0036379,GO:0040007,GO:0040008,GO:0040011,GO:0040012,GO:0040017,GO:0042060,GO:0042221,GO:0042278,GO:0042454,GO:0042623,GO:0042641,GO:0042692,GO:0042755,GO:0042802,GO:0042803,GO:0043009,GO:0043050,GO:0043167,GO:0043168,GO:0043226,GO:0043227,GO:0043228,GO:0043229,GO:0043231,GO:0043232,GO:0043233,GO:0043234,GO:0043292,GO:0043462,GO:0043623,GO:0043933,GO:0044057,GO:0044085,GO:0044087,GO:0044237,GO:0044238,GO:0044248,GO:0044270,GO:0044281,GO:0044291,GO:0044422,GO:0044424,GO:0044427,GO:0044428,GO:0044430,GO:0044444,GO:0044446,GO:0044448,GO:0044449,GO:0044464,GO:0044699,GO:0044700,GO:0044702,GO:0044706,GO:0044707,GO:0044708,GO:0044710,GO:0044763,GO:0044767,GO:0045214,GO:0045595,GO:0046034,GO:0046128,GO:0046130,GO:0046434,GO:0046483,GO:0046620,GO:0046700,GO:0046982,GO:0046983,GO:0048017,GO:0048468,GO:0048477,GO:0048513,GO:0048518,GO:0048568,GO:0048609,GO:0048610,GO:0048638,GO:0048644,GO:0048646,GO:0048666,GO:0048699,GO:0048729,GO:0048731,GO:0048738,GO:0048739,GO:0048741,GO:0048747,GO:0048856,GO:0048869,GO:0048870,GO:0050789,GO:0050790,GO:0050793,GO:0050794,GO:0050821,GO:0050879,GO:0050881,GO:0050896,GO:0051015,GO:0051128,GO:0051146,GO:0051147,GO | Myosin heavy chain |

| | | |
|---|---|---|
| comp220891_c3_seq4.p1 | | Inherit from opiNOG: Endonuclease/Exonuclease/phosphatase family |
| comp220891_c3_seq4.p2 | | Inherit from KOG: Retrotransposon protein |
| comp220908_c0_seq3.p1 | | small nuclear RNA activating complex, polypeptide 4, 190kDa |
| comp218472_c0_seq1.p1 | | Inherit from biNOG: ankyrin 3, node of Ranvier (ankyrin G) |
| comp220294_c0_seq4.p1 | GO:0000278,GO:0000279,GO:0003674,GO:0003824,GO:0004221,GO:0004843,GO:0005575,GO:0005622,GO:0005623,GO:0005634,GO:0005737,GO:0006464,GO:0007049,GO:0007088,GO:0007346,GO:0008150,GO:0008152,GO:0008233,GO:0008234,GO:0009892,GO:0009894,GO:0009895,GO:0009987,GO:0010564,GO:0010605,GO:0010639,GO:0010948,GO:0016579,GO:0016787,GO:0016788,GO:0016790,GO:0019222,GO:0019538,GO:0019783,GO:0022402,GO:0022403,GO:0030071,GO:0030162,GO:0031323,GO:0031324,GO:0031329,GO:0031330,GO:0031396,GO:0031397,GO:0031399,GO:0031400,GO:0032268,GO:0032269,GO:0032434,GO:0032435,GO:0033043,GO:0036211,GO:0042176,GO:0042177,GO:0043086,GO:0043170,GO:0043226,GO:0043227,GO:0043229,GO:0043231,GO:0043412,GO:0044092,GO:0044237,GO:0044238,GO:0044260,GO:0044267,GO:0044424,GO:0044464,GO:0044699,GO:0044763,GO:0045786,GO:0045839,GO:0045841,GO:0045861,GO:0048519,GO:0048523,GO:0050789,GO:0050790,GO:0050794,GO:0051128,GO:0051129,GO:0051246,GO:0051248,GO:0051322,GO:0051340,GO:0051352,GO:0051436,GO:0051438,GO:0051439,GO:0051444,GO:0051726,GO:0051783,GO:0051784,GO:0060255,GO:0060564,GO:0061136,GO:0065007,GO:0065009,GO:0070011,GO:0070646,GO:0070647,GO:0071704,GO:0080090,GO:0090231,GO:1901799,GO:1901976,GO:1901987,GO:1901988,GO:1901990,GO:1901991,GO:1902099,GO:1902100 | ubiquitin- specific peptidase |

| | | |
|---|---|---|
| comp215818_c1_seq4.p1 | GO:0000003,GO:0000226,GO:0000578,GO:0000902,GO:0000904,GO:0001505,GO:0001654,GO:0001754,GO:0003001,GO:0003002,GO:0003006,GO:0003008,GO:0003674,GO:0003774,GO:0003777,GO:0003824,GO:0005488,GO:0005515,GO:0005575,GO:0005622,GO:0005623,GO:0005737,GO:0005856,GO:0005871,GO:0005874,GO:0005875,GO:0006403,GO:0006810,GO:0006836,GO:0006886,GO:0006928,GO:0006935,GO:0006996,GO:0007005,GO:0007010,GO:0007017,GO:0007018,GO:0007028,GO:0007154,GO:0007163,GO:0007267,GO:0007268,GO:0007269,GO:0007275,GO:0007276,GO:0007281,GO:0007292,GO:0007300,GO:0007303,GO:0007308,GO:0007309,GO:0007310,GO:0007314,GO:0007315,GO:0007316,GO:0007317,GO:0007350,GO:0007351,GO:0007389,GO:0007399,GO:0007409,GO:0007411,GO:0007423,GO:0007517,GO:0007519,GO:0008017,GO:0008088,GO:0008092,GO:0008103,GO:0008104,GO:0008150,GO:0008152,GO:0008298,GO:0008358,GO:0008574,GO:0008595,GO:0009605,GO:0009653,GO:0009790,GO:0009798,GO:0009880,GO:0009887,GO:0009888,GO:0009948,GO:0009950,GO:0009952,GO:0009953,GO:0009987,GO:0009994,GO:0010970,GO:0014706,GO:0015031,GO:0015630,GO:0015631,GO:0016043,GO:0016325,GO:0016358,GO:0016462,GO:0016482,GO:0016787,GO:0016817,GO:0016818,GO:0017111,GO:0019094,GO:0019226,GO:0019227,GO:0019953,GO:0021700,GO:0022008,GO:0022412,GO:0022414,GO:0022607,GO:0023052,GO:0023061,GO:0030011,GO:0030030,GO:0030154,GO:0030182,GO:0030705,GO:0030951,GO:0030952,GO:0031175,GO:0032501,GO:0032502,GO:0032504,GO:0032879,GO:0032940,GO:0032989,GO:0032990,GO:0032991,GO:0033036,GO:0034613,GO:0034643,GO:0035282,GO:0035371,GO:0035637,GO:0040011,GO:0040023,GO:0042221,GO:0042330,GO:0042692,GO:0043226,GO:0043228,GO:0043229,GO:0043232,GO:0043234,GO:0044085,GO:0044087,GO:0044422,GO:0044424,GO:0044430,GO:0044446,GO:0044464,GO:0044699,GO:0044700,GO:0044702,GO:0044707,GO:0044763,GO:0044765,GO:0044767,GO:0045184,GO:0045451,GO:0045595,GO:0045995,GO:0046530,GO:0046903,GO:0046907,GO:0047497,GO:0048311,GO:0048312,GO:0048468,GO:0048469,GO:0048477,GO:0048513,GO:0048592,GO:0048599,GO:0048609,GO:0048610,GO:0048666,GO:0048667,GO:0048699,GO:0048731,GO:0048741,GO:0048747,GO:0048812,GO:0048813,GO:0048856,GO:0048858,GO:0048869,GO:0050789,GO:0050793,GO:0050794,GO:0050877,GO:0050896,GO:0051128,GO:0051146,GO:0051179,GO:0051234,GO:0051239,GO:0051640,GO:0051641,GO:0051646,GO:0051647,GO:0051649,GO:0051654,GO:0051656,GO:0055001,GO:0055002,GO:0060281,GO:0060284,GO:0060341,GO:0060537,GO:0060538,GO:0060810,GO:0060811,GO:0061061,GO:0061564,GO:0065007,GO:0065008,GO:0070727,GO:0071702,GO:0071840,GO:0072384,GO:0097485,GO:2000026,GO:2000241 | KISc |

| | | |
|---|---|---|
| comp215818_c1_seq4.p2 | GO:0000003,GO:0000226,GO:0000578,GO:0000902,GO:0000904,GO:0001505,GO:0001654,GO:0001754,GO:0003001,GO:0003002,GO:0003006,GO:0003008,GO:0003674,GO:0003774,GO:0003777,GO:0003824,GO:0005488,GO:0005515,GO:0005575,GO:0005622,GO:0005623,GO:0005737,GO:0005856,GO:0005871,GO:0005874,GO:0005875,GO:0006403,GO:0006810,GO:0006836,GO:0006886,GO:0006928,GO:0006935,GO:0006996,GO:0007005,GO:0007010,GO:0007017,GO:0007018,GO:0007028,GO:0007154,GO:0007163,GO:0007267,GO:0007268,GO:0007269,GO:0007275,GO:0007276,GO:0007281,GO:0007292,GO:0007300,GO:0007303,GO:0007308,GO:0007309,GO:0007310,GO:0007314,GO:0007315,GO:0007316,GO:0007317,GO:0007350,GO:0007351,GO:0007389,GO:0007399,GO:0007409,GO:0007411,GO:0007423,GO:0007517,GO:0007519,GO:0008017,GO:0008088,GO:0008092,GO:0008103,GO:0008104,GO:0008150,GO:0008152,GO:0008298,GO:0008358,GO:0008574,GO:0008595,GO:0009605,GO:0009653,GO:0009790,GO:0009798,GO:0009880,GO:0009887,GO:0009888,GO:0009948,GO:0009950,GO:0009952,GO:0009953,GO:0009987,GO:0009994,GO:0010970,GO:0014706,GO:0015031,GO:0015630,GO:0015631,GO:0016043,GO:0016325,GO:0016358,GO:0016462,GO:0016482,GO:0016787,GO:0016817,GO:0016818,GO:0017111,GO:0019094,GO:0019226,GO:0019227,GO:0019953,GO:0021700,GO:0022008,GO:0022412,GO:0022414,GO:0022607,GO:0023052,GO:0023061,GO:0030011,GO:0030030,GO:0030154,GO:0030182,GO:0030705,GO:0030951,GO:0030952,GO:0031175,GO:0032501,GO:0032502,GO:0032504,GO:0032879,GO:0032940,GO:0032989,GO:0032990,GO:0032991,GO:0033036,GO:0034613,GO:0034643,GO:0035282,GO:0035371,GO:0035637,GO:0040011,GO:0040023,GO:0042221,GO:0042330,GO:0042692,GO:0043226,GO:0043228,GO:0043229,GO:0043232,GO:0043234,GO:0044085,GO:0044087,GO:0044422,GO:0044424,GO:0044430,GO:0044446,GO:0044464,GO:0044699,GO:0044700,GO:0044702,GO:0044707,GO:0044763,GO:0044765,GO:0044767,GO:0045184,GO:0045451,GO:0045595,GO:0045995,GO:0046530,GO:0046903,GO:0046907,GO:0047497,GO:0048311,GO:0048312,GO:0048468,GO:0048469,GO:0048477,GO:0048513,GO:0048592,GO:0048599,GO:0048609,GO:0048610,GO:0048666,GO:0048667,GO:0048699,GO:0048731,GO:0048741,GO:0048747,GO:0048812,GO:0048813,GO:0048856,GO:0048858,GO:0048869,GO:0050789,GO:0050793,GO:0050794,GO:0050877,GO:0050896,GO:0051128,GO:0051146,GO:0051179,GO:0051234,GO:0051239,GO:0051640,GO:0051641,GO:0051646,GO:0051647,GO:0051649,GO:0051654,GO:0051656,GO:0055001,GO:0055002,GO:0060281,GO:0060284,GO:0060341,GO:0060537,GO:0060538,GO:0060810,GO:0060811,GO:0061061,GO:0061564,GO:0065007,GO:0065008,GO:0070727,GO:0071702,GO:0071840,GO:0072384,GO:0097485,GO:2000026,GO:2000241 | KISc |
| comp220613_c0_seq6.p1 | GO:0003674,GO:0003724,GO:0003824,GO:0004004,GO:0004386,GO:0005575,GO:0005622,GO:0005623,GO:0005634,GO:0005635,GO:0005694,GO:0005737,GO:0005829,GO:0006139,GO:0006152,GO:0006163,GO:0006195,GO:0006200,GO:0006725,GO:0006753,GO:0006793,GO:0006796,GO:0006807,GO:0008026,GO:0008150,GO:0008152,GO:0008186,GO:0009056,GO:0009116,GO:0009117,GO:0009119,GO:0009123,GO:0009125,GO:0009126,GO:0009128,GO:0009141,GO:0009143,GO:0009144,GO:0009146,GO:0009150,GO:0009154,GO:0009158,GO:0009161,GO:0009164,GO:0009166,GO:0009167,GO:0009169,GO:0009199,GO:0009203,GO:0009205,GO:0009207,GO:0009259,GO:0009261,GO:0009987,GO:0010525,GO:0010526,GO:0010528,GO:0010529,GO:0012505,GO:0016020,GO:0016462,GO:0016787,GO:0016817,GO:0016818,GO:0016887,GO:0017111,GO:0019439,GO:0019637,GO:0019693,GO:0031090,GO:0031965,GO:0031967,GO:0031975,GO:0034641,GO:0034655,GO:0042278,GO:0042454,GO:0042623,GO:0043226,GO:0043227,GO:0043228,GO:0043229,GO:0043231,GO:0043232,GO:0044237,GO:0044238,GO:0044248,GO:0044270,GO:0044281,GO:0044422,GO:0044424,GO:0044428,GO:0044444,GO:0044446,GO:0044464,GO:0044710,GO:0046034,GO:0046128,GO:0046130,GO:0046434,GO:0046483,GO:0046700,GO:0048519,GO:0048523,GO:0050789,GO:0050794,GO:0055086,GO:0065007,GO:0070035,GO:0070725,GO:0071704,GO:0072521,GO:0072523,GO:1901135,GO:1901136,GO:1901292,GO:1901360,GO:1901361,GO:1901564,GO:1901565,GO:1901575,GO:1901657,GO:1901658 | tudor domain containing 12 |

| | | |
|---|---|---|
| comp221215_c0_seq1.p1 | GO:0000226,GO:0000235,GO:0000902,GO:0000904,GO:0001558,GO:0001578,GO:0001709,GO:0001763,GO:0002009,GO:0002165,GO:0003674,GO:0003779,GO:0005488,GO:0005515,GO:0005575,GO:0005622,GO:0005623,GO:0005737,GO:0005818,GO:0005819,GO:0005856,GO:0005874,GO:0005876,GO:0005881,GO:0005912,GO:0006928,GO:0006935,GO:0006996,GO:0007010,GO:0007017,GO:0007026,GO:0007154,GO:0007267,GO:0007275,GO:0007399,GO:0007409,GO:0007411,GO:0007417,GO:0007420,GO:0007423,GO:0007424,GO:0007444,GO:0007472,GO:0007475,GO:0007476,GO:0007517,GO:0007552,GO:0007560,GO:0008017,GO:0008037,GO:0008038,GO:0008092,GO:0008150,GO:0008361,GO:0008587,GO:0009605,GO:0009653,GO:0009791,GO:0009886,GO:0009887,GO:0009888,GO:0009987,GO:0010639,GO:0010720,GO:0010769,GO:0010975,GO:0015630,GO:0015631,GO:0016043,GO:0016198,GO:0016199,GO:0016203,GO:0016204,GO:0016319,GO:0016358,GO:0022008,GO:0022603,GO:0022604,GO:0023052,GO:0030029,GO:0030030,GO:0030036,GO:0030054,GO:0030154,GO:0030175,GO:0030182,GO:0030307,GO:0030424,GO:0030426,GO:0030427,GO:0030516,GO:0030716,GO:0031110,GO:0031111,GO:0031114,GO:0031175,GO:0031344,GO:0031346,GO:0032501,GO:0032502,GO:0032535,GO:0032886,GO:0032989,GO:0032990,GO:0032991,GO:0033043,GO:0033267,GO:0035107,GO:0035114,GO:0035120,GO:0035146,GO:0035147,GO:0035148,GO:0035149,GO:0035152,GO:0035220,GO:0035239,GO:0035295,GO:0040008,GO:0040011,GO:0042221,GO:0042330,GO:0042995,GO:0043005,GO:0043226,GO:0043228,GO:0043229,GO:0043232,GO:0043234,GO:0043242,GO:0043244,GO:0044087,GO:0044295,GO:0044422,GO:0044424,GO:0044430,GO:0044444,GO:0044446,GO:0044463,GO:0044464,GO:0044699,GO:0044700,GO:0044707,GO:0044763,GO:0044767,GO:0045165,GO:0045169,GO:0045595,GO:0045597,GO:0045664,GO:0045773,GO:0045927,GO:0048468,GO:0048513,GO:0048518,GO:0048519,GO:0048522,GO:0048523,GO:0048563,GO:0048569,GO:0048638,GO:0048639,GO:0048646,GO:0048666,GO:0048667,GO:0048699,GO:0048707,GO:0048729,GO:0048731,GO:0048736,GO:0048737,GO:0048754,GO:0048812,GO:0048813,GO:0048856,GO:0048858,GO:0048869,GO:0050767,GO:0050769,GO:0050770,GO:0050772,GO:0050789,GO:0050793,GO:0050794,GO:0050896,GO:0051094,GO:0051128,GO:0051129,GO:0051130,GO:0051239,GO:0051489,GO:0051491,GO:0051493,GO:0051494,GO:0051960,GO:0060284,GO:0060429,GO:0060446,GO:0060491,GO:0060538,GO:0060541,GO:0060562,GO:0061061,GO:0061138,GO:0061387,GO:0061564,GO:0065007,GO:0065008,GO:0070161,GO:0070507,GO:0071840,GO:0090066,GO:0097458,GO:0097485,GO:1901879,GO:1901880,GO:2000026 | Spectrin repeat |

| | | |
|---|---|---|
| comp208375_c1_seq1.p1 | GO:0001775,GO:0003002,GO:0003674,GO:0005102,GO:0005114,GO:0005126,GO:0005160,GO:0005488,GO:0005515,GO:0005575,GO:0005622,GO:0005623,GO:0005737,GO:0005886,GO:0005887,GO:0006355,GO:0006928,GO:0006950,GO:0007154,GO:0007165,GO:0007166,GO:0007167,GO:0007178,GO:0007179,GO:0007275,GO:0007389,GO:0007596,GO:0007599,GO:0008150,GO:0008284,GO:0008360,GO:0009611,GO:0009719,GO:0009889,GO:0009891,GO:0009893,GO:0009953,GO:0009966,GO:0009967,GO:0009968,GO:0009987,GO:0010033,GO:0010468,GO:0010556,GO:0010557,GO:0010604,GO:0010628,GO:0010646,GO:0010647,GO:0010648,GO:0010717,GO:0010718,GO:0010720,GO:0010769,GO:0010770,GO:0016020,GO:0016021,GO:0016477,GO:0017015,GO:0019219,GO:0019222,GO:0022603,GO:0022604,GO:0023051,GO:0023052,GO:0023056,GO:0023057,GO:0030111,GO:0030168,GO:0030177,GO:0030510,GO:0030512,GO:0030514,GO:0031224,GO:0031226,GO:0031323,GO:0031325,GO:0031326,GO:0031328,GO:0032092,GO:0032386,GO:0032388,GO:0032501,GO:0032502,GO:0032879,GO:0032880,GO:0033157,GO:0035412,GO:0035413,GO:0040011,GO:0042060,GO:0042127,GO:0042221,GO:0042306,GO:0042307,GO:0043393,GO:0044093,GO:0044424,GO:0044425,GO:0044459,GO:0044464,GO:0044699,GO:0044700,GO:0044707,GO:0044763,GO:0045595,GO:0045597,GO:0045893,GO:0045935,GO:0046822,GO:0046824,GO:0048518,GO:0048519,GO:0048522,GO:0048523,GO:0048583,GO:0048584,GO:0048585,GO:0048870,GO:0050789,GO:0050793,GO:0050794,GO:0050817,GO:0050878,GO:0050896,GO:0051049,GO:0051050,GO:0051094,GO:0051098,GO:0051099,GO:0051128,GO:0051130,GO:0051171,GO:0051173,GO:0051179,GO:0051222,GO:0051223,GO:0051239,GO:0051252,GO:0051254,GO:0051674,GO:0051716,GO:0060255,GO:0060284,GO:0060341,GO:0060828,GO:0065007,GO:0065008,GO:0065009,GO:0070201,GO:0070848,GO:0070887,GO:0071310,GO:0071363,GO:0071495,GO:0071559,GO:0071560,GO:0071944,GO:0080090,GO:0090092,GO:0090101,GO:0090263,GO:0090287,GO:0090288,GO:0090316,GO:1900180,GO:2000026,GO:2000112,GO:2000736,GO:2001141 | BMP and activin membrane-bound inhibitor homolog (Xenopus laevis) |
| comp215876_c1_seq4.p1 | GO:0003674,GO:0003824,GO:0004465,GO:0005575,GO:0005622,GO:0005623,GO:0005737,GO:0007154,GO:0007165,GO:0007602,GO:0008150,GO:0008152,GO:0009314,GO:0009416,GO:0009581,GO:0009582,GO:0009583,GO:0009605,GO:0009628,GO:0009987,GO:0016298,GO:0016787,GO:0016788,GO:0023052,GO:0044424,GO:0044464,GO:0044699,GO:0044700,GO:0044763,GO:0050789,GO:0050794,GO:0050896,GO:0051606,GO:0051716,GO:0052689,GO:0065007 | Lipase (class 3) |
| comp210910_c1_seq1.p1 | | PGAM |

| | | |
|---|---|---|
| comp184398_c1_seq2.p1 | GO:0000003,GO:0000981,GO:0000982,GO:0001071,GO:0001077,GO:0001085,GO:0001228,GO:0001525,GO:0001558,GO:0001568,GO:0001666,GO:0001701,GO:0001890,GO:0001892,GO:0001894,GO:0001944,GO:0001974,GO:0002027,GO:0002262,GO:0002376,GO:0002520,GO:0003008,GO:0003674,GO:0003676,GO:0003677,GO:0003700,GO:0005488,GO:0005515,GO:0005575,GO:0005622,GO:0005623,GO:0005634,GO:0005654,GO:0005667,GO:0005737,GO:0005829,GO:0006139,GO:0006351,GO:0006355,GO:0006357,GO:0006366,GO:0006584,GO:0006725,GO:0006807,GO:0006915,GO:0006950,GO:0006974,GO:0006979,GO:0006996,GO:0007005,GO:0007154,GO:0007165,GO:0007275,GO:0007276,GO:0007292,GO:0007399,GO:0007568,GO:0007600,GO:0007601,GO:0008016,GO:0008134,GO:0008150,GO:0008152,GO:0008219,GO:0008340,GO:0009058,GO:0009059,GO:0009266,GO:0009267,GO:0009408,GO:0009605,GO:0009628,GO:0009653,GO:0009712,GO:0009719,GO:0009725,GO:0009790,GO:0009792,GO:0009889,GO:0009891,GO:0009893,GO:0009987,GO:0009991,GO:0010033,GO:0010243,GO:0010259,GO:0010286,GO:0010467,GO:0010468,GO:0010556,GO:0010557,GO:0010604,GO:0010628,GO:0010941,GO:0012501,GO:0014823,GO:0016043,GO:0016070,GO:0016265,GO:0018130,GO:0018958,GO:0019219,GO:0019222,GO:0019438,GO:0019899,GO:0019953,GO:0021700,GO:0022008,GO:0022414,GO:0023052,GO:0030097,GO:0030099,GO:0030154,GO:0030182,GO:0030218,GO:0030308,GO:0030323,GO:0030324,GO:0031323,GO:0031325,GO:0031326,GO:0031328,GO:0031667,GO:0031668,GO:0031669,GO:0031974,GO:0031981,GO:0032501,GO:0032502,GO:0032504,GO:0032774,GO:0032868,GO:0032869,GO:0032870,GO:0032991,GO:0033554,GO:0034101,GO:0034599,GO:0034641,GO:0034645,GO:0034654,GO:0035035,GO:0035295,GO:0036293,GO:0036294,GO:0040008,GO:0042221,GO:0042415,GO:0042592,GO:0042594,GO:0042692,GO:0042981,GO:0043009,GO:0043066,GO:0043067,GO:0043069,GO:0043129,GO:0043170,GO:0043226,GO:0043227,GO:0043229,GO:0043231,GO:0043233,GO:0043234,GO:0043434,GO:0043523,GO:0043618,GO:0043619,GO:0043620,GO:0044057,GO:0044237,GO:0044238,GO:0044249,GO:0044260,GO:0044271,GO:0044422,GO:0044424,GO:0044428,GO:0044444,GO:0044446,GO:0044451,GO:0044464,GO:0044699,GO:0044700,GO:0044702,GO:0044707,GO:0044763,GO:0044767,GO:0045165,GO:0045445,GO:0045893,GO:0045926,GO:0045935,GO:0045944,GO:0046483,GO:0048468,GO:0048469,GO:0048477,GO:0048513,GO:0048514,GO:0048518,GO:0048519,GO:0048522,GO:0048523,GO:0048534,GO:0048568,GO:0048609,GO:0048625,GO:0048646,GO:0048699,GO:0048731,GO:0048771,GO:0048856,GO:0048869,GO:0048871,GO:0048872,GO:0048875,GO:0048878,GO:0050789,GO:0050794,GO:0050877,GO:0050896,GO:0050953,GO:0051128,GO:0051171,GO:0051173,GO:0051239,GO:0051252,GO:0051254,GO:0051402,GO:0051716,GO:0051726,GO:0060249,GO:0060255,GO:0060438,GO:0060541,GO:0060548,GO:0061061,GO:0061418,GO:0061433,GO:0065007,GO:0065008,GO:0070013,GO:0070482,GO:0070887,GO:0070997,GO:0071310,GO:0071375,GO:0071417,GO:0071453,GO:0071456,GO:0071495,GO:0071496,GO:0071704,GO:0071840,GO:0072358,GO:0072359,GO:0080090,GO:0090304,GO:0097159,GO:0097285,GO:1901214,GO:1901360,GO:1901362,GO:1901363,GO:1901564,GO:1901576,GO:1901615,GO:1901652,GO:1901653,GO:1901698,GO:1901699,GO:1901700,GO:1901701,GO:2000112,GO:2001141 | Hypoxia inducible factor 1, alpha subunit (Basic helix-loop-helix transcription factor) |
| comp218218_c0_seq12.p1 | GO:0008150,GO:0009653,GO:0010171,GO:0032502,GO:0044699,GO:0044767,GO:0048856 | pogo transposable element with KRAB domain |
| comp213493_c1_seq5.p1 | | Reverse transcriptase (RNA-dependent DNA polymerase) |

| | | |
|---|---|---|
| comp215442_c0_seq3.p1 | GO:0000122,GO:0000278,GO:0001654,GO:0001745,GO:0001932,GO:0001933,GO:0002165,GO:0003674,GO:0003824,GO:0004721,GO:0004722,GO:0004723,GO:0004871,GO:0005488,GO:0005515,GO:0005575,GO:0005622,GO:0005623,GO:0005634,GO:0005654,GO:0005737,GO:0005829,GO:0006139,GO:0006351,GO:0006352,GO:0006355,GO:0006357,GO:0006366,GO:0006367,GO:0006464,GO:0006469,GO:0006470,GO:0006725,GO:0006793,GO:0006796,GO:0006807,GO:0006950,GO:0006979,GO:0007049,GO:0007050,GO:0007154,GO:0007165,GO:0007166,GO:0007167,GO:0007169,GO:0007178,GO:0007179,GO:0007275,GO:0007399,GO:0007423,GO:0007444,GO:0007472,GO:0007476,GO:0007552,GO:0007560,GO:0008150,GO:0008152,GO:0008286,GO:0009058,GO:0009059,GO:0009653,GO:0009719,GO:0009725,GO:0009791,GO:0009886,GO:0009887,GO:0009889,GO:0009890,GO:0009891,GO:0009892,GO:0009893,GO:0009966,GO:0009967,GO:0009968,GO:0009987,GO:0010033,GO:0010243,GO:0010467,GO:0010468,GO:0010556,GO:0010557,GO:0010558,GO:0010563,GO:0010604,GO:0010605,GO:0010627,GO:0010628,GO:0010629,GO:0010646,GO:0010647,GO:0010648,GO:0010740,GO:0010741,GO:0010990,GO:0010991,GO:0016055,GO:0016070,GO:0016311,GO:0016787,GO:0016788,GO:0016791,GO:0017015,GO:0018130,GO:0019219,GO:0019220,GO:0019221,GO:0019222,GO:0019438,GO:0019538,GO:0022008,GO:0022402,GO:0022603,GO:0023051,GO:0023052,GO:0023056,GO:0023057,GO:0030111,GO:0030154,GO:0030177,GO:0030512,GO:0031323,GO:0031324,GO:0031325,GO:0031326,GO:0031327,GO:0031328,GO:0031333,GO:0031399,GO:0031400,GO:0031974,GO:0031981,GO:0032268,GO:0032269,GO:0032501,GO:0032502,GO:0032774,GO:0032868,GO:0032869,GO:0032870,GO:0033192,GO:0033673,GO:0034097,GO:0034641,GO:0034645,GO:0034654,GO:0035107,GO:0035114,GO:0035120,GO:0035220,GO:0035970,GO:0036211,GO:0042221,GO:0042325,GO:0042326,GO:0042578,GO:0042995,GO:0043005,GO:0043086,GO:0043122,GO:0043123,GO:0043170,GO:0043226,GO:0043227,GO:0043229,GO:0043231,GO:0043233,GO:0043254,GO:0043405,GO:0043407,GO:0043408,GO:0043409,GO:0043412,GO:0043434,GO:0043549,GO:0044087,GO:0044092,GO:0044237,GO:0044238,GO:0044249,GO:0044260,GO:0044267,GO:0044271,GO:0044422,GO:0044424,GO:0044428,GO:0044444,GO:0044446,GO:0044464,GO:0044699,GO:0044700,GO:0044707,GO:0044763,GO:0044767,GO:0045595,GO:0045597,GO:0045664,GO:0045666,GO:0045676,GO:0045678,GO:0045786,GO:0045859,GO:0045892,GO:0045893,GO:0045934,GO:0045935,GO:0045936,GO:0046332,GO:0046483,GO:0046532,GO:0046534,GO:0046578,GO:0046580,GO:0048513,GO:0048518,GO:0048519,GO:0048522,GO:0048523,GO:0048563,GO:0048569,GO:0048583,GO:0048584,GO:0048585,GO:0048592,GO:0048699,GO:0048707,GO:0048731,GO:0048736,GO:0048737,GO:0048749,GO:0048856,GO:0048869,GO:0050767,GO:0050789,GO:0050790,GO:0050793,GO:0050794,GO:0050896,GO:0051056,GO:0051058,GO:0051094,GO:0051128,GO:0051129,GO:0051171,GO:0051172,GO:0051173,GO:0051174,GO:0051239,GO:0051240,GO:0051246,GO:0051248,GO:0051252,GO:0051253,GO:0051254,GO:0051338,GO:0051348,GO:0051716,GO:0051726,GO:0051960,GO:0060089,GO:0060255,GO:0060284,GO:0065007,GO:0065009,GO:0070013,GO:0070412,GO:0070848,GO:0070887,GO:0071310,GO:0071345,GO:0071363,GO:0071375,GO:0071417,GO:0071495,GO:0071559,GO:0071560,GO:0071704,GO:0071900,GO:0071901,GO:0080090,GO:0090092,GO:0090101,GO:0090287,GO:0090288,GO:0090304,GO:0097458,GO:1901360,GO:1901362,GO:1901576,GO:1901652,GO:1901653,GO:1901698,GO:1901699,GO:1901700,GO:1901701,GO:2000026,GO:2000027,GO:2000112,GO:2000113,GO:2001141 | protein phosphatase, Mg2 Mn2 dependent |
| comp218414_c0_seq16.p1 | | matrix metallopeptidase |
| comp220442_c0_seq4.p1 | | Inherit from KOG: ATPdependent RNA helicase |
| comp214548_c1_seq3.p1 | | Fatty acid amide hydrolase |

| | | |
|---|---|---|
| comp213703_c2_seq9.p1 | GO:0000122,GO:0000578,GO:0000902,GO:0000904,GO:0000981,GO:0000982,GO:0001071,GO:0001078,GO:0001227,GO:0001667,GO:0001709,GO:0003002,GO:0003674,GO:0003700,GO:0004871,GO:0004872,GO:0004879,GO:0005488,GO:0005515,GO:0005575,GO:0005622,GO:0005623,GO:0005634,GO:0006139,GO:0006351,GO:0006355,GO:0006357,GO:0006366,GO:0006725,GO:0006807,GO:0006928,GO:0007088,GO:0007154,GO:0007165,GO:0007275,GO:0007346,GO:0007350,GO:0007351,GO:0007354,GO:0007389,GO:0007399,GO:0007417,GO:0007424,GO:0007427,GO:0007517,GO:0008134,GO:0008150,GO:0008152,GO:0008544,GO:0008595,GO:0009058,GO:0009059,GO:0009653,GO:0009790,GO:0009798,GO:0009880,GO:0009888,GO:0009889,GO:0009890,GO:0009892,GO:0009948,GO:0009952,GO:0009987,GO:0010467,GO:0010468,GO:0010556,GO:0010558,GO:0010564,GO:0010605,GO:0010629,GO:0010631,GO:0016043,GO:0016070,GO:0016358,GO:0016477,GO:0018130,GO:0019219,GO:0019222,GO:0019438,GO:0022008,GO:0023052,GO:0030030,GO:0030154,GO:0030182,GO:0030522,GO:0031175,GO:0031323,GO:0031324,GO:0031326,GO:0031327,GO:0032501,GO:0032502,GO:0032774,GO:0032989,GO:0032990,GO:0033043,GO:0034641,GO:0034645,GO:0034654,GO:0035150,GO:0035151,GO:0035152,GO:0035282,GO:0035290,GO:0038023,GO:0040011,GO:0043170,GO:0043226,GO:0043227,GO:0043229,GO:0043231,GO:0043588,GO:0044237,GO:0044238,GO:0044249,GO:0044260,GO:0044271,GO:0044424,GO:0044464,GO:0044699,GO:0044700,GO:0044707,GO:0044763,GO:0044767,GO:0045165,GO:0045892,GO:0045934,GO:0046483,GO:0046845,GO:0048468,GO:0048513,GO:0048519,GO:0048523,GO:0048666,GO:0048667,GO:0048699,GO:0048731,GO:0048812,GO:0048813,GO:0048856,GO:0048858,GO:0048869,GO:0048870,GO:0050789,GO:0050794,GO:0050896,GO:0051128,GO:0051171,GO:0051172,GO:0051179,GO:0051252,GO:0051253,GO:0051674,GO:0051716,GO:0051726,GO:0051783,GO:0060089,GO:0060255,GO:0060541,GO:0061061,GO:0065007,GO:0065008,GO:0070491,GO:0071704,GO:0071840,GO:0080090,GO:0090066,GO:0090130,GO:0090132,GO:0090304,GO:1901360,GO:1901362,GO:1901576,GO:2000112,GO:2000113,GO:2001141 | branched duct epithelial cell fate determination, open tracheal system |
| comp207250_c0_seq1.p1 | GO:0003674,GO:0003824,GO:0005215,GO:0005388,GO:0005575,GO:0005623,GO:0005886,GO:0006139,GO:0006152,GO:0006163,GO:0006195,GO:0006200,GO:0006725,GO:0006753,GO:0006793,GO:0006796,GO:0006807,GO:0006810,GO:0006811,GO:0006812,GO:0006816,GO:0006873,GO:0006874,GO:0006875,GO:0008150,GO:0008152,GO:0008324,GO:0009056,GO:0009116,GO:0009117,GO:0009119,GO:0009123,GO:0009125,GO:0009126,GO:0009128,GO:0009141,GO:0009143,GO:0009144,GO:0009146,GO:0009150,GO:0009154,GO:0009158,GO:0009161,GO:0009164,GO:0009166,GO:0009167,GO:0009169,GO:0009199,GO:0009203,GO:0009205,GO:0009207,GO:0009259,GO:0009261,GO:0009987,GO:0015075,GO:0015085,GO:0015399,GO:0015405,GO:0015662,GO:0016020,GO:0016462,GO:0016787,GO:0016817,GO:0016818,GO:0016820,GO:0016887,GO:0017111,GO:0019439,GO:0019637,GO:0019693,GO:0019725,GO:0019829,GO:0022804,GO:0022857,GO:0022890,GO:0022891,GO:0022892,GO:0030001,GO:0030003,GO:0034220,GO:0034641,GO:0034655,GO:0042278,GO:0042454,GO:0042592,GO:0042623,GO:0042625,GO:0042626,GO:0043492,GO:0044237,GO:0044238,GO:0044248,GO:0044270,GO:0044281,GO:0044464,GO:0044699,GO:0044710,GO:0044763,GO:0044765,GO:0046034,GO:0046128,GO:0046130,GO:0046434,GO:0046483,GO:0046700,GO:0046873,GO:0048878,GO:0050801,GO:0051179,GO:0051234,GO:0055065,GO:0055074,GO:0055080,GO:0055082,GO:0055085,GO:0055086,GO:0065007,GO:0065008,GO:0070588,GO:0070838,GO:0071704,GO:0071944,GO:0072503,GO:0072507,GO:0072509,GO:0072511,GO:0072521,GO:0072523,GO:1901135,GO:1901136,GO:1901292,GO:1901360,GO:1901361,GO:1901564,GO:1901565,GO:1901575,GO:1901657,GO:1901658 | Plasma membrane calcium-transporting ATPase |

| | | |
|---|---|---|
| comp214987_c4_seq4.p1 | GO:0000226,GO:0000902,GO:0001578,GO:0001894,GO:0003674,GO:0003824,GO:0004672,GO:0004674,GO:0005488,GO:0005515,GO:0005575,GO:0005622,GO:0005623,GO:0005737,GO:0005829,GO:0006464,GO:0006468,GO:0006793,GO:0006796,GO:0006915,GO:0006950,GO:0006952,GO:0006996,GO:0007010,GO:0007015,GO:0007017,GO:0007154,GO:0007165,GO:0007243,GO:0007249,GO:0007252,GO:0007275,GO:0007399,GO:0007423,GO:0007424,GO:0008150,GO:0008152,GO:0008219,GO:0008356,GO:0008384,GO:0008385,GO:0008407,GO:0009653,GO:0009887,GO:0009987,GO:0010669,GO:0010941,GO:0010950,GO:0010952,GO:0012501,GO:0016043,GO:0016265,GO:0016301,GO:0016310,GO:0016322,GO:0016740,GO:0016772,GO:0016773,GO:0019222,GO:0019538,GO:0021700,GO:0022008,GO:0022416,GO:0023052,GO:0030029,GO:0030036,GO:0030154,GO:0030182,GO:0032501,GO:0032502,GO:0032989,GO:0032991,GO:0033043,GO:0035152,GO:0035160,GO:0035556,GO:0036211,GO:0042551,GO:0042592,GO:0042981,GO:0043067,GO:0043085,GO:0043170,GO:0043234,GO:0043280,GO:0043281,GO:0043412,GO:0043933,GO:0044093,GO:0044237,GO:0044238,GO:0044260,GO:0044267,GO:0044424,GO:0044444,GO:0044445,GO:0044464,GO:0044699,GO:0044700,GO:0044707,GO:0044763,GO:0044767,GO:0045035,GO:0048468,GO:0048469,GO:0048513,GO:0048666,GO:0048699,GO:0048731,GO:0048856,GO:0048869,GO:0048871,GO:0050789,GO:0050790,GO:0050794,GO:0050896,GO:0051128,GO:0051301,GO:0051336,GO:0051345,GO:0051493,GO:0051716,GO:0052547,GO:0052548,GO:0060249,GO:0060541,GO:0065007,GO:0065008,GO:0065009,GO:0071704,GO:0071822,GO:0071840,GO:2000116,GO:2001056 | STYKc |

| | | |
|---|---|---|
| comp217754_c1_seq2.p1 | GO:0000122,GO:0000988,GO:0000989,GO:0001085,GO:0001102,GO:0001654,GO:0001666,GO:0002088,GO:0002089,GO:0002262,GO:0002376,GO:0002520,GO:0003002,GO:0003407,GO:0003674,GO:0003712,GO:0003714,GO:0003824,GO:0004672,GO:0004674,GO:0005488,GO:0005515,GO:0005575,GO:0005622,GO:0005623,GO:0005634,GO:0005635,GO:0005654,GO:0005737,GO:0005813,GO:0005815,GO:0005856,GO:0006355,GO:0006357,GO:0006464,GO:0006468,GO:0006793,GO:0006796,GO:0006915,GO:0006950,GO:0006974,GO:0006978,GO:0006996,GO:0006997,GO:0007154,GO:0007165,GO:0007166,GO:0007167,GO:0007178,GO:0007179,GO:0007224,GO:0007267,GO:0007275,GO:0007389,GO:0007399,GO:0007423,GO:0007610,GO:0007626,GO:0007628,GO:0008134,GO:0008150,GO:0008152,GO:0008219,GO:0008284,GO:0008344,GO:0008630,GO:0009628,GO:0009653,GO:0009719,GO:0009790,GO:0009887,GO:0009889,GO:0009890,GO:0009891,GO:0009892,GO:0009893,GO:0009952,GO:0009966,GO:0009967,GO:0009968,GO:0009987,GO:0010033,GO:0010468,GO:0010556,GO:0010557,GO:0010558,GO:0010604,GO:0010605,GO:0010627,GO:0010628,GO:0010629,GO:0010646,GO:0010647,GO:0010648,GO:0010740,GO:0010842,GO:0010941,GO:0012501,GO:0012505,GO:0015630,GO:0016020,GO:0016032,GO:0016043,GO:0016265,GO:0016301,GO:0016310,GO:0016604,GO:0016605,GO:0016740,GO:0016772,GO:0016773,GO:0017015,GO:0018105,GO:0018107,GO:0018193,GO:0018209,GO:0018210,GO:0019048,GO:0019219,GO:0019222,GO:0019538,GO:0022008,GO:0022603,GO:0023051,GO:0023052,GO:0023056,GO:0023057,GO:0030097,GO:0030099,GO:0030154,GO:0030182,GO:0030218,GO:0030330,GO:0030510,GO:0030511,GO:0030514,GO:0030534,GO:0030575,GO:0030578,GO:0031076,GO:0031090,GO:0031128,GO:0031323,GO:0031324,GO:0031325,GO:0031326,GO:0031327,GO:0031328,GO:0031965,GO:0031967,GO:0031974,GO:0031975,GO:0031981,GO:0032092,GO:0032501,GO:0032502,GO:0032872,GO:0032874,GO:0033554,GO:0033613,GO:0034101,GO:0035556,GO:0035821,GO:0036211,GO:0036293,GO:0036294,GO:0042127,GO:0042221,GO:0042592,GO:0042770,GO:0042771,GO:0042772,GO:0042981,GO:0043010,GO:0043066,GO:0043067,GO:0043069,GO:0043170,GO:0043226,GO:0043227,GO:0043228,GO:0043229,GO:0043231,GO:0043232,GO:0043233,GO:0043388,GO:0043393,GO:0043408,GO:0043410,GO:0043412,GO:0043523,GO:0043524,GO:0044003,GO:0044093,GO:0044237,GO:0044238,GO:0044260,GO:0044267,GO:0044403,GO:0044419,GO:0044422,GO:0044424,GO:0044428,GO:0044430,GO:0044446,GO:0044451,GO:0044464,GO:0044699,GO:0044700,GO:0044707,GO:0044708,GO:0044763,GO:0044764,GO:0044767,GO:0045165,GO:0045168,GO:0045765,GO:0045766,GO:0045892,GO:0045893,GO:0045934,GO:0045935,GO:0045944,GO:0046328,GO:0046330,GO:0046790,GO:0048048,GO:0048513,GO:0048518,GO:0048519,GO:0048522,GO:0048523,GO:0048534,GO:0048562,GO:0048568,GO:0048583,GO:0048584,GO:0048585,GO:0048592,GO:0048593,GO:0048596,GO:0048598,GO:0048646,GO:0048699,GO:0048731,GO:0048856,GO:0048869,GO:0048872,GO:0050789,GO:0050793,GO:0050794,GO:0050879,GO:0050881,GO:0050882,GO:0050896,GO:0051090,GO:0051091,GO:0051094,GO:0051098,GO:0051099,GO:0051101,GO:0051171,GO:0051172,GO:0051173,GO:0051239,GO:0051252,GO:0051253,GO:0051254,GO:0051701,GO:0051704,GO:0051716,GO:0051726,GO:0051817,GO:0060041,GO:0060042,GO:0060059,GO:0060235,GO:0060255,GO:0060395,GO:0060548,GO:0061072,GO:0065007,GO:0065008,GO:0065009,GO:0070013,GO:0070302,GO:0070304,GO:0070482,GO:0070848,GO:0070887,GO:0071310,GO:0071363,GO:0071453,GO:0071456,GO:0071495,GO:0071559,GO:0071560,GO:0071704,GO:0071840,GO:0072331,GO:0072332,GO:0080090,GO:0080134,GO:0080135,GO:0090092,GO:0090100,GO:0090101,GO:0090287,GO:0097190,GO:0097193,GO:1901214,GO:1901215,GO:1901342,GO:2000026,GO:2000112,GO:2000113,GO:2001141 | Homeodomain interacting protein kinase 2 |
| comp210329_c0_seq1.p1 | | cysteine-rich, DPF motif domain containing 1 |
| comp215725_c0_seq14.p1 | | Zinc finger protein |

| | | |
|---|---|---|
| comp201482_c0_seq1.p1 | GO:0000082,GO:0000166,GO:0000278,GO:0000902,GO:0000904,GO:0001558,GO:0001882,GO:0001883,GO:0003674,GO:0003824,GO:0003924,GO:0005488,GO:0005515,GO:0005525,GO:0006139,GO:0006152,GO:0006163,GO:0006184,GO:0006195,GO:0006417,GO:0006725,GO:0006753,GO:0006793,GO:0006796,GO:0006807,GO:0006928,GO:0006935,GO:0006950,GO:0007049,GO:0007275,GO:0007399,GO:0007409,GO:0007411,GO:0007416,GO:0007444,GO:0007446,GO:0007517,GO:0007519,GO:0007528,GO:0008150,GO:0008152,GO:0008361,GO:0009056,GO:0009116,GO:0009117,GO:0009119,GO:0009141,GO:0009143,GO:0009144,GO:0009146,GO:0009150,GO:0009154,GO:0009164,GO:0009166,GO:0009199,GO:0009203,GO:0009205,GO:0009207,GO:0009259,GO:0009261,GO:0009314,GO:0009416,GO:0009453,GO:0009605,GO:0009628,GO:0009653,GO:0009888,GO:0009889,GO:0009891,GO:0009893,GO:0009894,GO:0009987,GO:0009991,GO:0010468,GO:0010506,GO:0010556,GO:0010557,GO:0010604,GO:0010608,GO:0014706,GO:0016043,GO:0016049,GO:0016462,GO:0016787,GO:0016817,GO:0016818,GO:0017076,GO:0017111,GO:0019001,GO:0019222,GO:0019439,GO:0019637,GO:0019693,GO:0022008,GO:0022402,GO:0022403,GO:0022607,GO:0022613,GO:0030030,GO:0030154,GO:0030182,GO:0030307,GO:0031175,GO:0031323,GO:0031325,GO:0031326,GO:0031328,GO:0031329,GO:0031667,GO:0032268,GO:0032270,GO:0032501,GO:0032502,GO:0032535,GO:0032549,GO:0032550,GO:0032553,GO:0032555,GO:0032561,GO:0032989,GO:0032990,GO:0034641,GO:0034655,GO:0035265,GO:0035639,GO:0036094,GO:0040007,GO:0040008,GO:0040011,GO:0042221,GO:0042254,GO:0042278,GO:0042330,GO:0042331,GO:0042454,GO:0042594,GO:0042692,GO:0043167,GO:0043168,GO:0044085,GO:0044087,GO:0044089,GO:0044237,GO:0044238,GO:0044248,GO:0044270,GO:0044281,GO:0044699,GO:0044707,GO:0044710,GO:0044763,GO:0044767,GO:0044770,GO:0044772,GO:0045727,GO:0045793,GO:0045927,GO:0046039,GO:0046128,GO:0046130,GO:0046434,GO:0046483,GO:0046700,GO:0048468,GO:0048513,GO:0048518,GO:0048522,GO:0048589,GO:0048666,GO:0048667,GO:0048699,GO:0048731,GO:0048741,GO:0048747,GO:0048812,GO:0048856,GO:0048858,GO:0048869,GO:0050789,GO:0050794,GO:0050808,GO:0050896,GO:0051124,GO:0051128,GO:0051146,GO:0051246,GO:0051247,GO:0051325,GO:0051329,GO:0055001,GO:0055002,GO:0055086,GO:0060255,GO:0060537,GO:0060538,GO:0061061,GO:0061564,GO:0065007,GO:0065008,GO:0071704,GO:0071840,GO:0072521,GO:0072523,GO:0080090,GO:0090066,GO:0090069,GO:0090070,GO:0097159,GO:0097485,GO:1901068,GO:1901069,GO:1901135,GO:1901136,GO:1901265,GO:1901292,GO:1901360,GO:1901361,GO:1901363,GO:1901564,GO:1901565,GO:1901575,GO:1901657,GO:1901658,GO:2000112,GO:2000377 | GTP-binding protein Rheb homolog |
| comp211949_c1_seq2.p1 | | UDP glucuronosyltransferase 5 family, polypeptide |

| | | |
|---|---|---|
| comp618580_c0_seq1.p1 | GO:0000003,GO:0000082,GO:0000083,GO:0000278,GO:0000307,GO:0000988,GO:0000989,GO:0001541,GO:0001547,GO:0001889,GO:0001932,GO:0003006,GO:0003674,GO:0003712,GO:0003713,GO:0003824,GO:0005102,GO:0005488,GO:0005515,GO:0005575,GO:0005622,GO:0005623,GO:0005634,GO:0005654,GO:0005737,GO:0005813,GO:0005815,GO:0005829,GO:0005856,GO:0006139,GO:0006259,GO:0006260,GO:0006261,GO:0006270,GO:0006355,GO:0006357,GO:0006464,GO:0006468,GO:0006725,GO:0006793,GO:0006796,GO:0006807,GO:0007049,GO:0007154,GO:0007165,GO:0007166,GO:0007275,GO:0007548,GO:0007584,GO:0008150,GO:0008152,GO:0008406,GO:0008585,GO:0009058,GO:0009059,GO:0009605,GO:0009719,GO:0009725,GO:0009889,GO:0009891,GO:0009893,GO:0009987,GO:0009991,GO:0010033,GO:0010243,GO:0010468,GO:0010556,GO:0010557,GO:0010604,GO:0010628,GO:0014070,GO:0014074,GO:0015630,GO:0016055,GO:0016301,GO:0016310,GO:0016538,GO:0016740,GO:0016772,GO:0019207,GO:0019219,GO:0019220,GO:0019222,GO:0019538,GO:0019887,GO:0022402,GO:0022403,GO:0022414,GO:0022602,GO:0023052,GO:0030234,GO:0030518,GO:0030521,GO:0030522,GO:0031099,GO:0031100,GO:0031323,GO:0031325,GO:0031326,GO:0031328,GO:0031399,GO:0031667,GO:0031668,GO:0031669,GO:0031670,GO:0031960,GO:0031974,GO:0031981,GO:0032268,GO:0032355,GO:0032501,GO:0032502,GO:0032504,GO:0032570,GO:0032991,GO:0033197,GO:0033273,GO:0033993,GO:0034097,GO:0034641,GO:0034645,GO:0035257,GO:0035258,GO:0036211,GO:0040007,GO:0042221,GO:0042325,GO:0042493,GO:0042698,GO:0043170,GO:0043226,GO:0043227,GO:0043228,GO:0043229,GO:0043231,GO:0043232,GO:0043233,GO:0043234,GO:0043412,GO:0043549,GO:0043627,GO:0044237,GO:0044238,GO:0044249,GO:0044260,GO:0044267,GO:0044422,GO:0044424,GO:0044428,GO:0044430,GO:0044444,GO:0044446,GO:0044464,GO:0044699,GO:0044700,GO:0044702,GO:0044707,GO:0044763,GO:0044767,GO:0044770,GO:0044772,GO:0045137,GO:0045471,GO:0045595,GO:0045597,GO:0045859,GO:0045893,GO:0045935,GO:0046483,GO:0046545,GO:0046660,GO:0048511,GO:0048513,GO:0048518,GO:0048522,GO:0048545,GO:0048589,GO:0048608,GO:0048609,GO:0048731,GO:0048856,GO:0050681,GO:0050789,GO:0050790,GO:0050793,GO:0050794,GO:0050896,GO:0051094,GO:0051171,GO:0051173,GO:0051174,GO:0051246,GO:0051252,GO:0051254,GO:0051325,GO:0051329,GO:0051338,GO:0051384,GO:0051385,GO:0051412,GO:0051427,GO:0051597,GO:0051716,GO:0051726,GO:0060255,GO:0061008,GO:0061458,GO:0065007,GO:0065009,GO:0070013,GO:0070887,GO:0071496,GO:0071704,GO:0080090,GO:0090304,GO:0097305,GO:1901360,GO:1901576,GO:1901654,GO:1901698,GO:1901700,GO:2000112,GO:2001141 | cyclin E1 |
| comp213602_c0_seq8.p1 | | Inherit from veNOG: spindle apparatus coiled-coil protein 1 |

| | | |
|---|---|---|
| comp216403_c2_seq2.p1 | GO:0000003,GO:0000146,GO:0000166,GO:0000302,GO:0001667,GO:0001701,GO:0001725,GO:0001756,GO:0001757,GO:0001778,GO:0001882,GO:0001883,GO:0002026,GO:0002027,GO:0002119,GO:0002164,GO:0003002,GO:0003006,GO:0003007,GO:0003008,GO:0003009,GO:0003010,GO:0003012,GO:0003013,GO:0003015,GO:0003205,GO:0003206,GO:0003208,GO:0003209,GO:0003228,GO:0003229,GO:0003230,GO:0003231,GO:0003674,GO:0003774,GO:0003779,GO:0003824,GO:0005198,GO:0005488,GO:0005515,GO:0005516,GO:0005524,GO:0005575,GO:0005622,GO:0005623,GO:0005634,GO:0005694,GO:0005700,GO:0005703,GO:0005730,GO:0005737,GO:0005794,GO:0005826,GO:0005829,GO:0005856,GO:0005859,GO:0005863,GO:0005911,GO:0005912,GO:0005924,GO:0005925,GO:0005938,GO:0006139,GO:0006140,GO:0006152,GO:0006163,GO:0006195,GO:0006200,GO:0006355,GO:0006357,GO:0006461,GO:0006725,GO:0006753,GO:0006793,GO:0006796,GO:0006807,GO:0006810,GO:0006897,GO:0006898,GO:0006928,GO:0006936,GO:0006941,GO:0006950,GO:0006979,GO:0006996,GO:0007009,GO:0007010,GO:0007015,GO:0007154,GO:0007165,GO:0007166,GO:0007167,GO:0007178,GO:0007275,GO:0007276,GO:0007292,GO:0007297,GO:0007298,GO:0007379,GO:0007389,GO:0007399,GO:0007413,GO:0007424,GO:0007427,GO:0007507,GO:0007512,GO:0007517,GO:0007519,GO:0007522,GO:0007525,GO:0007527,GO:0007610,GO:0007626,GO:0007631,GO:0008015,GO:0008016,GO:0008037,GO:0008038,GO:0008092,GO:0008150,GO:0008152,GO:0008217,GO:0008307,GO:0009056,GO:0009116,GO:0009117,GO:0009118,GO:0009119,GO:0009123,GO:0009125,GO:0009126,GO:0009128,GO:0009141,GO:0009143,GO:0009144,GO:0009146,GO:0009150,GO:0009154,GO:0009158,GO:0009161,GO:0009164,GO:0009166,GO:0009167,GO:0009169,GO:0009199,GO:0009203,GO:0009205,GO:0009207,GO:0009259,GO:0009261,GO:0009611,GO:0009653,GO:0009790,GO:0009791,GO:0009792,GO:0009880,GO:0009887,GO:0009888,GO:0009889,GO:0009894,GO:0009952,GO:0009987,GO:0010171,GO:0010468,GO:0010556,GO:0010608,GO:0010631,GO:0010927,GO:0014704,GO:0014706,GO:0014721,GO:0014823,GO:0014866,GO:0015629,GO:0016020,GO:0016043,GO:0016044,GO:0016055,GO:0016192,GO:0016459,GO:0016460,GO:0016462,GO:0016477,GO:0016787,GO:0016817,GO:0016818,GO:0016887,GO:0017076,GO:0017111,GO:0018991,GO:0019098,GO:0019219,GO:0019220,GO:0019222,GO:0019439,GO:0019637,GO:0019693,GO:0019953,GO:0022008,GO:0022412,GO:0022414,GO:0022603,GO:0022607,GO:0023052,GO:0030016,GO:0030017,GO:0030018,GO:0030029,GO:0030030,GO:0030036,GO:0030048,GO:0030049,GO:0030054,GO:0030055,GO:0030154,GO:0030182,GO:0030239,GO:0030240,GO:0030241,GO:0030509,GO:0030554,GO:0030707,GO:0030811,GO:0030898,GO:0030899,GO:0031032,GO:0031033,GO:0031034,GO:0031175,GO:0031323,GO:0031326,GO:0031329,GO:0031444,GO:0031647,GO:0031672,GO:0031674,GO:0031974,GO:0031981,GO:0032153,GO:0032155,GO:0032432,GO:0032501,GO:0032502,GO:0032504,GO:0032549,GO:0032550,GO:0032553,GO:0032555,GO:0032559,GO:0032956,GO:0032970,GO:0032982,GO:0032989,GO:0032991,GO:0033043,GO:0033057,GO:0033121,GO:0033275,GO:0034622,GO:0034641,GO:0034655,GO:0035050,GO:0035051,GO:0035282,GO:0035295,GO:0035556,GO:0035639,GO:0036094,GO:0036379,GO:0040007,GO:0040008,GO:0040011,GO:0040012,GO:0040017,GO:0042060,GO:0042221,GO:0042278,GO:0042454,GO:0042623,GO:0042641,GO:0042692,GO:0042755,GO:0042802,GO:0042803,GO:0043009,GO:0043050,GO:0043167,GO:0043168,GO:0043226,GO:0043227,GO:0043228,GO:0043229,GO:0043231,GO:0043232,GO:0043233,GO:0043234,GO:0043292,GO:0043462,GO:0043623,GO:0043933,GO:0044057,GO:0044085,GO:0044087,GO:0044237,GO:0044238,GO:0044248,GO:0044270,GO:0044281,GO:0044291,GO:0044422,GO:0044424,GO:0044427,GO:0044428,GO:0044430,GO:0044444,GO:0044446,GO:0044448,GO:0044449,GO:0044464,GO:0044699,GO:0044700,GO:0044702,GO:0044706,GO:0044707,GO:0044708,GO:0044710,GO:0044763,GO:0044767,GO:0045214,GO:0045595,GO:0046034,GO:0046128,GO:0046130,GO:0046434,GO:0046483,GO:0046620,GO:0046700,GO:0046982,GO:0046983,GO:0048017,GO:0048468,GO:0048477,GO:0048513,GO:0048518,GO:0048568,GO:0048609,GO:0048610,GO:0048638,GO:0048644,GO:0048646,GO:0048666,GO:0048699,GO:0048729,GO:0048731,GO:0048738,GO:0048739,GO:0048741,GO:0048747,GO:0048856,GO:0048869,GO:0048870,GO:0050789,GO:0050790,GO:0050793,GO:0050794,GO:0050821,GO:0050879,GO:0050881,GO:0050896,GO:0051015,GO:0051128,GO:0051146,GO:0051147,GO | Myosin heavy chain |

| | | |
|---|---|---|
| comp211180_c0_seq6.p1 | GO:0000902,GO:0000904,GO:0001838,GO:0001841,GO:0001843,GO:0001932,GO:0001934,GO:0002009,GO:0003674,GO:0004871,GO:0004872,GO:0005575,GO:0005622,GO:0005623,GO:0005737,GO:0005783,GO:0005886,GO:0006355,GO:0006357,GO:0006928,GO:0006935,GO:0007154,GO:0007165,GO:0007166,GO:0007167,GO:0007178,GO:0007275,GO:0007399,GO:0007409,GO:0007411,GO:0008150,GO:0009605,GO:0009653,GO:0009790,GO:0009792,GO:0009888,GO:0009889,GO:0009893,GO:0009966,GO:0009986,GO:0009987,GO:0010468,GO:0010556,GO:0010562,GO:0010604,GO:0010646,GO:0014020,GO:0015026,GO:0016020,GO:0016043,GO:0016331,GO:0019219,GO:0019220,GO:0019222,GO:0021915,GO:0022008,GO:0023051,GO:0023052,GO:0030030,GO:0030154,GO:0030182,GO:0030509,GO:0030510,GO:0031175,GO:0031323,GO:0031325,GO:0031326,GO:0031399,GO:0031401,GO:0032268,GO:0032270,GO:0032501,GO:0032502,GO:0032989,GO:0032990,GO:0035148,GO:0035239,GO:0035295,GO:0038023,GO:0040011,GO:0042221,GO:0042325,GO:0042327,GO:0042330,GO:0043009,GO:0043226,GO:0043227,GO:0043229,GO:0043231,GO:0044424,GO:0044444,GO:0044464,GO:0044699,GO:0044700,GO:0044707,GO:0044763,GO:0044767,GO:0045937,GO:0048468,GO:0048518,GO:0048522,GO:0048583,GO:0048598,GO:0048646,GO:0048666,GO:0048667,GO:0048699,GO:0048729,GO:0048731,GO:0048812,GO:0048856,GO:0048858,GO:0048869,GO:0050789,GO:0050794,GO:0050896,GO:0051171,GO:0051174,GO:0051246,GO:0051247,GO:0051252,GO:0051716,GO:0060089,GO:0060255,GO:0060429,GO:0060562,GO:0060606,GO:0061564,GO:0065007,GO:0071840,GO:0071944,GO:0072175,GO:0080090,GO:0090092,GO:0097485,GO:2000112,GO:2001141 | RGM domain family, member A |
| comp214589_c0_seq5.p1 | GO:0000003,GO:0000228,GO:0000793,GO:0000794,GO:0000795,GO:0001673,GO:0003674,GO:0003676,GO:0003677,GO:0003682,GO:0003696,GO:0003697,GO:0003824,GO:0005488,GO:0005515,GO:0005575,GO:0005622,GO:0005623,GO:0005634,GO:0005654,GO:0005694,GO:0005712,GO:0006139,GO:0006152,GO:0006163,GO:0006195,GO:0006200,GO:0006259,GO:0006281,GO:0006298,GO:0006310,GO:0006725,GO:0006753,GO:0006793,GO:0006796,GO:0006807,GO:0006950,GO:0006974,GO:0006996,GO:0007049,GO:0007126,GO:0007127,GO:0007129,GO:0007130,GO:0007131,GO:0007140,GO:0007143,GO:0007144,GO:0008104,GO:0008150,GO:0008152,GO:0009056,GO:0009116,GO:0009117,GO:0009119,GO:0009123,GO:0009125,GO:0009126,GO:0009128,GO:0009141,GO:0009143,GO:0009144,GO:0009146,GO:0009150,GO:0009154,GO:0009158,GO:0009161,GO:0009164,GO:0009166,GO:0009167,GO:0009169,GO:0009199,GO:0009203,GO:0009205,GO:0009207,GO:0009259,GO:0009261,GO:0009987,GO:0016043,GO:0016462,GO:0016787,GO:0016817,GO:0016818,GO:0016887,GO:0017111,GO:0019237,GO:0019439,GO:0019637,GO:0019693,GO:0022402,GO:0022607,GO:0031974,GO:0031981,GO:0032300,GO:0032390,GO:0032403,GO:0032404,GO:0032407,GO:0032991,GO:0033036,GO:0033554,GO:0034641,GO:0034655,GO:0035825,GO:0042278,GO:0042454,GO:0043073,GO:0043170,GO:0043226,GO:0043227,GO:0043228,GO:0043229,GO:0043231,GO:0043232,GO:0043233,GO:0043234,GO:0043565,GO:0043566,GO:0044085,GO:0044237,GO:0044238,GO:0044248,GO:0044260,GO:0044270,GO:0044281,GO:0044422,GO:0044424,GO:0044427,GO:0044428,GO:0044446,GO:0044454,GO:0044464,GO:0044699,GO:0044710,GO:0044763,GO:0046034,GO:0046128,GO:0046130,GO:0046434,GO:0046483,GO:0046700,GO:0048610,GO:0050896,GO:0051179,GO:0051276,GO:0051321,GO:0051716,GO:0055086,GO:0070013,GO:0070192,GO:0070193,GO:0071704,GO:0071840,GO:0072521,GO:0072523,GO:0090304,GO:0097159,GO:1901135,GO:1901136,GO:1901292,GO:1901360,GO:1901361,GO:1901363,GO:1901564,GO:1901565,GO:1901575,GO:1901657,GO:1901658 | MutL homolog 3 (E coli) |

| | | |
|---|---|---|
| comp216938_c0_seq6.p1 | GO:0000228,GO:0000775,GO:0000779,GO:0000780,GO:0000785,GO:0000792,GO:0000793,GO:0000794,GO:0000922,GO:0005575,GO:0005622,GO:0005623,GO:0005634,GO:0005694,GO:0005721,GO:0005819,GO:0005856,GO:0007275,GO:0007399,GO:0008150,GO:0008278,GO:0009987,GO:0015630,GO:0016322,GO:0021700,GO:0022008,GO:0030154,GO:0030182,GO:0031974,GO:0031981,GO:0032501,GO:0032502,GO:0032991,GO:0035327,GO:0042551,GO:0043226,GO:0043227,GO:0043228,GO:0043229,GO:0043231,GO:0043232,GO:0043233,GO:0043234,GO:0044422,GO:0044424,GO:0044427,GO:0044428,GO:0044430,GO:0044446,GO:0044454,GO:0044464,GO:0044699,GO:0044707,GO:0044763,GO:0044767,GO:0048468,GO:0048469,GO:0048666,GO:0048699,GO:0048731,GO:0048856,GO:0048869,GO:0070013 | Stromal antigen |
| comp219221_c0_seq3.p1 | GO:0000723,GO:0000781,GO:0000785,GO:0000791,GO:0000792,GO:0002165,GO:0003674,GO:0005488,GO:0005515,GO:0005575,GO:0005622,GO:0005623,GO:0005634,GO:0005694,GO:0005700,GO:0005705,GO:0006066,GO:0006139,GO:0006259,GO:0006355,GO:0006357,GO:0006629,GO:0006694,GO:0006697,GO:0006725,GO:0006807,GO:0006996,GO:0007275,GO:0007399,GO:0008150,GO:0008152,GO:0008202,GO:0008205,GO:0008610,GO:0009058,GO:0009791,GO:0009889,GO:0009987,GO:0010468,GO:0010556,GO:0010817,GO:0016043,GO:0016125,GO:0016126,GO:0016233,GO:0019219,GO:0019222,GO:0022008,GO:0030154,GO:0031323,GO:0031326,GO:0032200,GO:0032501,GO:0032502,GO:0032991,GO:0034641,GO:0034754,GO:0035012,GO:0042180,GO:0042181,GO:0042445,GO:0042446,GO:0042592,GO:0043170,GO:0043226,GO:0043227,GO:0043228,GO:0043229,GO:0043231,GO:0043232,GO:0043234,GO:0044237,GO:0044238,GO:0044249,GO:0044260,GO:0044281,GO:0044283,GO:0044422,GO:0044424,GO:0044427,GO:0044446,GO:0044464,GO:0044699,GO:0044707,GO:0044710,GO:0044711,GO:0044763,GO:0045455,GO:0045456,GO:0046165,GO:0046483,GO:0048731,GO:0048856,GO:0048869,GO:0050789,GO:0050794,GO:0051171,GO:0051252,GO:0051276,GO:0060249,GO:0060255,GO:0065007,GO:0065008,GO:0071704,GO:0071840,GO:0080090,GO:0090304,GO:1901360,GO:1901362,GO:1901576,GO:1901615,GO:1901617,GO:2000112,GO:2001141 | zinc finger MYM-type protein 4-like |
| comp219221_c0_seq3.p3 | GO:0000902,GO:0000904,GO:0003008,GO:0003674,GO:0003824,GO:0004721,GO:0004725,GO:0004728,GO:0004871,GO:0004872,GO:0004888,GO:0005001,GO:0005057,GO:0005575,GO:0005623,GO:0005886,GO:0006464,GO:0006470,GO:0006793,GO:0006796,GO:0006928,GO:0006935,GO:0007154,GO:0007165,GO:0007275,GO:0007399,GO:0007409,GO:0007411,GO:0007417,GO:0007424,GO:0007610,GO:0007611,GO:0007613,GO:0007616,GO:0008045,GO:0008150,GO:0008152,GO:0009605,GO:0009653,GO:0009987,GO:0016020,GO:0016043,GO:0016310,GO:0016311,GO:0016787,GO:0016788,GO:0016791,GO:0019198,GO:0019538,GO:0022008,GO:0023014,GO:0023052,GO:0030030,GO:0030154,GO:0030182,GO:0030424,GO:0031175,GO:0032501,GO:0032502,GO:0032989,GO:0032990,GO:0035335,GO:0036211,GO:0038023,GO:0040011,GO:0042221,GO:0042330,GO:0042578,GO:0042995,GO:0043005,GO:0043170,GO:0043412,GO:0044237,GO:0044238,GO:0044260,GO:0044267,GO:0044464,GO:0044699,GO:0044700,GO:0044707,GO:0044708,GO:0044763,GO:0044767,GO:0045177,GO:0048468,GO:0048666,GO:0048667,GO:0048699,GO:0048731,GO:0048812,GO:0048856,GO:0048858,GO:0048869,GO:0050789,GO:0050794,GO:0050877,GO:0050890,GO:0050896,GO:0051716,GO:0060089,GO:0060541,GO:0061564,GO:0065007,GO:0071704,GO:0071840,GO:0071944,GO:0097458,GO:0097485 | Tissue factor |

| | | |
|---|---|---|
| comp217807_c0_seq5.p1 | GO:0000122,GO:0000414,GO:0000415,GO:0000975,GO:0000988,GO:0000989,GO:0001067,GO:0003674,GO:0003676,GO:0003677,GO:0003712,GO:0003714,GO:0005488,GO:0005515,GO:0005575,GO:0005622,GO:0005623,GO:0005634,GO:0006325,GO:0006355,GO:0006357,GO:0006464,GO:0006513,GO:0006996,GO:0007275,GO:0007389,GO:0007507,GO:0008150,GO:0008152,GO:0009653,GO:0009798,GO:0009887,GO:0009889,GO:0009890,GO:0009892,GO:0009987,GO:0010390,GO:0010468,GO:0010556,GO:0010558,GO:0010605,GO:0010629,GO:0010639,GO:0016043,GO:0016567,GO:0016568,GO:0016569,GO:0016570,GO:0016574,GO:0019219,GO:0019222,GO:0019538,GO:0022603,GO:0030278,GO:0030279,GO:0030500,GO:0030502,GO:0031056,GO:0031057,GO:0031060,GO:0031061,GO:0031072,GO:0031323,GO:0031324,GO:0031326,GO:0031327,GO:0031399,GO:0031400,GO:0032268,GO:0032269,GO:0032446,GO:0032501,GO:0032502,GO:0033043,GO:0033044,GO:0033522,GO:0035518,GO:0036211,GO:0042476,GO:0042481,GO:0043170,GO:0043226,GO:0043227,GO:0043229,GO:0043231,GO:0043412,GO:0044212,GO:0044237,GO:0044238,GO:0044260,GO:0044267,GO:0044424,GO:0044464,GO:0044699,GO:0044707,GO:0044763,GO:0044767,GO:0045892,GO:0045934,GO:0048513,GO:0048519,GO:0048523,GO:0048731,GO:0048856,GO:0050789,GO:0050793,GO:0050794,GO:0051093,GO:0051128,GO:0051129,GO:0051171,GO:0051172,GO:0051239,GO:0051241,GO:0051246,GO:0051248,GO:0051252,GO:0051253,GO:0051276,GO:0051569,GO:0051572,GO:0060021,GO:0060255,GO:0065001,GO:0065007,GO:0070167,GO:0070168,GO:0070170,GO:0070171,GO:0070647,GO:0071704,GO:0071840,GO:0072358,GO:0072359,GO:0080090,GO:0097159,GO:1901363,GO:1902275,GO:2000026,GO:2000027,GO:2000112,GO:2000113,GO:2001141,GO:2001251 | BCL6 corepressor |

| | | |
|---|---|---|
| comp220758_c0_seq9.p1 | GO:0000122,GO:0000165,GO:0001775,GO:0001932,GO:0001934,GO:0002327,GO:0002376,GO:0002520,GO:0002521,GO:0002682,GO:0002902,GO:0002904,GO:0003674,GO:0004871,GO:0005083,GO:0005085,GO:0005088,GO:0005096,GO:0005099,GO:0005488,GO:0005509,GO:0005515,GO:0005575,GO:0005576,GO:0005578,GO:0005622,GO:0005623,GO:0005737,GO:0005886,GO:0005887,GO:0006140,GO:0006355,GO:0006357,GO:0006915,GO:0006950,GO:0007154,GO:0007165,GO:0007243,GO:0007264,GO:0007265,GO:0007275,GO:0008047,GO:0008150,GO:0008219,GO:0008289,GO:0008544,GO:0009118,GO:0009267,GO:0009605,GO:0009888,GO:0009889,GO:0009890,GO:0009892,GO:0009893,GO:0009894,GO:0009966,GO:0009968,GO:0009987,GO:0009991,GO:0010466,GO:0010468,GO:0010556,GO:0010558,GO:0010562,GO:0010604,GO:0010605,GO:0010629,GO:0010646,GO:0010648,GO:0010941,GO:0010942,GO:0010951,GO:0012501,GO:0016020,GO:0016021,GO:0016265,GO:0017016,GO:0017034,GO:0018996,GO:0019219,GO:0019220,GO:0019222,GO:0019899,GO:0019932,GO:0019933,GO:0019935,GO:0019992,GO:0023051,GO:0023052,GO:0023057,GO:0030097,GO:0030098,GO:0030154,GO:0030183,GO:0030234,GO:0030695,GO:0030742,GO:0030811,GO:0031012,GO:0031224,GO:0031226,GO:0031267,GO:0031323,GO:0031324,GO:0031325,GO:0031326,GO:0031327,GO:0031329,GO:0031399,GO:0031401,GO:0031644,GO:0031667,GO:0031668,GO:0031669,GO:0031929,GO:0032006,GO:0032007,GO:0032268,GO:0032270,GO:0032317,GO:0032318,GO:0032320,GO:0032487,GO:0032501,GO:0032502,GO:0032854,GO:0033121,GO:0033124,GO:0033554,GO:0035556,GO:0040002,GO:0042113,GO:0042303,GO:0042325,GO:0042327,GO:0042335,GO:0042594,GO:0042981,GO:0043065,GO:0043067,GO:0043068,GO:0043085,GO:0043086,GO:0043087,GO:0043154,GO:0043167,GO:0043169,GO:0043281,GO:0043547,GO:0043588,GO:0044057,GO:0044092,GO:0044093,GO:0044421,GO:0044424,GO:0044425,GO:0044459,GO:0044464,GO:0044699,GO:0044700,GO:0044707,GO:0044763,GO:0044767,GO:0045321,GO:0045595,GO:0045892,GO:0045934,GO:0045937,GO:0046578,GO:0046582,GO:0046649,GO:0046872,GO:0048167,GO:0048513,GO:0048518,GO:0048519,GO:0048522,GO:0048523,GO:0048534,GO:0048583,GO:0048585,GO:0048731,GO:0048856,GO:0048869,GO:0050789,GO:0050790,GO:0050793,GO:0050794,GO:0050804,GO:0050896,GO:0051020,GO:0051056,GO:0051171,GO:0051172,GO:0051174,GO:0051239,GO:0051246,GO:0051247,GO:0051252,GO:0051253,GO:0051336,GO:0051345,GO:0051346,GO:0051716,GO:0051969,GO:0052547,GO:0052548,GO:0060089,GO:0060102,GO:0060255,GO:0060589,GO:0065007,GO:0065008,GO:0065009,GO:0070228,GO:0070230,GO:0071496,GO:0071944,GO:0080090,GO:1900542,GO:1901532,GO:2000026,GO:2000106,GO:2000108,GO:2000112,GO:2000113,GO:2000116,GO:2000117,GO:2000973,GO:2001141 | Rap guanine nucleotide exchange factor (GEF) |
| comp198260_c0_seq4.p1 | GO:0005575,GO:0005622,GO:0005623,GO:0005634,GO:0005737,GO:0005886,GO:0016020,GO:0043226,GO:0043227,GO:0043229,GO:0043231,GO:0044424,GO:0044464,GO:0071944 | coiled-coil domain containing 62 |
| comp215994_c1_seq8.p1 | GO:0005575,GO:0005622,GO:0005623,GO:0005634,GO:0043226,GO:0043227,GO:0043229,GO:0043231,GO:0044424,GO:0044464 | Zinc finger protein |

| | | |
|---|---|---|
| comp218715_c1_seq2.p1 | GO:0000003,GO:0000226,GO:0000578,GO:0000902,GO:0000904,GO:0001505,GO:0001654,GO:0001754,GO:0003001,GO:0003002,GO:0003006,GO:0003008,GO:0003674,GO:0003774,GO:0003777,GO:0003824,GO:0005488,GO:0005515,GO:0005575,GO:0005622,GO:0005623,GO:0005737,GO:0005856,GO:0005871,GO:0005874,GO:0005875,GO:0006403,GO:0006810,GO:0006836,GO:0006886,GO:0006928,GO:0006935,GO:0006996,GO:0007005,GO:0007010,GO:0007017,GO:0007018,GO:0007028,GO:0007154,GO:0007163,GO:0007267,GO:0007268,GO:0007269,GO:0007275,GO:0007276,GO:0007281,GO:0007292,GO:0007300,GO:0007303,GO:0007308,GO:0007309,GO:0007310,GO:0007314,GO:0007315,GO:0007316,GO:0007317,GO:0007350,GO:0007351,GO:0007389,GO:0007399,GO:0007409,GO:0007411,GO:0007423,GO:0007517,GO:0007519,GO:0008017,GO:0008088,GO:0008092,GO:0008103,GO:0008104,GO:0008150,GO:0008152,GO:0008298,GO:0008358,GO:0008574,GO:0008595,GO:0009605,GO:0009653,GO:0009790,GO:0009798,GO:0009880,GO:0009887,GO:0009888,GO:0009948,GO:0009950,GO:0009952,GO:0009953,GO:0009987,GO:0009994,GO:0010970,GO:0014706,GO:0015031,GO:0015630,GO:0015631,GO:0016043,GO:0016325,GO:0016358,GO:0016462,GO:0016482,GO:0016787,GO:0016817,GO:0016818,GO:0017111,GO:0019094,GO:0019226,GO:0019227,GO:0019953,GO:0021700,GO:0022008,GO:0022412,GO:0022414,GO:0022607,GO:0023052,GO:0023061,GO:0030011,GO:0030030,GO:0030154,GO:0030182,GO:0030705,GO:0030951,GO:0030952,GO:0031175,GO:0032501,GO:0032502,GO:0032504,GO:0032879,GO:0032940,GO:0032989,GO:0032990,GO:0032991,GO:0033036,GO:0034613,GO:0034643,GO:0035282,GO:0035371,GO:0035637,GO:0040011,GO:0040023,GO:0042221,GO:0042330,GO:0042692,GO:0043226,GO:0043228,GO:0043229,GO:0043232,GO:0043234,GO:0044085,GO:0044087,GO:0044422,GO:0044424,GO:0044430,GO:0044446,GO:0044464,GO:0044699,GO:0044700,GO:0044702,GO:0044707,GO:0044763,GO:0044765,GO:0044767,GO:0045184,GO:0045451,GO:0045595,GO:0045995,GO:0046530,GO:0046903,GO:0046907,GO:0047497,GO:0048311,GO:0048312,GO:0048468,GO:0048469,GO:0048477,GO:0048513,GO:0048592,GO:0048599,GO:0048609,GO:0048610,GO:0048666,GO:0048667,GO:0048699,GO:0048731,GO:0048741,GO:0048747,GO:0048812,GO:0048813,GO:0048856,GO:0048858,GO:0048869,GO:0050789,GO:0050793,GO:0050794,GO:0050877,GO:0050896,GO:0051128,GO:0051146,GO:0051179,GO:0051234,GO:0051239,GO:0051640,GO:0051641,GO:0051646,GO:0051647,GO:0051649,GO:0051654,GO:0051656,GO:0055001,GO:0055002,GO:0060281,GO:0060284,GO:0060341,GO:0060537,GO:0060538,GO:0060810,GO:0060811,GO:0061061,GO:0061564,GO:0065007,GO:0065008,GO:0070727,GO:0071702,GO:0071840,GO:0072384,GO:0097485,GO:2000026,GO:2000241 | KISc |
| comp220695_c0_seq7.p1 | | chromosome 10 open reading frame 118 |
| comp208817_c2_seq1.p1 | GO:0003674,GO:0003824,GO:0005575,GO:0005622,GO:0005623,GO:0005737,GO:0005739,GO:0005777,GO:0005811,GO:0005856,GO:0005875,GO:0008150,GO:0008152,GO:0015630,GO:0016491,GO:0016614,GO:0016616,GO:0032991,GO:0042579,GO:0043226,GO:0043227,GO:0043228,GO:0043229,GO:0043231,GO:0043232,GO:0043234,GO:0044422,GO:0044424,GO:0044430,GO:0044444,GO:0044446,GO:0044464,GO:0044710,GO:0055114 | Hydroxysteroid dehydrogenase like 2 |
| comp213187_c0_seq1.p1 | GO:0003674,GO:0005215,GO:0005575,GO:0006810,GO:0006811,GO:0006820,GO:0008150,GO:0008509,GO:0008514,GO:0009987,GO:0015075,GO:0015711,GO:0016020,GO:0016021,GO:0022857,GO:0022891,GO:0022892,GO:0031224,GO:0034220,GO:0044425,GO:0044699,GO:0044763,GO:0044765,GO:0051179,GO:0051234,GO:0055085,GO:0071702 | Organic Anion Transporter Polypeptide (OATP) family |

| | | |
|---|---|---|
| comp218138_c0_seq2.p1 | GO:0003674,GO:0005080,GO:0005488,GO:0005515,GO:0005575,GO:0005622,GO:0005623,GO:0005737,GO:0005856,GO:0006996,GO:0007010,GO:0008092,GO:0008150,GO:0009653,GO:0009987,GO:0010927,GO:0016043,GO:0019899,GO:0019900,GO:0019901,GO:0022607,GO:0030016,GO:0030017,GO:0030018,GO:0030029,GO:0030036,GO:0030154,GO:0030239,GO:0031032,GO:0031143,GO:0031674,GO:0032502,GO:0032989,GO:0042692,GO:0042805,GO:0042995,GO:0043226,GO:0043228,GO:0043229,GO:0043232,GO:0043292,GO:0044085,GO:0044422,GO:0044424,GO:0044444,GO:0044449,GO:0044464,GO:0044699,GO:0044763,GO:0044767,GO:0045214,GO:0048468,GO:0048471,GO:0048646,GO:0048856,GO:0048869,GO:0051146,GO:0051371,GO:0051393,GO:0055001,GO:0055002,GO:0061061,GO:0070925,GO:0071840 | LIM domain binding 3 |
| comp208775_c0_seq5.p1 | | LRR_TYP |
| comp216693_c3_seq1.p1 | GO:0003674,GO:0003824,GO:0004175,GO:0004222,GO:0006508,GO:0008150,GO:0008152,GO:0008233,GO:0008237,GO:0016787,GO:0019538,GO:0043170,GO:0044238,GO:0070011,GO:0071704 | Thrombospondin type 1 domain |
| comp208826_c0_seq2.p1 | GO:0000003,GO:0005575,GO:0005622,GO:0005623,GO:0005737,GO:0006915,GO:0007276,GO:0007283,GO:0008150,GO:0008219,GO:0009987,GO:0012501,GO:0016265,GO:0019953,GO:0022414,GO:0032501,GO:0032504,GO:0044424,GO:0044464,GO:0044699,GO:0044702,GO:0044763,GO:0048232,GO:0048609 | Spermatogenesis associated 4 |
| comp199910_c0_seq2.p1 | | Inherit from KOG: leucine rich repeat containing |
| comp217171_c0_seq1.p1 | GO:0000323,GO:0000407,GO:0003674,GO:0005488,GO:0005515,GO:0005575,GO:0005622,GO:0005623,GO:0005737,GO:0005764,GO:0005768,GO:0005770,GO:0005773,GO:0005776,GO:0005829,GO:0006461,GO:0006914,GO:0006950,GO:0007154,GO:0008150,GO:0008152,GO:0009056,GO:0009267,GO:0009605,GO:0009966,GO:0009987,GO:0009991,GO:0010627,GO:0010646,GO:0016043,GO:0016236,GO:0019899,GO:0019900,GO:0019901,GO:0022607,GO:0023051,GO:0030016,GO:0030017,GO:0030278,GO:0030500,GO:0031430,GO:0031667,GO:0031668,GO:0031669,GO:0031672,GO:0032182,GO:0032872,GO:0033554,GO:0042594,GO:0043130,GO:0043226,GO:0043227,GO:0043228,GO:0043229,GO:0043231,GO:0043232,GO:0043292,GO:0043408,GO:0043933,GO:0044085,GO:0044237,GO:0044248,GO:0044422,GO:0044424,GO:0044444,GO:0044449,GO:0044464,GO:0044699,GO:0044763,GO:0045595,GO:0045596,GO:0045667,GO:0045668,GO:0048519,GO:0048523,GO:0048583,GO:0050789,GO:0050793,GO:0050794,GO:0050896,GO:0051019,GO:0051093,GO:0051239,GO:0051259,GO:0051716,GO:0065003,GO:0065007,GO:0070167,GO:0070271,GO:0070302,GO:0071496,GO:0071822,GO:0071840,GO:0080134,GO:0080135,GO:2000026 | Zinc finger, ZZ type |
| comp218425_c1_seq1.p1 | GO:0005575,GO:0016020 | GRAM domain containing 1B |
| comp219179_c0_seq4.p1 | GO:0003674,GO:0005215,GO:0005575,GO:0006810,GO:0006811,GO:0006812,GO:0006813,GO:0006814,GO:0006816,GO:0008150,GO:0008273,GO:0008324,GO:0009987,GO:0015075,GO:0015077,GO:0015079,GO:0015081,GO:0015085,GO:0015291,GO:0015297,GO:0015298,GO:0015368,GO:0015491,GO:0015672,GO:0016020,GO:0016021,GO:0022804,GO:0022821,GO:0022857,GO:0022890,GO:0022891,GO:0022892,GO:0030001,GO:0031224,GO:0034220,GO:0035725,GO:0044425,GO:0044699,GO:0044763,GO:0044765,GO:0046873,GO:0051179,GO:0051234,GO:0055085,GO:0070588,GO:0070838,GO:0071804,GO:0071805,GO:0072509,GO:0072511 | Sodium potassium calcium exchanger |

| | | |
|---|---|---|
| comp220405_c1_seq2.p1 | GO:0002165,GO:0003002,GO:0003674,GO:0003676,GO:0003677,GO:0005488,GO:0005575,GO:0005622,GO:0005623,GO:0005634,GO:0006355,GO:0007154,GO:0007165,GO:0007166,GO:0007275,GO:0007379,GO:0007389,GO:0007399,GO:0007444,GO:0007447,GO:0007450,GO:0007472,GO:0007476,GO:0007478,GO:0007480,GO:0007552,GO:0007560,GO:0008150,GO:0008586,GO:0008587,GO:0009653,GO:0009791,GO:0009886,GO:0009887,GO:0009889,GO:0009891,GO:0009893,GO:0009953,GO:0009966,GO:0009987,GO:0010468,GO:0010556,GO:0010557,GO:0010604,GO:0010628,GO:0010646,GO:0016055,GO:0019219,GO:0019222,GO:0022008,GO:0023051,GO:0023052,GO:0030154,GO:0030182,GO:0031323,GO:0031325,GO:0031326,GO:0031328,GO:0032501,GO:0032502,GO:0032991,GO:0035060,GO:0035107,GO:0035108,GO:0035114,GO:0035120,GO:0035127,GO:0035218,GO:0035220,GO:0035222,GO:0035282,GO:0042058,GO:0043226,GO:0043227,GO:0043229,GO:0043231,GO:0043234,GO:0044422,GO:0044424,GO:0044428,GO:0044446,GO:0044464,GO:0044699,GO:0044700,GO:0044707,GO:0044763,GO:0044767,GO:0045893,GO:0045935,GO:0046530,GO:0048190,GO:0048513,GO:0048518,GO:0048522,GO:0048563,GO:0048569,GO:0048583,GO:0048699,GO:0048707,GO:0048731,GO:0048736,GO:0048737,GO:0048856,GO:0048869,GO:0050789,GO:0050794,GO:0050896,GO:0051171,GO:0051173,GO:0051252,GO:0051254,GO:0051716,GO:0060173,GO:0060255,GO:0065007,GO:0070603,GO:0080090,GO:0090544,GO:0097159,GO:1901184,GO:1901363,GO:2000112,GO:2001141 | Trithorax group (trxG) protein required for embryonic segmentation, development of the notum and wing margin, and photoreceptor differentiation. Required for the activation of genes such as Antp, Ubx and Eve. Binds to DNA without specific affinity, suggesting that it is recruited to promoters by promoter-specific proteins. Essential component of the Brahma complex, a multiprotein complex which is the equivalent of the yeast SWI SNF complex and acts by remodeling the chromatin by catalyzing an ATP-dependent alteration in the structure of nucleosomal DNA. This complex can both serve as a transcriptional coactivator or corepressor, depending on the context. Acts as an essential coactivator for Zeste, which recruits the whole complex to specific genes. In contrast, it acts as a corepressor for Wg target genes, possibly via an interaction with Pan and Gro. It also acts as a negative regulator for proneural achaete-scute, when it is directly recruited by Pan and Chi. Also represses E2f activation |

| | | |
|---|---|---|
| comp212167_c0_seq8.p1 | GO:0000123,GO:0000975,GO:0001067,GO:0001071,GO:0003674,GO:0003676,GO:0003677,GO:0003700,GO:0005488,GO:0005515,GO:0005575,GO:0005622,GO:0005623,GO:0005634,GO:0005654,GO:0005671,GO:0006325,GO:0006338,GO:0006355,GO:0006357,GO:0006464,GO:0006473,GO:0006475,GO:0006996,GO:0008150,GO:0008152,GO:0009889,GO:0009891,GO:0009893,GO:0009987,GO:0010468,GO:0010556,GO:0010557,GO:0010604,GO:0010628,GO:0016043,GO:0016568,GO:0016569,GO:0016570,GO:0016573,GO:0018193,GO:0018205,GO:0018393,GO:0018394,GO:0019219,GO:0019222,GO:0019538,GO:0031323,GO:0031325,GO:0031326,GO:0031328,GO:0031974,GO:0031981,GO:0032991,GO:0036211,GO:0042802,GO:0042803,GO:0043170,GO:0043226,GO:0043227,GO:0043229,GO:0043231,GO:0043233,GO:0043234,GO:0043412,GO:0043543,GO:0044212,GO:0044237,GO:0044238,GO:0044260,GO:0044267,GO:0044422,GO:0044424,GO:0044428,GO:0044446,GO:0044451,GO:0044464,GO:0044699,GO:0044763,GO:0045893,GO:0045935,GO:0045944,GO:0046982,GO:0046983,GO:0048518,GO:0048522,GO:0050789,GO:0050794,GO:0051171,GO:0051173,GO:0051252,GO:0051254,GO:0051276,GO:0060255,GO:0065007,GO:0070013,GO:0071704,GO:0071840,GO:0080090,GO:0097159,GO:1901363,GO:2000112,GO:2001141 | GA binding protein transcription factor beta subunit |
| comp214874_c0_seq2.p1 | GO:0000166,GO:0001882,GO:0001883,GO:0003674,GO:0003824,GO:0003924,GO:0005488,GO:0005525,GO:0006139,GO:0006152,GO:0006163,GO:0006184,GO:0006195,GO:0006725,GO:0006753,GO:0006793,GO:0006796,GO:0006807,GO:0007154,GO:0007165,GO:0007264,GO:0008150,GO:0008152,GO:0009056,GO:0009116,GO:0009117,GO:0009119,GO:0009141,GO:0009143,GO:0009144,GO:0009146,GO:0009150,GO:0009154,GO:0009164,GO:0009166,GO:0009199,GO:0009203,GO:0009205,GO:0009207,GO:0009259,GO:0009261,GO:0009987,GO:0016462,GO:0016787,GO:0016817,GO:0016818,GO:0017076,GO:0017111,GO:0019001,GO:0019439,GO:0019637,GO:0019693,GO:0023052,GO:0032549,GO:0032550,GO:0032553,GO:0032555,GO:0032561,GO:0034641,GO:0034655,GO:0035556,GO:0035639,GO:0036094,GO:0042278,GO:0042454,GO:0043167,GO:0043168,GO:0044237,GO:0044238,GO:0044248,GO:0044270,GO:0044281,GO:0044699,GO:0044700,GO:0044710,GO:0044763,GO:0046039,GO:0046128,GO:0046130,GO:0046434,GO:0046483,GO:0046700,GO:0050789,GO:0050794,GO:0050896,GO:0051716,GO:0055086,GO:0065007,GO:0071704,GO:0072521,GO:0072523,GO:0097159,GO:1901068,GO:1901069,GO:1901135,GO:1901136,GO:1901265,GO:1901292,GO:1901360,GO:1901361,GO:1901363,GO:1901564,GO:1901565,GO:1901575,GO:1901657,GO:1901658 | GTP binding protein |
| comp208824_c0_seq2.p1 | | Inherit from opiNOG: Nucleotide-diphospho-sugar transferase |
| comp214073_c0_seq1.p1 | | Serum amyloid A-like 1 |

| | | |
|---|---|---|
| comp219916_c0_seq6.p1 | GO:0000122,GO:0000165,GO:0001775,GO:0001932,GO:0001934,GO:0002327,GO:0002376,GO:0002520,GO:0002521,GO:0002682,GO:0002902,GO:0002904,GO:0003674,GO:0004871,GO:0005083,GO:0005085,GO:0005088,GO:0005096,GO:0005099,GO:0005488,GO:0005509,GO:0005515,GO:0005575,GO:0005576,GO:0005578,GO:0005622,GO:0005623,GO:0005737,GO:0005886,GO:0005887,GO:0006140,GO:0006355,GO:0006357,GO:0006915,GO:0006950,GO:0007154,GO:0007165,GO:0007243,GO:0007264,GO:0007265,GO:0007275,GO:0008047,GO:0008150,GO:0008219,GO:0008289,GO:0008544,GO:0009118,GO:0009267,GO:0009605,GO:0009888,GO:0009889,GO:0009890,GO:0009892,GO:0009893,GO:0009894,GO:0009966,GO:0009968,GO:0009987,GO:0009991,GO:0010466,GO:0010468,GO:0010556,GO:0010558,GO:0010562,GO:0010604,GO:0010605,GO:0010629,GO:0010646,GO:0010648,GO:0010941,GO:0010942,GO:0010951,GO:0012501,GO:0016020,GO:0016021,GO:0016265,GO:0017016,GO:0017034,GO:0018996,GO:0019219,GO:0019220,GO:0019222,GO:0019899,GO:0019932,GO:0019933,GO:0019935,GO:0019992,GO:0023051,GO:0023052,GO:0023057,GO:0030097,GO:0030098,GO:0030154,GO:0030183,GO:0030234,GO:0030695,GO:0030742,GO:0030811,GO:0031012,GO:0031224,GO:0031226,GO:0031267,GO:0031323,GO:0031324,GO:0031325,GO:0031326,GO:0031327,GO:0031329,GO:0031399,GO:0031401,GO:0031644,GO:0031667,GO:0031668,GO:0031669,GO:0031929,GO:0032006,GO:0032007,GO:0032268,GO:0032270,GO:0032317,GO:0032318,GO:0032320,GO:0032487,GO:0032501,GO:0032502,GO:0032854,GO:0033121,GO:0033124,GO:0033554,GO:0035556,GO:0040002,GO:0042113,GO:0042303,GO:0042325,GO:0042327,GO:0042335,GO:0042594,GO:0042981,GO:0043065,GO:0043067,GO:0043068,GO:0043085,GO:0043086,GO:0043087,GO:0043154,GO:0043167,GO:0043169,GO:0043281,GO:0043547,GO:0043588,GO:0044057,GO:0044092,GO:0044093,GO:0044421,GO:0044424,GO:0044425,GO:0044459,GO:0044464,GO:0044699,GO:0044700,GO:0044707,GO:0044763,GO:0044767,GO:0045321,GO:0045595,GO:0045892,GO:0045934,GO:0045937,GO:0046578,GO:0046582,GO:0046649,GO:0046872,GO:0048167,GO:0048513,GO:0048518,GO:0048519,GO:0048522,GO:0048523,GO:0048534,GO:0048583,GO:0048585,GO:0048731,GO:0048856,GO:0048869,GO:0050789,GO:0050790,GO:0050793,GO:0050794,GO:0050804,GO:0050896,GO:0051020,GO:0051056,GO:0051171,GO:0051172,GO:0051174,GO:0051239,GO:0051246,GO:0051247,GO:0051252,GO:0051253,GO:0051336,GO:0051345,GO:0051346,GO:0051716,GO:0051969,GO:0052547,GO:0052548,GO:0060089,GO:0060102,GO:0060255,GO:0060589,GO:0065007,GO:0065008,GO:0065009,GO:0070228,GO:0070230,GO:0071496,GO:0071944,GO:0080090,GO:1900542,GO:1901532,GO:2000026,GO:2000106,GO:2000108,GO:2000112,GO:2000113,GO:2000116,GO:2000117,GO:2000973,GO:2001141 | Rap guanine nucleotide exchange factor (GEF) |
| comp216462_c0_seq3.p1 | GO:0003674,GO:0003774,GO:0003777,GO:0003824,GO:0005575,GO:0005622,GO:0005623,GO:0005737,GO:0005856,GO:0005858,GO:0005875,GO:0005929,GO:0005930,GO:0006139,GO:0006152,GO:0006163,GO:0006195,GO:0006200,GO:0006725,GO:0006753,GO:0006793,GO:0006796,GO:0006807,GO:0006928,GO:0007017,GO:0007018,GO:0008150,GO:0008152,GO:0009056,GO:0009116,GO:0009117,GO:0009119,GO:0009123,GO:0009125,GO:0009126,GO:0009128,GO:0009141,GO:0009143,GO:0009144,GO:0009146,GO:0009150,GO:0009154,GO:0009158,GO:0009161,GO:0009164,GO:0009166,GO:0009167,GO:0009169,GO:0009199,GO:0009203,GO:0009205,GO:0009207,GO:0009259,GO:0009261,GO:0009987,GO:0015630,GO:0016462,GO:0016787,GO:0016817,GO:0016818,GO:0016887,GO:0017111,GO:0019439,GO:0019637,GO:0019693,GO:0030286,GO:0031514,GO:0032838,GO:0032991,GO:0034641,GO:0034655,GO:0035085,GO:0042278,GO:0042454,GO:0042623,GO:0042995,GO:0043226,GO:0043227,GO:0043228,GO:0043229,GO:0043231,GO:0043232,GO:0043234,GO:0044237,GO:0044238,GO:0044248,GO:0044270,GO:0044281,GO:0044422,GO:0044424,GO:0044430,GO:0044441,GO:0044444,GO:0044446,GO:0044447,GO:0044463,GO:0044464,GO:0044699,GO:0044710,GO:0044763,GO:0046034,GO:0046128,GO:0046130,GO:0046434,GO:0046483,GO:0046700,GO:0055086,GO:0071704,GO:0072521,GO:0072523,GO:0097014,GO:1901135,GO:1901136,GO:1901292,GO:1901360,GO:1901361,GO:1901564,GO:1901565,GO:1901575,GO:1901657,GO:1901658 | Dynein, axonemal, heavy chain |

| | | |
|---|---|---|
| comp217672_c1_seq3.p1 | GO:0005575,GO:0005622,GO:0005623,GO:0005737,GO:0005739,GO:0005783,GO:0005788,GO:0006605,GO:0006612,GO:0006810,GO:0006886,GO:0007009,GO:0008104,GO:0008150,GO:0009987,GO:0015031,GO:0016043,GO:0016044,GO:0031974,GO:0033036,GO:0034613,GO:0043226,GO:0043227,GO:0043229,GO:0043231,GO:0043233,GO:0044422,GO:0044424,GO:0044432,GO:0044444,GO:0044446,GO:0044464,GO:0044699,GO:0044763,GO:0044765,GO:0045184,GO:0046907,GO:0051179,GO:0051234,GO:0051641,GO:0051649,GO:0061024,GO:0070013,GO:0070727,GO:0071702,GO:0071840,GO:0072657,GO:0072659,GO:0072661,GO:0090002,GO:0090150 | phosphofurin acidic cluster sorting protein |
| comp217678_c0_seq3.p1 | GO:0000768,GO:0000902,GO:0000904,GO:0003674,GO:0004872,GO:0005488,GO:0005515,GO:0005575,GO:0005623,GO:0005886,GO:0005887,GO:0006355,GO:0006928,GO:0006935,GO:0006949,GO:0007155,GO:0007275,GO:0007399,GO:0007409,GO:0007411,GO:0007517,GO:0007519,GO:0007520,GO:0008150,GO:0009605,GO:0009653,GO:0009888,GO:0009889,GO:0009987,GO:0010468,GO:0010556,GO:0014706,GO:0014902,GO:0016020,GO:0016021,GO:0016043,GO:0019219,GO:0019222,GO:0022008,GO:0022610,GO:0030030,GO:0030154,GO:0030182,GO:0031175,GO:0031224,GO:0031226,GO:0031323,GO:0031326,GO:0032501,GO:0032502,GO:0032989,GO:0032990,GO:0035914,GO:0040011,GO:0042221,GO:0042330,GO:0042592,GO:0042692,GO:0044425,GO:0044459,GO:0044464,GO:0044699,GO:0044707,GO:0044763,GO:0044767,GO:0045296,GO:0045595,GO:0045597,GO:0048468,GO:0048513,GO:0048518,GO:0048522,GO:0048646,GO:0048666,GO:0048667,GO:0048699,GO:0048731,GO:0048812,GO:0048856,GO:0048858,GO:0048869,GO:0048878,GO:0050789,GO:0050793,GO:0050794,GO:0050801,GO:0050839,GO:0050896,GO:0051094,GO:0051146,GO:0051147,GO:0051149,GO:0051171,GO:0051252,GO:0055065,GO:0055072,GO:0055076,GO:0055080,GO:0060255,GO:0060537,GO:0060538,GO:0061061,GO:0061564,GO:0065007,GO:0065008,GO:0071840,GO:0071944,GO:0080090,GO:0097485,GO:2000112,GO:2001141 | neogenin 1 |
| comp217678_c0_seq3.p2 | GO:0000902,GO:0000904,GO:0003002,GO:0003674,GO:0004871,GO:0004872,GO:0004888,GO:0005042,GO:0005575,GO:0005623,GO:0005886,GO:0006928,GO:0006935,GO:0007154,GO:0007165,GO:0007166,GO:0007275,GO:0007389,GO:0007399,GO:0007409,GO:0007411,GO:0007431,GO:0007432,GO:0008037,GO:0008038,GO:0008045,GO:0008150,GO:0008347,GO:0009605,GO:0009653,GO:0009887,GO:0009987,GO:0010160,GO:0010975,GO:0016020,GO:0016043,GO:0016198,GO:0016199,GO:0016358,GO:0016477,GO:0022008,GO:0023052,GO:0030030,GO:0030154,GO:0030182,GO:0030424,GO:0030425,GO:0031175,GO:0031344,GO:0032501,GO:0032502,GO:0032989,GO:0032990,GO:0035272,GO:0038007,GO:0038023,GO:0040011,GO:0042063,GO:0042221,GO:0042330,GO:0042995,GO:0043005,GO:0043025,GO:0044297,GO:0044464,GO:0044699,GO:0044700,GO:0044707,GO:0044763,GO:0044767,GO:0045595,GO:0045664,GO:0048468,GO:0048513,GO:0048645,GO:0048646,GO:0048666,GO:0048667,GO:0048699,GO:0048731,GO:0048732,GO:0048812,GO:0048813,GO:0048856,GO:0048858,GO:0048859,GO:0048869,GO:0048870,GO:0050767,GO:0050773,GO:0050789,GO:0050793,GO:0050794,GO:0050896,GO:0051128,GO:0051179,GO:0051239,GO:0051674,GO:0051716,GO:0051960,GO:0060089,GO:0060284,GO:0061564,GO:0065007,GO:0070983,GO:0071840,GO:0071944,GO:0097458,GO:0097485,GO:2000026 | Neogenin C-terminus |

| | | |
|---|---|---|
| comp217229_c4_seq38.p1 | GO:0001738,GO:0002009,GO:0003674,GO:0003824,GO:0004385,GO:0005488,GO:0005515,GO:0005575,GO:0005622,GO:0005623,GO:0005737,GO:0005886,GO:0005911,GO:0005912,GO:0005913,GO:0005918,GO:0005938,GO:0006139,GO:0006725,GO:0006753,GO:0006793,GO:0006796,GO:0006807,GO:0007043,GO:0007154,GO:0007163,GO:0007267,GO:0007275,GO:0008150,GO:0008152,GO:0009117,GO:0009653,GO:0009790,GO:0009888,GO:0009987,GO:0016020,GO:0016043,GO:0016301,GO:0016310,GO:0016324,GO:0016327,GO:0016331,GO:0016332,GO:0016740,GO:0016772,GO:0016776,GO:0019201,GO:0019205,GO:0019637,GO:0022607,GO:0023052,GO:0030054,GO:0030154,GO:0032501,GO:0032502,GO:0032991,GO:0034329,GO:0034330,GO:0034332,GO:0034333,GO:0034641,GO:0035003,GO:0035088,GO:0040003,GO:0042335,GO:0043296,GO:0043297,GO:0044085,GO:0044237,GO:0044238,GO:0044281,GO:0044424,GO:0044425,GO:0044444,GO:0044448,GO:0044459,GO:0044464,GO:0044699,GO:0044700,GO:0044707,GO:0044710,GO:0044763,GO:0044767,GO:0045165,GO:0045168,GO:0045177,GO:0045179,GO:0045186,GO:0045197,GO:0045216,GO:0046331,GO:0046483,GO:0046939,GO:0048598,GO:0048729,GO:0048856,GO:0048869,GO:0055086,GO:0060429,GO:0061245,GO:0070160,GO:0070161,GO:0071704,GO:0071840,GO:0071944,GO:1901360 | GuKc |
| comp191115_c0_seq2.p1 | | ankyrin repeat domain 5 |
| comp221163_c0_seq8.p1 | GO:0000226,GO:0001539,GO:0003008,GO:0005575,GO:0005622,GO:0005623,GO:0005737,GO:0005813,GO:0005814,GO:0005815,GO:0005856,GO:0006928,GO:0006996,GO:0007010,GO:0007017,GO:0007020,GO:0007275,GO:0007600,GO:0007605,GO:0008150,GO:0009653,GO:0009790,GO:0009987,GO:0015630,GO:0016043,GO:0031023,GO:0032501,GO:0032502,GO:0040011,GO:0043226,GO:0043228,GO:0043229,GO:0043232,GO:0044422,GO:0044424,GO:0044430,GO:0044444,GO:0044446,GO:0044450,GO:0044464,GO:0044699,GO:0044707,GO:0044763,GO:0044767,GO:0048598,GO:0048856,GO:0048870,GO:0050877,GO:0050954,GO:0051179,GO:0051297,GO:0051674,GO:0071840 | Pericentrin |
| comp208554_c0_seq1.p3 | GO:0005575,GO:0005622,GO:0005623,GO:0005737,GO:0005783,GO:0005789,GO:0012505,GO:0016020,GO:0016021,GO:0030176,GO:0031090,GO:0031224,GO:0031227,GO:0031300,GO:0031301,GO:0032991,GO:0042175,GO:0043226,GO:0043227,GO:0043229,GO:0043231,GO:0043234,GO:0044422,GO:0044424,GO:0044425,GO:0044432,GO:0044444,GO:0044446,GO:0044464,GO:0072546 | ER membrane protein complex subunit 6 |
| comp218546_c2_seq11.p1 | GO:0007275,GO:0008150,GO:0009653,GO:0009987,GO:0016043,GO:0030198,GO:0032501,GO:0032502,GO:0033333,GO:0033334,GO:0035107,GO:0043062,GO:0044699,GO:0044707,GO:0044763,GO:0044767,GO:0048736,GO:0048856,GO:0071711,GO:0071840 | G2F domain |
| comp218485_c1_seq1.p1 | GO:0001654,GO:0002165,GO:0003674,GO:0005488,GO:0005543,GO:0005575,GO:0005622,GO:0005623,GO:0005737,GO:0005886,GO:0006810,GO:0006886,GO:0007275,GO:0007423,GO:0007444,GO:0007472,GO:0007476,GO:0007552,GO:0007560,GO:0008104,GO:0008150,GO:0008289,GO:0008356,GO:0008586,GO:0008593,GO:0009653,GO:0009791,GO:0009886,GO:0009887,GO:0009966,GO:0009968,GO:0009987,GO:0010646,GO:0010648,GO:0015031,GO:0016020,GO:0016197,GO:0016324,GO:0023051,GO:0023057,GO:0030100,GO:0032501,GO:0032502,GO:0032879,GO:0033036,GO:0034613,GO:0035107,GO:0035114,GO:0035120,GO:0035220,GO:0043167,GO:0043168,GO:0044424,GO:0044425,GO:0044459,GO:0044464,GO:0044699,GO:0044707,GO:0044763,GO:0044765,GO:0044767,GO:0045035,GO:0045177,GO:0045184,GO:0045746,GO:0046907,GO:0048513,GO:0048519,GO:0048523,GO:0048563,GO:0048569,GO:0048583,GO:0048585,GO:0048707,GO:0048731,GO:0048736,GO:0048737,GO:0048749,GO:0048856,GO:0050789,GO:0050794,GO:0051049,GO:0051128,GO:0051179,GO:0051234,GO:0051301,GO:0051641,GO:0051649,GO:0060627,GO:0065007,GO:0070727,GO:0071702,GO:0071944 | DM14 |

| | | |
|---|---|---|
| comp210039_c0_seq2.p1 | GO:0000003,GO:0000226,GO:0000278,GO:0001654,GO:0001745,GO:0001763,GO:0002009,GO:0003006,GO:0003674,GO:0005198,GO:0005488,GO:0005515,GO:0005575,GO:0005622,GO:0005623,GO:0005634,GO:0005635,GO:0005638,GO:0005652,GO:0005737,GO:0005811,GO:0005813,GO:0005815,GO:0005819,GO:0005856,GO:0005875,GO:0005882,GO:0006325,GO:0006342,GO:0006355,GO:0006996,GO:0006997,GO:0006998,GO:0007010,GO:0007017,GO:0007049,GO:0007084,GO:0007097,GO:0007154,GO:0007267,GO:0007275,GO:0007399,GO:0007417,GO:0007423,GO:0007424,GO:0007430,GO:0007517,GO:0007548,GO:0007568,GO:0007569,GO:0007610,GO:0007626,GO:0008064,GO:0008104,GO:0008150,GO:0008344,GO:0008406,GO:0009653,GO:0009887,GO:0009888,GO:0009889,GO:0009890,GO:0009892,GO:0009987,GO:0010468,GO:0010556,GO:0010558,GO:0010605,GO:0010629,GO:0012505,GO:0015630,GO:0016043,GO:0016044,GO:0016458,GO:0019219,GO:0019222,GO:0022402,GO:0022414,GO:0023052,GO:0030154,GO:0030534,GO:0030832,GO:0030833,GO:0031023,GO:0031081,GO:0031323,GO:0031324,GO:0031326,GO:0031327,GO:0031468,GO:0031503,GO:0031967,GO:0031974,GO:0031975,GO:0031981,GO:0032271,GO:0032501,GO:0032502,GO:0032535,GO:0032956,GO:0032970,GO:0032991,GO:0033036,GO:0033043,GO:0034399,GO:0034613,GO:0034629,GO:0035239,GO:0035262,GO:0035295,GO:0035989,GO:0040003,GO:0040023,GO:0040029,GO:0042335,GO:0043226,GO:0043227,GO:0043228,GO:0043229,GO:0043231,GO:0043232,GO:0043233,GO:0043234,GO:0043254,GO:0044087,GO:0044422,GO:0044424,GO:0044428,GO:0044430,GO:0044444,GO:0044446,GO:0044464,GO:0044699,GO:0044700,GO:0044707,GO:0044708,GO:0044763,GO:0044767,GO:0045111,GO:0045137,GO:0045165,GO:0045168,GO:0045814,GO:0045892,GO:0045934,GO:0046331,GO:0048513,GO:0048519,GO:0048523,GO:0048546,GO:0048565,GO:0048592,GO:0048608,GO:0048644,GO:0048729,GO:0048731,GO:0048749,GO:0048754,GO:0048856,GO:0048869,GO:0050789,GO:0050794,GO:0050896,GO:0051128,GO:0051171,GO:0051172,GO:0051179,GO:0051234,GO:0051252,GO:0051253,GO:0051276,GO:0051297,GO:0051493,GO:0051640,GO:0051641,GO:0051647,GO:0051649,GO:0051656,GO:0051664,GO:0051668,GO:0055123,GO:0060255,GO:0060415,GO:0060429,GO:0060446,GO:0060541,GO:0060562,GO:0061024,GO:0061061,GO:0061138,GO:0061448,GO:0061458,GO:0065007,GO:0065008,GO:0070013,GO:0070727,GO:0070732,GO:0071763,GO:0071840,GO:0072686,GO:0080090,GO:0090066,GO:0090435,GO:2000112,GO:2000113,GO:2001141 | Intermediate filament protein |
| comp213497_c0_seq1.p1 | | ankyrin repeat and FYVE |
| comp212897_c1_seq2.p1 | GO:0000003,GO:0000375,GO:0000377,GO:0000398,GO:0002009,GO:0002119,GO:0002164,GO:0003006,GO:0003674,GO:0003676,GO:0003723,GO:0005488,GO:0005575,GO:0005622,GO:0005623,GO:0005634,GO:0005654,GO:0006139,GO:0006396,GO:0006397,GO:0006725,GO:0006807,GO:0007275,GO:0007276,GO:0007281,GO:0008150,GO:0008152,GO:0008380,GO:0009607,GO:0009615,GO:0009653,GO:0009790,GO:0009791,GO:0009792,GO:0009888,GO:0009987,GO:0010467,GO:0010564,GO:0010876,GO:0016070,GO:0016071,GO:0019915,GO:0019953,GO:0022412,GO:0022414,GO:0030154,GO:0031974,GO:0031981,GO:0032501,GO:0032502,GO:0032504,GO:0033036,GO:0034641,GO:0040007,GO:0040008,GO:0040009,GO:0040010,GO:0040011,GO:0040020,GO:0042127,GO:0043170,GO:0043226,GO:0043227,GO:0043229,GO:0043231,GO:0043233,GO:0044237,GO:0044238,GO:0044260,GO:0044422,GO:0044424,GO:0044428,GO:0044446,GO:0044464,GO:0044699,GO:0044702,GO:0044707,GO:0044763,GO:0044767,GO:0045927,GO:0046483,GO:0048468,GO:0048518,GO:0048609,GO:0048610,GO:0048729,GO:0048856,GO:0048869,GO:0050789,GO:0050794,GO:0050896,GO:0051179,GO:0051445,GO:0051704,GO:0051707,GO:0051726,GO:0060429,GO:0065007,GO:0070013,GO:0071704,GO:0090304,GO:0097159,GO:1901360,GO:1901363 | Heterogeneous nuclear ribonucleoprotein U-like 1 |

| | | |
|---|---|---|
| comp208853_c0_seq1.p1 | GO:0000975,GO:0001067,GO:0001071,GO:0003674,GO:0003676,GO:0003677,GO:0003700,GO:0005488,GO:0005575,GO:0005622,GO:0005623,GO:0005634,GO:0006355,GO:0006357,GO:0008150,GO:0009889,GO:0009891,GO:0009893,GO:0010468,GO:0010556,GO:0010557,GO:0010604,GO:0010628,GO:0019219,GO:0019222,GO:0031323,GO:0031325,GO:0031326,GO:0031328,GO:0043226,GO:0043227,GO:0043229,GO:0043231,GO:0044212,GO:0044424,GO:0044464,GO:0045893,GO:0045935,GO:0045944,GO:0048518,GO:0048522,GO:0050789,GO:0050794,GO:0051171,GO:0051173,GO:0051252,GO:0051254,GO:0060255,GO:0065007,GO:0080090,GO:0097159,GO:1901363,GO:2000112,GO:2001141 | transcription factor 20 |
| comp217574_c1_seq1.p1 | GO:0000902,GO:0000904,GO:0003674,GO:0005083,GO:0005096,GO:0005099,GO:0005100,GO:0005575,GO:0005622,GO:0005623,GO:0005634,GO:0005737,GO:0005829,GO:0006140,GO:0006928,GO:0006935,GO:0007154,GO:0007165,GO:0007264,GO:0007275,GO:0007399,GO:0007409,GO:0007411,GO:0008047,GO:0008150,GO:0009118,GO:0009605,GO:0009653,GO:0009894,GO:0009966,GO:0009987,GO:0010646,GO:0016043,GO:0019219,GO:0019220,GO:0019222,GO:0022008,GO:0023051,GO:0023052,GO:0030030,GO:0030154,GO:0030182,GO:0030234,GO:0030695,GO:0030811,GO:0031175,GO:0031323,GO:0031329,GO:0032318,GO:0032319,GO:0032320,GO:0032321,GO:0032501,GO:0032502,GO:0032989,GO:0032990,GO:0033121,GO:0033124,GO:0035023,GO:0035556,GO:0040011,GO:0042221,GO:0042330,GO:0043085,GO:0043087,GO:0043226,GO:0043227,GO:0043229,GO:0043231,GO:0043547,GO:0044093,GO:0044424,GO:0044444,GO:0044464,GO:0044699,GO:0044700,GO:0044707,GO:0044763,GO:0044767,GO:0046578,GO:0048468,GO:0048583,GO:0048666,GO:0048667,GO:0048699,GO:0048731,GO:0048812,GO:0048856,GO:0048858,GO:0048869,GO:0050789,GO:0050790,GO:0050794,GO:0050896,GO:0051056,GO:0051171,GO:0051174,GO:0051336,GO:0051345,GO:0051716,GO:0060589,GO:0061564,GO:0065007,GO:0065009,GO:0071840,GO:0080090,GO:0097485,GO:1900542 | Rho GTPase activating protein 39 |
| comp220864_c4_seq4.p1 | | Inherit from veNOG: kiaa2026 |
| comp209791_c0_seq2.p1 | GO:0003674,GO:0003824,GO:0004793,GO:0008150,GO:0008152,GO:0016829,GO:0016830,GO:0016832 | Beta-eliminating lyase |
| comp221069_c2_seq8.p1 | | transportin 3 |
| comp210725_c0_seq3.p1 | GO:0001678,GO:0003674,GO:0003824,GO:0004340,GO:0004396,GO:0005575,GO:0005622,GO:0005623,GO:0005737,GO:0005829,GO:0005975,GO:0005996,GO:0006006,GO:0006007,GO:0006091,GO:0006096,GO:0006793,GO:0006796,GO:0008150,GO:0008152,GO:0008865,GO:0009056,GO:0009987,GO:0016052,GO:0016301,GO:0016310,GO:0016740,GO:0016772,GO:0016773,GO:0019158,GO:0019200,GO:0019318,GO:0019320,GO:0019637,GO:0019725,GO:0033500,GO:0040011,GO:0042592,GO:0042593,GO:0044237,GO:0044238,GO:0044262,GO:0044424,GO:0044444,GO:0044464,GO:0044699,GO:0044723,GO:0044724,GO:0044763,GO:0046365,GO:0046835,GO:0048878,GO:0051156,GO:0055082,GO:0060361,GO:0065007,GO:0065008,GO:0071704,GO:1901135,GO:1901575 | Hexokinase |

| | | |
|---|---|---|
| comp214276_c1_seq1.p1 | GO:0000165,GO:0000902,GO:0003008,GO:0003674,GO:0005488,GO:0005515,GO:0005575,GO:0005622,GO:0005623,GO:0005737,GO:0006810,GO:0006928,GO:0006950,GO:0007017,GO:0007018,GO:0007154,GO:0007165,GO:0007243,GO:0007254,GO:0007267,GO:0007268,GO:0007275,GO:0007399,GO:0008088,GO:0008092,GO:0008150,GO:0009653,GO:0009966,GO:0009987,GO:0010627,GO:0010646,GO:0010970,GO:0016043,GO:0016192,GO:0019226,GO:0019894,GO:0019896,GO:0019899,GO:0019900,GO:0019901,GO:0022008,GO:0023051,GO:0023052,GO:0030030,GO:0030154,GO:0030182,GO:0030705,GO:0031098,GO:0031175,GO:0032501,GO:0032502,GO:0032872,GO:0032989,GO:0032990,GO:0033554,GO:0035556,GO:0035637,GO:0043408,GO:0044424,GO:0044464,GO:0044699,GO:0044700,GO:0044707,GO:0044763,GO:0044765,GO:0044767,GO:0046328,GO:0046907,GO:0048468,GO:0048489,GO:0048490,GO:0048491,GO:0048583,GO:0048666,GO:0048699,GO:0048731,GO:0048812,GO:0048856,GO:0048858,GO:0048869,GO:0050789,GO:0050794,GO:0050877,GO:0050896,GO:0051179,GO:0051234,GO:0051403,GO:0051640,GO:0051641,GO:0051648,GO:0051649,GO:0051650,GO:0051656,GO:0051716,GO:0065007,GO:0070302,GO:0071840,GO:0080134,GO:0080135,GO:0097479,GO:0097480 | PTB |
| comp148406_c0_seq2.p1 | | tripartite motif containing 3 |
| comp1481530_c0_seq1.p1 | | serine threonine-protein phosphatase 6 regulatory ankyrin repeat subunit |
| comp215571_c0_seq3.p1 | | K(lysine) acetyltransferase 2A |
| comp215608_c0_seq1.p1 | GO:0000003,GO:0001071,GO:0001654,GO:0001667,GO:0001708,GO:0001709,GO:0001742,GO:0001745,GO:0001751,GO:0001754,GO:0002165,GO:0003002,GO:0003006,GO:0003008,GO:0003674,GO:0003700,GO:0005575,GO:0005622,GO:0005623,GO:0005634,GO:0006355,GO:0006928,GO:0007275,GO:0007379,GO:0007389,GO:0007399,GO:0007423,GO:0007424,GO:0007427,GO:0007438,GO:0007444,GO:0007469,GO:0007472,GO:0007476,GO:0007517,GO:0007525,GO:0007548,GO:0007552,GO:0007560,GO:0007600,GO:0007605,GO:0008150,GO:0008406,GO:0008584,GO:0008586,GO:0009653,GO:0009791,GO:0009886,GO:0009887,GO:0009889,GO:0009890,GO:0009892,GO:0009987,GO:0010453,GO:0010468,GO:0010556,GO:0010558,GO:0010605,GO:0010629,GO:0010631,GO:0016477,GO:0019219,GO:0019222,GO:0022008,GO:0022414,GO:0030154,GO:0030182,GO:0030539,GO:0031323,GO:0031324,GO:0031326,GO:0031327,GO:0032501,GO:0032502,GO:0035107,GO:0035114,GO:0035120,GO:0035214,GO:0035220,GO:0035277,GO:0035282,GO:0035309,GO:0035310,GO:0040011,GO:0042659,GO:0043226,GO:0043227,GO:0043229,GO:0043231,GO:0044424,GO:0044464,GO:0044699,GO:0044707,GO:0044763,GO:0044767,GO:0045137,GO:0045165,GO:0045465,GO:0045466,GO:0045595,GO:0045892,GO:0045934,GO:0046530,GO:0046546,GO:0046661,GO:0046845,GO:0048098,GO:0048468,GO:0048513,GO:0048519,GO:0048523,GO:0048563,GO:0048569,GO:0048592,GO:0048608,GO:0048644,GO:0048699,GO:0048707,GO:0048731,GO:0048736,GO:0048737,GO:0048749,GO:0048806,GO:0048856,GO:0048869,GO:0048870,GO:0050789,GO:0050793,GO:0050794,GO:0050877,GO:0050954,GO:0051171,GO:0051172,GO:0051179,GO:0051252,GO:0051253,GO:0051674,GO:0060255,GO:0060541,GO:0060573,GO:0060581,GO:0061061,GO:0061458,GO:0065007,GO:0080090,GO:0090130,GO:0090132,GO:2000112,GO:2000113,GO:2001141 | Zinc finger, C2H2 type |
| comp219875_c0_seq3.p1 | | collagen, type XII, alpha 1 |
| comp220642_c3_seq3.p1 | | NACHT and WD repeat |

| | | |
|---|---|---|
| comp211426_c0_seq2.p1 | GO:0000122,GO:0000775,GO:0000785,GO:0000792,GO:0000975,GO:0001067,GO:0001071,GO:0001654,GO:0001775,GO:0001779,GO:0002376,GO:0002377,GO:0002440,GO:0002520,GO:0002521,GO:0002682,GO:0002684,GO:0002694,GO:0002696,GO:0002761,GO:0002763,GO:0003674,GO:0003676,GO:0003677,GO:0003700,GO:0005488,GO:0005515,GO:0005575,GO:0005622,GO:0005623,GO:0005634,GO:0005654,GO:0005667,GO:0005694,GO:0005721,GO:0005737,GO:0005886,GO:0006355,GO:0006357,GO:0006464,GO:0006468,GO:0006793,GO:0006796,GO:0006873,GO:0006874,GO:0006875,GO:0007204,GO:0007275,GO:0007399,GO:0007417,GO:0007420,GO:0007423,GO:0007498,GO:0008150,GO:0008152,GO:0009888,GO:0009889,GO:0009890,GO:0009891,GO:0009892,GO:0009893,GO:0009987,GO:0010468,GO:0010556,GO:0010557,GO:0010558,GO:0010604,GO:0010605,GO:0010628,GO:0010629,GO:0010941,GO:0016020,GO:0016049,GO:0016310,GO:0019219,GO:0019222,GO:0019538,GO:0019725,GO:0030003,GO:0030097,GO:0030098,GO:0030101,GO:0030154,GO:0030183,GO:0030217,GO:0030852,GO:0030854,GO:0030888,GO:0030900,GO:0031323,GO:0031324,GO:0031325,GO:0031326,GO:0031327,GO:0031328,GO:0031974,GO:0031981,GO:0032501,GO:0032502,GO:0032944,GO:0032991,GO:0032993,GO:0033077,GO:0036211,GO:0040007,GO:0040008,GO:0040014,GO:0040018,GO:0042110,GO:0042113,GO:0042127,GO:0042592,GO:0042802,GO:0042803,GO:0042981,GO:0043010,GO:0043067,GO:0043170,GO:0043226,GO:0043227,GO:0043228,GO:0043229,GO:0043231,GO:0043232,GO:0043233,GO:0043234,GO:0043412,GO:0043565,GO:0044212,GO:0044237,GO:0044238,GO:0044260,GO:0044267,GO:0044422,GO:0044424,GO:0044427,GO:0044428,GO:0044446,GO:0044451,GO:0044464,GO:0044699,GO:0044707,GO:0044763,GO:0044767,GO:0045321,GO:0045577,GO:0045580,GO:0045582,GO:0045595,GO:0045597,GO:0045619,GO:0045621,GO:0045637,GO:0045639,GO:0045658,GO:0045660,GO:0045892,GO:0045893,GO:0045927,GO:0045934,GO:0045935,GO:0045944,GO:0046634,GO:0046635,GO:0046637,GO:0046638,GO:0046649,GO:0046982,GO:0046983,GO:0048513,GO:0048518,GO:0048519,GO:0048522,GO:0048523,GO:0048534,GO:0048535,GO:0048537,GO:0048538,GO:0048541,GO:0048731,GO:0048732,GO:0048856,GO:0048869,GO:0048878,GO:0050670,GO:0050789,GO:0050793,GO:0050794,GO:0050801,GO:0050863,GO:0050864,GO:0050865,GO:0050867,GO:0050870,GO:0051094,GO:0051136,GO:0051138,GO:0051171,GO:0051172,GO:0051173,GO:0051239,GO:0051240,GO:0051249,GO:0051251,GO:0051252,GO:0051253,GO:0051254,GO:0051480,GO:0051481,GO:0055065,GO:0055074,GO:0055080,GO:0055082,GO:0060041,GO:0060255,GO:0065007,GO:0065008,GO:0070013,GO:0070663,GO:0071704,GO:0071944,GO:0072503,GO:0072507,GO:0080090,GO:0097159,GO:1901363,GO:1902105,GO:1902107,GO:2000026,GO:2000112,GO:2000113,GO:2001141 | IKAROS family zinc finger |
| comp216575_c3_seq1.p1 | | SANT |
| comp214826_c0_seq5.p1 | | cyclin J |

| | | |
|---|---|---|
| comp221132_c0_seq4.p1 | GO:0000122,GO:0000228,GO:0000785,GO:0000790,GO:0000975,GO:0000988,GO:0000989,GO:0001067,GO:0001076,GO:0001104,GO:0001105,GO:0001190,GO:0001558,GO:0001944,GO:0003674,GO:0003676,GO:0003677,GO:0003712,GO:0003713,GO:0003824,GO:0004386,GO:0005488,GO:0005575,GO:0005622,GO:0005623,GO:0005634,GO:0005654,GO:0005694,GO:0005856,GO:0006139,GO:0006152,GO:0006163,GO:0006195,GO:0006200,GO:0006259,GO:0006323,GO:0006325,GO:0006333,GO:0006334,GO:0006338,GO:0006351,GO:0006355,GO:0006357,GO:0006366,GO:0006725,GO:0006753,GO:0006793,GO:0006796,GO:0006807,GO:0006996,GO:0007275,GO:0008094,GO:0008150,GO:0008152,GO:0008285,GO:0009056,GO:0009058,GO:0009059,GO:0009116,GO:0009117,GO:0009119,GO:0009123,GO:0009125,GO:0009126,GO:0009128,GO:0009141,GO:0009143,GO:0009144,GO:0009146,GO:0009150,GO:0009154,GO:0009158,GO:0009161,GO:0009164,GO:0009166,GO:0009167,GO:0009169,GO:0009199,GO:0009203,GO:0009205,GO:0009207,GO:0009259,GO:0009261,GO:0009889,GO:0009890,GO:0009891,GO:0009892,GO:0009893,GO:0009987,GO:0010467,GO:0010468,GO:0010556,GO:0010557,GO:0010558,GO:0010604,GO:0010605,GO:0010628,GO:0010629,GO:0016043,GO:0016070,GO:0016462,GO:0016514,GO:0016568,GO:0016787,GO:0016817,GO:0016818,GO:0016887,GO:0017111,GO:0018130,GO:0019219,GO:0019222,GO:0019438,GO:0019439,GO:0019637,GO:0019693,GO:0022607,GO:0030154,GO:0030308,GO:0031323,GO:0031324,GO:0031325,GO:0031326,GO:0031327,GO:0031328,GO:0031497,GO:0031974,GO:0031981,GO:0032501,GO:0032502,GO:0032774,GO:0032991,GO:0034622,GO:0034641,GO:0034645,GO:0034654,GO:0034655,GO:0034728,GO:0035886,GO:0035887,GO:0040008,GO:0042127,GO:0042278,GO:0042454,GO:0042623,GO:0042692,GO:0043170,GO:0043226,GO:0043227,GO:0043228,GO:0043229,GO:0043231,GO:0043232,GO:0043233,GO:0043234,GO:0043933,GO:0044085,GO:0044212,GO:0044237,GO:0044238,GO:0044248,GO:0044249,GO:0044260,GO:0044270,GO:0044271,GO:0044281,GO:0044422,GO:0044424,GO:0044427,GO:0044428,GO:0044446,GO:0044454,GO:0044464,GO:0044699,GO:0044707,GO:0044710,GO:0044763,GO:0044767,GO:0045111,GO:0045892,GO:0045893,GO:0045926,GO:0045934,GO:0045935,GO:0045944,GO:0046034,GO:0046128,GO:0046130,GO:0046434,GO:0046483,GO:0046700,GO:0048518,GO:0048519,GO:0048522,GO:0048523,GO:0048731,GO:0048856,GO:0048869,GO:0050789,GO:0050794,GO:0051128,GO:0051145,GO:0051171,GO:0051172,GO:0051173,GO:0051252,GO:0051253,GO:0051254,GO:0051276,GO:0055086,GO:0060255,GO:0061061,GO:0065003,GO:0065004,GO:0065007,GO:0070013,GO:0070603,GO:0071103,GO:0071564,GO:0071565,GO:0071704,GO:0071778,GO:0071824,GO:0071840,GO:0072358,GO:0072359,GO:0072521,GO:0072523,GO:0080090,GO:0090304,GO:0090544,GO:0097159,GO:0097458,GO:1901135,GO:1901136,GO:1901292,GO:1901360,GO:1901361,GO:1901362,GO:1901363,GO:1901564,GO:1901565,GO:1901575,GO:1901576,GO:1901657,GO:1901658,GO:2000112,GO:2000113,GO:2001141 | SWI SNF related, matrix associated, actin dependent regulator of chromatin, subfamily a, member |
| comp216179_c0_seq4.p1 | | ZU5 |
| comp216845_c0_seq1.p1 | | |
| comp218025_c0_seq3.p1 | | Rho GTPase activating protein |
| comp217202_c0_seq20.p1 | | signal peptide, CUB domain, EGF-like 1 |
| comp217202_c0_seq20.p2 | | DEAD (Asp-Glu-Ala-Asp) box polypeptide 55 |
| comp216383_c1_seq12.p1 | | breakpoint cluster region |

| | | |
|---|---|---|
| comp218356_c0_seq22.p1 | GO:0000988,GO:0000989,GO:0003674,GO:0003712,GO:0003713,GO:0005575,GO:0005622,GO:0005623,GO:0005634,GO:0005654,GO:0006355,GO:0006357,GO:0006629,GO:0008150,GO:0008152,GO:0009889,GO:0009891,GO:0009893,GO:0009987,GO:0010468,GO:0010556,GO:0010557,GO:0010604,GO:0010628,GO:0019219,GO:0019222,GO:0030374,GO:0031323,GO:0031325,GO:0031326,GO:0031328,GO:0031974,GO:0031981,GO:0043226,GO:0043227,GO:0043229,GO:0043231,GO:0043233,GO:0044237,GO:0044238,GO:0044255,GO:0044281,GO:0044422,GO:0044424,GO:0044428,GO:0044446,GO:0044464,GO:0044710,GO:0045893,GO:0045935,GO:0045944,GO:0048518,GO:0048522,GO:0050789,GO:0050794,GO:0051171,GO:0051173,GO:0051252,GO:0051254,GO:0060255,GO:0065007,GO:0070013,GO:0071704,GO:0080090,GO:2000112,GO:2001141 | helicase with zinc finger 2, transcriptional coactivator |
| comp198501_c1_seq1.p1 | | guanylate cyclase |
| comp219249_c0_seq5.p3 | GO:0000228,GO:0000381,GO:0000785,GO:0000790,GO:0000791,GO:0000975,GO:0001047,GO:0001067,GO:0003674,GO:0003676,GO:0003677,GO:0003682,GO:0003723,GO:0003729,GO:0005488,GO:0005575,GO:0005576,GO:0005622,GO:0005623,GO:0005634,GO:0005681,GO:0005694,GO:0005719,GO:0006139,GO:0006351,GO:0006355,GO:0006357,GO:0006366,GO:0006461,GO:0006508,GO:0006509,GO:0006725,GO:0006807,GO:0008150,GO:0008152,GO:0009058,GO:0009059,GO:0009889,GO:0009891,GO:0009892,GO:0009893,GO:0009987,GO:0010033,GO:0010467,GO:0010468,GO:0010556,GO:0010557,GO:0010604,GO:0010605,GO:0010628,GO:0016043,GO:0016070,GO:0018130,GO:0019219,GO:0019222,GO:0019438,GO:0019538,GO:0022607,GO:0030529,GO:0031323,GO:0031324,GO:0031325,GO:0031326,GO:0031328,GO:0031974,GO:0031981,GO:0031982,GO:0031988,GO:0032774,GO:0032991,GO:0033119,GO:0033120,GO:0033619,GO:0034097,GO:0034641,GO:0034645,GO:0034654,GO:0042221,GO:0043170,GO:0043226,GO:0043227,GO:0043228,GO:0043229,GO:0043230,GO:0043231,GO:0043232,GO:0043233,GO:0043484,GO:0043933,GO:0044085,GO:0044212,GO:0044237,GO:0044238,GO:0044249,GO:0044260,GO:0044271,GO:0044421,GO:0044422,GO:0044424,GO:0044427,GO:0044428,GO:0044446,GO:0044454,GO:0044464,GO:0044530,GO:0044699,GO:0044763,GO:0045893,GO:0045934,GO:0045935,GO:0045944,GO:0046483,GO:0048024,GO:0048025,GO:0048026,GO:0048518,GO:0048519,GO:0048522,GO:0048523,GO:0050684,GO:0050685,GO:0050686,GO:0050789,GO:0050794,GO:0050896,GO:0051171,GO:0051172,GO:0051173,GO:0051252,GO:0051253,GO:0051254,GO:0051259,GO:0051260,GO:0051716,GO:0060255,GO:0065003,GO:0065007,GO:0065010,GO:0070013,GO:0070062,GO:0070271,GO:0070555,GO:0070887,GO:0071013,GO:0071310,GO:0071345,GO:0071347,GO:0071704,GO:0071822,GO:0071840,GO:0080090,GO:0090304,GO:0097159,GO:1901360,GO:1901362,GO:1901363,GO:1901576,GO:2000112,GO:2001141 | RNA binding motif protein |
| comp213031_c4_seq1.p1 | GO:0003674,GO:0003824,GO:0004842,GO:0005575,GO:0005622,GO:0005623,GO:0005634,GO:0005737,GO:0006355,GO:0006464,GO:0007498,GO:0008150,GO:0008152,GO:0008219,GO:0009888,GO:0009889,GO:0009987,GO:0010468,GO:0010556,GO:0012501,GO:0016020,GO:0016021,GO:0016265,GO:0016567,GO:0016874,GO:0016879,GO:0016881,GO:0019219,GO:0019222,GO:0019538,GO:0019787,GO:0031224,GO:0031323,GO:0031326,GO:0032446,GO:0032502,GO:0036211,GO:0043170,GO:0043226,GO:0043227,GO:0043229,GO:0043231,GO:0043412,GO:0044237,GO:0044238,GO:0044260,GO:0044267,GO:0044424,GO:0044425,GO:0044464,GO:0044699,GO:0044763,GO:0048856,GO:0050789,GO:0050794,GO:0051171,GO:0051252,GO:0060255,GO:0065007,GO:0070647,GO:0071704,GO:0080090,GO:2000112,GO:2001141 | Ring finger protein |

| | | |
|---|---|---|
| comp219124_c1_seq11.p1 | GO:0001726,GO:0001817,GO:0001818,GO:0001819,GO:0002218,GO:0002221,GO:0002230,GO:0002252,GO:0002253,GO:0002376,GO:0002682,GO:0002684,GO:0002697,GO:0002753,GO:0002757,GO:0002758,GO:0002764,GO:0002831,GO:0003674,GO:0003676,GO:0003677,GO:0003690,GO:0003723,GO:0003725,GO:0003727,GO:0005488,GO:0005515,GO:0005575,GO:0005622,GO:0005623,GO:0005737,GO:0005829,GO:0005856,GO:0005886,GO:0005911,GO:0005923,GO:0006355,GO:0006357,GO:0006950,GO:0006952,GO:0006955,GO:0007154,GO:0007165,GO:0008150,GO:0008270,GO:0009597,GO:0009607,GO:0009615,GO:0009889,GO:0009891,GO:0009892,GO:0009893,GO:0009987,GO:0010033,GO:0010468,GO:0010556,GO:0010557,GO:0010604,GO:0010605,GO:0010608,GO:0010628,GO:0010629,GO:0014070,GO:0015629,GO:0016020,GO:0016246,GO:0016441,GO:0016458,GO:0019219,GO:0019222,GO:0023052,GO:0030054,GO:0030334,GO:0030522,GO:0031047,GO:0031252,GO:0031253,GO:0031256,GO:0031323,GO:0031325,GO:0031326,GO:0031328,GO:0031347,GO:0031349,GO:0032386,GO:0032388,GO:0032479,GO:0032480,GO:0032481,GO:0032587,GO:0032647,GO:0032648,GO:0032727,GO:0032728,GO:0032879,GO:0032880,GO:0033157,GO:0034344,GO:0035194,GO:0035556,GO:0039528,GO:0039529,GO:0040012,GO:0040029,GO:0042221,GO:0042306,GO:0042307,GO:0042990,GO:0042993,GO:0042995,GO:0043167,GO:0043169,GO:0043226,GO:0043228,GO:0043229,GO:0043232,GO:0043296,GO:0043330,GO:0043331,GO:0043566,GO:0043900,GO:0044093,GO:0044424,GO:0044425,GO:0044444,GO:0044459,GO:0044463,GO:0044464,GO:0044699,GO:0044700,GO:0044763,GO:0045087,GO:0045088,GO:0045089,GO:0045893,GO:0045935,GO:0045944,GO:0046822,GO:0046824,GO:0046872,GO:0046914,GO:0048518,GO:0048519,GO:0048522,GO:0048583,GO:0048584,GO:0050688,GO:0050691,GO:0050776,GO:0050778,GO:0050789,GO:0050794,GO:0050896,GO:0051049,GO:0051050,GO:0051090,GO:0051091,GO:0051171,GO:0051173,GO:0051222,GO:0051223,GO:0051239,GO:0051240,GO:0051241,GO:0051252,GO:0051254,GO:0051270,GO:0051607,GO:0051704,GO:0051707,GO:0051716,GO:0060255,GO:0060341,GO:0065007,GO:0065009,GO:0070160,GO:0070201,GO:0071944,GO:0080090,GO:0080134,GO:0090316,GO:0097159,GO:1900180,GO:1901363,GO:1901698,GO:2000112,GO:2000145,GO:2001141 | DEAD (Asp-Glu-Ala-Asp) box polypeptide 58 |
| comp217845_c0_seq3.p1 | GO:0001662,GO:0002209,GO:0003674,GO:0003824,GO:0004175,GO:0004197,GO:0004221,GO:0004843,GO:0005102,GO:0005488,GO:0005515,GO:0005575,GO:0005622,GO:0005623,GO:0005737,GO:0005768,GO:0006464,GO:0006508,GO:0006511,GO:0006950,GO:0006952,GO:0007610,GO:0008150,GO:0008152,GO:0008233,GO:0008234,GO:0008593,GO:0009056,GO:0009057,GO:0009966,GO:0009968,GO:0009987,GO:0010469,GO:0010646,GO:0010648,GO:0016579,GO:0016787,GO:0016788,GO:0016790,GO:0019538,GO:0019783,GO:0019941,GO:0022898,GO:0023051,GO:0023057,GO:0030163,GO:0031644,GO:0032228,GO:0032409,GO:0032412,GO:0032501,GO:0032879,GO:0033555,GO:0034762,GO:0034765,GO:0035254,GO:0035255,GO:0036211,GO:0040027,GO:0040028,GO:0042596,GO:0043170,GO:0043226,GO:0043227,GO:0043229,GO:0043231,GO:0043269,GO:0043412,GO:0043632,GO:0044057,GO:0044237,GO:0044238,GO:0044248,GO:0044257,GO:0044260,GO:0044265,GO:0044267,GO:0044297,GO:0044424,GO:0044444,GO:0044464,GO:0044699,GO:0044707,GO:0044708,GO:0045746,GO:0048519,GO:0048523,GO:0048580,GO:0048581,GO:0048583,GO:0048585,GO:0050789,GO:0050793,GO:0050794,GO:0050804,GO:0050896,GO:0051049,GO:0051093,GO:0051239,GO:0051241,GO:0051603,GO:0051969,GO:0061062,GO:0061064,GO:0065007,GO:0065009,GO:0070011,GO:0070646,GO:0070647,GO:0071704,GO:0071947,GO:1900449,GO:1901575,GO:2000026,GO:2000311 | ubiquitin carboxyl-terminal hydrolase |

| | | |
|---|---|---|
| comp220250_c1_seq4.p1 | GO:0000003,GO:0000070,GO:0000228,GO:0000278,GO:0000280,GO:0000793,GO:0000794,GO:0000819,GO:0000902,GO:0001701,GO:0001756,GO:0001775,GO:0001932,GO:0002576,GO:0003002,GO:0003007,GO:0003008,GO:0003012,GO:0003013,GO:0003015,GO:0003300,GO:0003674,GO:0003779,GO:0003824,GO:0004672,GO:0004674,GO:0004687,GO:0005198,GO:0005200,GO:0005488,GO:0005509,GO:0005515,GO:0005516,GO:0005575,GO:0005576,GO:0005622,GO:0005623,GO:0005634,GO:0005694,GO:0005737,GO:0005794,GO:0005829,GO:0005856,GO:0005859,GO:0005865,GO:0006139,GO:0006259,GO:0006323,GO:0006461,GO:0006464,GO:0006468,GO:0006725,GO:0006793,GO:0006796,GO:0006807,GO:0006810,GO:0006887,GO:0006928,GO:0006936,GO:0006941,GO:0006950,GO:0006996,GO:0007010,GO:0007015,GO:0007049,GO:0007059,GO:0007067,GO:0007076,GO:0007275,GO:0007389,GO:0007498,GO:0007507,GO:0007512,GO:0007517,GO:0007596,GO:0007599,GO:0008015,GO:0008092,GO:0008150,GO:0008152,GO:0008307,GO:0009056,GO:0009057,GO:0009581,GO:0009582,GO:0009605,GO:0009611,GO:0009612,GO:0009628,GO:0009653,GO:0009790,GO:0009792,GO:0009887,GO:0009888,GO:0009952,GO:0009987,GO:0010035,GO:0010038,GO:0010927,GO:0014706,GO:0014866,GO:0014896,GO:0014897,GO:0015629,GO:0016043,GO:0016192,GO:0016301,GO:0016310,GO:0016459,GO:0016460,GO:0016740,GO:0016772,GO:0016773,GO:0019220,GO:0019222,GO:0019538,GO:0022402,GO:0022607,GO:0030016,GO:0030017,GO:0030018,GO:0030029,GO:0030036,GO:0030048,GO:0030049,GO:0030154,GO:0030163,GO:0030168,GO:0030239,GO:0030240,GO:0030241,GO:0030261,GO:0031032,GO:0031033,GO:0031034,GO:0031323,GO:0031399,GO:0031430,GO:0031433,GO:0031672,GO:0031674,GO:0031974,GO:0031981,GO:0032268,GO:0032501,GO:0032502,GO:0032940,GO:0032989,GO:0032991,GO:0033058,GO:0033275,GO:0034622,GO:0034641,GO:0035051,GO:0035282,GO:0035994,GO:0035995,GO:0036211,GO:0036379,GO:0040008,GO:0040011,GO:0040014,GO:0040015,GO:0040018,GO:0040027,GO:0040028,GO:0042060,GO:0042221,GO:0042325,GO:0042692,GO:0042805,GO:0043009,GO:0043056,GO:0043167,GO:0043169,GO:0043170,GO:0043226,GO:0043227,GO:0043228,GO:0043229,GO:0043231,GO:0043232,GO:0043233,GO:0043234,GO:0043292,GO:0043412,GO:0043549,GO:0043621,GO:0043623,GO:0043933,GO:0044085,GO:0044237,GO:0044238,GO:0044260,GO:0044267,GO:0044422,GO:0044424,GO:0044428,GO:0044430,GO:0044444,GO:0044446,GO:0044449,GO:0044464,GO:0044699,GO:0044707,GO:0044763,GO:0044765,GO:0044767,GO:0045214,GO:0045859,GO:0045926,GO:0045927,GO:0046483,GO:0046872,GO:0046903,GO:0048285,GO:0048468,GO:0048513,GO:0048518,GO:0048519,GO:0048580,GO:0048581,GO:0048644,GO:0048646,GO:0048729,GO:0048731,GO:0048738,GO:0048739,GO:0048747,GO:0048769,GO:0048856,GO:0048869,GO:0050789,GO:0050790,GO:0050793,GO:0050794,GO:0050817,GO:0050878,GO:0050896,GO:0050982,GO:0051015,GO:0051093,GO:0051146,GO:0051174,GO:0051179,GO:0051234,GO:0051239,GO:0051240,GO:0051241,GO:0051246,GO:0051276,GO:0051338,GO:0051592,GO:0051606,GO:0051641,GO:0051649,GO:0055001,GO:0055002,GO:0055003,GO:0055006,GO:0055007,GO:0055008,GO:0055013,GO:0060047,GO:0060048,GO:0060255,GO:0060415,GO:0060537,GO:0061053,GO:0061061,GO:0061062,GO:0061064,GO:0065003,GO:0065007,GO:0065008,GO:0065009,GO:0070013,GO:0070252,GO:0070271,GO:0070925,GO:0071103,GO:0071688,GO:0071704,GO:0071822,GO:0071840,GO:0072358,GO:0072359,GO:0080090,GO:0090304,GO:1901360,GO:1901575,GO:2000026 | myosin light chain kinase |

| | | |
|---|---|---|
| comp197451_c0_seq7.p1 | GO:0001071,GO:0001667,GO:0001763,GO:0002009,GO:0003674,GO:0003676,GO:0003677,GO:0003700,GO:0005488,GO:0005515,GO:0005575,GO:0005622,GO:0005623,GO:0005634,GO:0006355,GO:0006357,GO:0006928,GO:0007275,GO:0007424,GO:0007427,GO:0008150,GO:0009653,GO:0009888,GO:0009889,GO:0009891,GO:0009893,GO:0009987,GO:0010468,GO:0010556,GO:0010557,GO:0010604,GO:0010628,GO:0010631,GO:0016477,GO:0019219,GO:0019222,GO:0031323,GO:0031325,GO:0031326,GO:0031328,GO:0032501,GO:0032502,GO:0032991,GO:0035146,GO:0035147,GO:0035239,GO:0035295,GO:0040011,GO:0043226,GO:0043227,GO:0043229,GO:0043231,GO:0043234,GO:0043565,GO:0044424,GO:0044464,GO:0044699,GO:0044707,GO:0044763,GO:0044767,GO:0045893,GO:0045935,GO:0045944,GO:0046982,GO:0046983,GO:0048518,GO:0048522,GO:0048729,GO:0048731,GO:0048754,GO:0048856,GO:0048870,GO:0050789,GO:0050794,GO:0051171,GO:0051173,GO:0051179,GO:0051252,GO:0051254,GO:0051674,GO:0060255,GO:0060429,GO:0060446,GO:0060541,GO:0060562,GO:0061138,GO:0065007,GO:0080090,GO:0090130,GO:0090132,GO:0097159,GO:1901363,GO:2000112,GO:2001141 | PAS fold |
| comp220484_c1_seq8.p1 | GO:0000981,GO:0000982,GO:0001071,GO:0001077,GO:0001228,GO:0003008,GO:0003012,GO:0003299,GO:0003300,GO:0003674,GO:0003676,GO:0003677,GO:0003682,GO:0003700,GO:0005488,GO:0005515,GO:0005516,GO:0005575,GO:0005622,GO:0005623,GO:0005634,GO:0006139,GO:0006351,GO:0006355,GO:0006357,GO:0006366,GO:0006725,GO:0006807,GO:0006950,GO:0007275,GO:0008134,GO:0008150,GO:0008152,GO:0008277,GO:0009058,GO:0009059,GO:0009314,GO:0009416,GO:0009628,GO:0009790,GO:0009792,GO:0009889,GO:0009891,GO:0009893,GO:0009966,GO:0009987,GO:0010467,GO:0010468,GO:0010556,GO:0010557,GO:0010604,GO:0010628,GO:0010646,GO:0014887,GO:0014888,GO:0014896,GO:0014897,GO:0014898,GO:0016059,GO:0016070,GO:0018130,GO:0019219,GO:0019222,GO:0019438,GO:0019899,GO:0022400,GO:0023051,GO:0031323,GO:0031325,GO:0031326,GO:0031328,GO:0032101,GO:0032501,GO:0032502,GO:0032774,GO:0034641,GO:0034645,GO:0034654,GO:0042802,GO:0042803,GO:0042826,GO:0043170,GO:0043226,GO:0043227,GO:0043229,GO:0043231,GO:0043500,GO:0044237,GO:0044238,GO:0044249,GO:0044260,GO:0044271,GO:0044424,GO:0044464,GO:0044699,GO:0044707,GO:0044767,GO:0045893,GO:0045935,GO:0045944,GO:0046483,GO:0046983,GO:0048518,GO:0048522,GO:0048583,GO:0048856,GO:0050789,GO:0050794,GO:0050896,GO:0051171,GO:0051173,GO:0051252,GO:0051254,GO:0060255,GO:0065007,GO:0071704,GO:0080090,GO:0090304,GO:0097159,GO:1901360,GO:1901362,GO:1901363,GO:1901576,GO:2000112,GO:2001141 | Calmodulin binding transcription activator |
| comp192754_c1_seq2.p1 | | NACHT domain |

| | | |
|---|---|---|
| comp220548_c0_seq9.p1 | GO:0001101,GO:0001678,GO:0002791,GO:0002793,GO:0003674,GO:0003824,GO:0005488,GO:0005515,GO:0005575,GO:0005622,GO:0005623,GO:0005737,GO:0005739,GO:0005759,GO:0006629,GO:0006950,GO:0007275,GO:0007399,GO:0007417,GO:0007420,GO:0007507,GO:0007584,GO:0008150,GO:0008152,GO:0008260,GO:0008410,GO:0009056,GO:0009605,GO:0009719,GO:0009725,GO:0009743,GO:0009746,GO:0009749,GO:0009888,GO:0009987,GO:0009991,GO:0010033,GO:0010646,GO:0014823,GO:0016740,GO:0016782,GO:0019725,GO:0023051,GO:0031667,GO:0031974,GO:0032024,GO:0032501,GO:0032502,GO:0032879,GO:0033500,GO:0034284,GO:0042180,GO:0042182,GO:0042221,GO:0042493,GO:0042592,GO:0042593,GO:0042594,GO:0042802,GO:0042803,GO:0043226,GO:0043227,GO:0043229,GO:0043231,GO:0043233,GO:0044237,GO:0044238,GO:0044248,GO:0044255,GO:0044281,GO:0044282,GO:0044422,GO:0044424,GO:0044429,GO:0044444,GO:0044446,GO:0044464,GO:0044699,GO:0044707,GO:0044710,GO:0044712,GO:0044763,GO:0044767,GO:0045471,GO:0046883,GO:0046887,GO:0046950,GO:0046952,GO:0046983,GO:0048513,GO:0048518,GO:0048522,GO:0048731,GO:0048856,GO:0048878,GO:0050789,GO:0050794,GO:0050796,GO:0050896,GO:0051046,GO:0051047,GO:0051049,GO:0051050,GO:0051716,GO:0055082,GO:0060341,GO:0060612,GO:0061448,GO:0065007,GO:0065008,GO:0070013,GO:0070887,GO:0071229,GO:0071310,GO:0071322,GO:0071326,GO:0071331,GO:0071333,GO:0071704,GO:0072358,GO:0072359,GO:0090087,GO:0090276,GO:0090277,GO:0097305,GO:1901575,GO:1901700,GO:1901701 | Key enzyme for ketone body catabolism. Transfers the CoA moiety from succinate to acetoacetate. Formation of the enzyme-CoA intermediate proceeds via an unstable anhydride species formed between the carboxylate groups of the enzyme and substrate (By similarity) |
| comp210710_c0_seq6.p1 | GO:0005575,GO:0005622,GO:0005623,GO:0005737,GO:0005739,GO:0005759,GO:0031974,GO:0043226,GO:0043227,GO:0043229,GO:0043231,GO:0043233,GO:0044422,GO:0044424,GO:0044429,GO:0044444,GO:0044446,GO:0044464,GO:0070013 | aldehyde dehydrogenase 4 family, member A1 |
| comp217682_c0_seq1.p1 | | chromosome 4 open reading frame 22 |
| comp212971_c1_seq5.p1 | | inhibitor of growth protein 4-like |
| comp215949_c0_seq5.p1 | GO:0005575,GO:0005622,GO:0005623,GO:0005737,GO:0044424,GO:0044464 | zinc finger, CCHC domain containing 2 |
| comp212858_c0_seq2.p1 | GO:0000394,GO:0005575,GO:0005622,GO:0005623,GO:0005634,GO:0005730,GO:0006139,GO:0006388,GO:0006396,GO:0006399,GO:0006725,GO:0006807,GO:0008033,GO:0008150,GO:0008152,GO:0008380,GO:0009987,GO:0010467,GO:0016070,GO:0031974,GO:0031981,GO:0034470,GO:0034641,GO:0034660,GO:0043170,GO:0043226,GO:0043227,GO:0043228,GO:0043229,GO:0043231,GO:0043232,GO:0043233,GO:0044237,GO:0044238,GO:0044260,GO:0044422,GO:0044424,GO:0044428,GO:0044446,GO:0044464,GO:0046483,GO:0070013,GO:0071704,GO:0090304,GO:1901360 | tRNA splicing endonuclease 54 homolog (S. cerevisiae) |
| comp1721978_c0_seq1.p1 | | Reticulon 4 receptor |
| comp213671_c1_seq4.p1 | | HEAT repeat |

| | | |
|---|---|---|
| comp218792_c0_seq1.p1 | GO:0000003,GO:0000166,GO:0002119,GO:0002164,GO:0003674,GO:0003824,GO:0004312,GO:0004313,GO:0004314,GO:0004315,GO:0005488,GO:0005515,GO:0005575,GO:0005622,GO:0005623,GO:0005737,GO:0005739,GO:0005794,GO:0005811,GO:0005829,GO:0005835,GO:0005886,GO:0006082,GO:0006084,GO:0006091,GO:0006112,GO:0006520,GO:0006575,GO:0006629,GO:0006631,GO:0006633,GO:0006637,GO:0006638,GO:0006639,GO:0006641,GO:0006732,GO:0006766,GO:0006767,GO:0006793,GO:0006807,GO:0006915,GO:0007275,GO:0008144,GO:0008150,GO:0008152,GO:0008219,GO:0008610,GO:0009058,GO:0009108,GO:0009790,GO:0009791,GO:0009792,GO:0009893,GO:0009987,GO:0010033,GO:0010876,GO:0012501,GO:0015939,GO:0015980,GO:0016020,GO:0016023,GO:0016053,GO:0016265,GO:0016407,GO:0016417,GO:0016418,GO:0016419,GO:0016420,GO:0016740,GO:0016746,GO:0016747,GO:0019222,GO:0019432,GO:0019752,GO:0019915,GO:0031323,GO:0031325,GO:0031410,GO:0031982,GO:0031988,GO:0032501,GO:0032502,GO:0032787,GO:0032991,GO:0033036,GO:0034097,GO:0034641,GO:0035336,GO:0035337,GO:0035338,GO:0035383,GO:0035384,GO:0036094,GO:0040007,GO:0042221,GO:0042470,GO:0042587,GO:0042802,GO:0042803,GO:0043167,GO:0043168,GO:0043226,GO:0043227,GO:0043229,GO:0043231,GO:0043234,GO:0043436,GO:0043603,GO:0044237,GO:0044238,GO:0044249,GO:0044255,GO:0044281,GO:0044283,GO:0044424,GO:0044444,GO:0044445,GO:0044464,GO:0044699,GO:0044707,GO:0044710,GO:0044711,GO:0044763,GO:0044767,GO:0045017,GO:0046394,GO:0046460,GO:0046463,GO:0046486,GO:0046949,GO:0046983,GO:0048037,GO:0048518,GO:0048522,GO:0048770,GO:0048856,GO:0050661,GO:0050662,GO:0050789,GO:0050794,GO:0050896,GO:0051179,GO:0051186,GO:0051188,GO:0051716,GO:0055114,GO:0065007,GO:0070402,GO:0070670,GO:0070887,GO:0071310,GO:0071345,GO:0071353,GO:0071616,GO:0071704,GO:0071944,GO:0072330,GO:0097159,GO:1901265,GO:1901363,GO:1901564,GO:1901576 | fatty acid synthase |
| comp208854_c0_seq1.p1 | | Fibronectin type III domain |
| comp217941_c0_seq1.p1 | GO:0000323,GO:0003674,GO:0005215,GO:0005216,GO:0005244,GO:0005253,GO:0005342,GO:0005351,GO:0005402,GO:0005436,GO:0005575,GO:0005622,GO:0005623,GO:0005737,GO:0005764,GO:0005765,GO:0005773,GO:0005774,GO:0005783,GO:0005789,GO:0005886,GO:0005887,GO:0005903,GO:0006082,GO:0006725,GO:0006793,GO:0006796,GO:0006807,GO:0006810,GO:0006811,GO:0006812,GO:0006814,GO:0006817,GO:0006818,GO:0006820,GO:0006855,GO:0008021,GO:0008150,GO:0008152,GO:0008308,GO:0008324,GO:0008509,GO:0008514,GO:0008643,GO:0009987,GO:0012505,GO:0012506,GO:0015075,GO:0015077,GO:0015081,GO:0015103,GO:0015114,GO:0015136,GO:0015143,GO:0015144,GO:0015238,GO:0015267,GO:0015291,GO:0015293,GO:0015294,GO:0015295,GO:0015296,GO:0015321,GO:0015562,GO:0015672,GO:0015698,GO:0015711,GO:0015739,GO:0015747,GO:0015849,GO:0015893,GO:0015992,GO:0016020,GO:0016021,GO:0016023,GO:0016324,GO:0019534,GO:0022803,GO:0022804,GO:0022832,GO:0022836,GO:0022838,GO:0022839,GO:0022857,GO:0022890,GO:0022891,GO:0022892,GO:0030001,GO:0030135,GO:0030136,GO:0030659,GO:0030662,GO:0030665,GO:0030672,GO:0031090,GO:0031224,GO:0031226,GO:0031253,GO:0031410,GO:0031526,GO:0031982,GO:0031988,GO:0032879,GO:0034219,GO:0034220,GO:0034641,GO:0034762,GO:0034765,GO:0035435,GO:0035725,GO:0042175,GO:0042221,GO:0042493,GO:0042995,GO:0043226,GO:0043227,GO:0043229,GO:0043231,GO:0043269,GO:0044070,GO:0044237,GO:0044281,GO:0044341,GO:0044422,GO:0044424,GO:0044425,GO:0044432,GO:0044433,GO:0044437,GO:0044444,GO:0044446,GO:0044456,GO:0044459,GO:0044463,GO:0044464,GO:0044699,GO:0044710,GO:0044763,GO:0044765,GO:0045177,GO:0045202,GO:0046415,GO:0046483,GO:0046873,GO:0046942,GO:0046943,GO:0048471,GO:0050789,GO:0050794,GO:0050896,GO:0051049,GO:0051119,GO:0051179,GO:0051234,GO:0055085,GO:0065007,GO:0071702,GO:0071704,GO:0071705,GO:0071944,GO:0072521,GO:0090484,GO:1901264,GO:1901360,GO:1901476,GO:1901505,GO:1901564,GO:1901702,GO:1901998 | solute carrier family 17 (anion sugar transporter), member 5 |

| | | |
|---|---|---|
| comp219797_c1_seq4.p1 | GO:0000375,GO:0000377,GO:0000387,GO:0000398,GO:0000988,GO:0000989,GO:0003674,GO:0003676,GO:0003712,GO:0003713,GO:0003723,GO:0005488,GO:0005515,GO:0005575,GO:0005622,GO:0005623,GO:0005634,GO:0005654,GO:0005737,GO:0005829,GO:0006139,GO:0006396,GO:0006397,GO:0006461,GO:0006725,GO:0006807,GO:0007591,GO:0008150,GO:0008152,GO:0008380,GO:0009889,GO:0009987,GO:0010033,GO:0010467,GO:0010556,GO:0014070,GO:0015030,GO:0016043,GO:0016070,GO:0016071,GO:0016604,GO:0017069,GO:0019219,GO:0019222,GO:0022607,GO:0022613,GO:0022618,GO:0030374,GO:0030529,GO:0031323,GO:0031326,GO:0031974,GO:0031981,GO:0032501,GO:0032991,GO:0033993,GO:0034622,GO:0034641,GO:0034660,GO:0035075,GO:0036314,GO:0042221,GO:0042303,GO:0043170,GO:0043226,GO:0043227,GO:0043229,GO:0043231,GO:0043233,GO:0043933,GO:0044085,GO:0044237,GO:0044238,GO:0044260,GO:0044422,GO:0044424,GO:0044428,GO:0044444,GO:0044446,GO:0044451,GO:0044464,GO:0044699,GO:0044707,GO:0046483,GO:0050789,GO:0050794,GO:0050896,GO:0051171,GO:0051252,GO:0060255,GO:0065003,GO:0065007,GO:0070013,GO:0070271,GO:0071254,GO:0071704,GO:0071822,GO:0071826,GO:0071840,GO:0080090,GO:0090304,GO:0097159,GO:0097305,GO:1901360,GO:1901363,GO:1901654,GO:1901700,GO:2001141 | gem (nuclear organelle) associated protein 5 |
| comp207765_c1_seq3.p1 | GO:0005575,GO:0005622,GO:0005623,GO:0005737,GO:0005739,GO:0006461,GO:0006810,GO:0006891,GO:0006944,GO:0008150,GO:0009987,GO:0010468,GO:0010608,GO:0016020,GO:0016043,GO:0016044,GO:0016192,GO:0019222,GO:0022607,GO:0031647,GO:0043226,GO:0043227,GO:0043229,GO:0043231,GO:0043933,GO:0044085,GO:0044424,GO:0044444,GO:0044464,GO:0044699,GO:0044763,GO:0044765,GO:0046907,GO:0048193,GO:0050789,GO:0050821,GO:0051179,GO:0051234,GO:0051641,GO:0051649,GO:0060255,GO:0061024,GO:0061025,GO:0065003,GO:0065007,GO:0065008,GO:0070271,GO:0071822,GO:0071840 | N-ethylmaleimide-sensitive factor attachment protein, gamma |
| comp207381_c0_seq1.p1 | GO:0005575,GO:0016020 | TLC domain containing |
| comp204971_c0_seq2.p1 | GO:0000975,GO:0000976,GO:0000977,GO:0001012,GO:0001067,GO:0002165,GO:0003674,GO:0003676,GO:0003677,GO:0005488,GO:0005575,GO:0005622,GO:0005623,GO:0005634,GO:0006355,GO:0007275,GO:0007399,GO:0007417,GO:0007444,GO:0007472,GO:0007476,GO:0007552,GO:0007560,GO:0008150,GO:0009653,GO:0009791,GO:0009886,GO:0009887,GO:0009889,GO:0010468,GO:0010556,GO:0019219,GO:0019222,GO:0031323,GO:0031326,GO:0032501,GO:0032502,GO:0035107,GO:0035114,GO:0035120,GO:0035220,GO:0043226,GO:0043227,GO:0043229,GO:0043231,GO:0043565,GO:0044212,GO:0044424,GO:0044464,GO:0044699,GO:0044707,GO:0044767,GO:0048513,GO:0048563,GO:0048569,GO:0048707,GO:0048731,GO:0048736,GO:0048737,GO:0048856,GO:0050789,GO:0050794,GO:0051171,GO:0051252,GO:0060255,GO:0065007,GO:0080090,GO:0097159,GO:1901363,GO:2000112,GO:2001141 | finger homeodomain |
| comp186872_c0_seq1.p1 | GO:0001508,GO:0003008,GO:0005575,GO:0005622,GO:0005623,GO:0005737,GO:0005856,GO:0005886,GO:0005911,GO:0005923,GO:0007154,GO:0007155,GO:0007165,GO:0007272,GO:0007275,GO:0007399,GO:0008150,GO:0008366,GO:0009987,GO:0012505,GO:0014069,GO:0016020,GO:0016324,GO:0016327,GO:0019226,GO:0019228,GO:0022610,GO:0023052,GO:0030054,GO:0030425,GO:0031410,GO:0031982,GO:0032501,GO:0032502,GO:0035556,GO:0035637,GO:0042391,GO:0042552,GO:0042995,GO:0043005,GO:0043209,GO:0043218,GO:0043220,GO:0043226,GO:0043228,GO:0043229,GO:0043232,GO:0043296,GO:0044422,GO:0044424,GO:0044425,GO:0044430,GO:0044444,GO:0044446,GO:0044456,GO:0044459,GO:0044463,GO:0044464,GO:0044699,GO:0044700,GO:0044707,GO:0044763,GO:0045177,GO:0045202,GO:0048731,GO:0048856,GO:0050789,GO:0050794,GO:0050877,GO:0050896,GO:0051716,GO:0065007,GO:0065008,GO:0070160,GO:0071944,GO:0097458 | Multiple PDZ domain protein |

| | | |
|---|---|---|
| comp210494_c1_seq1.p1 | GO:0000003,GO:0000122,GO:0000228,GO:0000785,GO:0000790,GO:0000902,GO:0000904,GO:0000975,GO:0000976,GO:0000981,GO:0000988,GO:0000989,GO:0001067,GO:0001071,GO:0001525,GO:0001568,GO:0001655,GO:0001667,GO:0001701,GO:0001889,GO:0001890,GO:0001893,GO:0001944,GO:0002065,GO:0002067,GO:0002165,GO:0003006,GO:0003007,GO:0003205,GO:0003206,GO:0003208,GO:0003229,GO:0003231,GO:0003674,GO:0003676,GO:0003677,GO:0003682,GO:0003700,GO:0003707,GO:0003708,GO:0003712,GO:0003713,GO:0004871,GO:0004872,GO:0004879,GO:0004884,GO:0004886,GO:0005102,GO:0005488,GO:0005500,GO:0005515,GO:0005575,GO:0005622,GO:0005623,GO:0005634,GO:0005654,GO:0005667,GO:0005694,GO:0005700,GO:0006066,GO:0006139,GO:0006351,GO:0006352,GO:0006355,GO:0006357,GO:0006366,GO:0006367,GO:0006629,GO:0006725,GO:0006766,GO:0006807,GO:0006928,GO:0006950,GO:0006952,GO:0006954,GO:0007154,GO:0007165,GO:0007275,GO:0007276,GO:0007281,GO:0007292,GO:0007297,GO:0007298,GO:0007399,GO:0007417,GO:0007420,GO:0007422,GO:0007431,GO:0007435,GO:0007494,GO:0007507,GO:0007517,GO:0007519,GO:0007552,GO:0007559,GO:0007565,GO:0007566,GO:0007568,GO:0007584,GO:0008134,GO:0008150,GO:0008152,GO:0008202,GO:0008203,GO:0008219,GO:0008230,GO:0008232,GO:0008283,GO:0008285,GO:0008289,GO:0009058,GO:0009059,GO:0009605,GO:0009611,GO:0009653,GO:0009719,GO:0009725,GO:0009755,GO:0009790,GO:0009791,GO:0009792,GO:0009886,GO:0009887,GO:0009888,GO:0009889,GO:0009890,GO:0009891,GO:0009892,GO:0009893,GO:0009966,GO:0009987,GO:0009991,GO:0010033,GO:0010035,GO:0010243,GO:0010269,GO:0010467,GO:0010468,GO:0010469,GO:0010556,GO:0010557,GO:0010558,GO:0010604,GO:0010605,GO:0010628,GO:0010629,GO:0010631,GO:0010646,GO:0010861,GO:0010876,GO:0010941,GO:0010942,GO:0012501,GO:0012502,GO:0014070,GO:0014706,GO:0014855,GO:0016032,GO:0016043,GO:0016070,GO:0016125,GO:0016265,GO:0016271,GO:0016319,GO:0016322,GO:0016358,GO:0016477,GO:0018130,GO:0019048,GO:0019219,GO:0019222,GO:0019438,GO:0019840,GO:0019915,GO:0019953,GO:0021700,GO:0022008,GO:0022412,GO:0022414,GO:0022603,GO:0022612,GO:0023051,GO:0023052,GO:0030030,GO:0030154,GO:0030182,GO:0030374,GO:0030375,GO:0030424,GO:0030425,GO:0030518,GO:0030522,GO:0030545,GO:0030546,GO:0030707,GO:0030850,GO:0030855,GO:0031099,GO:0031102,GO:0031103,GO:0031175,GO:0031323,GO:0031324,GO:0031325,GO:0031326,GO:0031327,GO:0031328,GO:0031490,GO:0031641,GO:0031644,GO:0031667,GO:0031960,GO:0031974,GO:0031981,GO:0032501,GO:0032502,GO:0032504,GO:0032526,GO:0032774,GO:0032868,GO:0032869,GO:0032870,GO:0032989,GO:0032990,GO:0032991,GO:0033002,GO:0033036,GO:0033189,GO:0033273,GO:0033280,GO:0033554,GO:0033993,GO:0034641,GO:0034645,GO:0034654,GO:0035051,GO:0035070,GO:0035071,GO:0035072,GO:0035075,GO:0035076,GO:0035078,GO:0035081,GO:0035257,GO:0035265,GO:0035272,GO:0035357,GO:0035821,GO:0036314,GO:0036315,GO:0038023,GO:0040007,GO:0040011,GO:0040034,GO:0042127,GO:0042221,GO:0042551,GO:0042562,GO:0042692,GO:0042802,GO:0042803,GO:0042809,GO:0042974,GO:0042981,GO:0042995,GO:0043005,GO:0043009,GO:0043065,GO:0043067,GO:0043068,GO:0043170,GO:0043226,GO:0043227,GO:0043228,GO:0043229,GO:0043231,GO:0043232,GO:0043233,GO:0043234,GO:0043235,GO:0043401,GO:0043434,GO:0043565,GO:0043566,GO:0044003,GO:0044057,GO:0044093,GO:0044212,GO:0044237,GO:0044238,GO:0044249,GO:0044255,GO:0044260,GO:0044271,GO:0044281,GO:0044323,GO:0044403,GO:0044419,GO:0044422,GO:0044424,GO:0044427,GO:0044428,GO:0044446,GO:0044451,GO:0044454,GO:0044464,GO:0044699,GO:0044700,GO:0044702,GO:0044703,GO:0044706,GO:0044707,GO:0044710,GO:0044763,GO:0044764,GO:0044767,GO:0045471,GO:0045595,GO:0045596,GO:0045892,GO:0045893,GO:0045934,GO:0045935,GO:0045944,GO:0046483,GO:0046966,GO:0046982,GO:0046983,GO:0048102,GO:0048384,GO:0048468,GO:0048469,GO:0048477,GO:0048513,GO:0048514,GO:0048518,GO:0048519,GO:0048522,GO:0048523,GO:0048545,GO:0048565,GO:0048583,GO:0048608,GO:0048609,GO:0048610,GO:0048644,GO:0048646,GO:0048666,GO:0048667,GO:0048678,GO:0048699,GO:0048707,GO:0048729,GO:0048731,GO:0048732,GO:0048738,GO:0048812,GO:0048813,GO:0048856,GO:0048858,GO:0048869,GO:0048870,GO:0050789,GO:0050793,GO:0050794,GO:0050896,GO:0051093,GO:0051146,GO:0051171,GO:0051172,GO:0051173,GO:0051179,GO:0051239,GO | Retinoid x receptor |
| comp219143_c0_seq1.p1 | | Zinc finger protein |
| comp218196_c2_seq7.p1 | GO:0005575,GO:0005622,GO:0005623,GO:0005634,GO:0043226,GO:0043227,GO:0043229,GO:0043231,GO:0044424,GO:0044464 | Pbx knotted 1 homeobox 2 |
| comp220794_c1_seq8.p1 | | Zinc finger, C2HC type |
| comp220794_c1_seq8.p2 | | myelin transcription factor 1 |
| comp212064_c0_seq15.p1 | | coiled-coil domain containing 41 |

| | | |
|---|---|---|
| comp211478_c0_seq1.p1 | GO:0003674,GO:0003676,GO:0003677,GO:0003684,GO:0005488,GO:0006139,GO:0006259,GO:0006281,GO:0006289,GO:0006725,GO:0006807,GO:0006950,GO:0006974,GO:0008150,GO:0008152,GO:0009987,GO:0033554,GO:0034641,GO:0043170,GO:0044237,GO:0044238,GO:0044260,GO:0044699,GO:0044763,GO:0046483,GO:0050896,GO:0051716,GO:0071704,GO:0090304,GO:0097159,GO:1901360,GO:1901363 | excision repair protein rad23 |
| comp204738_c0_seq1.p1 | GO:0000003,GO:0000075,GO:0000077,GO:0000082,GO:0000084,GO:0000209,GO:0000278,GO:0000502,GO:0002119,GO:0002164,GO:0002376,GO:0002474,GO:0002478,GO:0002479,GO:0003674,GO:0003824,GO:0004175,GO:0005488,GO:0005515,GO:0005575,GO:0005622,GO:0005623,GO:0005634,GO:0005654,GO:0005737,GO:0005829,GO:0005838,GO:0006139,GO:0006464,GO:0006508,GO:0006511,GO:0006521,GO:0006725,GO:0006807,GO:0006915,GO:0006950,GO:0006974,GO:0006977,GO:0007049,GO:0007093,GO:0007154,GO:0007165,GO:0007275,GO:0007346,GO:0007568,GO:0008150,GO:0008152,GO:0008219,GO:0008233,GO:0008340,GO:0008541,GO:0009056,GO:0009057,GO:0009790,GO:0009791,GO:0009792,GO:0009892,GO:0009893,GO:0009987,GO:0010259,GO:0010467,GO:0010498,GO:0010564,GO:0010565,GO:0010604,GO:0010605,GO:0010941,GO:0010948,GO:0012501,GO:0016032,GO:0016070,GO:0016071,GO:0016265,GO:0016567,GO:0016787,GO:0019222,GO:0019538,GO:0019882,GO:0019884,GO:0019941,GO:0022402,GO:0022403,GO:0022624,GO:0023052,GO:0030163,GO:0030330,GO:0031145,GO:0031323,GO:0031324,GO:0031325,GO:0031396,GO:0031397,GO:0031398,GO:0031399,GO:0031400,GO:0031401,GO:0031570,GO:0031571,GO:0031974,GO:0031981,GO:0032268,GO:0032269,GO:0032270,GO:0032446,GO:0032501,GO:0032502,GO:0032991,GO:0033238,GO:0033554,GO:0034641,GO:0035556,GO:0036211,GO:0040007,GO:0042590,GO:0042770,GO:0042981,GO:0043067,GO:0043085,GO:0043086,GO:0043161,GO:0043170,GO:0043226,GO:0043227,GO:0043229,GO:0043231,GO:0043233,GO:0043234,GO:0043412,GO:0043632,GO:0044092,GO:0044093,GO:0044237,GO:0044238,GO:0044248,GO:0044257,GO:0044260,GO:0044265,GO:0044267,GO:0044281,GO:0044403,GO:0044419,GO:0044422,GO:0044424,GO:0044428,GO:0044444,GO:0044446,GO:0044464,GO:0044699,GO:0044700,GO:0044707,GO:0044710,GO:0044763,GO:0044764,GO:0044767,GO:0044770,GO:0044772,GO:0044773,GO:0044774,GO:0044783,GO:0046483,GO:0048002,GO:0048518,GO:0048519,GO:0048522,GO:0048523,GO:0048856,GO:0050789,GO:0050790,GO:0050794,GO:0050896,GO:0051171,GO:0051246,GO:0051247,GO:0051248,GO:0051320,GO:0051325,GO:0051329,GO:0051340,GO:0051351,GO:0051352,GO:0051436,GO:0051437,GO:0051438,GO:0051439,GO:0051443,GO:0051444,GO:0051603,GO:0051704,GO:0051716,GO:0051726,GO:0060255,GO:0065007,GO:0065009,GO:0070011,GO:0070013,GO:0070647,GO:0071156,GO:0071158,GO:0071704,GO:0072331,GO:0072395,GO:0072401,GO:0072413,GO:0072422,GO:0072431,GO:0080090,GO:0090068,GO:0090304,GO:1901360,GO:1901575,GO:1901987,GO:1901988,GO:1901990,GO:1901991,GO:2000045,GO:2000134 | proteasome (prosome, macropain) 26S subunit, non-ATPase, 3 |
| comp217555_c0_seq9.p1 | | Family with sequence similarity 46, member |
| comp220213_c0_seq8.p1 | GO:0005575,GO:0005622,GO:0005623,GO:0005634,GO:0005654,GO:0006139,GO:0006351,GO:0006352,GO:0006354,GO:0006366,GO:0006367,GO:0006368,GO:0006725,GO:0006807,GO:0008150,GO:0008152,GO:0009058,GO:0009059,GO:0009987,GO:0010467,GO:0016032,GO:0016070,GO:0018130,GO:0019438,GO:0031974,GO:0031981,GO:0032774,GO:0034641,GO:0034645,GO:0034654,GO:0043170,GO:0043226,GO:0043227,GO:0043229,GO:0043231,GO:0043233,GO:0044237,GO:0044238,GO:0044249,GO:0044260,GO:0044271,GO:0044403,GO:0044419,GO:0044422,GO:0044424,GO:0044428,GO:0044446,GO:0044464,GO:0044764,GO:0046483,GO:0051704,GO:0070013,GO:0071704,GO:0090304,GO:1901360,GO:1901362,GO:1901576 | General transcription factor IIE, polypeptide 1, alpha 56kDa |
| comp220213_c0_seq8.p2 | | |
| comp221022_c0_seq3.p1 | | adenylate kinase domain containing 1 |

| | | |
|---|---|---|
| comp221022_c0_seq3.p2 | GO:0000323,GO:0005575,GO:0005576,GO:0005615,GO:0005622,GO:0005623,GO:0005737,GO:0005764,GO:0005773,GO:0005775,GO:0005794,GO:0005796,GO:0005886,GO:0005887,GO:0005975,GO:0006022,GO:0006023,GO:0006024,GO:0006026,GO:0006027,GO:0006029,GO:0006082,GO:0006790,GO:0006807,GO:0008150,GO:0008152,GO:0009056,GO:0009057,GO:0009058,GO:0009059,GO:0009100,GO:0009987,GO:0016020,GO:0016021,GO:0019538,GO:0030203,GO:0030204,GO:0031224,GO:0031226,GO:0031974,GO:0043170,GO:0043202,GO:0043226,GO:0043227,GO:0043229,GO:0043231,GO:0043233,GO:0043436,GO:0044237,GO:0044238,GO:0044260,GO:0044281,GO:0044421,GO:0044422,GO:0044424,GO:0044425,GO:0044431,GO:0044437,GO:0044444,GO:0044446,GO:0044459,GO:0044464,GO:0044710,GO:0050654,GO:0070013,GO:0071704,GO:0071944,GO:1901135,GO:1901136,GO:1901137,GO:1901564,GO:1901565,GO:1901566,GO:1901575,GO:1901576 | Cell surface proteoglycan that bears heparan sulfate (By similarity) |
| comp213356_c0_seq2.p1 | GO:0000323,GO:0003674,GO:0005215,GO:0005216,GO:0005244,GO:0005253,GO:0005342,GO:0005351,GO:0005402,GO:0005436,GO:0005575,GO:0005622,GO:0005623,GO:0005737,GO:0005764,GO:0005765,GO:0005773,GO:0005774,GO:0005783,GO:0005789,GO:0005886,GO:0005887,GO:0005903,GO:0006082,GO:0006725,GO:0006793,GO:0006796,GO:0006807,GO:0006810,GO:0006811,GO:0006812,GO:0006814,GO:0006817,GO:0006818,GO:0006820,GO:0006855,GO:0008021,GO:0008150,GO:0008152,GO:0008308,GO:0008324,GO:0008509,GO:0008514,GO:0008643,GO:0009987,GO:0012505,GO:0012506,GO:0015075,GO:0015077,GO:0015081,GO:0015103,GO:0015114,GO:0015136,GO:0015143,GO:0015144,GO:0015238,GO:0015267,GO:0015291,GO:0015293,GO:0015294,GO:0015295,GO:0015296,GO:0015321,GO:0015562,GO:0015672,GO:0015698,GO:0015711,GO:0015739,GO:0015747,GO:0015849,GO:0015893,GO:0015992,GO:0016020,GO:0016021,GO:0016023,GO:0016324,GO:0019534,GO:0022803,GO:0022804,GO:0022832,GO:0022836,GO:0022838,GO:0022839,GO:0022857,GO:0022890,GO:0022891,GO:0022892,GO:0030001,GO:0030135,GO:0030136,GO:0030659,GO:0030662,GO:0030665,GO:0030672,GO:0031090,GO:0031224,GO:0031226,GO:0031253,GO:0031410,GO:0031526,GO:0031982,GO:0031988,GO:0032879,GO:0034219,GO:0034220,GO:0034641,GO:0034762,GO:0034765,GO:0035435,GO:0035725,GO:0042175,GO:0042221,GO:0042493,GO:0042995,GO:0043226,GO:0043227,GO:0043229,GO:0043231,GO:0043269,GO:0044070,GO:0044237,GO:0044281,GO:0044341,GO:0044422,GO:0044424,GO:0044425,GO:0044432,GO:0044433,GO:0044437,GO:0044444,GO:0044446,GO:0044456,GO:0044459,GO:0044463,GO:0044464,GO:0044699,GO:0044710,GO:0044763,GO:0044765,GO:0045177,GO:0045202,GO:0046415,GO:0046483,GO:0046873,GO:0046942,GO:0046943,GO:0048471,GO:0050789,GO:0050794,GO:0050896,GO:0051049,GO:0051119,GO:0051179,GO:0051234,GO:0055085,GO:0065007,GO:0071702,GO:0071704,GO:0071705,GO:0071944,GO:0072521,GO:0090484,GO:1901264,GO:1901360,GO:1901476,GO:1901505,GO:1901564,GO:1901702,GO:1901998 | solute carrier family 17 (anion sugar transporter), member 5 |
| comp208542_c1_seq8.p1 | GO:0000022,GO:0000226,GO:0000278,GO:0005575,GO:0005622,GO:0005623,GO:0005634,GO:0005737,GO:0005783,GO:0006996,GO:0007010,GO:0007017,GO:0007049,GO:0007051,GO:0007052,GO:0008150,GO:0009987,GO:0016043,GO:0022402,GO:0043226,GO:0043227,GO:0043229,GO:0043231,GO:0043933,GO:0044424,GO:0044444,GO:0044464,GO:0044699,GO:0044763,GO:0051231,GO:0071822,GO:0071840 | S-phase cyclin A-associated protein in the ER |
| comp208542_c1_seq8.p2 | GO:0005575,GO:0005622,GO:0005623,GO:0005634,GO:0005737,GO:0005783,GO:0043226,GO:0043227,GO:0043229,GO:0043231,GO:0044424,GO:0044444,GO:0044464 | cyclin A-associated protein in the ER |
| comp194549_c0_seq2.p1 | | ANK |
| comp211966_c0_seq3.p1 | | leucine rich repeat containing 14 |
| comp218975_c0_seq6.p1 | | |

| | | |
|---|---|---|
| comp218975_c0_seq6.p2 | GO:0001654,GO:0001745,GO:0001751,GO:0001754,GO:0002164,GO:0003674,GO:0003824,GO:0005488,GO:0005515,GO:0005575,GO:0005622,GO:0005623,GO:0005634,GO:0005635,GO:0005737,GO:0005811,GO:0005856,GO:0005874,GO:0005875,GO:0005881,GO:0006139,GO:0006152,GO:0006163,GO:0006195,GO:0006200,GO:0006725,GO:0006753,GO:0006793,GO:0006796,GO:0006807,GO:0006810,GO:0006869,GO:0006928,GO:0007017,GO:0007018,GO:0007097,GO:0007275,GO:0007399,GO:0007423,GO:0007424,GO:0007517,GO:0007522,GO:0007523,GO:0008150,GO:0008152,GO:0009056,GO:0009116,GO:0009117,GO:0009119,GO:0009123,GO:0009125,GO:0009126,GO:0009128,GO:0009141,GO:0009143,GO:0009144,GO:0009146,GO:0009150,GO:0009154,GO:0009158,GO:0009161,GO:0009164,GO:0009166,GO:0009167,GO:0009169,GO:0009199,GO:0009203,GO:0009205,GO:0009207,GO:0009259,GO:0009261,GO:0009653,GO:0009791,GO:0009887,GO:0009987,GO:0010876,GO:0010970,GO:0012505,GO:0015630,GO:0016043,GO:0016044,GO:0016462,GO:0016787,GO:0016817,GO:0016818,GO:0016887,GO:0017111,GO:0019439,GO:0019637,GO:0019693,GO:0022008,GO:0030154,GO:0030182,GO:0030286,GO:0030705,GO:0031887,GO:0031967,GO:0031975,GO:0032501,GO:0032502,GO:0032991,GO:0033036,GO:0034641,GO:0034655,GO:0035148,GO:0035149,GO:0035152,GO:0035239,GO:0035295,GO:0040011,GO:0040023,GO:0042051,GO:0042278,GO:0042454,GO:0042461,GO:0042462,GO:0042623,GO:0043226,GO:0043227,GO:0043228,GO:0043229,GO:0043231,GO:0043232,GO:0043234,GO:0044237,GO:0044238,GO:0044248,GO:0044270,GO:0044281,GO:0044422,GO:0044424,GO:0044428,GO:0044430,GO:0044444,GO:0044446,GO:0044464,GO:0044699,GO:0044707,GO:0044710,GO:0044763,GO:0044765,GO:0044767,GO:0046034,GO:0046128,GO:0046130,GO:0046434,GO:0046483,GO:0046530,GO:0046700,GO:0046907,GO:0048468,GO:0048471,GO:0048513,GO:0048569,GO:0048592,GO:0048646,GO:0048666,GO:0048699,GO:0048731,GO:0048749,GO:0048856,GO:0048869,GO:0051179,GO:0051234,GO:0051640,GO:0051641,GO:0051647,GO:0051649,GO:0051656,GO:0055086,GO:0060361,GO:0060541,GO:0061024,GO:0061061,GO:0065007,GO:0065008,GO:0071702,GO:0071704,GO:0071840,GO:0072521,GO:0072523,GO:1901135,GO:1901136,GO:1901292,GO:1901360,GO:1901361,GO:1901564,GO:1901565,GO:1901575,GO:1901657,GO:1901658 | |
| comp214150_c0_seq2.p1 | GO:0000138,GO:0000149,GO:0000938,GO:0003674,GO:0005488,GO:0005515,GO:0005575,GO:0005622,GO:0005623,GO:0005737,GO:0005768,GO:0005794,GO:0005795,GO:0005797,GO:0005802,GO:0007275,GO:0007398,GO:0007439,GO:0008150,GO:0009653,GO:0009790,GO:0009888,GO:0009987,GO:0010008,GO:0010668,GO:0016020,GO:0016023,GO:0017016,GO:0017137,GO:0019899,GO:0019905,GO:0030154,GO:0031090,GO:0031267,GO:0031410,GO:0031982,GO:0031984,GO:0031985,GO:0031988,GO:0032501,GO:0032502,GO:0032588,GO:0032991,GO:0043226,GO:0043227,GO:0043229,GO:0043231,GO:0043234,GO:0044422,GO:0044424,GO:0044431,GO:0044433,GO:0044440,GO:0044444,GO:0044446,GO:0044464,GO:0044699,GO:0044707,GO:0044763,GO:0044767,GO:0048513,GO:0048565,GO:0048566,GO:0048568,GO:0048611,GO:0048729,GO:0048731,GO:0048856,GO:0048869,GO:0051020,GO:0055123 | vacuolar protein sorting 52 homolog (S. cerevisiae) |
| comp211989_c2_seq59.p1 | GO:0000257,GO:0003674,GO:0003824,GO:0004551,GO:0006139,GO:0006725,GO:0006807,GO:0008150,GO:0008152,GO:0009987,GO:0016462,GO:0016787,GO:0016810,GO:0016815,GO:0016817,GO:0016818,GO:0034641,GO:0044237,GO:0044238,GO:0046483,GO:0047710,GO:0071704,GO:1901360 | Nitrilase and fragile histidine triad fusion protein |

| | | |
|---|---|---|
| comp211989_c2_seq59.p2 | GO:0002376,GO:0002520,GO:0003008,GO:0003012,GO:0003013,GO:0003015,GO:0003674,GO:0003824,GO:0005575,GO:0005622,GO:0005623,GO:0005737,GO:0005739,GO:0005740,GO:0005743,GO:0005886,GO:0006091,GO:0006119,GO:0006461,GO:0006629,GO:0006644,GO:0006650,GO:0006655,GO:0006793,GO:0006796,GO:0006936,GO:0006941,GO:0006996,GO:0007005,GO:0007006,GO:0007007,GO:0007275,GO:0007507,GO:0007517,GO:0007519,GO:0008015,GO:0008150,GO:0008152,GO:0008374,GO:0008610,GO:0008654,GO:0009058,GO:0009888,GO:0009966,GO:0009968,GO:0009987,GO:0010257,GO:0010646,GO:0010648,GO:0014706,GO:0015980,GO:0016020,GO:0016043,GO:0016044,GO:0016310,GO:0016740,GO:0016746,GO:0016747,GO:0019637,GO:0019866,GO:0022607,GO:0022900,GO:0022904,GO:0023051,GO:0023057,GO:0030097,GO:0030111,GO:0030178,GO:0031090,GO:0031966,GO:0031967,GO:0031975,GO:0032048,GO:0032049,GO:0032501,GO:0032502,GO:0032981,GO:0033108,GO:0034622,GO:0035965,GO:0042407,GO:0042773,GO:0042775,GO:0043226,GO:0043227,GO:0043229,GO:0043231,GO:0043623,GO:0043933,GO:0044085,GO:0044237,GO:0044238,GO:0044249,GO:0044255,GO:0044281,GO:0044422,GO:0044424,GO:0044429,GO:0044444,GO:0044446,GO:0044464,GO:0044699,GO:0044707,GO:0044710,GO:0044763,GO:0044767,GO:0045017,GO:0045333,GO:0046471,GO:0046474,GO:0046486,GO:0047184,GO:0048513,GO:0048519,GO:0048523,GO:0048534,GO:0048583,GO:0048585,GO:0048731,GO:0048738,GO:0048856,GO:0050789,GO:0050794,GO:0055114,GO:0060047,GO:0060048,GO:0060537,GO:0060538,GO:0060828,GO:0061024,GO:0061061,GO:0065003,GO:0065007,GO:0070271,GO:0071704,GO:0071822,GO:0071840,GO:0071944,GO:0072358,GO:0072359,GO:0090090,GO:0090407,GO:0097031,GO:1901576 | tafazzin |
| comp214029_c0_seq1.p1 | | NADP-dependent oxidoreductase domain containing 1 |
| comp220611_c0_seq19.p1 | GO:0001654,GO:0001736,GO:0001738,GO:0001745,GO:0002009,GO:0002165,GO:0003002,GO:0003008,GO:0003674,GO:0003824,GO:0004672,GO:0004674,GO:0006464,GO:0006468,GO:0006793,GO:0006796,GO:0007154,GO:0007163,GO:0007164,GO:0007267,GO:0007268,GO:0007275,GO:0007389,GO:0007423,GO:0007444,GO:0007472,GO:0007474,GO:0007476,GO:0007552,GO:0007560,GO:0008150,GO:0008152,GO:0008582,GO:0008586,GO:0009605,GO:0009628,GO:0009629,GO:0009653,GO:0009791,GO:0009886,GO:0009887,GO:0009888,GO:0009966,GO:0009968,GO:0009987,GO:0010646,GO:0010648,GO:0016202,GO:0016301,GO:0016310,GO:0016318,GO:0016740,GO:0016772,GO:0016773,GO:0019226,GO:0019538,GO:0023051,GO:0023052,GO:0023057,GO:0030111,GO:0030178,GO:0032501,GO:0032502,GO:0035107,GO:0035114,GO:0035120,GO:0035220,GO:0035637,GO:0036211,GO:0040008,GO:0040011,GO:0042067,GO:0042330,GO:0042332,GO:0043170,GO:0043412,GO:0044087,GO:0044089,GO:0044237,GO:0044238,GO:0044260,GO:0044267,GO:0044699,GO:0044700,GO:0044707,GO:0044763,GO:0044767,GO:0045595,GO:0045887,GO:0045927,GO:0048513,GO:0048518,GO:0048519,GO:0048522,GO:0048523,GO:0048563,GO:0048569,GO:0048583,GO:0048585,GO:0048592,GO:0048634,GO:0048638,GO:0048639,GO:0048641,GO:0048707,GO:0048729,GO:0048731,GO:0048736,GO:0048737,GO:0048742,GO:0048749,GO:0048856,GO:0050789,GO:0050793,GO:0050794,GO:0050803,GO:0050807,GO:0050877,GO:0050896,GO:0051094,GO:0051128,GO:0051130,GO:0051147,GO:0051153,GO:0051239,GO:0051960,GO:0051962,GO:0051963,GO:0051965,GO:0060284,GO:0060429,GO:0065007,GO:0065008,GO:0071704,GO:1901861,GO:2000026 | Protein tyrosine kinase |
| comp211930_c0_seq1.p1 | | GNS1/SUR4 family |
| comp220400_c0_seq6.p1 | | Inherit from veNOG: Nucleoporin 54kDa |
| comp220051_c0_seq2.p1 | GO:0003674,GO:0005083,GO:0005085,GO:0005088,GO:0005488,GO:0005515,GO:0005575,GO:0005622,GO:0005623,GO:0005737,GO:0005829,GO:0008150,GO:0016020,GO:0017112,GO:0019222,GO:0030234,GO:0030695,GO:0032794,GO:0032879,GO:0032880,GO:0044424,GO:0044444,GO:0044464,GO:0050789,GO:0050790,GO:0051049,GO:0051223,GO:0060589,GO:0065007,GO:0065009,GO:0070201 | VPS9 |

| | | |
|---|---|---|
| comp217843_c0_seq2.p1 | | Collagen triple helix repeat (20 copies) |
| comp217843_c0_seq2.p2 | | formin homology 2 domain containing 1 |
| comp221018_c0_seq2.p1 | | Protein of unknown function (DUF3643) |
| comp217348_c0_seq9.p1 | GO:0000003,GO:0000902,GO:0000904,GO:0001654,GO:0001655,GO:0001700,GO:0001745,GO:0001763,GO:0002009,GO:0003002,GO:0003006,GO:0003156,GO:0003674,GO:0003824,GO:0004672,GO:0004674,GO:0004675,GO:0004871,GO:0004872,GO:0004888,GO:0005024,GO:0005026,GO:0005488,GO:0005515,GO:0005575,GO:0005623,GO:0005886,GO:0005887,GO:0006464,GO:0006468,GO:0006793,GO:0006796,GO:0006928,GO:0006935,GO:0007154,GO:0007165,GO:0007166,GO:0007167,GO:0007178,GO:0007179,GO:0007275,GO:0007276,GO:0007281,GO:0007389,GO:0007391,GO:0007399,GO:0007409,GO:0007411,GO:0007423,GO:0007424,GO:0007428,GO:0007442,GO:0007443,GO:0008150,GO:0008152,GO:0008258,GO:0008356,GO:0009605,GO:0009653,GO:0009719,GO:0009790,GO:0009792,GO:0009887,GO:0009888,GO:0009892,GO:0009953,GO:0009987,GO:0010033,GO:0010468,GO:0010605,GO:0010629,GO:0016020,GO:0016021,GO:0016043,GO:0016301,GO:0016310,GO:0016331,GO:0016358,GO:0016740,GO:0016772,GO:0016773,GO:0017002,GO:0017145,GO:0019199,GO:0019222,GO:0019538,GO:0019827,GO:0019953,GO:0022008,GO:0022412,GO:0022414,GO:0022603,GO:0023052,GO:0030030,GO:0030154,GO:0030182,GO:0030718,GO:0031175,GO:0031224,GO:0031226,GO:0032403,GO:0032501,GO:0032502,GO:0032504,GO:0032924,GO:0032989,GO:0032990,GO:0032991,GO:0035239,GO:0035295,GO:0036211,GO:0038023,GO:0040011,GO:0042078,GO:0042221,GO:0042330,GO:0043170,GO:0043234,GO:0043235,GO:0043412,GO:0044237,GO:0044238,GO:0044260,GO:0044267,GO:0044425,GO:0044459,GO:0044464,GO:0044699,GO:0044700,GO:0044702,GO:0044707,GO:0044763,GO:0044767,GO:0045595,GO:0045596,GO:0045704,GO:0045705,GO:0048179,GO:0048185,GO:0048468,GO:0048513,GO:0048519,GO:0048523,GO:0048546,GO:0048565,GO:0048592,GO:0048598,GO:0048609,GO:0048610,GO:0048619,GO:0048666,GO:0048667,GO:0048699,GO:0048729,GO:0048731,GO:0048749,GO:0048754,GO:0048812,GO:0048813,GO:0048856,GO:0048858,GO:0048863,GO:0048864,GO:0048869,GO:0050789,GO:0050793,GO:0050794,GO:0050896,GO:0051093,GO:0051239,GO:0051241,GO:0051301,GO:0051716,GO:0055123,GO:0060089,GO:0060255,GO:0060429,GO:0060446,GO:0060541,GO:0060562,GO:0061138,GO:0061326,GO:0061333,GO:0061525,GO:0061564,GO:0065007,GO:0070848,GO:0070887,GO:0071310,GO:0071363,GO:0071495,GO:0071559,GO:0071560,GO:0071704,GO:0071840,GO:0071944,GO:0072001,GO:0072002,GO:0097485,GO:2000026,GO:2000027 | Activin receptor |
| comp214542_c0_seq4.p1 | GO:0000003,GO:0002119,GO:0002164,GO:0005575,GO:0005622,GO:0005623,GO:0005737,GO:0005886,GO:0006461,GO:0006810,GO:0006897,GO:0006898,GO:0006996,GO:0007010,GO:0007275,GO:0008150,GO:0009653,GO:0009791,GO:0009987,GO:0010927,GO:0016020,GO:0016043,GO:0016192,GO:0022607,GO:0030029,GO:0030036,GO:0030154,GO:0030239,GO:0031032,GO:0031033,GO:0031034,GO:0032501,GO:0032502,GO:0032989,GO:0034622,GO:0040007,GO:0042692,GO:0043226,GO:0043228,GO:0043229,GO:0043232,GO:0043292,GO:0043623,GO:0043933,GO:0044085,GO:0044422,GO:0044424,GO:0044444,GO:0044449,GO:0044464,GO:0044699,GO:0044707,GO:0044763,GO:0044767,GO:0048468,GO:0048646,GO:0048856,GO:0048869,GO:0051146,GO:0051179,GO:0051234,GO:0055001,GO:0055002,GO:0055120,GO:0061061,GO:0065003,GO:0070271,GO:0070925,GO:0071688,GO:0071822,GO:0071840,GO:0071944 | Actin |
| comp215566_c2_seq3.p1 | GO:0002165,GO:0007275,GO:0007444,GO:0007472,GO:0007476,GO:0007552,GO:0007560,GO:0008150,GO:0009653,GO:0009791,GO:0009886,GO:0009887,GO:0032501,GO:0032502,GO:0035107,GO:0035114,GO:0035120,GO:0035220,GO:0044699,GO:0044707,GO:0044767,GO:0048513,GO:0048563,GO:0048569,GO:0048707,GO:0048731,GO:0048736,GO:0048737,GO:0048856 | Pfam:Yippee |

| | | |
|---|---|---|
| comp202469_c0_seq1.p1 | GO:0006950,GO:0006970,GO:0008150,GO:0009628,GO:0009651,GO:0042592,GO:0048878,GO:0050801,GO:0050896,GO:0055081,GO:0065007,GO:0065008 | solute carrier family 26, member 6 |
| comp214348_c0_seq2.p1 | GO:0001085,GO:0003674,GO:0003676,GO:0003723,GO:0003824,GO:0005488,GO:0005515,GO:0006417,GO:0007610,GO:0007635,GO:0008134,GO:0008150,GO:0008152,GO:0008168,GO:0009889,GO:0009890,GO:0009892,GO:0010468,GO:0010556,GO:0010558,GO:0010605,GO:0010608,GO:0016740,GO:0016741,GO:0017069,GO:0017148,GO:0019222,GO:0031323,GO:0031324,GO:0031326,GO:0031327,GO:0032259,GO:0032268,GO:0032269,GO:0042048,GO:0042221,GO:0044708,GO:0048519,GO:0048523,GO:0050789,GO:0050794,GO:0050896,GO:0051246,GO:0051248,GO:0060255,GO:0065007,GO:0080090,GO:0097159,GO:1901363,GO:2000112,GO:2000113 | methyltransferase activity |
| comp217433_c1_seq2.p1 | GO:0000975,GO:0001067,GO:0001071,GO:0002376,GO:0002520,GO:0003002,GO:0003674,GO:0003676,GO:0003677,GO:0003680,GO:0003682,GO:0003700,GO:0003824,GO:0005488,GO:0005515,GO:0005575,GO:0005622,GO:0005623,GO:0005634,GO:0005654,GO:0006139,GO:0006259,GO:0006304,GO:0006305,GO:0006306,GO:0006325,GO:0006351,GO:0006355,GO:0006357,GO:0006366,GO:0006461,GO:0006464,GO:0006473,GO:0006475,GO:0006479,GO:0006725,GO:0006807,GO:0006996,GO:0007275,GO:0007389,GO:0008150,GO:0008152,GO:0008168,GO:0008170,GO:0008213,GO:0008270,GO:0008276,GO:0008285,GO:0008757,GO:0009058,GO:0009059,GO:0009790,GO:0009889,GO:0009891,GO:0009893,GO:0009952,GO:0009987,GO:0010467,GO:0010468,GO:0010556,GO:0010557,GO:0010604,GO:0010628,GO:0016043,GO:0016070,GO:0016278,GO:0016279,GO:0016568,GO:0016569,GO:0016570,GO:0016571,GO:0016573,GO:0016740,GO:0016741,GO:0018022,GO:0018023,GO:0018024,GO:0018130,GO:0018193,GO:0018205,GO:0018393,GO:0018394,GO:0019219,GO:0019222,GO:0019438,GO:0019538,GO:0022607,GO:0030097,GO:0031056,GO:0031060,GO:0031323,GO:0031325,GO:0031326,GO:0031328,GO:0031399,GO:0031974,GO:0031981,GO:0032259,GO:0032268,GO:0032409,GO:0032411,GO:0032501,GO:0032502,GO:0032774,GO:0032879,GO:0032991,GO:0033043,GO:0033044,GO:0034641,GO:0034645,GO:0034654,GO:0034708,GO:0034968,GO:0035097,GO:0035162,GO:0036211,GO:0040029,GO:0042054,GO:0042127,GO:0042393,GO:0042800,GO:0042802,GO:0042803,GO:0043167,GO:0043169,GO:0043170,GO:0043226,GO:0043227,GO:0043229,GO:0043231,GO:0043233,GO:0043234,GO:0043412,GO:0043414,GO:0043543,GO:0043565,GO:0043933,GO:0043967,GO:0043984,GO:0044085,GO:0044093,GO:0044212,GO:0044237,GO:0044238,GO:0044249,GO:0044260,GO:0044267,GO:0044271,GO:0044422,GO:0044424,GO:0044428,GO:0044446,GO:0044451,GO:0044464,GO:0044665,GO:0044699,GO:0044707,GO:0044728,GO:0044763,GO:0044767,GO:0045322,GO:0045893,GO:0045935,GO:0045944,GO:0046483,GO:0046872,GO:0046914,GO:0046983,GO:0048513,GO:0048518,GO:0048519,GO:0048522,GO:0048523,GO:0048534,GO:0048568,GO:0048583,GO:0048584,GO:0048731,GO:0048856,GO:0050789,GO:0050794,GO:0051049,GO:0051050,GO:0051128,GO:0051171,GO:0051173,GO:0051246,GO:0051252,GO:0051254,GO:0051276,GO:0051568,GO:0051569,GO:0060255,GO:0065003,GO:0065007,GO:0065009,GO:0070013,GO:0070271,GO:0070577,GO:0071339,GO:0071704,GO:0071822,GO:0071840,GO:0080090,GO:0080182,GO:0090304,GO:0097159,GO:1901360,GO:1901362,GO:1901363,GO:1901576,GO:1902275,GO:2000112,GO:2001023,GO:2001025,GO:2001038,GO:2001040,GO:2001141 | myeloid lymphoid or mixed-lineage leukemia (trithorax homolog, Drosophila) |
| comp217433_c1_seq2.p2 | GO:0000910,GO:0001932,GO:0001934,GO:0003674,GO:0003824,GO:0004672,GO:0004674,GO:0004702,GO:0004871,GO:0005057,GO:0006464,GO:0006468,GO:0006793,GO:0006796,GO:0007049,GO:0007154,GO:0007165,GO:0008150,GO:0008152,GO:0008283,GO:0009893,GO:0009987,GO:0010562,GO:0010604,GO:0010799,GO:0010800,GO:0016301,GO:0016310,GO:0016740,GO:0016772,GO:0016773,GO:0019220,GO:0019222,GO:0019538,GO:0022402,GO:0023014,GO:0023052,GO:0031323,GO:0031325,GO:0031399,GO:0031401,GO:0032268,GO:0032270,GO:0032879,GO:0036211,GO:0042325,GO:0042327,GO:0042801,GO:0043170,GO:0043412,GO:0044237,GO:0044238,GO:0044260,GO:0044267,GO:0044699,GO:0044700,GO:0044763,GO:0045937,GO:0048518,GO:0048522,GO:0050789,GO:0050794,GO:0050896,GO:0051174,GO:0051246,GO:0051247,GO:0051301,GO:0051716,GO:0060089,GO:0060255,GO:0065007,GO:0071704,GO:0080090 | Polo kinase kinase |

| | | |
|---|---|---|
| comp221139_c0_seq6.p1 | | SEA |
| comp219846_c0_seq3.p1 | | meningioma expressed antigen 5 (hyaluronidase) |
| comp219846_c0_seq3.p2 | | multicellular organismal development |
| comp215538_c0_seq1.p1 | GO:0000166,GO:0000287,GO:0001882,GO:0001883,GO:0003674,GO:0003824,GO:0004611,GO:0004613,GO:0005488,GO:0005525,GO:0005575,GO:0005622,GO:0005623,GO:0005737,GO:0005739,GO:0005759,GO:0005829,GO:0005975,GO:0005996,GO:0006006,GO:0006066,GO:0006071,GO:0006082,GO:0006090,GO:0006094,GO:0006107,GO:0006114,GO:0006464,GO:0006473,GO:0006475,GO:0006629,GO:0006638,GO:0006639,GO:0006641,GO:0008150,GO:0008152,GO:0008610,GO:0009058,GO:0009719,GO:0009725,GO:0009743,GO:0009746,GO:0009749,GO:0009987,GO:0010033,GO:0010243,GO:0014823,GO:0016051,GO:0016829,GO:0016830,GO:0016831,GO:0017076,GO:0017144,GO:0019001,GO:0019003,GO:0019318,GO:0019319,GO:0019400,GO:0019401,GO:0019432,GO:0019538,GO:0019751,GO:0019752,GO:0030145,GO:0031406,GO:0031974,GO:0032549,GO:0032550,GO:0032553,GO:0032555,GO:0032561,GO:0032787,GO:0032868,GO:0033500,GO:0034284,GO:0034637,GO:0035639,GO:0036094,GO:0036211,GO:0042221,GO:0042592,GO:0042593,GO:0043167,GO:0043168,GO:0043169,GO:0043170,GO:0043226,GO:0043227,GO:0043228,GO:0043229,GO:0043231,GO:0043232,GO:0043233,GO:0043292,GO:0043412,GO:0043434,GO:0043436,GO:0043543,GO:0043648,GO:0044237,GO:0044238,GO:0044249,GO:0044255,GO:0044260,GO:0044262,GO:0044267,GO:0044281,GO:0044283,GO:0044422,GO:0044424,GO:0044429,GO:0044444,GO:0044446,GO:0044449,GO:0044464,GO:0044710,GO:0044711,GO:0044723,GO:0045017,GO:0046165,GO:0046173,GO:0046327,GO:0046364,GO:0046460,GO:0046463,GO:0046486,GO:0046872,GO:0046914,GO:0048878,GO:0050896,GO:0055120,GO:0065007,GO:0065008,GO:0070013,GO:0071704,GO:0097159,GO:1901265,GO:1901363,GO:1901576,GO:1901615,GO:1901617,GO:1901652,GO:1901698,GO:1901700 | phosphoenolpyruvate carboxykinase |
| comp214452_c0_seq1.p1 | | G1 to S phase transition 1 |

| | | |
|---|---|---|
| comp214452_c0_seq1.p2 | GO:0000003,GO:0000082,GO:0000166,GO:0000184,GO:0000278,GO:0000956,GO:0001882,GO:0001883,GO:0002119,GO:0002164,GO:0003674,GO:0003676,GO:0003723,GO:0003747,GO:0003824,GO:0003924,GO:0005488,GO:0005525,GO:0005575,GO:0005622,GO:0005623,GO:0005737,GO:0005829,GO:0006139,GO:0006152,GO:0006163,GO:0006184,GO:0006195,GO:0006401,GO:0006402,GO:0006412,GO:0006415,GO:0006464,GO:0006479,GO:0006725,GO:0006753,GO:0006793,GO:0006796,GO:0006807,GO:0006810,GO:0006897,GO:0006898,GO:0006915,GO:0007049,GO:0007275,GO:0008079,GO:0008135,GO:0008150,GO:0008152,GO:0008213,GO:0008219,GO:0009056,GO:0009057,GO:0009058,GO:0009059,GO:0009116,GO:0009117,GO:0009119,GO:0009141,GO:0009143,GO:0009144,GO:0009146,GO:0009150,GO:0009154,GO:0009164,GO:0009166,GO:0009199,GO:0009203,GO:0009205,GO:0009207,GO:0009259,GO:0009261,GO:0009790,GO:0009791,GO:0009792,GO:0009987,GO:0010467,GO:0012501,GO:0016043,GO:0016070,GO:0016071,GO:0016192,GO:0016265,GO:0016462,GO:0016787,GO:0016817,GO:0016818,GO:0017076,GO:0017111,GO:0019001,GO:0019439,GO:0019538,GO:0019637,GO:0019693,GO:0022402,GO:0022403,GO:0022411,GO:0032259,GO:0032501,GO:0032502,GO:0032549,GO:0032550,GO:0032553,GO:0032555,GO:0032561,GO:0032984,GO:0034641,GO:0034645,GO:0034655,GO:0035639,GO:0036094,GO:0036211,GO:0040007,GO:0040008,GO:0040009,GO:0040010,GO:0040011,GO:0040014,GO:0040018,GO:0042278,GO:0042454,GO:0043167,GO:0043168,GO:0043170,GO:0043241,GO:0043412,GO:0043414,GO:0043624,GO:0043933,GO:0044237,GO:0044238,GO:0044248,GO:0044249,GO:0044260,GO:0044265,GO:0044267,GO:0044270,GO:0044281,GO:0044424,GO:0044444,GO:0044464,GO:0044699,GO:0044707,GO:0044710,GO:0044763,GO:0044767,GO:0044770,GO:0044772,GO:0045927,GO:0046039,GO:0046128,GO:0046130,GO:0046434,GO:0046483,GO:0046700,GO:0048518,GO:0048856,GO:0050789,GO:0051179,GO:0051234,GO:0051239,GO:0051240,GO:0051325,GO:0051329,GO:0055086,GO:0065007,GO:0071704,GO:0071822,GO:0071840,GO:0072521,GO:0072523,GO:0090304,GO:0097159,GO:1901068,GO:1901069,GO:1901135,GO:1901136,GO:1901265,GO:1901292,GO:1901360,GO:1901361,GO:1901363,GO:1901564,GO:1901565,GO:1901575,GO:1901576,GO:1901657,GO:1901658 | G1 to S phase transition |
| comp214608_c0_seq1.p1 | | Viral A-type inclusion protein |
| comp220309_c0_seq5.p1 | | Zinc finger protein |
| comp218949_c1_seq13.p1 | | alpha/beta hydrolase fold |
| comp218949_c1_seq13.p2 | | Inherit from opiNOG: protein Hydra magnipapillata |
| comp217005_c1_seq2.p1 | GO:0000003,GO:0003674,GO:0005488,GO:0005515,GO:0005520,GO:0005575,GO:0005576,GO:0005615,GO:0005622,GO:0005623,GO:0005634,GO:0007154,GO:0007165,GO:0007275,GO:0007276,GO:0007292,GO:0008150,GO:0009790,GO:0009792,GO:0009987,GO:0016942,GO:0019838,GO:0019953,GO:0022414,GO:0023052,GO:0032501,GO:0032502,GO:0032504,GO:0032991,GO:0040008,GO:0040009,GO:0040010,GO:0040011,GO:0040014,GO:0040015,GO:0042567,GO:0043226,GO:0043227,GO:0043229,GO:0043231,GO:0043234,GO:0044421,GO:0044424,GO:0044464,GO:0044699,GO:0044700,GO:0044702,GO:0044707,GO:0044763,GO:0044767,GO:0045926,GO:0045927,GO:0048477,GO:0048518,GO:0048519,GO:0048609,GO:0048856,GO:0050789,GO:0050794,GO:0050896,GO:0051239,GO:0051241,GO:0051716,GO:0065007 | Insulin-like growth factor binding protein, acid labile subunit |

| | | |
|---|---|---|
| comp208701_c0_seq2.p1 | GO:0000166,GO:0001882,GO:0001883,GO:0002376,GO:0003674,GO:0003824,GO:0004672,GO:0004674,GO:0005488,GO:0005515,GO:0005516,GO:0005524,GO:0005575,GO:0005622,GO:0005623,GO:0005737,GO:0005829,GO:0005856,GO:0006417,GO:0006464,GO:0006468,GO:0006793,GO:0006796,GO:0006915,GO:0006917,GO:0006950,GO:0006952,GO:0006955,GO:0007154,GO:0007165,GO:0007243,GO:0008150,GO:0008152,GO:0008219,GO:0009889,GO:0009890,GO:0009892,GO:0009894,GO:0009966,GO:0009987,GO:0010033,GO:0010468,GO:0010469,GO:0010506,GO:0010556,GO:0010558,GO:0010605,GO:0010608,GO:0010646,GO:0010941,GO:0010942,GO:0012501,GO:0012502,GO:0015629,GO:0016265,GO:0016301,GO:0016310,GO:0016740,GO:0016772,GO:0016773,GO:0017076,GO:0017148,GO:0019222,GO:0019538,GO:0022898,GO:0023051,GO:0023052,GO:0030554,GO:0031323,GO:0031324,GO:0031326,GO:0031327,GO:0031329,GO:0032268,GO:0032269,GO:0032409,GO:0032412,GO:0032549,GO:0032550,GO:0032553,GO:0032555,GO:0032559,GO:0032879,GO:0034097,GO:0034341,GO:0034762,GO:0034765,GO:0035556,GO:0035639,GO:0036094,GO:0036211,GO:0042221,GO:0042981,GO:0043065,GO:0043066,GO:0043067,GO:0043068,GO:0043069,GO:0043167,GO:0043168,GO:0043170,GO:0043226,GO:0043228,GO:0043229,GO:0043232,GO:0043269,GO:0043412,GO:0044237,GO:0044238,GO:0044260,GO:0044267,GO:0044424,GO:0044444,GO:0044464,GO:0044699,GO:0044700,GO:0044763,GO:0045087,GO:0046777,GO:0048518,GO:0048519,GO:0048522,GO:0048523,GO:0048583,GO:0050789,GO:0050794,GO:0050896,GO:0051049,GO:0051246,GO:0051248,GO:0051716,GO:0060255,GO:0060548,GO:0065007,GO:0065009,GO:0070887,GO:0071310,GO:0071345,GO:0071346,GO:0071704,GO:0080090,GO:0097159,GO:0097190,GO:1900449,GO:1901265,GO:1901363,GO:2000112,GO:2000113,GO:2000310 | death-associated protein kinase 1 |
| comp220835_c1_seq4.p1 | | FERM, RhoGEF and pleckstrin domain protein 2 |
| comp198525_c0_seq2.p1 | | Ankyrin repeat |
| comp84898_c0_seq1.p1 | | histamine receptor H4 |
| comp218273_c1_seq2.p1 | GO:0000103,GO:0000166,GO:0001501,GO:0001882,GO:0001883,GO:0002165,GO:0003674,GO:0003824,GO:0004020,GO:0004779,GO:0004781,GO:0005488,GO:0005524,GO:0005575,GO:0005622,GO:0005623,GO:0005737,GO:0005829,GO:0006082,GO:0006139,GO:0006163,GO:0006164,GO:0006725,GO:0006753,GO:0006790,GO:0006793,GO:0006796,GO:0006805,GO:0006807,GO:0006950,GO:0007275,GO:0007444,GO:0007472,GO:0007476,GO:0007552,GO:0007560,GO:0007568,GO:0007596,GO:0007599,GO:0008150,GO:0008152,GO:0008340,GO:0009058,GO:0009116,GO:0009117,GO:0009119,GO:0009150,GO:0009152,GO:0009163,GO:0009165,GO:0009259,GO:0009260,GO:0009410,GO:0009611,GO:0009653,GO:0009791,GO:0009886,GO:0009887,GO:0009987,GO:0010259,GO:0016301,GO:0016310,GO:0016740,GO:0016772,GO:0016773,GO:0016779,GO:0017076,GO:0018130,GO:0019438,GO:0019637,GO:0019693,GO:0030554,GO:0032501,GO:0032502,GO:0032549,GO:0032550,GO:0032553,GO:0032555,GO:0032559,GO:0033865,GO:0033866,GO:0033875,GO:0034030,GO:0034032,GO:0034033,GO:0034035,GO:0034036,GO:0034641,GO:0034654,GO:0035107,GO:0035114,GO:0035120,GO:0035220,GO:0035639,GO:0036094,GO:0042060,GO:0042221,GO:0042278,GO:0042451,GO:0042455,GO:0043167,GO:0043168,GO:0043436,GO:0044237,GO:0044238,GO:0044249,GO:0044271,GO:0044272,GO:0044281,GO:0044424,GO:0044444,GO:0044464,GO:0044699,GO:0044707,GO:0044710,GO:0044763,GO:0044767,GO:0046128,GO:0046129,GO:0046390,GO:0046483,GO:0048513,GO:0048563,GO:0048569,GO:0048707,GO:0048731,GO:0048736,GO:0048737,GO:0048856,GO:0050427,GO:0050428,GO:0050817,GO:0050878,GO:0050896,GO:0051716,GO:0055086,GO:0060348,GO:0065007,GO:0065008,GO:0070566,GO:0070887,GO:0071466,GO:0071704,GO:0072521,GO:0072522,GO:0090407,GO:0097159,GO:1901135,GO:1901137,GO:1901265,GO:1901293,GO:1901360,GO:1901362,GO:1901363,GO:1901564,GO:1901566,GO:1901576,GO:1901657,GO:1901659 | 3'-phosphoadenosine 5'-phosphosulfate synthase |

| | | |
|---|---|---|
| comp211847_c2_seq1.p1 | GO:0000003,GO:0000086,GO:0000139,GO:0000166,GO:0000278,GO:0001666,GO:0001882,GO:0001883,GO:0001885,GO:0002064,GO:0003158,GO:0003674,GO:0003824,GO:0005215,GO:0005488,GO:0005524,GO:0005575,GO:0005622,GO:0005623,GO:0005737,GO:0005739,GO:0005794,GO:0005886,GO:0005887,GO:0005911,GO:0006139,GO:0006152,GO:0006163,GO:0006195,GO:0006200,GO:0006629,GO:0006725,GO:0006753,GO:0006793,GO:0006796,GO:0006807,GO:0006810,GO:0006855,GO:0006950,GO:0007049,GO:0007275,GO:0007399,GO:0007417,GO:0007584,GO:0007589,GO:0007595,GO:0007623,GO:0008150,GO:0008152,GO:0008283,GO:0009056,GO:0009116,GO:0009117,GO:0009119,GO:0009123,GO:0009125,GO:0009126,GO:0009128,GO:0009141,GO:0009143,GO:0009144,GO:0009146,GO:0009150,GO:0009154,GO:0009158,GO:0009161,GO:0009164,GO:0009166,GO:0009167,GO:0009169,GO:0009199,GO:0009203,GO:0009205,GO:0009207,GO:0009259,GO:0009261,GO:0009314,GO:0009410,GO:0009605,GO:0009628,GO:0009719,GO:0009725,GO:0009888,GO:0009986,GO:0009987,GO:0009991,GO:0010033,GO:0010035,GO:0010038,GO:0010212,GO:0010941,GO:0012505,GO:0014045,GO:0014070,GO:0015238,GO:0015399,GO:0015405,GO:0015562,GO:0015893,GO:0016020,GO:0016021,GO:0016324,GO:0016462,GO:0016787,GO:0016817,GO:0016818,GO:0016820,GO:0016887,GO:0017076,GO:0017111,GO:0019439,GO:0019637,GO:0019693,GO:0022402,GO:0022414,GO:0022804,GO:0022857,GO:0030054,GO:0030154,GO:0030554,GO:0030855,GO:0030879,GO:0031090,GO:0031224,GO:0031226,GO:0031667,GO:0031960,GO:0032501,GO:0032502,GO:0032504,GO:0032549,GO:0032550,GO:0032553,GO:0032555,GO:0032559,GO:0032941,GO:0033273,GO:0033280,GO:0033993,GO:0034097,GO:0034612,GO:0034641,GO:0034655,GO:0035639,GO:0036094,GO:0036293,GO:0042221,GO:0042278,GO:0042391,GO:0042454,GO:0042493,GO:0042623,GO:0042626,GO:0043167,GO:0043168,GO:0043226,GO:0043227,GO:0043229,GO:0043231,GO:0043492,GO:0044237,GO:0044238,GO:0044248,GO:0044255,GO:0044270,GO:0044281,GO:0044422,GO:0044424,GO:0044425,GO:0044431,GO:0044444,GO:0044446,GO:0044459,GO:0044464,GO:0044699,GO:0044707,GO:0044710,GO:0044763,GO:0044765,GO:0044767,GO:0044770,GO:0044772,GO:0045177,GO:0045446,GO:0046034,GO:0046128,GO:0046130,GO:0046434,GO:0046483,GO:0046581,GO:0046685,GO:0046686,GO:0046700,GO:0046903,GO:0048468,GO:0048511,GO:0048513,GO:0048519,GO:0048523,GO:0048545,GO:0048609,GO:0048731,GO:0048732,GO:0048856,GO:0048869,GO:0050789,GO:0050794,GO:0050878,GO:0050896,GO:0051179,GO:0051234,GO:0051384,GO:0051716,GO:0055085,GO:0055086,GO:0060429,GO:0060548,GO:0060856,GO:0061028,GO:0065007,GO:0065008,GO:0070482,GO:0070887,GO:0071310,GO:0071345,GO:0071356,GO:0071704,GO:0071944,GO:0072089,GO:0072521,GO:0072523,GO:0090484,GO:0097159,GO:0097305,GO:1901135,GO:1901136,GO:1901265,GO:1901292,GO:1901360,GO:1901361,GO:1901363,GO:1901564,GO:1901565,GO:1901575,GO:1901657,GO:1901658,GO:1901700 | ATP-binding cassette, subfamily B (MDR TAP), member |
| comp213345_c0_seq1.p1 | GO:0000003,GO:0000026,GO:0000030,GO:0002119,GO:0002164,GO:0003674,GO:0003824,GO:0005575,GO:0005622,GO:0005623,GO:0005737,GO:0005783,GO:0005789,GO:0005975,GO:0006464,GO:0006486,GO:0006487,GO:0006488,GO:0006810,GO:0006897,GO:0006898,GO:0007275,GO:0007399,GO:0008150,GO:0008152,GO:0009058,GO:0009059,GO:0009100,GO:0009101,GO:0009791,GO:0009987,GO:0012505,GO:0016020,GO:0016192,GO:0016740,GO:0016757,GO:0016758,GO:0018193,GO:0018196,GO:0018279,GO:0019538,GO:0022008,GO:0030154,GO:0031090,GO:0032501,GO:0032502,GO:0034645,GO:0036211,GO:0040007,GO:0040011,GO:0042175,GO:0043170,GO:0043226,GO:0043227,GO:0043229,GO:0043231,GO:0043412,GO:0043413,GO:0043687,GO:0044237,GO:0044238,GO:0044249,GO:0044260,GO:0044267,GO:0044422,GO:0044424,GO:0044425,GO:0044432,GO:0044444,GO:0044446,GO:0044464,GO:0044699,GO:0044707,GO:0044723,GO:0044763,GO:0048731,GO:0048856,GO:0048869,GO:0051179,GO:0051234,GO:0070085,GO:0071704,GO:1901135,GO:1901137,GO:1901576 | asparagine-linked glycosylation 11, alpha-1,2-mannosyltransferase homolog (yeast) |

| | | |
|---|---|---|
| comp219769_c0_seq6.p1 | GO:0000003,GO:0000166,GO:0001882,GO:0001883,GO:0001947,GO:0002009,GO:0002026,GO:0002027,GO:0002119,GO:0002164,GO:0003007,GO:0003008,GO:0003012,GO:0003143,GO:0003674,GO:0003676,GO:0003677,GO:0003682,GO:0003824,GO:0005215,GO:0005388,GO:0005488,GO:0005509,GO:0005515,GO:0005524,GO:0005575,GO:0005622,GO:0005623,GO:0005634,GO:0005635,GO:0005737,GO:0005783,GO:0005789,GO:0005793,GO:0005811,GO:0005886,GO:0005887,GO:0006139,GO:0006152,GO:0006163,GO:0006195,GO:0006200,GO:0006461,GO:0006725,GO:0006753,GO:0006793,GO:0006796,GO:0006807,GO:0006810,GO:0006811,GO:0006812,GO:0006816,GO:0006873,GO:0006874,GO:0006875,GO:0006897,GO:0006898,GO:0006915,GO:0006937,GO:0006942,GO:0006950,GO:0006984,GO:0006986,GO:0006996,GO:0007005,GO:0007010,GO:0007154,GO:0007155,GO:0007165,GO:0007267,GO:0007268,GO:0007274,GO:0007275,GO:0007276,GO:0007292,GO:0007368,GO:0007389,GO:0007507,GO:0007568,GO:0007596,GO:0007599,GO:0007610,GO:0007626,GO:0007629,GO:0008016,GO:0008150,GO:0008152,GO:0008219,GO:0008324,GO:0008340,GO:0008344,GO:0008544,GO:0008637,GO:0009056,GO:0009116,GO:0009117,GO:0009119,GO:0009123,GO:0009125,GO:0009126,GO:0009128,GO:0009141,GO:0009143,GO:0009144,GO:0009146,GO:0009150,GO:0009154,GO:0009158,GO:0009161,GO:0009164,GO:0009166,GO:0009167,GO:0009169,GO:0009199,GO:0009203,GO:0009205,GO:0009207,GO:0009259,GO:0009261,GO:0009605,GO:0009611,GO:0009612,GO:0009628,GO:0009653,GO:0009719,GO:0009725,GO:0009790,GO:0009791,GO:0009792,GO:0009799,GO:0009855,GO:0009887,GO:0009888,GO:0009987,GO:0010033,GO:0010243,GO:0010259,GO:0010460,GO:0010927,GO:0012501,GO:0012505,GO:0012506,GO:0014724,GO:0014819,GO:0015075,GO:0015085,GO:0015399,GO:0015405,GO:0015662,GO:0016020,GO:0016021,GO:0016043,GO:0016192,GO:0016265,GO:0016462,GO:0016528,GO:0016529,GO:0016787,GO:0016817,GO:0016818,GO:0016820,GO:0016887,GO:0017076,GO:0017111,GO:0018996,GO:0019226,GO:0019439,GO:0019637,GO:0019693,GO:0019725,GO:0019829,GO:0019953,GO:0022414,GO:0022607,GO:0022610,GO:0022804,GO:0022857,GO:0022890,GO:0022891,GO:0022892,GO:0023052,GO:0030001,GO:0030003,GO:0030016,GO:0030017,GO:0030029,GO:0030036,GO:0030104,GO:0030154,GO:0030239,GO:0030322,GO:0030534,GO:0030554,GO:0030968,GO:0031032,GO:0031033,GO:0031034,GO:0031090,GO:0031094,GO:0031095,GO:0031224,GO:0031226,GO:0031410,GO:0031446,GO:0031448,GO:0031490,GO:0031672,GO:0031673,GO:0031674,GO:0031965,GO:0031967,GO:0031975,GO:0031982,GO:0031988,GO:0032469,GO:0032470,GO:0032471,GO:0032501,GO:0032502,GO:0032504,GO:0032549,GO:0032550,GO:0032553,GO:0032555,GO:0032559,GO:0032844,GO:0032879,GO:0032989,GO:0032991,GO:0033017,GO:0033554,GO:0034220,GO:0034620,GO:0034622,GO:0034641,GO:0034655,GO:0034702,GO:0034703,GO:0034704,GO:0034976,GO:0035050,GO:0035239,GO:0035295,GO:0035556,GO:0035637,GO:0035639,GO:0035966,GO:0035967,GO:0036094,GO:0040007,GO:0040024,GO:0042060,GO:0042175,GO:0042221,GO:0042278,GO:0042303,GO:0042391,GO:0042454,GO:0042493,GO:0042592,GO:0042623,GO:0042625,GO:0042626,GO:0042692,GO:0043167,GO:0043168,GO:0043169,GO:0043226,GO:0043227,GO:0043228,GO:0043229,GO:0043231,GO:0043232,GO:0043234,GO:0043292,GO:0043434,GO:0043492,GO:0043566,GO:0043588,GO:0043623,GO:0043933,GO:0044057,GO:0044085,GO:0044237,GO:0044238,GO:0044248,GO:0044270,GO:0044281,GO:0044422,GO:0044424,GO:0044425,GO:0044428,GO:0044432,GO:0044444,GO:0044446,GO:0044449,GO:0044459,GO:0044464,GO:0044699,GO:0044700,GO:0044702,GO:0044707,GO:0044708,GO:0044710,GO:0044763,GO:0044765,GO:0044767,GO:0045822,GO:0045823,GO:0045932,GO:0045933,GO:0045988,GO:0045989,GO:0046034,GO:0046128,GO:0046130,GO:0046434,GO:0046483,GO:0046700,GO:0046872,GO:0046873,GO:0046907,GO:0048468,GO:0048471,GO:0048477,GO:0048513,GO:0048518,GO:0048519,GO:0048562,GO:0048568,GO:0048598,GO:0048609,GO:0048646,GO:0048729,GO:0048731,GO:0048856,GO:0048869,GO:0048878,GO:0050789,GO:0050794,GO:0050801,GO:0050817,GO:0050877,GO:0050878,GO:0050896,GO:0051146,GO:0051179,GO:0051234,GO:0051235,GO:0051239,GO:0051240,GO:0051241,GO:0051282,GO:0051560,GO:0051561,GO:0051640,GO:0051641,GO:0051646,GO:0051649,GO:0051651,GO:0051657,GO:0051659,GO:0051716,GO:0055001,GO:0055002,GO:0055065,GO:0055074,GO:0055080,GO:0055082,GO:0055085,GO:0055086,GO:0055119,GO:0060429,GO | ATPase CA transporting cardiac muscle |

| | | |
|---|---|---|
| comp210899_c0_seq5.p1 | GO:0000003,GO:0000226,GO:0000578,GO:0001736,GO:0001737,GO:0001738,GO:0002009,GO:0002165,GO:0003002,GO:0003006,GO:0003674,GO:0005488,GO:0005543,GO:0005575,GO:0005622,GO:0005623,GO:0005737,GO:0005768,GO:0005769,GO:0005886,GO:0006403,GO:0006810,GO:0006897,GO:0006906,GO:0006944,GO:0006950,GO:0006996,GO:0007010,GO:0007017,GO:0007028,GO:0007163,GO:0007164,GO:0007275,GO:0007276,GO:0007281,GO:0007292,GO:0007308,GO:0007309,GO:0007314,GO:0007315,GO:0007316,GO:0007350,GO:0007351,GO:0007389,GO:0007399,GO:0007444,GO:0007472,GO:0007476,GO:0007552,GO:0007560,GO:0007596,GO:0007599,GO:0008103,GO:0008150,GO:0008270,GO:0008289,GO:0008298,GO:0008358,GO:0008544,GO:0008595,GO:0009611,GO:0009653,GO:0009790,GO:0009791,GO:0009798,GO:0009880,GO:0009886,GO:0009887,GO:0009888,GO:0009913,GO:0009948,GO:0009952,GO:0009987,GO:0009994,GO:0010008,GO:0016020,GO:0016043,GO:0016044,GO:0016050,GO:0016192,GO:0016197,GO:0016325,GO:0019094,GO:0019953,GO:0021700,GO:0022008,GO:0022412,GO:0022414,GO:0022607,GO:0030030,GO:0030100,GO:0030154,GO:0030182,GO:0030855,GO:0030951,GO:0030952,GO:0031090,GO:0031901,GO:0032266,GO:0032501,GO:0032502,GO:0032504,GO:0032879,GO:0033036,GO:0034058,GO:0035088,GO:0035091,GO:0035107,GO:0035114,GO:0035120,GO:0035220,GO:0035282,GO:0035315,GO:0035316,GO:0035317,GO:0042060,GO:0043167,GO:0043168,GO:0043169,GO:0043226,GO:0043227,GO:0043229,GO:0043231,GO:0043588,GO:0044085,GO:0044422,GO:0044424,GO:0044440,GO:0044444,GO:0044446,GO:0044464,GO:0044699,GO:0044702,GO:0044707,GO:0044763,GO:0044765,GO:0044767,GO:0045451,GO:0046872,GO:0046907,GO:0046914,GO:0048284,GO:0048468,GO:0048469,GO:0048477,GO:0048513,GO:0048563,GO:0048569,GO:0048599,GO:0048609,GO:0048610,GO:0048699,GO:0048707,GO:0048729,GO:0048731,GO:0048736,GO:0048737,GO:0048856,GO:0048869,GO:0050789,GO:0050794,GO:0050817,GO:0050878,GO:0050896,GO:0051049,GO:0051128,GO:0051179,GO:0051234,GO:0051641,GO:0051649,GO:0060429,GO:0060627,GO:0060810,GO:0060811,GO:0061024,GO:0061025,GO:0061245,GO:0065007,GO:0065008,GO:0070727,GO:0071840,GO:0071944,GO:1901981 | Zinc finger, FYVE domain containing 20 |
| comp213922_c0_seq1.p1 | | Kelch motif |
| comp217430_c1_seq2.p1 | GO:0003008,GO:0003009,GO:0003012,GO:0003674,GO:0005575,GO:0005622,GO:0005623,GO:0005737,GO:0005886,GO:0005887,GO:0006928,GO:0006936,GO:0006937,GO:0006941,GO:0008150,GO:0009987,GO:0010876,GO:0010959,GO:0015459,GO:0016020,GO:0016021,GO:0016247,GO:0016477,GO:0019915,GO:0031224,GO:0031226,GO:0032501,GO:0032879,GO:0033036,GO:0040011,GO:0043226,GO:0043228,GO:0043229,GO:0043232,GO:0043266,GO:0043269,GO:0043292,GO:0044057,GO:0044422,GO:0044424,GO:0044425,GO:0044444,GO:0044449,GO:0044459,GO:0044464,GO:0044699,GO:0044707,GO:0044763,GO:0048870,GO:0050789,GO:0050879,GO:0050881,GO:0051049,GO:0051179,GO:0051239,GO:0051674,GO:0055120,GO:0065007,GO:0071944,GO:0090257 | unc-93 homolog A (C. elegans) |
| comp214398_c0_seq1.p1 | GO:0000910,GO:0000922,GO:0003674,GO:0005488,GO:0005515,GO:0005575,GO:0005622,GO:0005623,GO:0005634,GO:0005737,GO:0005768,GO:0005813,GO:0005815,GO:0005816,GO:0005819,GO:0005825,GO:0005856,GO:0006810,GO:0007049,GO:0008150,GO:0009987,GO:0010008,GO:0015630,GO:0016020,GO:0016192,GO:0016197,GO:0017016,GO:0017137,GO:0019899,GO:0022402,GO:0030306,GO:0030496,GO:0031090,GO:0031267,GO:0032153,GO:0032154,GO:0032155,GO:0032456,GO:0032991,GO:0042802,GO:0042803,GO:0043226,GO:0043227,GO:0043228,GO:0043229,GO:0043231,GO:0043232,GO:0043234,GO:0044422,GO:0044424,GO:0044430,GO:0044440,GO:0044444,GO:0044446,GO:0044450,GO:0044464,GO:0044699,GO:0044763,GO:0044765,GO:0046907,GO:0046983,GO:0051020,GO:0051179,GO:0051234,GO:0051301,GO:0051641,GO:0051649,GO:0055037,GO:0055038 | RAB11 family interacting protein 3 (class II) |
| comp215894_c0_seq5.p1 | | Inherit from biNOG: AT rich interactive domain 5B (MRF1-like) |

| | | |
|---|---|---|
| comp212499_c0_seq1.p1 | | Zinc finger protein |
| comp196191_c0_seq1.p1 | GO:0000323,GO:0000902,GO:0000904,GO:0001932,GO:0001934,GO:0001948,GO:0002162,GO:0003008,GO:0003674,GO:0005198,GO:0005200,GO:0005488,GO:0005509,GO:0005515,GO:0005539,GO:0005575,GO:0005576,GO:0005578,GO:0005604,GO:0005605,GO:0005615,GO:0005622,GO:0005623,GO:0005737,GO:0005764,GO:0005773,GO:0005775,GO:0005794,GO:0005796,GO:0005886,GO:0005975,GO:0006022,GO:0006023,GO:0006024,GO:0006026,GO:0006027,GO:0006029,GO:0006082,GO:0006140,GO:0006355,GO:0006357,GO:0006461,GO:0006790,GO:0006807,GO:0006928,GO:0006935,GO:0007009,GO:0007154,GO:0007165,GO:0007166,GO:0007167,GO:0007169,GO:0007171,GO:0007186,GO:0007213,GO:0007267,GO:0007268,GO:0007275,GO:0007399,GO:0007409,GO:0007411,GO:0007416,GO:0007517,GO:0007519,GO:0007528,GO:0008047,GO:0008104,GO:0008150,GO:0008152,GO:0008201,GO:0008582,GO:0009056,GO:0009057,GO:0009058,GO:0009059,GO:0009100,GO:0009118,GO:0009605,GO:0009653,GO:0009790,GO:0009792,GO:0009888,GO:0009889,GO:0009891,GO:0009893,GO:0009894,GO:0009966,GO:0009986,GO:0009987,GO:0010468,GO:0010469,GO:0010556,GO:0010557,GO:0010562,GO:0010604,GO:0010628,GO:0010646,GO:0010941,GO:0010942,GO:0014706,GO:0016020,GO:0016043,GO:0016044,GO:0016202,GO:0019207,GO:0019209,GO:0019219,GO:0019220,GO:0019222,GO:0019226,GO:0019538,GO:0019887,GO:0021700,GO:0022008,GO:0022607,GO:0023051,GO:0023052,GO:0030030,GO:0030154,GO:0030182,GO:0030203,GO:0030204,GO:0030234,GO:0030295,GO:0030296,GO:0030297,GO:0030545,GO:0030546,GO:0030548,GO:0030811,GO:0031012,GO:0031175,GO:0031323,GO:0031325,GO:0031326,GO:0031328,GO:0031329,GO:0031344,GO:0031346,GO:0031399,GO:0031401,GO:0031406,GO:0031974,GO:0032147,GO:0032268,GO:0032270,GO:0032314,GO:0032318,GO:0032319,GO:0032320,GO:0032321,GO:0032501,GO:0032502,GO:0032989,GO:0032990,GO:0033036,GO:0033121,GO:0033124,GO:0033674,GO:0033691,GO:0034613,GO:0034622,GO:0035020,GO:0035023,GO:0035374,GO:0035637,GO:0040008,GO:0040011,GO:0042221,GO:0042325,GO:0042327,GO:0042330,GO:0042551,GO:0042692,GO:0042981,GO:0043065,GO:0043067,GO:0043068,GO:0043085,GO:0043087,GO:0043112,GO:0043113,GO:0043167,GO:0043168,GO:0043169,GO:0043170,GO:0043202,GO:0043226,GO:0043227,GO:0043229,GO:0043231,GO:0043233,GO:0043236,GO:0043394,GO:0043395,GO:0043436,GO:0043523,GO:0043525,GO:0043547,GO:0043549,GO:0043623,GO:0043933,GO:0044085,GO:0044087,GO:0044089,GO:0044093,GO:0044237,GO:0044238,GO:0044260,GO:0044281,GO:0044420,GO:0044421,GO:0044422,GO:0044424,GO:0044431,GO:0044437,GO:0044444,GO:0044446,GO:0044464,GO:0044699,GO:0044700,GO:0044707,GO:0044710,GO:0044763,GO:0044767,GO:0045161,GO:0045162,GO:0045202,GO:0045213,GO:0045595,GO:0045859,GO:0045860,GO:0045887,GO:0045893,GO:0045927,GO:0045935,GO:0045937,GO:0045944,GO:0046578,GO:0046872,GO:0048468,GO:0048469,GO:0048513,GO:0048518,GO:0048522,GO:0048583,GO:0048634,GO:0048638,GO:0048639,GO:0048641,GO:0048666,GO:0048667,GO:0048699,GO:0048731,GO:0048741,GO:0048742,GO:0048747,GO:0048812,GO:0048856,GO:0048858,GO:0048869,GO:0050654,GO:0050730,GO:0050731,GO:0050789,GO:0050790,GO:0050793,GO:0050794,GO:0050803,GO:0050807,GO:0050808,GO:0050840,GO:0050877,GO:0050896,GO:0051056,GO:0051094,GO:0051128,GO:0051130,GO:0051146,GO:0051147,GO:0051153,GO:0051171,GO:0051173,GO:0051174,GO:0051179,GO:0051239,GO:0051246,GO:0051247,GO:0051252,GO:0051254,GO:0051336,GO:0051338,GO:0051345,GO:0051347,GO:0051489,GO:0051491,GO:0051641,GO:0051716,GO:0051960,GO:0051962,GO:0051963,GO:0051965,GO:0055001,GO:0055002,GO:0060255,GO:0060284,GO:0060491,GO:0060537,GO:0060538,GO:0061024,GO:0061061,GO:0061097,GO:0061098,GO:0061564,GO:0065003,GO:0065007,GO:0065008,GO:0065009,GO:0070013,GO:0070271,GO:0070727,GO:0071704,GO:0071822,GO:0071840,GO:0071944,GO:0072657,GO:0080090,GO:0097367,GO:0097485,GO:1900542,GO:1901135,GO:1901136,GO:1901137,GO:1901214,GO:1901216,GO:1901564,GO:1901565,GO:1901566,GO:1901575,GO:1901576,GO:1901681,GO:1901861,GO:2000026,GO:2000112,GO:2000273,GO:2000539,GO:2000541,GO:2001141 | Kazal-type serine protease inhibitor domain |
| comp216630_c0_seq7.p1 | GO:0005575,GO:0016020 | leucine rich repeat and Ig domain containing 4 |

| | | |
|---|---|---|
| comp218016_c0_seq2.p1 | GO:0000003,GO:0000166,GO:0001882,GO:0001883,GO:0003674,GO:0003824,GO:0005488,GO:0005515,GO:0005524,GO:0005575,GO:0005622,GO:0005623,GO:0005634,GO:0005737,GO:0005777,GO:0005778,GO:0005829,GO:0006139,GO:0006152,GO:0006163,GO:0006195,GO:0006200,GO:0006605,GO:0006625,GO:0006725,GO:0006753,GO:0006793,GO:0006796,GO:0006807,GO:0006810,GO:0006886,GO:0006996,GO:0007017,GO:0007031,GO:0007275,GO:0007276,GO:0007283,GO:0007399,GO:0007417,GO:0007422,GO:0008104,GO:0008150,GO:0008152,GO:0009056,GO:0009116,GO:0009117,GO:0009119,GO:0009123,GO:0009125,GO:0009126,GO:0009128,GO:0009141,GO:0009143,GO:0009144,GO:0009146,GO:0009150,GO:0009154,GO:0009158,GO:0009161,GO:0009164,GO:0009166,GO:0009167,GO:0009169,GO:0009199,GO:0009203,GO:0009205,GO:0009207,GO:0009259,GO:0009261,GO:0009987,GO:0010001,GO:0015031,GO:0016020,GO:0016043,GO:0016462,GO:0016482,GO:0016558,GO:0016787,GO:0016817,GO:0016818,GO:0016887,GO:0017038,GO:0017076,GO:0017111,GO:0019439,GO:0019637,GO:0019693,GO:0019953,GO:0021782,GO:0022008,GO:0022412,GO:0022414,GO:0030154,GO:0030554,GO:0031090,GO:0031903,GO:0032403,GO:0032501,GO:0032502,GO:0032504,GO:0032549,GO:0032550,GO:0032553,GO:0032555,GO:0032559,GO:0033036,GO:0033365,GO:0034613,GO:0034641,GO:0034655,GO:0035639,GO:0036094,GO:0040008,GO:0040009,GO:0040010,GO:0042063,GO:0042278,GO:0042454,GO:0042579,GO:0042623,GO:0043167,GO:0043168,GO:0043226,GO:0043227,GO:0043229,GO:0043231,GO:0043574,GO:0044237,GO:0044238,GO:0044248,GO:0044270,GO:0044281,GO:0044422,GO:0044424,GO:0044438,GO:0044439,GO:0044444,GO:0044446,GO:0044464,GO:0044699,GO:0044702,GO:0044707,GO:0044710,GO:0044743,GO:0044763,GO:0044765,GO:0044767,GO:0045184,GO:0045927,GO:0046034,GO:0046128,GO:0046130,GO:0046434,GO:0046483,GO:0046700,GO:0046907,GO:0048137,GO:0048232,GO:0048468,GO:0048518,GO:0048609,GO:0048610,GO:0048731,GO:0048856,GO:0048869,GO:0050789,GO:0051179,GO:0051234,GO:0051301,GO:0051640,GO:0051641,GO:0051649,GO:0055085,GO:0055086,GO:0060151,GO:0060152,GO:0065002,GO:0065007,GO:0070727,GO:0071702,GO:0071704,GO:0071806,GO:0071840,GO:0072521,GO:0072523,GO:0072594,GO:0072662,GO:0072663,GO:0097159,GO:1901135,GO:1901136,GO:1901265,GO:1901292,GO:1901360,GO:1901361,GO:1901363,GO:1901564,GO:1901565,GO:1901575,GO:1901657,GO:1901658 | Peroxisomal biogenesis factor 1 |
| comp202032_c0_seq1.p1 | GO:0003674,GO:0003824,GO:0004672,GO:0004674,GO:0004683,GO:0005575,GO:0006464,GO:0006468,GO:0006793,GO:0006796,GO:0008150,GO:0008152,GO:0009987,GO:0016020,GO:0016301,GO:0016310,GO:0016740,GO:0016772,GO:0016773,GO:0019538,GO:0036211,GO:0043170,GO:0043412,GO:0044237,GO:0044238,GO:0044260,GO:0044267,GO:0071704 | calcium-dependent protein kinase |

| | | |
|---|---|---|
| comp198423_c0_seq1.p1 | GO:0003674,GO:0003779,GO:0005198,GO:0005488,GO:0005515,GO:0005575,GO:0005622,GO:0005623,GO:0005634,GO:0005635,GO:0005640,GO:0005654,GO:0005737,GO:0005739,GO:0005783,GO:0005789,GO:0005794,GO:0005819,GO:0005856,GO:0005886,GO:0005912,GO:0005924,GO:0005925,GO:0006928,GO:0006996,GO:0006997,GO:0007010,GO:0007030,GO:0007097,GO:0007275,GO:0007399,GO:0007417,GO:0007420,GO:0008092,GO:0008104,GO:0008150,GO:0009314,GO:0009416,GO:0009628,GO:0009987,GO:0010769,GO:0010975,GO:0012505,GO:0015630,GO:0016020,GO:0016021,GO:0016043,GO:0016528,GO:0016529,GO:0019867,GO:0019899,GO:0022008,GO:0022603,GO:0022604,GO:0030016,GO:0030017,GO:0030018,GO:0030027,GO:0030029,GO:0030048,GO:0030054,GO:0030055,GO:0030100,GO:0030154,GO:0030175,GO:0030334,GO:0030335,GO:0030425,GO:0030496,GO:0031022,GO:0031090,GO:0031224,GO:0031252,GO:0031253,GO:0031256,GO:0031258,GO:0031344,GO:0031527,GO:0031674,GO:0031965,GO:0031967,GO:0031968,GO:0031974,GO:0031975,GO:0031981,GO:0032501,GO:0032502,GO:0032507,GO:0032879,GO:0032991,GO:0033017,GO:0033036,GO:0033365,GO:0034504,GO:0034613,GO:0034992,GO:0034993,GO:0040012,GO:0040017,GO:0040023,GO:0042175,GO:0042692,GO:0042802,GO:0042803,GO:0042995,GO:0043005,GO:0043197,GO:0043226,GO:0043227,GO:0043228,GO:0043229,GO:0043231,GO:0043232,GO:0043233,GO:0043234,GO:0043292,GO:0043578,GO:0044309,GO:0044422,GO:0044424,GO:0044425,GO:0044428,GO:0044429,GO:0044430,GO:0044432,GO:0044444,GO:0044446,GO:0044449,GO:0044456,GO:0044459,GO:0044463,GO:0044464,GO:0044699,GO:0044707,GO:0044763,GO:0044767,GO:0045185,GO:0045202,GO:0045211,GO:0045595,GO:0045664,GO:0045807,GO:0046983,GO:0048259,GO:0048260,GO:0048471,GO:0048513,GO:0048518,GO:0048522,GO:0048699,GO:0048731,GO:0048814,GO:0048856,GO:0048869,GO:0050767,GO:0050773,GO:0050789,GO:0050793,GO:0050794,GO:0050896,GO:0051049,GO:0051050,GO:0051128,GO:0051130,GO:0051179,GO:0051234,GO:0051235,GO:0051239,GO:0051270,GO:0051272,GO:0051457,GO:0051640,GO:0051641,GO:0051642,GO:0051647,GO:0051649,GO:0051651,GO:0051656,GO:0051960,GO:0060284,GO:0060627,GO:0061061,GO:0065007,GO:0065008,GO:0070013,GO:0070161,GO:0070727,GO:0071840,GO:0071944,GO:0072595,GO:0090286,GO:0090292,GO:0097060,GO:0097458,GO:2000026,GO:2000145,GO:2000147 | spectrin repeat containing, nuclear envelope 1 |

| | | |
|---|---|---|
| comp215789_c0_seq2.p1 | GO:0000003,GO:0000375,GO:0000377,GO:0000398,GO:0000578,GO:0003002,GO:0003006,GO:0003674,GO:0003676,GO:0003723,GO:0003727,GO:0003729,GO:0005488,GO:0005515,GO:0005575,GO:0005622,GO:0005623,GO:0005634,GO:0005654,GO:0005681,GO:0005737,GO:0005856,GO:0006139,GO:0006396,GO:0006397,GO:0006403,GO:0006417,GO:0006725,GO:0006807,GO:0006950,GO:0006996,GO:0007028,GO:0007275,GO:0007276,GO:0007281,GO:0007292,GO:0007308,GO:0007309,GO:0007310,GO:0007311,GO:0007314,GO:0007315,GO:0007316,GO:0007350,GO:0007351,GO:0007389,GO:0008150,GO:0008152,GO:0008187,GO:0008266,GO:0008298,GO:0008358,GO:0008380,GO:0008595,GO:0009611,GO:0009790,GO:0009792,GO:0009798,GO:0009880,GO:0009889,GO:0009948,GO:0009950,GO:0009952,GO:0009953,GO:0009966,GO:0009967,GO:0009987,GO:0009994,GO:0010467,GO:0010468,GO:0010556,GO:0010608,GO:0010627,GO:0010646,GO:0010647,GO:0010740,GO:0015629,GO:0016043,GO:0016070,GO:0016071,GO:0019094,GO:0019219,GO:0019222,GO:0019953,GO:0021700,GO:0022412,GO:0022414,GO:0022607,GO:0023051,GO:0023056,GO:0030154,GO:0030529,GO:0031323,GO:0031326,GO:0031974,GO:0031981,GO:0032268,GO:0032501,GO:0032502,GO:0032504,GO:0032991,GO:0033036,GO:0034641,GO:0035282,GO:0040008,GO:0040014,GO:0040018,GO:0042060,GO:0043170,GO:0043226,GO:0043227,GO:0043228,GO:0043229,GO:0043231,GO:0043232,GO:0043233,GO:0043234,GO:0043484,GO:0044085,GO:0044237,GO:0044238,GO:0044260,GO:0044422,GO:0044424,GO:0044428,GO:0044446,GO:0044464,GO:0044699,GO:0044702,GO:0044707,GO:0044763,GO:0044767,GO:0045451,GO:0045927,GO:0046425,GO:0046427,GO:0046483,GO:0048468,GO:0048469,GO:0048477,GO:0048518,GO:0048522,GO:0048583,GO:0048584,GO:0048599,GO:0048609,GO:0048610,GO:0048856,GO:0048869,GO:0050789,GO:0050794,GO:0050896,GO:0051171,GO:0051179,GO:0051239,GO:0051240,GO:0051246,GO:0051252,GO:0051276,GO:0051641,GO:0060255,GO:0060810,GO:0060811,GO:0065007,GO:0070013,GO:0070727,GO:0071013,GO:0071704,GO:0071840,GO:0080090,GO:0090304,GO:0097159,GO:1901360,GO:1901363,GO:2000112 | heterogeneous nuclear ribonucleoprotein |
| comp212189_c1_seq16.p1 | | Sodium/hydrogen exchanger family |

| | | |
|---|---|---|
| comp218495_c1_seq2.p1 | GO:0000003,GO:0000070,GO:0000226,GO:0000228,GO:0000236,GO:0000278,GO:0000280,GO:0000724,GO:0000725,GO:0000775,GO:0000785,GO:0000790,GO:0000793,GO:0000794,GO:0000795,GO:0000798,GO:0000800,GO:0000819,GO:0000910,GO:0000920,GO:0000922,GO:0001736,GO:0001737,GO:0001738,GO:0002009,GO:0002165,GO:0003006,GO:0003674,GO:0003682,GO:0003774,GO:0003777,GO:0003824,GO:0004672,GO:0004674,GO:0005488,GO:0005515,GO:0005575,GO:0005576,GO:0005578,GO:0005604,GO:0005622,GO:0005623,GO:0005634,GO:0005654,GO:0005694,GO:0005737,GO:0005819,GO:0005856,GO:0006139,GO:0006259,GO:0006275,GO:0006281,GO:0006302,GO:0006310,GO:0006323,GO:0006464,GO:0006468,GO:0006725,GO:0006793,GO:0006796,GO:0006807,GO:0006810,GO:0006897,GO:0006898,GO:0006950,GO:0006974,GO:0006996,GO:0007010,GO:0007017,GO:0007049,GO:0007051,GO:0007052,GO:0007059,GO:0007062,GO:0007064,GO:0007067,GO:0007091,GO:0007109,GO:0007126,GO:0007154,GO:0007164,GO:0007165,GO:0007267,GO:0007275,GO:0007399,GO:0007444,GO:0007472,GO:0007476,GO:0007548,GO:0007552,GO:0007560,GO:0007568,GO:0008150,GO:0008152,GO:0008156,GO:0008278,GO:0008340,GO:0008544,GO:0009653,GO:0009790,GO:0009791,GO:0009792,GO:0009886,GO:0009887,GO:0009888,GO:0009889,GO:0009890,GO:0009892,GO:0009913,GO:0009987,GO:0010259,GO:0010556,GO:0010558,GO:0010564,GO:0010605,GO:0010948,GO:0015630,GO:0016043,GO:0016192,GO:0016301,GO:0016310,GO:0016363,GO:0016462,GO:0016740,GO:0016772,GO:0016773,GO:0016787,GO:0016817,GO:0016818,GO:0017111,GO:0019219,GO:0019222,GO:0019538,GO:0019827,GO:0022008,GO:0022402,GO:0022403,GO:0022414,GO:0023052,GO:0030030,GO:0030154,GO:0030182,GO:0030261,GO:0030855,GO:0030893,GO:0031012,GO:0031323,GO:0031324,GO:0031326,GO:0031327,GO:0031974,GO:0031981,GO:0032403,GO:0032501,GO:0032502,GO:0032506,GO:0032875,GO:0032876,GO:0032991,GO:0033554,GO:0034399,GO:0034641,GO:0034991,GO:0035107,GO:0035114,GO:0035120,GO:0035220,GO:0035315,GO:0035316,GO:0035317,GO:0036033,GO:0036211,GO:0040002,GO:0040008,GO:0040009,GO:0040010,GO:0040011,GO:0040035,GO:0042335,GO:0043170,GO:0043226,GO:0043227,GO:0043228,GO:0043229,GO:0043231,GO:0043232,GO:0043233,GO:0043234,GO:0043412,GO:0043588,GO:0043933,GO:0044237,GO:0044238,GO:0044260,GO:0044267,GO:0044420,GO:0044421,GO:0044422,GO:0044424,GO:0044427,GO:0044428,GO:0044430,GO:0044446,GO:0044454,GO:0044464,GO:0044699,GO:0044700,GO:0044707,GO:0044763,GO:0044767,GO:0044770,GO:0044772,GO:0044784,GO:0045165,GO:0045168,GO:0045502,GO:0045595,GO:0045596,GO:0045927,GO:0045934,GO:0046331,GO:0046483,GO:0048285,GO:0048468,GO:0048513,GO:0048518,GO:0048519,GO:0048523,GO:0048563,GO:0048569,GO:0048608,GO:0048610,GO:0048699,GO:0048707,GO:0048729,GO:0048731,GO:0048736,GO:0048737,GO:0048806,GO:0048856,GO:0048863,GO:0048864,GO:0048869,GO:0050789,GO:0050793,GO:0050794,GO:0050896,GO:0051052,GO:0051053,GO:0051093,GO:0051171,GO:0051172,GO:0051179,GO:0051234,GO:0051276,GO:0051301,GO:0051321,GO:0051716,GO:0051726,GO:0060255,GO:0060429,GO:0061458,GO:0065007,GO:0070013,GO:0071103,GO:0071704,GO:0071822,GO:0071840,GO:0080090,GO:0090304,GO:0090329,GO:1901360,GO:2000104,GO:2000112,GO:2000113 | Structural maintenance of chromosomes |
| comp218495_c1_seq2.p2 | GO:0000070,GO:0000278,GO:0000280,GO:0000819,GO:0006996,GO:0007049,GO:0007059,GO:0007062,GO:0007064,GO:0007067,GO:0008150,GO:0009987,GO:0016043,GO:0022402,GO:0044699,GO:0044763,GO:0048285,GO:0051276,GO:0071840 | structural maintenance of chromosomes 3 |
| comp216851_c0_seq8.p1 | | conserved hypothetical protein |
| comp218923_c0_seq4.p1 | GO:0000902,GO:0003341,GO:0005575,GO:0005622,GO:0005623,GO:0005929,GO:0006928,GO:0007017,GO:0007018,GO:0008150,GO:0009653,GO:0009987,GO:0016043,GO:0030030,GO:0032502,GO:0032989,GO:0032990,GO:0042995,GO:0043226,GO:0043227,GO:0043229,GO:0043231,GO:0044424,GO:0044464,GO:0044699,GO:0044763,GO:0044767,GO:0048856,GO:0048858,GO:0048869,GO:0060271,GO:0071840 | primary ciliary dyskinesia protein |

| | | |
|---|---|---|
| comp189912_c0_seq1.p1 | GO:0003674,GO:0004857,GO:0004866,GO:0005575,GO:0005576,GO:0008150,GO:0010466,GO:0010951,GO:0019222,GO:0030234,GO:0030414,GO:0043086,GO:0044092,GO:0050789,GO:0050790,GO:0051336,GO:0051346,GO:0052547,GO:0052548,GO:0061134,GO:0061135,GO:0065007,GO:0065009 | Kazal-type serine protease inhibitor domain |
| comp148501_c0_seq2.p1 | | ankyrin repeat domain 11 |
| comp148501_c0_seq2.p2 | GO:0000132,GO:0000226,GO:0000278,GO:0000902,GO:0000904,GO:0001539,GO:0001654,GO:0001655,GO:0001701,GO:0001754,GO:0001822,GO:0001894,GO:0001895,GO:0001944,GO:0002064,GO:0003002,GO:0003007,GO:0003008,GO:0003205,GO:0003206,GO:0003279,GO:0003382,GO:0005575,GO:0005622,GO:0005623,GO:0005737,GO:0005813,GO:0005814,GO:0005815,GO:0005856,GO:0005929,GO:0005930,GO:0005932,GO:0006810,GO:0006928,GO:0006996,GO:0007010,GO:0007017,GO:0007018,GO:0007049,GO:0007154,GO:0007163,GO:0007165,GO:0007166,GO:0007224,GO:0007275,GO:0007368,GO:0007369,GO:0007389,GO:0007399,GO:0007417,GO:0007420,GO:0007423,GO:0007507,GO:0007600,GO:0007605,GO:0008104,GO:0008150,GO:0008152,GO:0008589,GO:0009314,GO:0009416,GO:0009628,GO:0009653,GO:0009790,GO:0009792,GO:0009799,GO:0009855,GO:0009887,GO:0009888,GO:0009893,GO:0009952,GO:0009953,GO:0009966,GO:0009967,GO:0009987,GO:0010467,GO:0010604,GO:0010646,GO:0010647,GO:0010927,GO:0010941,GO:0010970,GO:0015630,GO:0016043,GO:0016485,GO:0019222,GO:0019538,GO:0021510,GO:0021511,GO:0021513,GO:0021537,GO:0022008,GO:0022402,GO:0022603,GO:0022607,GO:0023051,GO:0023052,GO:0023056,GO:0030010,GO:0030030,GO:0030031,GO:0030154,GO:0030162,GO:0030182,GO:0030323,GO:0030324,GO:0030326,GO:0030705,GO:0030855,GO:0030900,GO:0030990,GO:0030992,GO:0031323,GO:0031325,GO:0031503,GO:0031512,GO:0031513,GO:0031514,GO:0032268,GO:0032270,GO:0032391,GO:0032501,GO:0032502,GO:0032838,GO:0032989,GO:0032990,GO:0032991,GO:0033036,GO:0035058,GO:0035085,GO:0035107,GO:0035108,GO:0035113,GO:0035295,GO:0040001,GO:0040011,GO:0042073,GO:0042384,GO:0042461,GO:0042462,GO:0042481,GO:0042487,GO:0042592,GO:0042733,GO:0042981,GO:0042995,GO:0043009,GO:0043067,GO:0043170,GO:0043226,GO:0043227,GO:0043228,GO:0043229,GO:0043231,GO:0043232,GO:0043234,GO:0044085,GO:0044238,GO:0044422,GO:0044424,GO:0044430,GO:0044441,GO:0044444,GO:0044446,GO:0044447,GO:0044450,GO:0044463,GO:0044464,GO:0044699,GO:0044700,GO:0044707,GO:0044763,GO:0044765,GO:0044767,GO:0044782,GO:0045177,GO:0045494,GO:0045862,GO:0045880,GO:0046530,GO:0046907,GO:0048468,GO:0048513,GO:0048518,GO:0048522,GO:0048568,GO:0048583,GO:0048584,GO:0048592,GO:0048598,GO:0048646,GO:0048666,GO:0048699,GO:0048731,GO:0048736,GO:0048853,GO:0048854,GO:0048856,GO:0048858,GO:0048869,GO:0048870,GO:0048871,GO:0048872,GO:0048873,GO:0050789,GO:0050793,GO:0050794,GO:0050877,GO:0050896,GO:0050954,GO:0051179,GO:0051234,GO:0051239,GO:0051246,GO:0051247,GO:0051293,GO:0051294,GO:0051604,GO:0051640,GO:0051641,GO:0051649,GO:0051653,GO:0051656,GO:0051674,GO:0051716,GO:0060021,GO:0060173,GO:0060249,GO:0060255,GO:0060271,GO:0060411,GO:0060426,GO:0060429,GO:0060541,GO:0060785,GO:0061326,GO:0065007,GO:0065008,GO:0070925,GO:0071214,GO:0071478,GO:0071482,GO:0071704,GO:0071840,GO:0072001,GO:0072006,GO:0072009,GO:0072014,GO:0072019,GO:0072073,GO:0072080,GO:0072358,GO:0072359,GO:0072372,GO:0080090,GO:0097014,GO:2000026,GO:2000027 | Intraflagellar transport 88 homolog (Chlamydomonas) |

| | | |
|---|---|---|
| comp148501_c0_seq2.p4 | GO:0000003,GO:0000723,GO:0000781,GO:0001516,GO:0002039,GO:0003008,GO:0003674,GO:0003720,GO:0003824,GO:0003964,GO:0005488,GO:0005515,GO:0005575,GO:0005622,GO:0005623,GO:0005634,GO:0005635,GO:0005694,GO:0005697,GO:0005737,GO:0005783,GO:0005829,GO:0005856,GO:0005884,GO:0006082,GO:0006139,GO:0006259,GO:0006457,GO:0006629,GO:0006631,GO:0006633,GO:0006636,GO:0006690,GO:0006692,GO:0006693,GO:0006725,GO:0006807,GO:0006996,GO:0007030,GO:0007154,GO:0007165,GO:0007275,GO:0007600,GO:0008150,GO:0008152,GO:0008610,GO:0009058,GO:0009790,GO:0009792,GO:0009893,GO:0009987,GO:0010468,GO:0010604,GO:0010628,GO:0010941,GO:0012505,GO:0015629,GO:0016020,GO:0016043,GO:0016053,GO:0016740,GO:0016772,GO:0016779,GO:0016853,GO:0016860,GO:0019222,GO:0019233,GO:0019538,GO:0019752,GO:0023052,GO:0030016,GO:0030529,GO:0031072,GO:0031090,GO:0031965,GO:0031967,GO:0031975,GO:0032200,GO:0032501,GO:0032502,GO:0032787,GO:0032991,GO:0033559,GO:0034061,GO:0034641,GO:0042026,GO:0042592,GO:0042995,GO:0043005,GO:0043025,GO:0043170,GO:0043226,GO:0043227,GO:0043228,GO:0043229,GO:0043231,GO:0043232,GO:0043234,GO:0043292,GO:0043436,GO:0044237,GO:0044238,GO:0044249,GO:0044255,GO:0044260,GO:0044267,GO:0044281,GO:0044283,GO:0044297,GO:0044422,GO:0044424,GO:0044427,GO:0044428,GO:0044430,GO:0044444,GO:0044446,GO:0044464,GO:0044699,GO:0044700,GO:0044707,GO:0044710,GO:0044711,GO:0044763,GO:0044767,GO:0046394,GO:0046456,GO:0046457,GO:0046483,GO:0048471,GO:0048518,GO:0048519,GO:0048523,GO:0048856,GO:0050220,GO:0050789,GO:0050794,GO:0050877,GO:0050896,GO:0051082,GO:0051276,GO:0051716,GO:0051879,GO:0060249,GO:0060255,GO:0060548,GO:0061077,GO:0065007,GO:0065008,GO:0070389,GO:0071704,GO:0071840,GO:0072330,GO:0090304,GO:0097458,GO:1901360,GO:1901568,GO:1901570,GO:1901576 | Prostaglandin E synthase 3 (Cytosolic) |
| comp202720_c1_seq3.p1 | GO:0000003,GO:0003008,GO:0003012,GO:0003674,GO:0005198,GO:0005215,GO:0005246,GO:0005277,GO:0005326,GO:0005488,GO:0005515,GO:0005575,GO:0005622,GO:0005623,GO:0005856,GO:0005886,GO:0006810,GO:0006811,GO:0006812,GO:0006836,GO:0006936,GO:0006996,GO:0007010,GO:0007154,GO:0007267,GO:0007268,GO:0007275,GO:0007517,GO:0007519,GO:0007528,GO:0008092,GO:0008150,GO:0008307,GO:0008324,GO:0008582,GO:0009653,GO:0009888,GO:0009987,GO:0010927,GO:0014706,GO:0015075,GO:0015870,GO:0016010,GO:0016013,GO:0016020,GO:0016043,GO:0016202,GO:0016247,GO:0019226,GO:0019725,GO:0022607,GO:0022857,GO:0022891,GO:0022892,GO:0022898,GO:0023052,GO:0030029,GO:0030036,GO:0030054,GO:0030154,GO:0030239,GO:0031032,GO:0032409,GO:0032410,GO:0032412,GO:0032413,GO:0032501,GO:0032502,GO:0032879,GO:0032989,GO:0032991,GO:0034220,GO:0034762,GO:0034765,GO:0035637,GO:0040008,GO:0040009,GO:0040010,GO:0040012,GO:0040013,GO:0040014,GO:0040017,GO:0040018,GO:0042383,GO:0042592,GO:0042692,GO:0043058,GO:0043059,GO:0043226,GO:0043228,GO:0043229,GO:0043232,GO:0043234,GO:0043269,GO:0043900,GO:0044085,GO:0044087,GO:0044092,GO:0044424,GO:0044425,GO:0044459,GO:0044464,GO:0044699,GO:0044700,GO:0044707,GO:0044763,GO:0044765,GO:0044767,GO:0045202,GO:0045214,GO:0045595,GO:0045927,GO:0046662,GO:0046716,GO:0048468,GO:0048513,GO:0048518,GO:0048519,GO:0048583,GO:0048634,GO:0048638,GO:0048641,GO:0048646,GO:0048731,GO:0048741,GO:0048742,GO:0048747,GO:0048856,GO:0048869,GO:0050789,GO:0050793,GO:0050794,GO:0050795,GO:0050803,GO:0050807,GO:0050808,GO:0050877,GO:0051049,GO:0051051,GO:0051128,GO:0051146,GO:0051147,GO:0051153,GO:0051179,GO:0051234,GO:0051239,GO:0051240,GO:0051960,GO:0051963,GO:0055001,GO:0055002,GO:0055085,GO:0060249,GO:0060284,GO:0060537,GO:0060538,GO:0061061,GO:0065007,GO:0065008,GO:0065009,GO:0070925,GO:0071702,GO:0071705,GO:0071840,GO:0071944,GO:1901374,GO:1901375,GO:1901861,GO:2000026,GO:2000241 | Syntrophin, beta 1 (Dystrophin-associated protein A1, 59kDa, basic component 1) |

| | | |
|---|---|---|
| comp221021_c0_seq3.p1 | GO:0001932,GO:0001934,GO:0003674,GO:0005575,GO:0005912,GO:0008047,GO:0008150,GO:0009893,GO:0010562,GO:0010604,GO:0019207,GO:0019209,GO:0019220,GO:0019222,GO:0019887,GO:0030054,GO:0030234,GO:0030295,GO:0030296,GO:0031323,GO:0031325,GO:0031399,GO:0031401,GO:0032268,GO:0032270,GO:0033674,GO:0042325,GO:0042327,GO:0043085,GO:0043549,GO:0044093,GO:0045859,GO:0045860,GO:0045937,GO:0048518,GO:0048522,GO:0050730,GO:0050731,GO:0050789,GO:0050790,GO:0050794,GO:0051174,GO:0051246,GO:0051247,GO:0051338,GO:0051347,GO:0060255,GO:0061097,GO:0061098,GO:0065007,GO:0065009,GO:0070161,GO:0080090 | apoptosis stimulating of P53 |
| comp220202_c0_seq4.p1 | GO:0000003,GO:0000182,GO:0000323,GO:0000902,GO:0000904,GO:0000975,GO:0001016,GO:0001025,GO:0001030,GO:0001031,GO:0001032,GO:0001067,GO:0001101,GO:0001156,GO:0001932,GO:0001934,GO:0001936,GO:0001938,GO:0002119,GO:0002164,GO:0003006,GO:0003008,GO:0003674,GO:0003676,GO:0003677,GO:0005488,GO:0005515,GO:0005575,GO:0005622,GO:0005623,GO:0005737,GO:0005764,GO:0005773,GO:0005829,GO:0006355,GO:0006359,GO:0006928,GO:0006935,GO:0006950,GO:0006974,GO:0007049,GO:0007050,GO:0007154,GO:0007165,GO:0007166,GO:0007167,GO:0007169,GO:0007267,GO:0007268,GO:0007275,GO:0007399,GO:0007409,GO:0007411,GO:0007548,GO:0007568,GO:0008134,GO:0008150,GO:0008284,GO:0008286,GO:0008340,GO:0008361,GO:0008406,GO:0008582,GO:0009605,GO:0009653,GO:0009719,GO:0009725,GO:0009791,GO:0009889,GO:0009891,GO:0009893,GO:0009966,GO:0009967,GO:0009987,GO:0009991,GO:0010033,GO:0010243,GO:0010259,GO:0010468,GO:0010556,GO:0010557,GO:0010562,GO:0010604,GO:0010628,GO:0010646,GO:0010647,GO:0016043,GO:0016049,GO:0016202,GO:0019219,GO:0019220,GO:0019222,GO:0019226,GO:0022008,GO:0022402,GO:0022414,GO:0023051,GO:0023052,GO:0023056,GO:0030030,GO:0030154,GO:0030182,GO:0031175,GO:0031323,GO:0031325,GO:0031326,GO:0031328,GO:0031399,GO:0031401,GO:0031667,GO:0031668,GO:0031669,GO:0031929,GO:0031931,GO:0032006,GO:0032008,GO:0032268,GO:0032270,GO:0032501,GO:0032502,GO:0032535,GO:0032868,GO:0032869,GO:0032870,GO:0032989,GO:0032990,GO:0032991,GO:0033554,GO:0033674,GO:0035556,GO:0035637,GO:0040007,GO:0040008,GO:0040011,GO:0040024,GO:0042127,GO:0042221,GO:0042325,GO:0042327,GO:0042330,GO:0043085,GO:0043200,GO:0043226,GO:0043227,GO:0043229,GO:0043231,GO:0043234,GO:0043434,GO:0043549,GO:0043565,GO:0044087,GO:0044093,GO:0044212,GO:0044424,GO:0044444,GO:0044464,GO:0044699,GO:0044700,GO:0044707,GO:0044763,GO:0044767,GO:0045137,GO:0045595,GO:0045786,GO:0045859,GO:0045860,GO:0045886,GO:0045893,GO:0045926,GO:0045935,GO:0045937,GO:0045945,GO:0048468,GO:0048513,GO:0048518,GO:0048519,GO:0048522,GO:0048523,GO:0048583,GO:0048584,GO:0048608,GO:0048634,GO:0048638,GO:0048640,GO:0048641,GO:0048666,GO:0048667,GO:0048699,GO:0048731,GO:0048742,GO:0048812,GO:0048856,GO:0048858,GO:0048869,GO:0050678,GO:0050679,GO:0050789,GO:0050790,GO:0050793,GO:0050794,GO:0050803,GO:0050807,GO:0050877,GO:0050896,GO:0051093,GO:0051128,GO:0051129,GO:0051147,GO:0051153,GO:0051171,GO:0051173,GO:0051174,GO:0051239,GO:0051246,GO:0051247,GO:0051252,GO:0051254,GO:0051338,GO:0051347,GO:0051716,GO:0051726,GO:0051960,GO:0051961,GO:0051963,GO:0051964,GO:0060255,GO:0060284,GO:0061458,GO:0061564,GO:0065007,GO:0065008,GO:0065009,GO:0070887,GO:0071229,GO:0071230,GO:0071310,GO:0071375,GO:0071417,GO:0071495,GO:0071496,GO:0071840,GO:0071889,GO:0071900,GO:0071902,GO:0080084,GO:0080090,GO:0090066,GO:0097159,GO:0097485,GO:1901363,GO:1901652,GO:1901653,GO:1901698,GO:1901699,GO:1901700,GO:1901701,GO:1901861,GO:2000026,GO:2000112,GO:2001141 | regulatory associated protein of MTOR, complex 1 |
| comp217857_c3_seq1.p1 | | ankyrin repeat domain 5 |
| comp212877_c0_seq1.p1 | GO:0005575,GO:0005622,GO:0005623,GO:0005929,GO:0006928,GO:0008150,GO:0009987,GO:0016020,GO:0030317,GO:0031514,GO:0040011,GO:0042995,GO:0043226,GO:0043227,GO:0043229,GO:0043231,GO:0044424,GO:0044464,GO:0044699,GO:0044763,GO:0048870,GO:0051179,GO:0051674 | solute carrier family 9, subfamily C (Na - transporting carboxylic acid decarboxylase), member 1 |

| | | |
|---|---|---|
| comp218046_c1_seq8.p1 | | sorbin and SH3 domain containing 3 |
| comp212358_c0_seq2.p1 | | CRAL_TRIO_N |
| comp204927_c0_seq2.p1 | GO:0003674,GO:0003824,GO:0004721,GO:0004725,GO:0004871,GO:0004872,GO:0004888,GO:0005001,GO:0005102,GO:0005488,GO:0005515,GO:0005575,GO:0005623,GO:0005886,GO:0005887,GO:0006464,GO:0006470,GO:0006793,GO:0006796,GO:0007154,GO:0007155,GO:0007157,GO:0007165,GO:0007166,GO:0007167,GO:0007185,GO:0007275,GO:0007399,GO:0007416,GO:0008150,GO:0008152,GO:0009987,GO:0010720,GO:0010769,GO:0010975,GO:0016020,GO:0016021,GO:0016043,GO:0016044,GO:0016311,GO:0016337,GO:0016787,GO:0016788,GO:0016791,GO:0019198,GO:0019538,GO:0022008,GO:0022603,GO:0022604,GO:0022607,GO:0022610,GO:0023052,GO:0030154,GO:0030182,GO:0031224,GO:0031226,GO:0031344,GO:0031346,GO:0032501,GO:0032502,GO:0035335,GO:0036211,GO:0038023,GO:0042578,GO:0043170,GO:0043412,GO:0044085,GO:0044091,GO:0044237,GO:0044238,GO:0044260,GO:0044267,GO:0044425,GO:0044459,GO:0044464,GO:0044699,GO:0044700,GO:0044707,GO:0044763,GO:0045595,GO:0045597,GO:0045664,GO:0048518,GO:0048522,GO:0048699,GO:0048731,GO:0048814,GO:0048856,GO:0048869,GO:0050767,GO:0050769,GO:0050773,GO:0050775,GO:0050789,GO:0050793,GO:0050794,GO:0050808,GO:0050839,GO:0050896,GO:0051094,GO:0051128,GO:0051130,GO:0051239,GO:0051716,GO:0051960,GO:0060089,GO:0060284,GO:0061024,GO:0065007,GO:0071704,GO:0071709,GO:0071840,GO:0071944,GO:0097090,GO:0097105,GO:2000026 | protein tyrosine phosphatase receptor type |
| comp214114_c1_seq1.p1 | GO:0000139,GO:0000226,GO:0000785,GO:0005575,GO:0005622,GO:0005623,GO:0005694,GO:0005737,GO:0005794,GO:0005795,GO:0005802,GO:0005813,GO:0005815,GO:0005819,GO:0005856,GO:0006996,GO:0007010,GO:0007017,GO:0007049,GO:0007098,GO:0008150,GO:0009987,GO:0012505,GO:0015630,GO:0016020,GO:0016043,GO:0022402,GO:0031023,GO:0031090,GO:0031984,GO:0031985,GO:0032580,GO:0032991,GO:0043226,GO:0043227,GO:0043228,GO:0043229,GO:0043231,GO:0043232,GO:0043234,GO:0044422,GO:0044424,GO:0044427,GO:0044430,GO:0044431,GO:0044444,GO:0044446,GO:0044464,GO:0044699,GO:0044763,GO:0048471,GO:0051297,GO:0071840 | Golgin A1 |
| comp212744_c3_seq11.p1 | GO:0000228,GO:0005575,GO:0005622,GO:0005623,GO:0005634,GO:0005694,GO:0006139,GO:0006259,GO:0006260,GO:0006261,GO:0006325,GO:0006338,GO:0006351,GO:0006355,GO:0006725,GO:0006807,GO:0006996,GO:0008150,GO:0008152,GO:0008623,GO:0009058,GO:0009059,GO:0009889,GO:0009987,GO:0010467,GO:0010468,GO:0010556,GO:0016043,GO:0016070,GO:0016568,GO:0016590,GO:0018130,GO:0019219,GO:0019222,GO:0019438,GO:0031010,GO:0031323,GO:0031326,GO:0031974,GO:0031981,GO:0032774,GO:0032991,GO:0034641,GO:0034645,GO:0034654,GO:0043170,GO:0043226,GO:0043227,GO:0043228,GO:0043229,GO:0043231,GO:0043232,GO:0043233,GO:0043234,GO:0044237,GO:0044238,GO:0044249,GO:0044260,GO:0044271,GO:0044422,GO:0044424,GO:0044428,GO:0044446,GO:0044464,GO:0044699,GO:0044763,GO:0046483,GO:0050789,GO:0050794,GO:0051171,GO:0051252,GO:0051276,GO:0060255,GO:0065007,GO:0070013,GO:0070603,GO:0071704,GO:0071840,GO:0080090,GO:0090304,GO:1901360,GO:1901362,GO:1901576,GO:2000112,GO:2001141 | Bromodomain adjacent to zinc finger domain, 1A |

| | | |
|---|---|---|
| comp218511_c0_seq10.p1 | GO:0000003,GO:0000070,GO:0000226,GO:0000228,GO:0000278,GO:0000280,GO:0000775,GO:0000785,GO:0000793,GO:0000796,GO:0000803,GO:0000819,GO:0002119,GO:0002164,GO:0003674,GO:0003676,GO:0003677,GO:0005488,GO:0005515,GO:0005575,GO:0005622,GO:0005623,GO:0005634,GO:0005694,GO:0005737,GO:0005856,GO:0005875,GO:0006139,GO:0006259,GO:0006323,GO:0006725,GO:0006807,GO:0006928,GO:0006996,GO:0007010,GO:0007017,GO:0007049,GO:0007051,GO:0007052,GO:0007059,GO:0007062,GO:0007067,GO:0007076,GO:0007126,GO:0007275,GO:0007399,GO:0007549,GO:0008150,GO:0008152,GO:0008278,GO:0009790,GO:0009791,GO:0009792,GO:0009987,GO:0010032,GO:0010468,GO:0015630,GO:0016043,GO:0016477,GO:0019222,GO:0022008,GO:0022402,GO:0030154,GO:0030261,GO:0031974,GO:0031981,GO:0032501,GO:0032502,GO:0032991,GO:0034641,GO:0040011,GO:0040029,GO:0040039,GO:0042464,GO:0043170,GO:0043226,GO:0043227,GO:0043228,GO:0043229,GO:0043231,GO:0043232,GO:0043233,GO:0043234,GO:0043933,GO:0044237,GO:0044238,GO:0044260,GO:0044422,GO:0044424,GO:0044427,GO:0044428,GO:0044430,GO:0044446,GO:0044464,GO:0044699,GO:0044707,GO:0044763,GO:0044767,GO:0045132,GO:0046483,GO:0046536,GO:0046982,GO:0046983,GO:0048285,GO:0048610,GO:0048731,GO:0048856,GO:0048869,GO:0048870,GO:0050789,GO:0051179,GO:0051276,GO:0051321,GO:0051383,GO:0051674,GO:0060255,GO:0065007,GO:0070013,GO:0070192,GO:0071103,GO:0071704,GO:0071822,GO:0071840,GO:0090304,GO:0097159,GO:1901360,GO:1901363 | Structural maintenance of chromosomes |
| comp218511_c0_seq10.p2 | GO:0000003,GO:0002064,GO:0003006,GO:0003674,GO:0003824,GO:0005575,GO:0005623,GO:0005886,GO:0006629,GO:0006928,GO:0007009,GO:0007275,GO:0007276,GO:0007281,GO:0007283,GO:0007286,GO:0007291,GO:0007349,GO:0008150,GO:0008152,GO:0008354,GO:0009653,GO:0009888,GO:0009987,GO:0016020,GO:0016043,GO:0016044,GO:0016477,GO:0016740,GO:0016746,GO:0016747,GO:0019953,GO:0022412,GO:0022414,GO:0030154,GO:0030258,GO:0030855,GO:0032501,GO:0032502,GO:0032504,GO:0040011,GO:0044237,GO:0044238,GO:0044255,GO:0044464,GO:0044699,GO:0044702,GO:0044707,GO:0044710,GO:0044763,GO:0044767,GO:0048232,GO:0048468,GO:0048515,GO:0048609,GO:0048610,GO:0048646,GO:0048856,GO:0048869,GO:0048870,GO:0051179,GO:0051674,GO:0060429,GO:0061024,GO:0071617,GO:0071704,GO:0071840,GO:0071944 | MBOAT, membrane-bound O-acyltransferase family |
| comp208339_c0_seq1.p1 | GO:0003674,GO:0003824,GO:0004518,GO:0004519,GO:0004521,GO:0004523,GO:0004540,GO:0005575,GO:0005622,GO:0005623,GO:0005634,GO:0006139,GO:0006259,GO:0006260,GO:0006401,GO:0006725,GO:0006807,GO:0008150,GO:0008152,GO:0009056,GO:0009057,GO:0009058,GO:0009059,GO:0009987,GO:0016070,GO:0016787,GO:0016788,GO:0016891,GO:0016893,GO:0019439,GO:0032299,GO:0032991,GO:0034641,GO:0034645,GO:0034655,GO:0043170,GO:0043226,GO:0043227,GO:0043229,GO:0043231,GO:0043234,GO:0044237,GO:0044238,GO:0044248,GO:0044249,GO:0044260,GO:0044265,GO:0044270,GO:0044424,GO:0044464,GO:0046483,GO:0046700,GO:0071704,GO:0090304,GO:0090305,GO:1901360,GO:1901361,GO:1901575,GO:1901576 | Endonuclease that specifically degrades the RNA of RNA- DNA hybrids (By similarity) |

| | | |
|---|---|---|
| comp193912_c0_seq14.p2 | GO:0000003,GO:0000060,GO:0000375,GO:0000377,GO:0000381,GO:0000398,GO:0002009,GO:0002119,GO:0002164,GO:0003006,GO:0003674,GO:0003676,GO:0003723,GO:0003729,GO:0005215,GO:0005487,GO:0005488,GO:0005575,GO:0005622,GO:0005623,GO:0005634,GO:0005681,GO:0005737,GO:0006139,GO:0006396,GO:0006397,GO:0006605,GO:0006606,GO:0006725,GO:0006807,GO:0006810,GO:0006886,GO:0006913,GO:0006950,GO:0007275,GO:0007276,GO:0007281,GO:0007548,GO:0008104,GO:0008150,GO:0008152,GO:0008380,GO:0009653,GO:0009790,GO:0009791,GO:0009792,GO:0009888,GO:0009893,GO:0009987,GO:0010467,GO:0010468,GO:0010564,GO:0010604,GO:0015031,GO:0016070,GO:0016071,GO:0016482,GO:0017038,GO:0017069,GO:0018996,GO:0019219,GO:0019222,GO:0019953,GO:0022412,GO:0022414,GO:0030154,GO:0030529,GO:0031323,GO:0031325,GO:0032386,GO:0032388,GO:0032501,GO:0032502,GO:0032504,GO:0032879,GO:0032880,GO:0032991,GO:0033036,GO:0033120,GO:0033157,GO:0033365,GO:0033554,GO:0034504,GO:0034613,GO:0034641,GO:0036002,GO:0040007,GO:0040011,GO:0040020,GO:0040035,GO:0042127,GO:0042303,GO:0043170,GO:0043226,GO:0043227,GO:0043229,GO:0043231,GO:0043484,GO:0044237,GO:0044238,GO:0044260,GO:0044422,GO:0044424,GO:0044428,GO:0044446,GO:0044464,GO:0044699,GO:0044702,GO:0044707,GO:0044744,GO:0044763,GO:0044765,GO:0044767,GO:0045184,GO:0045292,GO:0045935,GO:0046483,GO:0046907,GO:0048024,GO:0048468,GO:0048513,GO:0048518,GO:0048522,GO:0048608,GO:0048609,GO:0048610,GO:0048729,GO:0048731,GO:0048806,GO:0048856,GO:0048869,GO:0050684,GO:0050789,GO:0050794,GO:0050896,GO:0051049,GO:0051050,GO:0051169,GO:0051170,GO:0051171,GO:0051173,GO:0051179,GO:0051222,GO:0051223,GO:0051234,GO:0051252,GO:0051254,GO:0051445,GO:0051641,GO:0051649,GO:0051716,GO:0051726,GO:0060255,GO:0060341,GO:0060429,GO:0061458,GO:0065007,GO:0070201,GO:0070727,GO:0071011,GO:0071013,GO:0071702,GO:0071704,GO:0080090,GO:0090304,GO:0090316,GO:0097159,GO:1901360,GO:1901363 | RNA binding motif protein 22 |
| comp193912_c0_seq14.p3 | GO:0003674,GO:0003676,GO:0003723,GO:0005102,GO:0005488,GO:0005515,GO:0005575,GO:0005622,GO:0005623,GO:0005856,GO:0016020,GO:0019898,GO:0030529,GO:0032991,GO:0043226,GO:0043228,GO:0043229,GO:0043232,GO:0044424,GO:0044425,GO:0044464,GO:0050811,GO:0097159,GO:1901363 | janus kinase and microtubule interacting protein |

| | | |
|---|---|---|
| comp218570_c0_seq1.p1 | GO:0000323,GO:0003674,GO:0004871,GO:0004872,GO:0004888,GO:0005215,GO:0005319,GO:0005488,GO:0005496,GO:0005575,GO:0005576,GO:0005622,GO:0005623,GO:0005634,GO:0005635,GO:0005737,GO:0005764,GO:0005765,GO:0005768,GO:0005770,GO:0005773,GO:0005774,GO:0005783,GO:0005794,GO:0005886,GO:0005887,GO:0005975,GO:0006082,GO:0006464,GO:0006486,GO:0006629,GO:0006810,GO:0006869,GO:0006897,GO:0006914,GO:0007034,GO:0007041,GO:0007154,GO:0007165,GO:0007610,GO:0007626,GO:0007628,GO:0008150,GO:0008152,GO:0008202,GO:0008206,GO:0008289,GO:0008344,GO:0009056,GO:0009058,GO:0009059,GO:0009100,GO:0009101,GO:0009719,GO:0009725,GO:0009892,GO:0009894,GO:0009895,GO:0009987,GO:0010008,GO:0010033,GO:0010035,GO:0010038,GO:0010506,GO:0010507,GO:0010646,GO:0010648,GO:0010876,GO:0012505,GO:0014070,GO:0015248,GO:0015485,GO:0015850,GO:0015918,GO:0016020,GO:0016021,GO:0016192,GO:0016241,GO:0016242,GO:0019222,GO:0019538,GO:0019752,GO:0022892,GO:0023052,GO:0030301,GO:0030534,GO:0031090,GO:0031224,GO:0031226,GO:0031323,GO:0031324,GO:0031329,GO:0031330,GO:0031902,GO:0031967,GO:0031975,GO:0031982,GO:0032101,GO:0032102,GO:0032104,GO:0032105,GO:0032107,GO:0032108,GO:0032501,GO:0032787,GO:0032870,GO:0032934,GO:0033036,GO:0033344,GO:0033993,GO:0034645,GO:0036094,GO:0036211,GO:0038023,GO:0042221,GO:0042493,GO:0042592,GO:0042632,GO:0043170,GO:0043178,GO:0043226,GO:0043227,GO:0043229,GO:0043231,GO:0043412,GO:0043413,GO:0043436,GO:0044237,GO:0044238,GO:0044248,GO:0044249,GO:0044260,GO:0044267,GO:0044281,GO:0044422,GO:0044424,GO:0044425,GO:0044428,GO:0044437,GO:0044440,GO:0044444,GO:0044446,GO:0044459,GO:0044464,GO:0044699,GO:0044700,GO:0044707,GO:0044708,GO:0044710,GO:0044723,GO:0044763,GO:0044765,GO:0045121,GO:0046686,GO:0046907,GO:0048471,GO:0048519,GO:0048523,GO:0048545,GO:0048583,GO:0048585,GO:0048878,GO:0050789,GO:0050794,GO:0050896,GO:0051179,GO:0051234,GO:0051641,GO:0051649,GO:0051716,GO:0055088,GO:0055092,GO:0055094,GO:0055098,GO:0060089,GO:0065007,GO:0065008,GO:0070085,GO:0070887,GO:0071310,GO:0071383,GO:0071396,GO:0071402,GO:0071404,GO:0071407,GO:0071495,GO:0071702,GO:0071704,GO:0071944,GO:0080134,GO:0080135,GO:0097159,GO:1901135,GO:1901137,GO:1901360,GO:1901576 | Niemann-Pick disease, type C1 |
| comp88028_c0_seq2.p1 | GO:0001655,GO:0001822,GO:0005575,GO:0005622,GO:0005623,GO:0005737,GO:0005783,GO:0007275,GO:0008150,GO:0032501,GO:0032502,GO:0043226,GO:0043227,GO:0043229,GO:0043231,GO:0044424,GO:0044444,GO:0044464,GO:0044699,GO:0044707,GO:0044767,GO:0048513,GO:0048731,GO:0048793,GO:0048856,GO:0072001 | protein kinase C substrate 80K-H |
| comp219335_c1_seq7.p1 | GO:0000226,GO:0003674,GO:0003824,GO:0004738,GO:0004739,GO:0005575,GO:0005622,GO:0005623,GO:0005737,GO:0005739,GO:0005759,GO:0005811,GO:0005856,GO:0005875,GO:0006082,GO:0006084,GO:0006085,GO:0006086,GO:0006090,GO:0006091,GO:0006099,GO:0006637,GO:0006732,GO:0006793,GO:0006996,GO:0007010,GO:0007017,GO:0007275,GO:0007568,GO:0008150,GO:0008152,GO:0008340,GO:0009058,GO:0009060,GO:0009108,GO:0009790,GO:0009792,GO:0009889,GO:0009987,GO:0010259,GO:0010510,GO:0010565,GO:0015630,GO:0015980,GO:0016043,GO:0016491,GO:0016624,GO:0016903,GO:0019216,GO:0019217,GO:0019222,GO:0019752,GO:0031122,GO:0031323,GO:0031326,GO:0031974,GO:0032501,GO:0032502,GO:0032787,GO:0032991,GO:0035383,GO:0035384,GO:0040008,GO:0040009,GO:0040010,GO:0043226,GO:0043227,GO:0043228,GO:0043229,GO:0043231,GO:0043232,GO:0043233,GO:0043234,GO:0043436,GO:0043933,GO:0044237,GO:0044238,GO:0044249,GO:0044281,GO:0044422,GO:0044424,GO:0044429,GO:0044430,GO:0044444,GO:0044446,GO:0044464,GO:0044699,GO:0044707,GO:0044710,GO:0044763,GO:0044767,GO:0045254,GO:0045333,GO:0045927,GO:0048518,GO:0048856,GO:0050789,GO:0050794,GO:0050812,GO:0051174,GO:0051186,GO:0051188,GO:0051193,GO:0051196,GO:0055114,GO:0065007,GO:0070013,GO:0071616,GO:0071704,GO:0071822,GO:0071840,GO:0080090,GO:1901576 | Pyruvate dehydrogenase (Lipoamide) beta |

| | | |
|---|---|---|
| comp215642_c0_seq1.p1 | GO:0000003,GO:0000075,GO:0000082,GO:0000084,GO:0000166,GO:0000278,GO:0000785,GO:0000910,GO:0001882,GO:0001883,GO:0003006,GO:0003674,GO:0003676,GO:0003677,GO:0003682,GO:0003697,GO:0005488,GO:0005515,GO:0005524,GO:0005575,GO:0005622,GO:0005623,GO:0005634,GO:0005654,GO:0005656,GO:0005694,GO:0006139,GO:0006152,GO:0006163,GO:0006195,GO:0006200,GO:0006259,GO:0006260,GO:0006261,GO:0006267,GO:0006268,GO:0006271,GO:0006275,GO:0006277,GO:0006725,GO:0006753,GO:0006793,GO:0006796,GO:0006807,GO:0007049,GO:0007154,GO:0007267,GO:0007275,GO:0007276,GO:0007292,GO:0007304,GO:0007306,GO:0007307,GO:0007548,GO:0008150,GO:0008152,GO:0009056,GO:0009058,GO:0009059,GO:0009116,GO:0009117,GO:0009119,GO:0009123,GO:0009125,GO:0009126,GO:0009128,GO:0009141,GO:0009143,GO:0009144,GO:0009146,GO:0009150,GO:0009154,GO:0009158,GO:0009161,GO:0009164,GO:0009166,GO:0009167,GO:0009169,GO:0009199,GO:0009203,GO:0009205,GO:0009207,GO:0009259,GO:0009261,GO:0009653,GO:0009790,GO:0009792,GO:0009889,GO:0009987,GO:0010556,GO:0010564,GO:0010927,GO:0010948,GO:0016043,GO:0017076,GO:0018130,GO:0019219,GO:0019222,GO:0019438,GO:0019439,GO:0019637,GO:0019693,GO:0019953,GO:0022402,GO:0022403,GO:0022412,GO:0022414,GO:0022607,GO:0022616,GO:0023052,GO:0030154,GO:0030174,GO:0030554,GO:0030703,GO:0030707,GO:0031323,GO:0031326,GO:0031974,GO:0031981,GO:0032392,GO:0032501,GO:0032502,GO:0032504,GO:0032508,GO:0032549,GO:0032550,GO:0032553,GO:0032555,GO:0032559,GO:0032989,GO:0032991,GO:0032993,GO:0033260,GO:0034622,GO:0034641,GO:0034645,GO:0034654,GO:0034655,GO:0035639,GO:0036094,GO:0036387,GO:0036388,GO:0040035,GO:0042278,GO:0042454,GO:0042555,GO:0043167,GO:0043168,GO:0043170,GO:0043226,GO:0043227,GO:0043228,GO:0043229,GO:0043231,GO:0043232,GO:0043233,GO:0043234,GO:0043566,GO:0043933,GO:0044085,GO:0044237,GO:0044238,GO:0044248,GO:0044249,GO:0044260,GO:0044270,GO:0044271,GO:0044281,GO:0044422,GO:0044424,GO:0044427,GO:0044428,GO:0044446,GO:0044451,GO:0044464,GO:0044699,GO:0044700,GO:0044702,GO:0044707,GO:0044710,GO:0044763,GO:0044767,GO:0044770,GO:0044772,GO:0044786,GO:0045165,GO:0045168,GO:0046034,GO:0046128,GO:0046130,GO:0046331,GO:0046434,GO:0046483,GO:0046700,GO:0048468,GO:0048477,GO:0048513,GO:0048519,GO:0048523,GO:0048608,GO:0048609,GO:0048610,GO:0048646,GO:0048731,GO:0048806,GO:0048856,GO:0048869,GO:0050789,GO:0050794,GO:0051052,GO:0051171,GO:0051301,GO:0051320,GO:0051325,GO:0051329,GO:0051726,GO:0055086,GO:0060255,GO:0061458,GO:0065003,GO:0065004,GO:0065007,GO:0070013,GO:0071103,GO:0071704,GO:0071824,GO:0071840,GO:0071897,GO:0072521,GO:0072523,GO:0080090,GO:0090304,GO:0090329,GO:0097159,GO:1901135,GO:1901136,GO:1901265,GO:1901292,GO:1901360,GO:1901361,GO:1901362,GO:1901363,GO:1901564,GO:1901565,GO:1901575,GO:1901576,GO:1901657,GO:1901658,GO:1901987,GO:1901988,GO:1902299,GO:2000112 | minichromosome maintenance complex component 6 |

| | | |
|---|---|---|
| comp217615_c0_seq4.p1 | GO:0000003,GO:0000041,GO:0000902,GO:0000904,GO:0001667,GO:0001763,GO:0001837,GO:0002009,GO:0003006,GO:0003674,GO:0005215,GO:0005385,GO:0005575,GO:0005622,GO:0005623,GO:0005737,GO:0005783,GO:0005886,GO:0005887,GO:0006810,GO:0006811,GO:0006812,GO:0006829,GO:0006873,GO:0006875,GO:0006882,GO:0006928,GO:0007275,GO:0007276,GO:0007280,GO:0007369,GO:0007399,GO:0007417,GO:0007424,GO:0007498,GO:0007506,GO:0007548,GO:0008150,GO:0008324,GO:0008354,GO:0008406,GO:0009653,GO:0009790,GO:0009887,GO:0009888,GO:0009986,GO:0009987,GO:0010035,GO:0010038,GO:0010043,GO:0014031,GO:0015075,GO:0016020,GO:0016021,GO:0016043,GO:0016477,GO:0019725,GO:0019953,GO:0022008,GO:0022412,GO:0022414,GO:0022857,GO:0022890,GO:0022891,GO:0022892,GO:0030001,GO:0030003,GO:0030027,GO:0030154,GO:0031224,GO:0031226,GO:0031252,GO:0031253,GO:0031256,GO:0031258,GO:0032501,GO:0032502,GO:0032504,GO:0032989,GO:0034220,GO:0035146,GO:0035147,GO:0035239,GO:0035262,GO:0035295,GO:0040011,GO:0042074,GO:0042221,GO:0042592,GO:0042995,GO:0043226,GO:0043227,GO:0043229,GO:0043231,GO:0044424,GO:0044425,GO:0044444,GO:0044459,GO:0044463,GO:0044464,GO:0044699,GO:0044702,GO:0044707,GO:0044763,GO:0044765,GO:0044767,GO:0045137,GO:0046873,GO:0046915,GO:0046916,GO:0048468,GO:0048513,GO:0048598,GO:0048608,GO:0048609,GO:0048610,GO:0048729,GO:0048731,GO:0048754,GO:0048762,GO:0048856,GO:0048863,GO:0048864,GO:0048869,GO:0048870,GO:0048878,GO:0050801,GO:0050896,GO:0051179,GO:0051234,GO:0051674,GO:0055065,GO:0055069,GO:0055076,GO:0055080,GO:0055082,GO:0055085,GO:0060429,GO:0060446,GO:0060485,GO:0060541,GO:0060562,GO:0061138,GO:0061458,GO:0065007,GO:0065008,GO:0071577,GO:0071578,GO:0071840,GO:0071944,GO:0072503,GO:0072507,GO:0072509,GO:0072511 | solute carrier family 39 (Zinc transporter), member |
| comp219739_c2_seq4.p1 | GO:0000075,GO:0000077,GO:0000902,GO:0000904,GO:0002165,GO:0002376,GO:0002520,GO:0003674,GO:0005102,GO:0005488,GO:0005515,GO:0005575,GO:0005622,GO:0005623,GO:0005634,GO:0006139,GO:0006259,GO:0006323,GO:0006325,GO:0006333,GO:0006334,GO:0006338,GO:0006351,GO:0006355,GO:0006357,GO:0006725,GO:0006807,GO:0006950,GO:0006974,GO:0006996,GO:0007049,GO:0007093,GO:0007095,GO:0007154,GO:0007165,GO:0007267,GO:0007275,GO:0007346,GO:0007399,GO:0008150,GO:0008152,GO:0008593,GO:0009058,GO:0009059,GO:0009653,GO:0009791,GO:0009889,GO:0009966,GO:0009967,GO:0009987,GO:0010033,GO:0010467,GO:0010468,GO:0010556,GO:0010564,GO:0010646,GO:0010647,GO:0010948,GO:0014070,GO:0016043,GO:0016070,GO:0016358,GO:0016568,GO:0016589,GO:0016922,GO:0018130,GO:0019219,GO:0019222,GO:0019438,GO:0022008,GO:0022402,GO:0022607,GO:0023051,GO:0023052,GO:0023056,GO:0030030,GO:0030097,GO:0030154,GO:0030182,GO:0030518,GO:0030522,GO:0031010,GO:0031175,GO:0031323,GO:0031326,GO:0031497,GO:0031570,GO:0031572,GO:0032501,GO:0032502,GO:0032774,GO:0032989,GO:0032990,GO:0032991,GO:0033554,GO:0033993,GO:0034622,GO:0034641,GO:0034645,GO:0034654,GO:0034728,GO:0035073,GO:0035075,GO:0035076,GO:0035210,GO:0036314,GO:0036315,GO:0042221,GO:0042766,GO:0043170,GO:0043226,GO:0043227,GO:0043229,GO:0043231,GO:0043234,GO:0043933,GO:0044085,GO:0044237,GO:0044238,GO:0044249,GO:0044260,GO:0044271,GO:0044422,GO:0044424,GO:0044428,GO:0044446,GO:0044464,GO:0044699,GO:0044700,GO:0044707,GO:0044763,GO:0044767,GO:0044773,GO:0044774,GO:0045165,GO:0045168,GO:0045747,GO:0046331,GO:0046483,GO:0048468,GO:0048513,GO:0048518,GO:0048519,GO:0048522,GO:0048523,GO:0048534,GO:0048583,GO:0048584,GO:0048666,GO:0048667,GO:0048699,GO:0048731,GO:0048812,GO:0048813,GO:0048856,GO:0048858,GO:0048869,GO:0050789,GO:0050794,GO:0050896,GO:0051171,GO:0051252,GO:0051276,GO:0051716,GO:0051726,GO:0060255,GO:0065003,GO:0065004,GO:0065007,GO:0070603,GO:0070887,GO:0071103,GO:0071310,GO:0071390,GO:0071396,GO:0071407,GO:0071704,GO:0071824,GO:0071840,GO:0080090,GO:0090304,GO:0097305,GO:0097306,GO:1901360,GO:1901362,GO:1901576,GO:1901654,GO:1901655,GO:1901700,GO:1901701,GO:1901987,GO:1901988,GO:1901990,GO:1901991,GO:2000112,GO:2001141 | Fetal alzheimer antigen |
| comp218375_c0_seq5.p1 | | dipeptidyl-peptidase 10 (non-functional) |

| | | |
|---|---|---|
| comp209120_c0_seq7.p1 | | Autophagy-related protein 11 |
| comp220975_c0_seq2.p1 | GO:0000122,GO:0000981,GO:0001071,GO:0001654,GO:0002165,GO:0003674,GO:0003700,GO:0003705,GO:0005575,GO:0005622,GO:0005623,GO:0005634,GO:0006139,GO:0006351,GO:0006355,GO:0006357,GO:0006366,GO:0006725,GO:0006807,GO:0007275,GO:0007423,GO:0007444,GO:0007478,GO:0007480,GO:0007552,GO:0007560,GO:0008150,GO:0008152,GO:0009058,GO:0009059,GO:0009653,GO:0009791,GO:0009886,GO:0009887,GO:0009889,GO:0009890,GO:0009892,GO:0009987,GO:0010467,GO:0010468,GO:0010556,GO:0010558,GO:0010605,GO:0010629,GO:0016070,GO:0018130,GO:0019219,GO:0019222,GO:0019438,GO:0031323,GO:0031324,GO:0031326,GO:0031327,GO:0032501,GO:0032502,GO:0032774,GO:0034641,GO:0034645,GO:0034654,GO:0035107,GO:0035108,GO:0035114,GO:0035120,GO:0035127,GO:0035218,GO:0043170,GO:0043226,GO:0043227,GO:0043229,GO:0043231,GO:0044237,GO:0044238,GO:0044249,GO:0044260,GO:0044271,GO:0044424,GO:0044464,GO:0044699,GO:0044707,GO:0044767,GO:0045892,GO:0045934,GO:0046483,GO:0048513,GO:0048519,GO:0048523,GO:0048563,GO:0048569,GO:0048707,GO:0048731,GO:0048736,GO:0048737,GO:0048749,GO:0048856,GO:0050789,GO:0050794,GO:0051171,GO:0051172,GO:0051252,GO:0051253,GO:0060173,GO:0060255,GO:0065007,GO:0071704,GO:0080090,GO:0090304,GO:1901360,GO:1901362,GO:1901576,GO:2000112,GO:2000113,GO:2001141 | Zinc finger, C2H2 type |
| comp199801_c0_seq1.p1 | | Coiled-coil-helix-coiled-coil-helix domain-containing protein |
| comp218302_c0_seq16.p1 | GO:0000902,GO:0003674,GO:0005488,GO:0005515,GO:0005575,GO:0005622,GO:0005623,GO:0005737,GO:0005768,GO:0005769,GO:0006810,GO:0006897,GO:0006944,GO:0006996,GO:0008150,GO:0009653,GO:0009987,GO:0010927,GO:0016023,GO:0016043,GO:0016044,GO:0016192,GO:0017016,GO:0017137,GO:0018996,GO:0019899,GO:0022607,GO:0030030,GO:0030031,GO:0030139,GO:0031267,GO:0031410,GO:0031982,GO:0031988,GO:0032501,GO:0032502,GO:0032794,GO:0032989,GO:0032990,GO:0042303,GO:0042384,GO:0043226,GO:0043227,GO:0043229,GO:0043231,GO:0044085,GO:0044424,GO:0044444,GO:0044464,GO:0044699,GO:0044707,GO:0044763,GO:0044767,GO:0044782,GO:0048646,GO:0048856,GO:0048858,GO:0048869,GO:0051020,GO:0051179,GO:0051234,GO:0055037,GO:0060271,GO:0061024,GO:0061025,GO:0070925,GO:0071840 | Rabaptin, RAB GTPase binding effector protein 1 |
| comp217538_c0_seq7.p1 | | kinetochore associated 1 |
| comp220013_c1_seq1.p1 | | BACK |

| | | |
|---|---|---|
| comp209457_c0_seq1.p1 | GO:0000003,GO:0000082,GO:0000084,GO:0000228,GO:0000278,GO:0000722,GO:0000723,GO:0000910,GO:0002009,GO:0002119,GO:0002164,GO:0003006,GO:0003674,GO:0003824,GO:0003887,GO:0003896,GO:0003899,GO:0005575,GO:0005622,GO:0005623,GO:0005634,GO:0005654,GO:0005657,GO:0005658,GO:0005694,GO:0006139,GO:0006259,GO:0006260,GO:0006261,GO:0006269,GO:0006270,GO:0006271,GO:0006310,GO:0006312,GO:0006725,GO:0006807,GO:0006810,GO:0006897,GO:0006898,GO:0006996,GO:0007049,GO:0007275,GO:0007399,GO:0007548,GO:0008150,GO:0008152,GO:0009058,GO:0009059,GO:0009653,GO:0009790,GO:0009791,GO:0009792,GO:0009888,GO:0009987,GO:0010833,GO:0016043,GO:0016070,GO:0016192,GO:0016740,GO:0016772,GO:0016779,GO:0018130,GO:0019438,GO:0022008,GO:0022402,GO:0022403,GO:0022414,GO:0022616,GO:0030154,GO:0030894,GO:0031974,GO:0031981,GO:0032200,GO:0032201,GO:0032501,GO:0032502,GO:0032774,GO:0032991,GO:0032993,GO:0033260,GO:0034061,GO:0034062,GO:0034641,GO:0034645,GO:0034654,GO:0040007,GO:0040035,GO:0042592,GO:0043170,GO:0043226,GO:0043227,GO:0043228,GO:0043229,GO:0043231,GO:0043232,GO:0043233,GO:0043234,GO:0043596,GO:0043601,GO:0044237,GO:0044238,GO:0044249,GO:0044260,GO:0044271,GO:0044422,GO:0044424,GO:0044427,GO:0044428,GO:0044446,GO:0044454,GO:0044464,GO:0044699,GO:0044707,GO:0044763,GO:0044767,GO:0044770,GO:0044772,GO:0044786,GO:0046483,GO:0048513,GO:0048608,GO:0048729,GO:0048731,GO:0048806,GO:0048856,GO:0048869,GO:0051179,GO:0051234,GO:0051276,GO:0051301,GO:0051320,GO:0051325,GO:0051329,GO:0060249,GO:0060429,GO:0061458,GO:0065007,GO:0065008,GO:0070013,GO:0071704,GO:0071840,GO:0090304,GO:1901360,GO:1901362,GO:1901576 | primase, DNA, polypeptide 1 (49kDa) |
| comp206999_c0_seq8.p1 | GO:0000122,GO:0000578,GO:0000902,GO:0000904,GO:0000981,GO:0000982,GO:0001071,GO:0001078,GO:0001227,GO:0001667,GO:0001709,GO:0003002,GO:0003674,GO:0003700,GO:0004871,GO:0004872,GO:0004879,GO:0005488,GO:0005515,GO:0005575,GO:0005622,GO:0005623,GO:0005634,GO:0006139,GO:0006351,GO:0006355,GO:0006357,GO:0006366,GO:0006725,GO:0006807,GO:0006928,GO:0007088,GO:0007154,GO:0007165,GO:0007275,GO:0007346,GO:0007350,GO:0007351,GO:0007354,GO:0007389,GO:0007399,GO:0007417,GO:0007424,GO:0007427,GO:0007517,GO:0008134,GO:0008150,GO:0008152,GO:0008544,GO:0008595,GO:0009058,GO:0009059,GO:0009653,GO:0009790,GO:0009798,GO:0009880,GO:0009888,GO:0009889,GO:0009890,GO:0009892,GO:0009948,GO:0009952,GO:0009987,GO:0010467,GO:0010468,GO:0010556,GO:0010558,GO:0010564,GO:0010605,GO:0010629,GO:0010631,GO:0016043,GO:0016070,GO:0016358,GO:0016477,GO:0018130,GO:0019219,GO:0019222,GO:0019438,GO:0022008,GO:0023052,GO:0030030,GO:0030154,GO:0030182,GO:0030522,GO:0031175,GO:0031323,GO:0031324,GO:0031326,GO:0031327,GO:0032501,GO:0032502,GO:0032774,GO:0032989,GO:0032990,GO:0033043,GO:0034641,GO:0034645,GO:0034654,GO:0035150,GO:0035151,GO:0035152,GO:0035282,GO:0035290,GO:0038023,GO:0040011,GO:0043170,GO:0043226,GO:0043227,GO:0043229,GO:0043231,GO:0043588,GO:0044237,GO:0044238,GO:0044249,GO:0044260,GO:0044271,GO:0044424,GO:0044464,GO:0044699,GO:0044700,GO:0044707,GO:0044763,GO:0044767,GO:0045165,GO:0045892,GO:0045934,GO:0046483,GO:0046845,GO:0048468,GO:0048513,GO:0048519,GO:0048523,GO:0048666,GO:0048667,GO:0048699,GO:0048731,GO:0048812,GO:0048813,GO:0048856,GO:0048858,GO:0048869,GO:0048870,GO:0050789,GO:0050794,GO:0050896,GO:0051128,GO:0051171,GO:0051172,GO:0051179,GO:0051252,GO:0051253,GO:0051674,GO:0051716,GO:0051726,GO:0051783,GO:0060089,GO:0060255,GO:0060541,GO:0061061,GO:0065007,GO:0065008,GO:0070491,GO:0071704,GO:0071840,GO:0080090,GO:0090066,GO:0090130,GO:0090132,GO:0090304,GO:1901360,GO:1901362,GO:1901576,GO:2000112,GO:2000113,GO:2001141 | branched duct epithelial cell fate determination, open tracheal system |

| | | |
|---|---|---|
| comp208993_c1_seq10.p1 | GO:0003674,GO:0005488,GO:0005515,GO:0005575,GO:0005622,GO:0005623,GO:0005737,GO:0005829,GO:0007154,GO:0007165,GO:0007264,GO:0007275,GO:0008150,GO:0009790,GO:0009792,GO:0009966,GO:0009987,GO:0010646,GO:0017016,GO:0017048,GO:0019899,GO:0023051,GO:0023052,GO:0031267,GO:0032501,GO:0032502,GO:0035023,GO:0035556,GO:0040008,GO:0040009,GO:0040010,GO:0044424,GO:0044444,GO:0044464,GO:0044699,GO:0044700,GO:0044707,GO:0044763,GO:0044767,GO:0045927,GO:0046578,GO:0048518,GO:0048583,GO:0048856,GO:0050789,GO:0050794,GO:0050896,GO:0051020,GO:0051056,GO:0051716,GO:0065007 | Rho GTPase activating protein 20 |
| comp215526_c1_seq1.p1 | GO:0000323,GO:0005575,GO:0005622,GO:0005623,GO:0005737,GO:0005764,GO:0005765,GO:0005768,GO:0005770,GO:0005773,GO:0005774,GO:0005819,GO:0005856,GO:0010008,GO:0015630,GO:0016020,GO:0031090,GO:0031902,GO:0032991,GO:0043226,GO:0043227,GO:0043228,GO:0043229,GO:0043231,GO:0043232,GO:0043234,GO:0044422,GO:0044424,GO:0044430,GO:0044437,GO:0044440,GO:0044444,GO:0044446,GO:0044464 | ADP-ribosylation factor-like 8B |
| comp211195_c0_seq1.p1 | GO:0000003,GO:0003674,GO:0003824,GO:0005215,GO:0005488,GO:0005515,GO:0005575,GO:0005622,GO:0005623,GO:0005737,GO:0005739,GO:0005740,GO:0005743,GO:0005746,GO:0005750,GO:0005811,GO:0005856,GO:0005875,GO:0006091,GO:0006119,GO:0006122,GO:0006793,GO:0006796,GO:0006810,GO:0006811,GO:0006812,GO:0006818,GO:0007275,GO:0008121,GO:0008150,GO:0008152,GO:0008324,GO:0009060,GO:0009790,GO:0009792,GO:0009987,GO:0015075,GO:0015077,GO:0015078,GO:0015630,GO:0015672,GO:0015980,GO:0015992,GO:0016020,GO:0016310,GO:0016491,GO:0016679,GO:0016681,GO:0019866,GO:0022857,GO:0022890,GO:0022891,GO:0022892,GO:0022900,GO:0022904,GO:0031090,GO:0031966,GO:0031967,GO:0031975,GO:0032403,GO:0032501,GO:0032502,GO:0032991,GO:0034220,GO:0040011,GO:0040019,GO:0042773,GO:0042775,GO:0043226,GO:0043227,GO:0043228,GO:0043229,GO:0043231,GO:0043232,GO:0043234,GO:0044237,GO:0044281,GO:0044422,GO:0044424,GO:0044425,GO:0044429,GO:0044430,GO:0044444,GO:0044446,GO:0044455,GO:0044464,GO:0044699,GO:0044707,GO:0044710,GO:0044763,GO:0044765,GO:0044767,GO:0045275,GO:0045333,GO:0045995,GO:0048518,GO:0048856,GO:0050789,GO:0050793,GO:0051094,GO:0051179,GO:0051234,GO:0051239,GO:0055085,GO:0055114,GO:0065007,GO:0070469,GO:2000026 | ubiquinol-cytochrome c reductase core protein II |
| comp211195_c0_seq1.p2 | | |
| comp213723_c1_seq1.p1 | | BRCT domain protein |
| comp221121_c2_seq1.p1 | GO:0005575,GO:0005622,GO:0005623,GO:0005737,GO:0005794,GO:0043226,GO:0043227,GO:0043229,GO:0043231,GO:0044424,GO:0044444,GO:0044464 | A kinase (PRKA) anchor protein (yotiao) 9 |
| comp213611_c0_seq20.p1 | | WD domain, G-beta repeat |
| comp218270_c0_seq3.p1 | GO:0005575,GO:0005622,GO:0005623,GO:0005737,GO:0005829,GO:0044424,GO:0044444,GO:0044464 | ATP GTP binding protein-like |
| comp221225_c0_seq4.p1 | | Retrotransposon protein |
| comp221225_c0_seq4.p2 | | ZnF_C2HC |

| | | |
|---|---|---|
| comp220591_c1_seq6.p1 | GO:0003674,GO:0005083,GO:0005096,GO:0005099,GO:0005488,GO:0005515,GO:0006140,GO:0007154,GO:0007165,GO:0007264,GO:0007265,GO:0007275,GO:0007568,GO:0008047,GO:0008150,GO:0008340,GO:0009118,GO:0009894,GO:0009966,GO:0009987,GO:0010259,GO:0010646,GO:0017123,GO:0019219,GO:0019220,GO:0019222,GO:0023051,GO:0023052,GO:0030234,GO:0030695,GO:0030811,GO:0031323,GO:0031329,GO:0032315,GO:0032318,GO:0032320,GO:0032484,GO:0032485,GO:0032501,GO:0032502,GO:0032852,GO:0032856,GO:0032859,GO:0032879,GO:0032880,GO:0033121,GO:0033124,GO:0035556,GO:0043085,GO:0043087,GO:0043547,GO:0044093,GO:0044699,GO:0044700,GO:0044707,GO:0044763,GO:0044767,GO:0046578,GO:0046982,GO:0046983,GO:0048583,GO:0050789,GO:0050790,GO:0050794,GO:0050896,GO:0051056,GO:0051171,GO:0051174,GO:0051336,GO:0051345,GO:0051716,GO:0060178,GO:0060341,GO:0060589,GO:0065007,GO:0065009,GO:0080090,GO:1900542 | Ral GTPase activating protein, beta subunit (Non-catalytic) |
| comp220591_c1_seq6.p2 | GO:0003674,GO:0005083,GO:0005096,GO:0005099,GO:0005488,GO:0005515,GO:0006140,GO:0007154,GO:0007165,GO:0007264,GO:0007265,GO:0007275,GO:0007568,GO:0008047,GO:0008150,GO:0008340,GO:0009118,GO:0009894,GO:0009966,GO:0009987,GO:0010259,GO:0010646,GO:0017123,GO:0019219,GO:0019220,GO:0019222,GO:0023051,GO:0023052,GO:0030234,GO:0030695,GO:0030811,GO:0031323,GO:0031329,GO:0032315,GO:0032318,GO:0032320,GO:0032484,GO:0032485,GO:0032501,GO:0032502,GO:0032852,GO:0032856,GO:0032859,GO:0032879,GO:0032880,GO:0033121,GO:0033124,GO:0035556,GO:0043085,GO:0043087,GO:0043547,GO:0044093,GO:0044699,GO:0044700,GO:0044707,GO:0044763,GO:0044767,GO:0046578,GO:0046982,GO:0046983,GO:0048583,GO:0050789,GO:0050790,GO:0050794,GO:0050896,GO:0051056,GO:0051171,GO:0051174,GO:0051336,GO:0051345,GO:0051716,GO:0060178,GO:0060341,GO:0060589,GO:0065007,GO:0065009,GO:0080090,GO:1900542 | Ral GTPase activating protein, beta subunit (Non-catalytic) |
| comp207425_c0_seq4.p1 | GO:0001505,GO:0001667,GO:0003001,GO:0003008,GO:0003674,GO:0005070,GO:0005488,GO:0005515,GO:0005575,GO:0005622,GO:0005623,GO:0005634,GO:0005737,GO:0005768,GO:0005769,GO:0005829,GO:0005929,GO:0006810,GO:0006836,GO:0006897,GO:0006928,GO:0006950,GO:0007034,GO:0007041,GO:0007154,GO:0007165,GO:0007166,GO:0007167,GO:0007169,GO:0007173,GO:0007243,GO:0007259,GO:0007267,GO:0007268,GO:0007269,GO:0007275,GO:0007424,GO:0007427,GO:0008022,GO:0008150,GO:0008269,GO:0008333,GO:0009611,GO:0009966,GO:0009967,GO:0009968,GO:0009987,GO:0010008,GO:0010627,GO:0010631,GO:0010646,GO:0010647,GO:0010648,GO:0010741,GO:0016020,GO:0016043,GO:0016044,GO:0016192,GO:0016197,GO:0016477,GO:0016482,GO:0019226,GO:0023051,GO:0023052,GO:0023056,GO:0023057,GO:0023061,GO:0030674,GO:0031090,GO:0031901,GO:0032182,GO:0032501,GO:0032502,GO:0032940,GO:0032991,GO:0033565,GO:0035556,GO:0035591,GO:0035637,GO:0038127,GO:0040011,GO:0042058,GO:0042059,GO:0042060,GO:0042995,GO:0043130,GO:0043226,GO:0043227,GO:0043229,GO:0043231,GO:0043234,GO:0043408,GO:0043409,GO:0044422,GO:0044424,GO:0044440,GO:0044444,GO:0044446,GO:0044464,GO:0044699,GO:0044700,GO:0044707,GO:0044763,GO:0044765,GO:0046425,GO:0046903,GO:0046907,GO:0048013,GO:0048488,GO:0048489,GO:0048518,GO:0048519,GO:0048522,GO:0048523,GO:0048583,GO:0048584,GO:0048585,GO:0048731,GO:0048856,GO:0048870,GO:0050789,GO:0050794,GO:0050877,GO:0050896,GO:0051179,GO:0051234,GO:0051640,GO:0051641,GO:0051648,GO:0051649,GO:0051650,GO:0051656,GO:0051674,GO:0051716,GO:0060090,GO:0060541,GO:0061024,GO:0065007,GO:0065008,GO:0071840,GO:0090130,GO:0090132,GO:0097479,GO:0097480,GO:1901184,GO:1901185 | signal transducing adaptor molecule (SH3 domain and ITAM motif) |

| | | |
|---|---|---|
| comp210945_c0_seq2.p1 | GO:0000003,GO:0000075,GO:0000723,GO:0000781,GO:0003674,GO:0003676,GO:0003677,GO:0003697,GO:0003824,GO:0004518,GO:0004527,GO:0005488,GO:0005575,GO:0005622,GO:0005623,GO:0005634,GO:0005694,GO:0005813,GO:0005815,GO:0005856,GO:0006139,GO:0006259,GO:0006260,GO:0006278,GO:0006281,GO:0006289,GO:0006725,GO:0006807,GO:0006950,GO:0006974,GO:0006996,GO:0007004,GO:0008150,GO:0008152,GO:0008408,GO:0008409,GO:0009058,GO:0009059,GO:0009987,GO:0010564,GO:0010833,GO:0010948,GO:0015630,GO:0016043,GO:0016233,GO:0016787,GO:0016788,GO:0022616,GO:0031627,GO:0031848,GO:0031860,GO:0032200,GO:0033554,GO:0034641,GO:0034645,GO:0042592,GO:0043170,GO:0043226,GO:0043227,GO:0043228,GO:0043229,GO:0043231,GO:0043232,GO:0043247,GO:0043566,GO:0044237,GO:0044238,GO:0044249,GO:0044260,GO:0044422,GO:0044424,GO:0044427,GO:0044430,GO:0044446,GO:0044464,GO:0044699,GO:0044763,GO:0046483,GO:0048519,GO:0048523,GO:0050789,GO:0050794,GO:0050896,GO:0051276,GO:0051716,GO:0051726,GO:0060249,GO:0065007,GO:0065008,GO:0071704,GO:0071840,GO:0090304,GO:0090305,GO:0097159,GO:1901360,GO:1901363,GO:1901576,GO:1901987,GO:1901988 | DNA cross-link repair 1B |
| comp220310_c1_seq18.p1 | GO:0005575,GO:0005622,GO:0005623,GO:0005634,GO:0005737,GO:0043226,GO:0043227,GO:0043229,GO:0043231,GO:0044424,GO:0044464 | zinc finger protein 318 |
| comp20601_c0_seq1.p1 | GO:0000975,GO:0000976,GO:0000977,GO:0001012,GO:0001067,GO:0002165,GO:0003674,GO:0003676,GO:0003677,GO:0005488,GO:0005575,GO:0005622,GO:0005623,GO:0005634,GO:0006355,GO:0007275,GO:0007399,GO:0007417,GO:0007444,GO:0007472,GO:0007476,GO:0007552,GO:0007560,GO:0008150,GO:0009653,GO:0009791,GO:0009886,GO:0009887,GO:0009889,GO:0010468,GO:0010556,GO:0019219,GO:0019222,GO:0031323,GO:0031326,GO:0032501,GO:0032502,GO:0035107,GO:0035114,GO:0035120,GO:0035220,GO:0043226,GO:0043227,GO:0043229,GO:0043231,GO:0043565,GO:0044212,GO:0044424,GO:0044464,GO:0044699,GO:0044707,GO:0044767,GO:0048513,GO:0048563,GO:0048569,GO:0048707,GO:0048731,GO:0048736,GO:0048737,GO:0048856,GO:0050789,GO:0050794,GO:0051171,GO:0051252,GO:0060255,GO:0065007,GO:0080090,GO:0097159,GO:1901363,GO:2000112,GO:2001141 | Homeobox domain |
| comp207023_c0_seq7.p1 | | Kazal-type serine protease inhibitor domain |
| comp254358_c0_seq1.p1 | GO:0002118,GO:0002121,GO:0003674,GO:0005488,GO:0005515,GO:0005575,GO:0005622,GO:0005623,GO:0005634,GO:0006355,GO:0006357,GO:0007275,GO:0007399,GO:0007417,GO:0007420,GO:0007424,GO:0007444,GO:0007610,GO:0008056,GO:0008150,GO:0008283,GO:0009889,GO:0009892,GO:0010468,GO:0010556,GO:0010605,GO:0010629,GO:0019219,GO:0019222,GO:0031323,GO:0031326,GO:0032501,GO:0032502,GO:0035214,GO:0035218,GO:0035220,GO:0043226,GO:0043227,GO:0043229,GO:0043231,GO:0044424,GO:0044464,GO:0044699,GO:0044707,GO:0044767,GO:0048513,GO:0048519,GO:0048731,GO:0048856,GO:0050789,GO:0050794,GO:0050896,GO:0051171,GO:0051252,GO:0051704,GO:0051705,GO:0060255,GO:0060541,GO:0065007,GO:0080090,GO:2000112,GO:2001141 | Inherit from inNOG: Zinc finger protein |
| comp215456_c3_seq4.p1 | | axonemal dynein light chain domain containing 1 |
| comp217717_c2_seq6.p1 | | GRB10 interacting GYF protein 1 |
| comp208493_c0_seq1.p1 | | chromosome 14 open reading frame 166B |
| comp216564_c1_seq1.p1 | | guanylate cyclase |
| comp216564_c1_seq1.p2 | | family with sequence similarity 92, member A1 |

| | | |
|---|---|---|
| comp207929_c0_seq4.p1 | GO:0000041,GO:0003674,GO:0005215,GO:0005385,GO:0006810,GO:0006811,GO:0006812,GO:0006829,GO:0008150,GO:0008324,GO:0009987,GO:0010035,GO:0010038,GO:0010043,GO:0015075,GO:0022857,GO:0022890,GO:0022891,GO:0022892,GO:0030001,GO:0034220,GO:0042221,GO:0044699,GO:0044763,GO:0044765,GO:0046873,GO:0046915,GO:0050896,GO:0051179,GO:0051234,GO:0055085,GO:0071577,GO:0072509,GO:0072511 | Cation efflux family |
| comp197793_c0_seq2.p1 | GO:0000578,GO:0000975,GO:0001067,GO:0001568,GO:0001655,GO:0001657,GO:0001932,GO:0001933,GO:0001944,GO:0002376,GO:0003002,GO:0003674,GO:0003676,GO:0003677,GO:0003682,GO:0004871,GO:0005057,GO:0005072,GO:0005102,GO:0005126,GO:0005160,GO:0005488,GO:0005515,GO:0005575,GO:0005622,GO:0005623,GO:0005634,GO:0005737,GO:0005829,GO:0006355,GO:0006357,GO:0006915,GO:0006950,GO:0006955,GO:0007154,GO:0007155,GO:0007165,GO:0007166,GO:0007167,GO:0007178,GO:0007275,GO:0007352,GO:0007389,GO:0007507,GO:0008150,GO:0008219,GO:0009653,GO:0009719,GO:0009725,GO:0009790,GO:0009798,GO:0009880,GO:0009889,GO:0009892,GO:0009950,GO:0009953,GO:0009966,GO:0009968,GO:0009987,GO:0010033,GO:0010466,GO:0010468,GO:0010556,GO:0010563,GO:0010605,GO:0010646,GO:0010648,GO:0010941,GO:0010951,GO:0010990,GO:0010991,GO:0012501,GO:0014070,GO:0016265,GO:0017015,GO:0019219,GO:0019220,GO:0019222,GO:0019229,GO:0022610,GO:0023051,GO:0023052,GO:0023057,GO:0030154,GO:0030509,GO:0030510,GO:0030512,GO:0030514,GO:0030617,GO:0031323,GO:0031324,GO:0031326,GO:0031333,GO:0031399,GO:0031400,GO:0031589,GO:0032268,GO:0032269,GO:0032501,GO:0032502,GO:0033612,GO:0033993,GO:0034405,GO:0034616,GO:0034713,GO:0035051,GO:0035295,GO:0035886,GO:0042221,GO:0042312,GO:0042325,GO:0042326,GO:0042692,GO:0042981,GO:0043066,GO:0043067,GO:0043069,GO:0043086,GO:0043154,GO:0043226,GO:0043227,GO:0043229,GO:0043231,GO:0043254,GO:0043281,GO:0043627,GO:0044057,GO:0044087,GO:0044092,GO:0044212,GO:0044424,GO:0044444,GO:0044464,GO:0044699,GO:0044700,GO:0044707,GO:0044763,GO:0044767,GO:0045907,GO:0045908,GO:0045936,GO:0048468,GO:0048513,GO:0048514,GO:0048518,GO:0048519,GO:0048523,GO:0048545,GO:0048583,GO:0048585,GO:0048731,GO:0048856,GO:0048869,GO:0050789,GO:0050790,GO:0050794,GO:0050896,GO:0051128,GO:0051129,GO:0051145,GO:0051171,GO:0051174,GO:0051239,GO:0051240,GO:0051241,GO:0051246,GO:0051248,GO:0051252,GO:0051336,GO:0051346,GO:0051716,GO:0052547,GO:0052548,GO:0055001,GO:0055006,GO:0060089,GO:0060255,GO:0060393,GO:0060394,GO:0060548,GO:0060947,GO:0060948,GO:0060976,GO:0060977,GO:0061061,GO:0065007,GO:0065009,GO:0070696,GO:0070697,GO:0070698,GO:0072001,GO:0072358,GO:0072359,GO:0080090,GO:0090092,GO:0090101,GO:0090287,GO:0090288,GO:0097084,GO:0097159,GO:1901363,GO:2000112,GO:2000116,GO:2000117,GO:2001141 | SMAD family member 6 |
| comp216979_c0_seq8.p1 | | Inherit from KOG: Retrotransposon protein |
| comp218951_c2_seq1.p1 | | Enhancer of |

| | | |
|---|---|---|
| comp221102_c2_seq12.p1 | GO:0000003,GO:0000122,GO:0000578,GO:0000988,GO:0000989,GO:0001071,GO:0001558,GO:0001654,GO:0001700,GO:0002165,GO:0003002,GO:0003006,GO:0003674,GO:0003676,GO:0003677,GO:0003700,GO:0003712,GO:0005488,GO:0005515,GO:0005575,GO:0005622,GO:0005623,GO:0005634,GO:0006355,GO:0006357,GO:0007275,GO:0007276,GO:0007292,GO:0007304,GO:0007306,GO:0007350,GO:0007351,GO:0007354,GO:0007362,GO:0007389,GO:0007423,GO:0007444,GO:0007447,GO:0007450,GO:0007472,GO:0007474,GO:0007476,GO:0007552,GO:0007560,GO:0008134,GO:0008150,GO:0008595,GO:0009653,GO:0009790,GO:0009791,GO:0009792,GO:0009798,GO:0009880,GO:0009886,GO:0009887,GO:0009889,GO:0009890,GO:0009892,GO:0009948,GO:0009952,GO:0009953,GO:0009987,GO:0010468,GO:0010556,GO:0010558,GO:0010605,GO:0010629,GO:0010927,GO:0016043,GO:0019219,GO:0019222,GO:0019953,GO:0022412,GO:0022414,GO:0030154,GO:0030308,GO:0030703,GO:0030707,GO:0031323,GO:0031324,GO:0031326,GO:0031327,GO:0032501,GO:0032502,GO:0032504,GO:0032989,GO:0035107,GO:0035114,GO:0035120,GO:0035220,GO:0035222,GO:0035282,GO:0040008,GO:0043226,GO:0043227,GO:0043229,GO:0043231,GO:0043565,GO:0044424,GO:0044464,GO:0044699,GO:0044702,GO:0044707,GO:0044763,GO:0044767,GO:0045892,GO:0045926,GO:0045934,GO:0046843,GO:0048190,GO:0048468,GO:0048477,GO:0048513,GO:0048519,GO:0048523,GO:0048563,GO:0048569,GO:0048592,GO:0048609,GO:0048610,GO:0048646,GO:0048707,GO:0048731,GO:0048736,GO:0048737,GO:0048856,GO:0048869,GO:0050789,GO:0050794,GO:0051128,GO:0051171,GO:0051172,GO:0051252,GO:0051253,GO:0060255,GO:0065007,GO:0070491,GO:0071840,GO:0080090,GO:0097159,GO:1901363,GO:2000112,GO:2000113,GO:2001141 | Inherit from biNOG: capicua homolog (Drosophila) |
| comp220925_c1_seq9.p1 | GO:0003674,GO:0003824,GO:0004672,GO:0004674,GO:0004693,GO:0005575,GO:0005622,GO:0005623,GO:0005694,GO:0005700,GO:0005703,GO:0006464,GO:0006468,GO:0006793,GO:0006796,GO:0006810,GO:0006897,GO:0006909,GO:0007049,GO:0007275,GO:0007399,GO:0008150,GO:0008152,GO:0008353,GO:0009987,GO:0016192,GO:0016301,GO:0016310,GO:0016740,GO:0016772,GO:0016773,GO:0019538,GO:0022008,GO:0030154,GO:0032501,GO:0032502,GO:0036211,GO:0043170,GO:0043226,GO:0043228,GO:0043229,GO:0043232,GO:0043412,GO:0044237,GO:0044238,GO:0044260,GO:0044267,GO:0044422,GO:0044424,GO:0044427,GO:0044446,GO:0044464,GO:0044699,GO:0044707,GO:0044763,GO:0044765,GO:0048731,GO:0048856,GO:0048869,GO:0051179,GO:0051234,GO:0071704,GO:0097472 | S_TKc |
| comp212325_c3_seq1.p1 | | ATPase, H transporting, lysosomal accessory protein 2 |
| comp207985_c0_seq1.p1 | GO:0003674,GO:0005215,GO:0005487,GO:0005575,GO:0005622,GO:0005623,GO:0005634,GO:0005635,GO:0005643,GO:0005654,GO:0005737,GO:0005829,GO:0005975,GO:0006139,GO:0006403,GO:0006405,GO:0006406,GO:0006605,GO:0006606,GO:0006611,GO:0006725,GO:0006807,GO:0006810,GO:0006886,GO:0006913,GO:0007154,GO:0007165,GO:0007166,GO:0008104,GO:0008150,GO:0008152,GO:0008643,GO:0008645,GO:0009987,GO:0010033,GO:0010467,GO:0010827,GO:0012505,GO:0015031,GO:0015749,GO:0015758,GO:0015931,GO:0016020,GO:0016021,GO:0016032,GO:0016070,GO:0016071,GO:0016482,GO:0017038,GO:0019221,GO:0023052,GO:0031224,GO:0031967,GO:0031974,GO:0031975,GO:0031981,GO:0032879,GO:0032991,GO:0033036,GO:0033365,GO:0034097,GO:0034504,GO:0034613,GO:0034641,GO:0042221,GO:0043170,GO:0043226,GO:0043227,GO:0043229,GO:0043231,GO:0043233,GO:0043234,GO:0044237,GO:0044238,GO:0044260,GO:0044281,GO:0044403,GO:0044419,GO:0044422,GO:0044424,GO:0044425,GO:0044428,GO:0044444,GO:0044446,GO:0044464,GO:0044699,GO:0044700,GO:0044710,GO:0044744,GO:0044763,GO:0044764,GO:0044765,GO:0045184,GO:0046483,GO:0046907,GO:0046930,GO:0050657,GO:0050658,GO:0050789,GO:0050794,GO:0050896,GO:0051028,GO:0051049,GO:0051168,GO:0051169,GO:0051170,GO:0051179,GO:0051234,GO:0051236,GO:0051641,GO:0051649,GO:0051704,GO:0051716,GO:0051726,GO:0055085,GO:0065007,GO:0070013,GO:0070727,GO:0070887,GO:0071310,GO:0071345,GO:0071702,GO:0071704,GO:0071705,GO:0090304,GO:1901360 | nucleoporin 214kDa |

| | | |
|---|---|---|
| comp207429_c1_seq1.p1 | | von Willebrand factor type A domain |
| comp215417_c0_seq3.p1 | | kelch-like |
| comp215417_c0_seq3.p2 | | Zinc finger protein |
| comp205553_c0_seq2.p1 | GO:0000003,GO:0000018,GO:0000279,GO:0003674,GO:0003676,GO:0003677,GO:0003824,GO:0005488,GO:0005575,GO:0005622,GO:0005623,GO:0005634,GO:0005654,GO:0005694,GO:0006139,GO:0006259,GO:0006310,GO:0006311,GO:0006325,GO:0006355,GO:0006357,GO:0006464,GO:0006479,GO:0006725,GO:0006807,GO:0006996,GO:0007049,GO:0007126,GO:0007127,GO:0007128,GO:0007276,GO:0007283,GO:0008150,GO:0008152,GO:0008168,GO:0008170,GO:0008213,GO:0008276,GO:0008757,GO:0009889,GO:0009891,GO:0009893,GO:0009987,GO:0010468,GO:0010520,GO:0010556,GO:0010557,GO:0010564,GO:0010604,GO:0010628,GO:0010844,GO:0010845,GO:0016043,GO:0016278,GO:0016279,GO:0016568,GO:0016569,GO:0016570,GO:0016571,GO:0016740,GO:0016741,GO:0018022,GO:0018024,GO:0018193,GO:0018205,GO:0019219,GO:0019222,GO:0019538,GO:0019953,GO:0022402,GO:0022403,GO:0022414,GO:0031323,GO:0031325,GO:0031326,GO:0031328,GO:0031974,GO:0031981,GO:0032259,GO:0032501,GO:0032504,GO:0034641,GO:0034968,GO:0035822,GO:0036211,GO:0040020,GO:0042054,GO:0042800,GO:0043170,GO:0043226,GO:0043227,GO:0043228,GO:0043229,GO:0043231,GO:0043232,GO:0043233,GO:0043412,GO:0043414,GO:0043565,GO:0044237,GO:0044238,GO:0044260,GO:0044267,GO:0044422,GO:0044424,GO:0044428,GO:0044446,GO:0044464,GO:0044699,GO:0044702,GO:0044763,GO:0045836,GO:0045893,GO:0045911,GO:0045935,GO:0045944,GO:0046483,GO:0048232,GO:0048518,GO:0048522,GO:0048609,GO:0048610,GO:0050789,GO:0050794,GO:0051052,GO:0051054,GO:0051171,GO:0051173,GO:0051252,GO:0051254,GO:0051276,GO:0051321,GO:0051324,GO:0051445,GO:0051568,GO:0051726,GO:0060255,GO:0060631,GO:0060903,GO:0065007,GO:0070013,GO:0071704,GO:0071840,GO:0080090,GO:0090068,GO:0090304,GO:0097159,GO:1901360,GO:1901363,GO:2000112,GO:2001141 | Histone methyltransferase that specifically trimethylates 'Lys-4' of histone H3 during meiotic prophase and is essential for proper meiotic progression. Does not have the ability to mono- and dimethylate 'Lys-4' of histone H3. H3 'Lys-4' methylation represents a specific tag for epigenetic transcriptional activation. Plays a central role in the transcriptional activation of genes during early meiotic prophase |

| | | |
|---|---|---|
| comp208323_c0_seq1.p1 | GO:0000003,GO:0000075,GO:0000077,GO:0000082,GO:0000084,GO:0000209,GO:0000278,GO:0000502,GO:0002119,GO:0002164,GO:0002376,GO:0002474,GO:0002478,GO:0002479,GO:0005575,GO:0005622,GO:0005623,GO:0005634,GO:0005654,GO:0005737,GO:0005829,GO:0005838,GO:0005856,GO:0005875,GO:0006139,GO:0006461,GO:0006464,GO:0006508,GO:0006511,GO:0006521,GO:0006725,GO:0006807,GO:0006915,GO:0006950,GO:0006974,GO:0006977,GO:0007049,GO:0007093,GO:0007154,GO:0007165,GO:0007275,GO:0007346,GO:0007568,GO:0008150,GO:0008152,GO:0008219,GO:0008340,GO:0008541,GO:0009056,GO:0009057,GO:0009790,GO:0009791,GO:0009792,GO:0009892,GO:0009893,GO:0009987,GO:0010259,GO:0010467,GO:0010498,GO:0010564,GO:0010565,GO:0010604,GO:0010605,GO:0010941,GO:0010948,GO:0012501,GO:0015630,GO:0016032,GO:0016043,GO:0016070,GO:0016071,GO:0016265,GO:0016567,GO:0019222,GO:0019538,GO:0019882,GO:0019884,GO:0019941,GO:0022402,GO:0022403,GO:0022607,GO:0022624,GO:0023052,GO:0030154,GO:0030163,GO:0030330,GO:0031145,GO:0031323,GO:0031324,GO:0031325,GO:0031396,GO:0031397,GO:0031398,GO:0031399,GO:0031400,GO:0031401,GO:0031570,GO:0031571,GO:0031974,GO:0031981,GO:0032268,GO:0032269,GO:0032270,GO:0032446,GO:0032501,GO:0032502,GO:0032991,GO:0033238,GO:0033554,GO:0034622,GO:0034641,GO:0035556,GO:0036211,GO:0042590,GO:0042770,GO:0042981,GO:0043067,GO:0043085,GO:0043086,GO:0043161,GO:0043170,GO:0043226,GO:0043227,GO:0043228,GO:0043229,GO:0043231,GO:0043232,GO:0043233,GO:0043234,GO:0043248,GO:0043412,GO:0043623,GO:0043632,GO:0043933,GO:0044085,GO:0044092,GO:0044093,GO:0044237,GO:0044238,GO:0044248,GO:0044257,GO:0044260,GO:0044265,GO:0044267,GO:0044281,GO:0044403,GO:0044419,GO:0044422,GO:0044424,GO:0044428,GO:0044430,GO:0044444,GO:0044446,GO:0044464,GO:0044699,GO:0044700,GO:0044707,GO:0044710,GO:0044763,GO:0044764,GO:0044767,GO:0044770,GO:0044772,GO:0044773,GO:0044774,GO:0044783,GO:0046483,GO:0048002,GO:0048518,GO:0048519,GO:0048522,GO:0048523,GO:0048856,GO:0048863,GO:0048869,GO:0050789,GO:0050790,GO:0050794,GO:0050896,GO:0051171,GO:0051246,GO:0051247,GO:0051248,GO:0051320,GO:0051325,GO:0051329,GO:0051340,GO:0051351,GO:0051352,GO:0051436,GO:0051437,GO:0051438,GO:0051439,GO:0051443,GO:0051444,GO:0051603,GO:0051704,GO:0051716,GO:0051726,GO:0060255,GO:0065003,GO:0065007,GO:0065009,GO:0070013,GO:0070271,GO:0070647,GO:0071156,GO:0071158,GO:0071704,GO:0071822,GO:0071840,GO:0072331,GO:0072395,GO:0072401,GO:0072413,GO:0072422,GO:0072431,GO:0080090,GO:0090068,GO:0090304,GO:1901360,GO:1901575,GO:1901987,GO:1901988,GO:1901990,GO:1901991,GO:2000045,GO:2000134 | proteasome (prosome, macropain) 26S subunit, non-ATPase, 11 |
| comp220468_c0_seq5.p1 | GO:0000003,GO:0001667,GO:0003006,GO:0006928,GO:0007276,GO:0007292,GO:0007297,GO:0007298,GO:0008150,GO:0009987,GO:0010631,GO:0016477,GO:0019953,GO:0022412,GO:0022414,GO:0030154,GO:0030707,GO:0032501,GO:0032502,GO:0032504,GO:0040011,GO:0044699,GO:0044702,GO:0044707,GO:0044763,GO:0044767,GO:0048468,GO:0048477,GO:0048609,GO:0048610,GO:0048856,GO:0048869,GO:0048870,GO:0051179,GO:0051674,GO:0090130,GO:0090132 | TBC |
| comp219654_c1_seq5.p1 | | Inherit from NOG: testis specific 10 |
| comp216392_c1_seq7.p1 | | Protein of unknown function (DUF3657) |

| | | |
|---|---|---|
| comp210442_c0_seq6.p1 | GO:0000975,GO:0000976,GO:0000977,GO:0000981,GO:0001012,GO:0001067,GO:0001071,GO:0003674,GO:0003676,GO:0003677,GO:0003700,GO:0004871,GO:0004872,GO:0004879,GO:0005488,GO:0005575,GO:0005622,GO:0005623,GO:0005634,GO:0006109,GO:0006139,GO:0006351,GO:0006355,GO:0006357,GO:0006366,GO:0006725,GO:0006807,GO:0006950,GO:0007154,GO:0007165,GO:0008150,GO:0008152,GO:0009058,GO:0009059,GO:0009605,GO:0009889,GO:0009987,GO:0009991,GO:0010467,GO:0010468,GO:0010556,GO:0010675,GO:0010906,GO:0016070,GO:0018130,GO:0019219,GO:0019222,GO:0019438,GO:0023052,GO:0030522,GO:0031323,GO:0031326,GO:0031667,GO:0032774,GO:0032881,GO:0034641,GO:0034645,GO:0034654,GO:0038023,GO:0042592,GO:0042594,GO:0042632,GO:0043170,GO:0043226,GO:0043227,GO:0043229,GO:0043231,GO:0043467,GO:0043565,GO:0044212,GO:0044237,GO:0044238,GO:0044249,GO:0044260,GO:0044271,GO:0044424,GO:0044464,GO:0044699,GO:0044700,GO:0044763,GO:0046483,GO:0048878,GO:0050789,GO:0050794,GO:0050896,GO:0051171,GO:0051252,GO:0051716,GO:0055088,GO:0055090,GO:0055092,GO:0060089,GO:0060255,GO:0065007,GO:0065008,GO:0070328,GO:0070873,GO:0071704,GO:0080090,GO:0090304,GO:0097159,GO:1901360,GO:1901362,GO:1901363,GO:1901576,GO:2000112,GO:2001141 | receptor |
| comp215134_c0_seq9.p1 | GO:0000428,GO:0003674,GO:0003824,GO:0003899,GO:0005575,GO:0005622,GO:0005623,GO:0005634,GO:0005654,GO:0005730,GO:0005736,GO:0006139,GO:0006351,GO:0006352,GO:0006353,GO:0006354,GO:0006360,GO:0006361,GO:0006362,GO:0006363,GO:0006725,GO:0006807,GO:0008150,GO:0008152,GO:0009058,GO:0009059,GO:0009303,GO:0009987,GO:0010467,GO:0016070,GO:0016740,GO:0016772,GO:0016779,GO:0018130,GO:0019438,GO:0030880,GO:0031974,GO:0031981,GO:0032774,GO:0032991,GO:0034062,GO:0034641,GO:0034645,GO:0034654,GO:0043170,GO:0043226,GO:0043227,GO:0043228,GO:0043229,GO:0043231,GO:0043232,GO:0043233,GO:0043234,GO:0044237,GO:0044238,GO:0044249,GO:0044260,GO:0044271,GO:0044422,GO:0044424,GO:0044428,GO:0044446,GO:0044452,GO:0044464,GO:0046483,GO:0055029,GO:0070013,GO:0071704,GO:0090304,GO:1901360,GO:1901362,GO:1901576 | polymerase (RNA) I polypeptide A |
| comp219388_c0_seq3.p1 | | |
| comp214778_c0_seq1.p1 | | flightless I homolog (Drosophila) |

| | | |
|---|---|---|
| comp220328_c0_seq3.p1 | GO:0000003,GO:0001655,GO:0001822,GO:0002376,GO:0002474,GO:0003674,GO:0005488,GO:0005515,GO:0005575,GO:0005622,GO:0005623,GO:0005634,GO:0005737,GO:0005829,GO:0006464,GO:0006473,GO:0006475,GO:0006508,GO:0006511,GO:0006915,GO:0006950,GO:0006974,GO:0006996,GO:0007029,GO:0007049,GO:0007126,GO:0007127,GO:0007129,GO:0007130,GO:0007154,GO:0007165,GO:0007275,GO:0007276,GO:0007283,GO:0007399,GO:0007417,GO:0007420,GO:0008104,GO:0008150,GO:0008152,GO:0008219,GO:0008630,GO:0009056,GO:0009057,GO:0009790,GO:0009892,GO:0009894,GO:0009895,GO:0009987,GO:0010468,GO:0010605,GO:0010608,GO:0012501,GO:0016043,GO:0016044,GO:0016265,GO:0018193,GO:0018205,GO:0018393,GO:0018394,GO:0019222,GO:0019538,GO:0019882,GO:0019941,GO:0019953,GO:0022402,GO:0022414,GO:0022607,GO:0023052,GO:0030162,GO:0030163,GO:0030323,GO:0030324,GO:0031323,GO:0031324,GO:0031329,GO:0031330,GO:0031593,GO:0031647,GO:0032182,GO:0032268,GO:0032269,GO:0032403,GO:0032434,GO:0032435,GO:0032501,GO:0032502,GO:0032504,GO:0032991,GO:0033036,GO:0033365,GO:0033554,GO:0034613,GO:0034976,GO:0035295,GO:0035556,GO:0036211,GO:0042127,GO:0042176,GO:0042177,GO:0042771,GO:0043021,GO:0043022,GO:0043130,GO:0043170,GO:0043226,GO:0043227,GO:0043229,GO:0043231,GO:0043234,GO:0043412,GO:0043543,GO:0043632,GO:0044085,GO:0044237,GO:0044238,GO:0044248,GO:0044257,GO:0044260,GO:0044265,GO:0044267,GO:0044424,GO:0044444,GO:0044445,GO:0044464,GO:0044699,GO:0044700,GO:0044702,GO:0044707,GO:0044763,GO:0044767,GO:0045048,GO:0045861,GO:0048002,GO:0048232,GO:0048513,GO:0048519,GO:0048523,GO:0048609,GO:0048610,GO:0048731,GO:0048856,GO:0050789,GO:0050794,GO:0050821,GO:0050896,GO:0051179,GO:0051205,GO:0051246,GO:0051248,GO:0051276,GO:0051321,GO:0051603,GO:0051641,GO:0051668,GO:0051716,GO:0060255,GO:0060541,GO:0061024,GO:0061136,GO:0065007,GO:0065008,GO:0070059,GO:0070192,GO:0070193,GO:0070628,GO:0070727,GO:0071704,GO:0071816,GO:0071818,GO:0071840,GO:0072001,GO:0072331,GO:0072332,GO:0072379,GO:0080090,GO:0097190,GO:0097193,GO:1901575,GO:1901799 | BCL2-associated athanogene 6 |
| comp220328_c0_seq3.p2 | | inositol polyphosphate multikinase |
| comp211517_c1_seq3.p1 | | NHL repeat |
| comp84022_c0_seq1.p1 | GO:0000815,GO:0000910,GO:0002119,GO:0002164,GO:0003674,GO:0005488,GO:0005543,GO:0005546,GO:0005575,GO:0005622,GO:0005623,GO:0005737,GO:0005768,GO:0005769,GO:0005770,GO:0005829,GO:0006810,GO:0006897,GO:0006909,GO:0007034,GO:0007041,GO:0007049,GO:0007275,GO:0007568,GO:0008150,GO:0008289,GO:0008333,GO:0008340,GO:0009790,GO:0009791,GO:0009792,GO:0009987,GO:0010008,GO:0010259,GO:0010876,GO:0016020,GO:0016043,GO:0016044,GO:0016192,GO:0016197,GO:0016482,GO:0018996,GO:0019915,GO:0022402,GO:0030496,GO:0031090,GO:0031902,GO:0032501,GO:0032502,GO:0032535,GO:0032991,GO:0033036,GO:0035091,GO:0040007,GO:0040008,GO:0040009,GO:0040010,GO:0040011,GO:0040014,GO:0040018,GO:0042303,GO:0043167,GO:0043168,GO:0043226,GO:0043227,GO:0043229,GO:0043231,GO:0043234,GO:0044422,GO:0044424,GO:0044425,GO:0044440,GO:0044444,GO:0044446,GO:0044464,GO:0044699,GO:0044707,GO:0044763,GO:0044765,GO:0044767,GO:0045927,GO:0046907,GO:0048518,GO:0048856,GO:0050789,GO:0051036,GO:0051179,GO:0051234,GO:0051239,GO:0051240,GO:0051301,GO:0051641,GO:0051649,GO:0061024,GO:0065007,GO:0065008,GO:0071840,GO:0090066,GO:1901981 | core component of the endosomal sorting required for transport complex III (ESCRT-III) which is involved in multivesicular bodies (MVBs) formation and sorting of endosomal cargo proteins into MVBs. MVBs contain intraluminal vesicles (ILVs) that are generated by invagination and scission from the limiting membrane of the endosome and mostly are delivered to lysosomes enabling degradation of membrane proteins, such as stimulated growth factor receptors, lysosomal enzymes and lipids |

| | | |
|---|---|---|
| comp215961_c0_seq11.p1 | GO:0000902,GO:0001736,GO:0001737,GO:0001738,GO:0002009,GO:0002165,GO:0005575,GO:0005622,GO:0005623,GO:0005737,GO:0005886,GO:0005929,GO:0005930,GO:0006996,GO:0007010,GO:0007163,GO:0007164,GO:0007275,GO:0007399,GO:0007444,GO:0007472,GO:0007476,GO:0007552,GO:0007560,GO:0008150,GO:0008360,GO:0008544,GO:0009653,GO:0009791,GO:0009886,GO:0009887,GO:0009888,GO:0009913,GO:0009987,GO:0016020,GO:0016043,GO:0016476,GO:0022008,GO:0022603,GO:0022604,GO:0022606,GO:0030010,GO:0030030,GO:0030154,GO:0030182,GO:0030855,GO:0031514,GO:0032185,GO:0032501,GO:0032502,GO:0032838,GO:0032879,GO:0032880,GO:0032989,GO:0032990,GO:0035085,GO:0035107,GO:0035114,GO:0035120,GO:0035220,GO:0035315,GO:0035316,GO:0035317,GO:0035320,GO:0042995,GO:0043226,GO:0043227,GO:0043229,GO:0043231,GO:0043588,GO:0044422,GO:0044424,GO:0044441,GO:0044444,GO:0044446,GO:0044463,GO:0044464,GO:0044699,GO:0044707,GO:0044763,GO:0044767,GO:0045995,GO:0048513,GO:0048563,GO:0048569,GO:0048699,GO:0048707,GO:0048729,GO:0048731,GO:0048736,GO:0048737,GO:0048856,GO:0048858,GO:0048869,GO:0050789,GO:0050793,GO:0050794,GO:0051128,GO:0051239,GO:0060271,GO:0060429,GO:0061162,GO:0061339,GO:0065007,GO:0065008,GO:0071840,GO:0071944,GO:0097014,GO:2000026 | WD repeat containing planar cell polarity effector |
| comp215961_c0_seq11.p2 | | WD repeat containing planar cell polarity effector |
| comp204963_c0_seq1.p1 | | protein serine kinase H1 |
| comp217018_c3_seq6.p1 | GO:0003674,GO:0005215,GO:0005575,GO:0005623,GO:0005886,GO:0006810,GO:0006811,GO:0006812,GO:0006814,GO:0006818,GO:0006820,GO:0006821,GO:0008150,GO:0008324,GO:0008509,GO:0008510,GO:0009987,GO:0015075,GO:0015077,GO:0015081,GO:0015103,GO:0015106,GO:0015291,GO:0015293,GO:0015294,GO:0015296,GO:0015297,GO:0015301,GO:0015672,GO:0015698,GO:0015701,GO:0015711,GO:0015992,GO:0016020,GO:0016021,GO:0022804,GO:0022857,GO:0022890,GO:0022891,GO:0022892,GO:0030001,GO:0031224,GO:0034220,GO:0035725,GO:0044425,GO:0044464,GO:0044699,GO:0044763,GO:0044765,GO:0046873,GO:0051179,GO:0051234,GO:0055085,GO:0071702,GO:0071944 | HCO3- transporter family |
| comp211293_c0_seq18.p1 | GO:0000086,GO:0000226,GO:0000278,GO:0005575,GO:0005622,GO:0005623,GO:0005737,GO:0005813,GO:0005814,GO:0005815,GO:0005829,GO:0005856,GO:0006996,GO:0007010,GO:0007017,GO:0007049,GO:0007098,GO:0007099,GO:0008150,GO:0009987,GO:0010564,GO:0015630,GO:0016043,GO:0022402,GO:0031023,GO:0032465,GO:0043226,GO:0043228,GO:0043229,GO:0043232,GO:0044422,GO:0044424,GO:0044430,GO:0044444,GO:0044446,GO:0044450,GO:0044464,GO:0044699,GO:0044763,GO:0044770,GO:0044772,GO:0050789,GO:0050794,GO:0051297,GO:0051298,GO:0051302,GO:0051726,GO:0065007,GO:0071840 | centriolar coiled coil protein 110kDa |
| comp218113_c0_seq2.p1 | | NEL-like 2 (chicken) |
| comp220781_c1_seq4.p1 | | Inherit from NOG: pathogenesis |
| comp220781_c1_seq4.p5 | | |
| comp220781_c1_seq4.p6 | | |
| comp220781_c1_seq4.p7 | | formin homology 2 domain containing 1 |
| comp220781_c1_seq4.p9 | | |

| | | |
|---|---|---|
| comp220906_c0_seq6.p1 | GO:0000226,GO:0003674,GO:0005198,GO:0005488,GO:0005515,GO:0005575,GO:0005622,GO:0005623,GO:0005737,GO:0005813,GO:0005814,GO:0005815,GO:0005856,GO:0005886,GO:0005929,GO:0006996,GO:0007010,GO:0007017,GO:0008092,GO:0008104,GO:0008150,GO:0009987,GO:0015629,GO:0015630,GO:0016020,GO:0016043,GO:0019894,GO:0031023,GO:0031514,GO:0033036,GO:0035253,GO:0042995,GO:0043226,GO:0043227,GO:0043228,GO:0043229,GO:0043231,GO:0043232,GO:0044422,GO:0044424,GO:0044430,GO:0044441,GO:0044444,GO:0044446,GO:0044450,GO:0044463,GO:0044464,GO:0044699,GO:0044763,GO:0051179,GO:0051297,GO:0071840,GO:0071944 | Ciliary rootlet coiled-coil, rootletin |
| comp220906_c0_seq6.p2 | GO:0000226,GO:0003674,GO:0005198,GO:0005488,GO:0005515,GO:0005575,GO:0005622,GO:0005623,GO:0005737,GO:0005813,GO:0005814,GO:0005815,GO:0005856,GO:0005886,GO:0005929,GO:0006996,GO:0007010,GO:0007017,GO:0008092,GO:0008104,GO:0008150,GO:0009987,GO:0015629,GO:0015630,GO:0016020,GO:0016043,GO:0019894,GO:0031023,GO:0031514,GO:0033036,GO:0035253,GO:0042995,GO:0043226,GO:0043227,GO:0043228,GO:0043229,GO:0043231,GO:0043232,GO:0044422,GO:0044424,GO:0044430,GO:0044441,GO:0044444,GO:0044446,GO:0044450,GO:0044463,GO:0044464,GO:0044699,GO:0044763,GO:0051179,GO:0051297,GO:0071840,GO:0071944 | Ciliary rootlet coiled-coil, rootletin |
| comp213527_c2_seq1.p1 | GO:0000003,GO:0000084,GO:0000226,GO:0000228,GO:0000278,GO:0000722,GO:0000723,GO:0003006,GO:0003674,GO:0003824,GO:0003887,GO:0005575,GO:0005622,GO:0005623,GO:0005634,GO:0005654,GO:0005657,GO:0005694,GO:0005856,GO:0005875,GO:0006139,GO:0006259,GO:0006260,GO:0006261,GO:0006271,GO:0006281,GO:0006283,GO:0006284,GO:0006289,GO:0006297,GO:0006310,GO:0006312,GO:0006725,GO:0006807,GO:0006950,GO:0006974,GO:0006996,GO:0007010,GO:0007017,GO:0007049,GO:0007051,GO:0007052,GO:0007275,GO:0007548,GO:0008150,GO:0008152,GO:0009058,GO:0009059,GO:0009790,GO:0009792,GO:0009987,GO:0010833,GO:0015630,GO:0016043,GO:0016740,GO:0016772,GO:0016779,GO:0022402,GO:0022403,GO:0022414,GO:0022616,GO:0030894,GO:0031974,GO:0031981,GO:0032200,GO:0032201,GO:0032501,GO:0032502,GO:0032991,GO:0032993,GO:0033260,GO:0033554,GO:0034061,GO:0034641,GO:0034645,GO:0040008,GO:0040009,GO:0040010,GO:0040011,GO:0040035,GO:0042575,GO:0042592,GO:0043170,GO:0043226,GO:0043227,GO:0043228,GO:0043229,GO:0043231,GO:0043232,GO:0043233,GO:0043234,GO:0043596,GO:0043601,GO:0043625,GO:0043933,GO:0044237,GO:0044238,GO:0044249,GO:0044260,GO:0044422,GO:0044424,GO:0044427,GO:0044428,GO:0044430,GO:0044446,GO:0044454,GO:0044464,GO:0044699,GO:0044707,GO:0044763,GO:0044767,GO:0044786,GO:0045927,GO:0046483,GO:0048513,GO:0048518,GO:0048608,GO:0048731,GO:0048806,GO:0048856,GO:0050789,GO:0050896,GO:0051276,GO:0051320,GO:0051325,GO:0051329,GO:0051716,GO:0060249,GO:0061458,GO:0065007,GO:0065008,GO:0070013,GO:0071704,GO:0071822,GO:0071840,GO:0090304,GO:1901360,GO:1901576 | polymerase (DNA directed), delta 2, accessory subunit |
| comp220804_c1_seq3.p1 | | coiled-coil domain containing 88A |

| | | |
|---|---|---|
| comp9905_c0_seq1.p1 | GO:0000003,GO:0001558,GO:0001653,GO:0001664,GO:0001990,GO:0001992,GO:0002027,GO:0002118,GO:0002125,GO:0002791,GO:0003008,GO:0003013,GO:0003014,GO:0003044,GO:0003073,GO:0003084,GO:0003674,GO:0004871,GO:0004872,GO:0004888,GO:0004930,GO:0005000,GO:0005080,GO:0005102,GO:0005488,GO:0005515,GO:0005575,GO:0005622,GO:0005623,GO:0005737,GO:0005768,GO:0005886,GO:0005887,GO:0005981,GO:0006091,GO:0006109,GO:0006873,GO:0006874,GO:0006875,GO:0006950,GO:0007154,GO:0007165,GO:0007166,GO:0007186,GO:0007202,GO:0007204,GO:0007275,GO:0007320,GO:0007399,GO:0007417,GO:0007420,GO:0007517,GO:0007519,GO:0007610,GO:0007617,GO:0007618,GO:0007620,GO:0007621,GO:0007625,GO:0008015,GO:0008016,GO:0008150,GO:0008152,GO:0008217,GO:0008284,GO:0008528,GO:0009414,GO:0009415,GO:0009605,GO:0009628,GO:0009719,GO:0009725,GO:0009888,GO:0009889,GO:0009891,GO:0009893,GO:0009894,GO:0009896,GO:0009987,GO:0009991,GO:0010033,GO:0010035,GO:0010460,GO:0010517,GO:0010518,GO:0010565,GO:0010604,GO:0010646,GO:0010648,GO:0010675,GO:0010676,GO:0010863,GO:0010906,GO:0010907,GO:0012506,GO:0014048,GO:0014049,GO:0014070,GO:0014706,GO:0014902,GO:0016020,GO:0016021,GO:0016023,GO:0017046,GO:0019098,GO:0019216,GO:0019217,GO:0019222,GO:0019229,GO:0019722,GO:0019725,GO:0019899,GO:0019900,GO:0019901,GO:0019932,GO:0021537,GO:0022414,GO:0023051,GO:0023052,GO:0023057,GO:0030003,GO:0030154,GO:0030307,GO:0030659,GO:0030900,GO:0031090,GO:0031224,GO:0031226,GO:0031323,GO:0031325,GO:0031326,GO:0031328,GO:0031329,GO:0031331,GO:0031392,GO:0031394,GO:0031410,GO:0031644,GO:0031645,GO:0031668,GO:0031893,GO:0031894,GO:0031960,GO:0031982,GO:0031988,GO:0032501,GO:0032502,GO:0032504,GO:0032844,GO:0032846,GO:0032847,GO:0032849,GO:0032879,GO:0032881,GO:0032890,GO:0032892,GO:0033057,GO:0033218,GO:0033554,GO:0033993,GO:0035176,GO:0035556,GO:0035809,GO:0035810,GO:0035813,GO:0035815,GO:0035914,GO:0038023,GO:0040008,GO:0042127,GO:0042221,GO:0042277,GO:0042304,GO:0042562,GO:0042592,GO:0042631,GO:0042692,GO:0042711,GO:0042713,GO:0043084,GO:0043085,GO:0043226,GO:0043227,GO:0043229,GO:0043231,GO:0043269,GO:0043270,GO:0043467,GO:0043470,GO:0043471,GO:0044057,GO:0044060,GO:0044062,GO:0044070,GO:0044093,GO:0044237,GO:0044422,GO:0044424,GO:0044425,GO:0044433,GO:0044444,GO:0044446,GO:0044459,GO:0044464,GO:0044699,GO:0044700,GO:0044702,GO:0044703,GO:0044706,GO:0044707,GO:0044708,GO:0044763,GO:0044767,GO:0045723,GO:0045777,GO:0045819,GO:0045823,GO:0045834,GO:0045907,GO:0045913,GO:0045923,GO:0045924,GO:0045927,GO:0046883,GO:0046889,GO:0046890,GO:0048513,GO:0048518,GO:0048519,GO:0048522,GO:0048523,GO:0048545,GO:0048609,GO:0048731,GO:0048856,GO:0048869,GO:0048878,GO:0050789,GO:0050790,GO:0050794,GO:0050801,GO:0050878,GO:0050886,GO:0050896,GO:0051046,GO:0051047,GO:0051049,GO:0051050,GO:0051128,GO:0051146,GO:0051239,GO:0051240,GO:0051241,GO:0051336,GO:0051345,GO:0051384,GO:0051385,GO:0051412,GO:0051459,GO:0051480,GO:0051703,GO:0051704,GO:0051705,GO:0051716,GO:0051952,GO:0051954,GO:0051955,GO:0051957,GO:0051969,GO:0051970,GO:0055065,GO:0055074,GO:0055080,GO:0055082,GO:0060089,GO:0060180,GO:0060191,GO:0060193,GO:0060255,GO:0060341,GO:0060537,GO:0060538,GO:0060746,GO:0061061,GO:0065007,GO:0065008,GO:0065009,GO:0070873,GO:0070875,GO:0070887,GO:0071214,GO:0071462,GO:0071496,GO:0071944,GO:0072503,GO:0072507,GO:0080090,GO:0090087,GO:0090276,GO:0097305,GO:1900274,GO:1901654,GO:1901700,GO:1901701,GO:2000021,GO:2001279,GO:2001280 | arginine vasopressin receptor 1A |
| comp220419_c1_seq3.p1 | GO:0005575,GO:0005622,GO:0005623,GO:0005634,GO:0043226,GO:0043227,GO:0043229,GO:0043231,GO:0044424,GO:0044464 | nuclear GTPase, germinal center associated |

| | | |
|---|---|---|
| comp214730_c0_seq1.p1 | GO:0003674,GO:0003824,GO:0004176,GO:0005575,GO:0005622,GO:0005623,GO:0005737,GO:0005739,GO:0005740,GO:0005743,GO:0006139,GO:0006152,GO:0006163,GO:0006195,GO:0006200,GO:0006725,GO:0006753,GO:0006793,GO:0006796,GO:0006807,GO:0008150,GO:0008152,GO:0008233,GO:0009056,GO:0009116,GO:0009117,GO:0009119,GO:0009123,GO:0009125,GO:0009126,GO:0009128,GO:0009141,GO:0009143,GO:0009144,GO:0009146,GO:0009150,GO:0009154,GO:0009158,GO:0009161,GO:0009164,GO:0009166,GO:0009167,GO:0009169,GO:0009199,GO:0009203,GO:0009205,GO:0009207,GO:0009259,GO:0009261,GO:0009987,GO:0016020,GO:0016462,GO:0016787,GO:0016817,GO:0016818,GO:0016887,GO:0017111,GO:0019439,GO:0019637,GO:0019693,GO:0019866,GO:0031090,GO:0031966,GO:0031967,GO:0031975,GO:0034641,GO:0034655,GO:0042278,GO:0042454,GO:0042623,GO:0043226,GO:0043227,GO:0043229,GO:0043231,GO:0044237,GO:0044238,GO:0044248,GO:0044270,GO:0044281,GO:0044422,GO:0044424,GO:0044429,GO:0044444,GO:0044446,GO:0044464,GO:0044710,GO:0046034,GO:0046128,GO:0046130,GO:0046434,GO:0046483,GO:0046700,GO:0055086,GO:0070011,GO:0071704,GO:0072521,GO:0072523,GO:1901135,GO:1901136,GO:1901292,GO:1901360,GO:1901361,GO:1901564,GO:1901565,GO:1901575,GO:1901657,GO:1901658 | Peptidase family M41 |
| comp218122_c1_seq1.p1 | GO:0000278,GO:0000280,GO:0002376,GO:0002478,GO:0002495,GO:0002504,GO:0003674,GO:0003676,GO:0003677,GO:0003774,GO:0003824,GO:0005488,GO:0005575,GO:0005622,GO:0005623,GO:0005737,GO:0005813,GO:0005815,GO:0005819,GO:0005829,GO:0005856,GO:0005871,GO:0005873,GO:0005875,GO:0006928,GO:0006950,GO:0006996,GO:0007017,GO:0007018,GO:0007049,GO:0007067,GO:0007596,GO:0007599,GO:0008150,GO:0008152,GO:0008283,GO:0009611,GO:0009987,GO:0015630,GO:0016043,GO:0016462,GO:0016787,GO:0016817,GO:0016818,GO:0017111,GO:0019882,GO:0019884,GO:0019886,GO:0022402,GO:0032501,GO:0032991,GO:0042060,GO:0043226,GO:0043228,GO:0043229,GO:0043232,GO:0043234,GO:0044422,GO:0044424,GO:0044430,GO:0044444,GO:0044446,GO:0044464,GO:0044699,GO:0044707,GO:0044763,GO:0048002,GO:0048285,GO:0050817,GO:0050878,GO:0050896,GO:0065007,GO:0065008,GO:0071840,GO:0097159,GO:1901363 | Kinesin family member 15 |
| comp212496_c3_seq2.p1 | GO:0003333,GO:0003674,GO:0005215,GO:0005342,GO:0005575,GO:0005623,GO:0005886,GO:0005887,GO:0006810,GO:0006811,GO:0006812,GO:0006813,GO:0006820,GO:0006821,GO:0006865,GO:0006884,GO:0007610,GO:0007638,GO:0008150,GO:0008324,GO:0008509,GO:0008514,GO:0009605,GO:0009612,GO:0009628,GO:0009987,GO:0010646,GO:0015075,GO:0015077,GO:0015079,GO:0015103,GO:0015108,GO:0015171,GO:0015291,GO:0015293,GO:0015294,GO:0015296,GO:0015377,GO:0015379,GO:0015672,GO:0015698,GO:0015711,GO:0015849,GO:0016020,GO:0016021,GO:0019725,GO:0022804,GO:0022820,GO:0022857,GO:0022890,GO:0022891,GO:0022892,GO:0023051,GO:0030001,GO:0031224,GO:0031226,GO:0031644,GO:0032228,GO:0034220,GO:0042592,GO:0044057,GO:0044425,GO:0044459,GO:0044464,GO:0044699,GO:0044708,GO:0044763,GO:0044765,GO:0046873,GO:0046942,GO:0046943,GO:0050789,GO:0050794,GO:0050804,GO:0050896,GO:0051179,GO:0051234,GO:0051239,GO:0051969,GO:0055085,GO:0065007,GO:0065008,GO:0071702,GO:0071705,GO:0071804,GO:0071805,GO:0071944 | Solute carrier family 12 (Potassium chloride transporters), member |

| | | |
|---|---|---|
| comp213153_c0_seq2.p1 | GO:0000003,GO:0000122,GO:0000186,GO:0000578,GO:0000902,GO:0000904,GO:0001501,GO:0001503,GO:0001525,GO:0001541,GO:0001568,GO:0001649,GO:0001654,GO:0001655,GO:0001656,GO:0001657,GO:0001658,GO:0001666,GO:0001667,GO:0001700,GO:0001701,GO:0001703,GO:0001704,GO:0001705,GO:0001707,GO:0001708,GO:0001709,GO:0001710,GO:0001712,GO:0001715,GO:0001745,GO:0001759,GO:0001763,GO:0001822,GO:0001823,GO:0001837,GO:0001838,GO:0001841,GO:0001843,GO:0001889,GO:0001932,GO:0001933,GO:0001934,GO:0001935,GO:0001936,GO:0001938,GO:0001944,GO:0001947,GO:0001958,GO:0002009,GO:0002040,GO:0002043,GO:0002062,GO:0002064,GO:0002065,GO:0002066,GO:0002067,GO:0002068,GO:0002088,GO:0002089,GO:0002119,GO:0002164,GO:0002165,GO:0002168,GO:0002244,GO:0002262,GO:0002320,GO:0002376,GO:0002520,GO:0002521,GO:0002573,GO:0002682,GO:0002683,GO:0002694,GO:0002695,GO:0003002,GO:0003006,GO:0003007,GO:0003008,GO:0003014,GO:0003128,GO:0003129,GO:0003130,GO:0003133,GO:0003134,GO:0003139,GO:0003143,GO:0003146,GO:0003154,GO:0003156,GO:0003158,GO:0003159,GO:0003170,GO:0003171,GO:0003179,GO:0003181,GO:0003197,GO:0003203,GO:0003205,GO:0003279,GO:0003303,GO:0003307,GO:0003308,GO:0003309,GO:0003323,GO:0003337,GO:0003338,GO:0003342,GO:0003343,GO:0003382,GO:0003401,GO:0003674,GO:0003824,GO:0004745,GO:0005102,GO:0005125,GO:0005126,GO:0005160,GO:0005488,GO:0005515,GO:0005518,GO:0005539,GO:0005575,GO:0005576,GO:0005578,GO:0005615,GO:0005622,GO:0005623,GO:0005737,GO:0006029,GO:0006275,GO:0006355,GO:0006357,GO:0006461,GO:0006464,GO:0006468,GO:0006469,GO:0006793,GO:0006796,GO:0006915,GO:0006928,GO:0006935,GO:0006950,GO:0006952,GO:0006954,GO:0006955,GO:0007088,GO:0007154,GO:0007165,GO:0007166,GO:0007167,GO:0007178,GO:0007179,GO:0007182,GO:0007219,GO:0007224,GO:0007267,GO:0007275,GO:0007276,GO:0007281,GO:0007292,GO:0007304,GO:0007306,GO:0007308,GO:0007309,GO:0007310,GO:0007313,GO:0007346,GO:0007350,GO:0007351,GO:0007352,GO:0007354,GO:0007368,GO:0007369,GO:0007378,GO:0007389,GO:0007391,GO:0007392,GO:0007393,GO:0007398,GO:0007399,GO:0007417,GO:0007420,GO:0007423,GO:0007424,GO:0007425,GO:0007427,GO:0007440,GO:0007442,GO:0007443,GO:0007444,GO:0007446,GO:0007447,GO:0007448,GO:0007449,GO:0007450,GO:0007455,GO:0007458,GO:0007472,GO:0007473,GO:0007474,GO:0007476,GO:0007479,GO:0007498,GO:0007500,GO:0007501,GO:0007507,GO:0007516,GO:0007517,GO:0007548,GO:0007552,GO:0007560,GO:0008047,GO:0008083,GO:0008104,GO:0008150,GO:0008152,GO:0008201,GO:0008219,GO:0008283,GO:0008284,GO:0008285,GO:0008356,GO:0008361,GO:0008406,GO:0008585,GO:0008586,GO:0008595,GO:0009100,GO:0009605,GO:0009607,GO:0009611,GO:0009612,GO:0009620,GO:0009628,GO:0009653,GO:0009719,GO:0009725,GO:0009755,GO:0009790,GO:0009791,GO:0009792,GO:0009798,GO:0009799,GO:0009855,GO:0009880,GO:0009886,GO:0009887,GO:0009888,GO:0009889,GO:0009890,GO:0009891,GO:0009892,GO:0009893,GO:0009948,GO:0009950,GO:0009952,GO:0009953,GO:0009954,GO:0009966,GO:0009967,GO:0009968,GO:0009986,GO:0009987,GO:0009994,GO:0009996,GO:0010002,GO:0010004,GO:0010033,GO:0010092,GO:0010159,GO:0010453,GO:0010454,GO:0010463,GO:0010464,GO:0010468,GO:0010470,GO:0010556,GO:0010557,GO:0010558,GO:0010562,GO:0010563,GO:0010564,GO:0010565,GO:0010566,GO:0010594,GO:0010595,GO:0010604,GO:0010605,GO:0010608,GO:0010627,GO:0010628,GO:0010629,GO:0010631,GO:0010632,GO:0010634,GO:0010639,GO:0010646,GO:0010647,GO:0010648,GO:0010668,GO:0010712,GO:0010714,GO:0010717,GO:0010718,GO:0010720,GO:0010721,GO:0010740,GO:0010741,GO:0010769,GO:0010770,GO:0010862,GO:0010894,GO:0010921,GO:0010922,GO:0010927,GO:0010941,GO:0010942,GO:0010948,GO:0012501,GO:0014013,GO:0014014,GO:0014015,GO:0014020,GO:0014031,GO:0014070,GO:0014706,GO:0016015,GO:0016043,GO:0016202,GO:0016265,GO:0016310,GO:0016331,GO:0016477,GO:0016491,GO:0016614,GO:0016616,GO:0017145,GO:0019208,GO:0019211,GO:0019216,GO:0019218,GO:0019219,GO:0019220,GO:0019222,GO:0019538,GO:0019827,GO:0019953,GO:0021532,GO:0021536,GO:0021537,GO:0021700,GO:0021871,GO:0021904,GO:0021915,GO:0021978,GO:0021983,GO:0021984,GO:0022008,GO:0022412,GO:0022414,GO:0022602,GO:0022603,GO:0022604,GO:0022607,GO:0022612,GO:0023051,GO:0023052,GO:0023056,GO:0023057,GO:0030097,GO:0030099,GO:0030111,GO | bone morpho-genetic protein |
| comp415458_c0_seq1.p1 | GO:0007275,GO:0007399,GO:0008150,GO:0009987,GO:0010721,GO:0022008,GO:0030154,GO:0032501,GO:0032502,GO:0044699,GO:0044707,GO:0044763,GO:0045595,GO:0045596,GO:0048519,GO:0048523,GO:0048699,GO:0048731,GO:0048856,GO:0048869,GO:0050767,GO:0050768,GO:0050789,GO:0050793,GO:0050794,GO:0051093,GO:0051239,GO:0051960,GO:0060284,GO:0065007,GO:2000026 | ORANGE |

| | | |
|---|---|---|
| comp215008_c0_seq6.p1 | GO:0000003,GO:0000122,GO:0000375,GO:0000377,GO:0000398,GO:0000988,GO:0000989,GO:0001071,GO:0001076,GO:0001104,GO:0001106,GO:0001191,GO:0002119,GO:0002164,GO:0003006,GO:0003674,GO:0003700,GO:0003712,GO:0003713,GO:0003714,GO:0005488,GO:0005515,GO:0005575,GO:0005622,GO:0005623,GO:0005634,GO:0005681,GO:0006139,GO:0006351,GO:0006354,GO:0006355,GO:0006357,GO:0006366,GO:0006396,GO:0006397,GO:0006725,GO:0006807,GO:0007275,GO:0007548,GO:0008150,GO:0008152,GO:0008380,GO:0009058,GO:0009059,GO:0009653,GO:0009790,GO:0009791,GO:0009792,GO:0009889,GO:0009890,GO:0009892,GO:0009987,GO:0010171,GO:0010467,GO:0010468,GO:0010556,GO:0010558,GO:0010564,GO:0010605,GO:0010608,GO:0010629,GO:0016070,GO:0016071,GO:0016246,GO:0016441,GO:0016458,GO:0018130,GO:0019219,GO:0019222,GO:0019438,GO:0022414,GO:0030529,GO:0031047,GO:0031323,GO:0031324,GO:0031326,GO:0031327,GO:0032501,GO:0032502,GO:0032774,GO:0032991,GO:0034641,GO:0034645,GO:0034654,GO:0035194,GO:0040007,GO:0040011,GO:0040020,GO:0040029,GO:0040035,GO:0042127,GO:0043170,GO:0043226,GO:0043227,GO:0043229,GO:0043231,GO:0044237,GO:0044238,GO:0044249,GO:0044260,GO:0044271,GO:0044422,GO:0044424,GO:0044428,GO:0044446,GO:0044464,GO:0044699,GO:0044707,GO:0044763,GO:0044767,GO:0045892,GO:0045934,GO:0046483,GO:0048513,GO:0048519,GO:0048523,GO:0048608,GO:0048731,GO:0048806,GO:0048856,GO:0050789,GO:0050794,GO:0051171,GO:0051172,GO:0051252,GO:0051253,GO:0051445,GO:0051726,GO:0060255,GO:0061458,GO:0065007,GO:0070064,GO:0071011,GO:0071013,GO:0071704,GO:0080090,GO:0090304,GO:1901360,GO:1901362,GO:1901576,GO:2000112,GO:2000113,GO:2001141 | Transcription elongation regulator 1 |
| comp215008_c0_seq6.p2 | GO:0000166,GO:0003674,GO:0003824,GO:0004470,GO:0005488,GO:0005575,GO:0005622,GO:0005623,GO:0005737,GO:0005739,GO:0005811,GO:0005813,GO:0005815,GO:0005829,GO:0005856,GO:0005975,GO:0005996,GO:0006006,GO:0006082,GO:0006094,GO:0006107,GO:0006108,GO:0006139,GO:0006725,GO:0006732,GO:0006733,GO:0006734,GO:0006753,GO:0006793,GO:0006796,GO:0006807,GO:0007154,GO:0007267,GO:0008150,GO:0008152,GO:0009058,GO:0009117,GO:0009987,GO:0015630,GO:0016051,GO:0016491,GO:0016614,GO:0016615,GO:0016616,GO:0019318,GO:0019319,GO:0019362,GO:0019637,GO:0019674,GO:0019752,GO:0023052,GO:0030060,GO:0030154,GO:0032502,GO:0034641,GO:0036094,GO:0043226,GO:0043227,GO:0043228,GO:0043229,GO:0043231,GO:0043232,GO:0043436,GO:0043648,GO:0044237,GO:0044238,GO:0044281,GO:0044422,GO:0044424,GO:0044430,GO:0044444,GO:0044446,GO:0044464,GO:0044699,GO:0044700,GO:0044710,GO:0044723,GO:0044763,GO:0045165,GO:0045168,GO:0046331,GO:0046364,GO:0046483,GO:0046496,GO:0048037,GO:0048869,GO:0050662,GO:0051186,GO:0051287,GO:0055086,GO:0055114,GO:0071704,GO:0072524,GO:0097159,GO:1901265,GO:1901360,GO:1901363,GO:1901564,GO:1901576 | malate dehydrogenase 1, NAD (soluble) |
| comp216189_c0_seq1.p1 | | ARM |
| comp217206_c0_seq2.p1 | GO:0003674,GO:0005488,GO:0005515,GO:0008150,GO:0019899,GO:0019902,GO:0040011 | protein phosphatase 1, regulatory subunit 21 |
| comp214397_c0_seq2.p1 | | Zinc finger protein |

| | | |
|---|---|---|
| comp214789_c1_seq4.p1 | GO:0000003,GO:0000075,GO:0000077,GO:0000080,GO:0000082,GO:0000084,GO:0000151,GO:0000152,GO:0000278,GO:0000280,GO:0002119,GO:0002164,GO:0003006,GO:0003674,GO:0003824,GO:0004842,GO:0005575,GO:0005622,GO:0005623,GO:0005634,GO:0005654,GO:0005737,GO:0005829,GO:0006464,GO:0006508,GO:0006511,GO:0006513,GO:0006810,GO:0006897,GO:0006898,GO:0006915,GO:0006928,GO:0006950,GO:0006974,GO:0006996,GO:0007049,GO:0007050,GO:0007067,GO:0007093,GO:0007095,GO:0007154,GO:0007165,GO:0007166,GO:0007219,GO:0007275,GO:0007346,GO:0007548,GO:0007568,GO:0008150,GO:0008152,GO:0008219,GO:0008283,GO:0008285,GO:0008340,GO:0008361,GO:0008406,GO:0009056,GO:0009057,GO:0009653,GO:0009790,GO:0009791,GO:0009792,GO:0009887,GO:0009893,GO:0009987,GO:0010259,GO:0010498,GO:0010564,GO:0010604,GO:0010941,GO:0010948,GO:0012501,GO:0016043,GO:0016192,GO:0016265,GO:0016477,GO:0016567,GO:0016874,GO:0016879,GO:0016881,GO:0019005,GO:0019222,GO:0019538,GO:0019787,GO:0019941,GO:0022402,GO:0022403,GO:0022414,GO:0023052,GO:0030163,GO:0031145,GO:0031146,GO:0031323,GO:0031325,GO:0031396,GO:0031398,GO:0031399,GO:0031401,GO:0031461,GO:0031570,GO:0031572,GO:0031974,GO:0031981,GO:0032268,GO:0032270,GO:0032446,GO:0032501,GO:0032502,GO:0032535,GO:0032991,GO:0033554,GO:0035556,GO:0036211,GO:0040007,GO:0040008,GO:0040011,GO:0040014,GO:0040015,GO:0042127,GO:0042981,GO:0043066,GO:0043067,GO:0043069,GO:0043085,GO:0043161,GO:0043170,GO:0043226,GO:0043227,GO:0043229,GO:0043231,GO:0043233,GO:0043234,GO:0043412,GO:0043632,GO:0044093,GO:0044237,GO:0044238,GO:0044248,GO:0044257,GO:0044260,GO:0044265,GO:0044267,GO:0044422,GO:0044424,GO:0044428,GO:0044444,GO:0044446,GO:0044464,GO:0044699,GO:0044700,GO:0044707,GO:0044763,GO:0044767,GO:0044770,GO:0044772,GO:0044773,GO:0044774,GO:0045137,GO:0045786,GO:0045926,GO:0045930,GO:0048285,GO:0048513,GO:0048518,GO:0048519,GO:0048522,GO:0048523,GO:0048608,GO:0048731,GO:0048856,GO:0048870,GO:0050789,GO:0050790,GO:0050794,GO:0050896,GO:0051179,GO:0051234,GO:0051239,GO:0051241,GO:0051246,GO:0051247,GO:0051318,GO:0051320,GO:0051325,GO:0051329,GO:0051340,GO:0051351,GO:0051437,GO:0051438,GO:0051439,GO:0051443,GO:0051603,GO:0051674,GO:0051716,GO:0051726,GO:0060255,GO:0060548,GO:0061458,GO:0065007,GO:0065008,GO:0065009,GO:0070013,GO:0070647,GO:0071704,GO:0071840,GO:0080090,GO:0090066,GO:0097190,GO:0097193,GO:1901575,GO:1901987,GO:1901988,GO:1901990,GO:1901991 | Cullin 1 |

| | | |
|---|---|---|
| comp214789_c1_seq4.p2 | GO:0000003,GO:0000075,GO:0000077,GO:0000080,GO:0000082,GO:0000084,GO:0000151,GO:0000152,GO:0000278,GO:0000280,GO:0002119,GO:0002164,GO:0003006,GO:0003674,GO:0003824,GO:0004842,GO:0005575,GO:0005622,GO:0005623,GO:0005634,GO:0005654,GO:0005737,GO:0005829,GO:0006464,GO:0006508,GO:0006511,GO:0006513,GO:0006810,GO:0006897,GO:0006898,GO:0006915,GO:0006928,GO:0006950,GO:0006974,GO:0006996,GO:0007049,GO:0007050,GO:0007067,GO:0007093,GO:0007095,GO:0007154,GO:0007165,GO:0007166,GO:0007219,GO:0007275,GO:0007346,GO:0007548,GO:0007568,GO:0008150,GO:0008152,GO:0008219,GO:0008283,GO:0008285,GO:0008340,GO:0008361,GO:0008406,GO:0009056,GO:0009057,GO:0009653,GO:0009790,GO:0009791,GO:0009792,GO:0009887,GO:0009893,GO:0009987,GO:0010259,GO:0010498,GO:0010564,GO:0010604,GO:0010941,GO:0010948,GO:0012501,GO:0016043,GO:0016192,GO:0016265,GO:0016477,GO:0016567,GO:0016874,GO:0016879,GO:0016881,GO:0019005,GO:0019222,GO:0019538,GO:0019787,GO:0019941,GO:0022402,GO:0022403,GO:0022414,GO:0023052,GO:0030163,GO:0031145,GO:0031146,GO:0031323,GO:0031325,GO:0031396,GO:0031398,GO:0031399,GO:0031401,GO:0031461,GO:0031570,GO:0031572,GO:0031974,GO:0031981,GO:0032268,GO:0032270,GO:0032446,GO:0032501,GO:0032502,GO:0032535,GO:0032991,GO:0033554,GO:0035556,GO:0036211,GO:0040007,GO:0040008,GO:0040011,GO:0040014,GO:0040015,GO:0042127,GO:0042981,GO:0043066,GO:0043067,GO:0043069,GO:0043085,GO:0043161,GO:0043170,GO:0043226,GO:0043227,GO:0043229,GO:0043231,GO:0043233,GO:0043234,GO:0043412,GO:0043632,GO:0044093,GO:0044237,GO:0044238,GO:0044248,GO:0044257,GO:0044260,GO:0044265,GO:0044267,GO:0044422,GO:0044424,GO:0044428,GO:0044444,GO:0044446,GO:0044464,GO:0044699,GO:0044700,GO:0044707,GO:0044763,GO:0044767,GO:0044770,GO:0044772,GO:0044773,GO:0044774,GO:0045137,GO:0045786,GO:0045926,GO:0045930,GO:0048285,GO:0048513,GO:0048518,GO:0048519,GO:0048522,GO:0048523,GO:0048608,GO:0048731,GO:0048856,GO:0048870,GO:0050789,GO:0050790,GO:0050794,GO:0050896,GO:0051179,GO:0051234,GO:0051239,GO:0051241,GO:0051246,GO:0051247,GO:0051318,GO:0051320,GO:0051325,GO:0051329,GO:0051340,GO:0051351,GO:0051437,GO:0051438,GO:0051439,GO:0051443,GO:0051603,GO:0051674,GO:0051716,GO:0051726,GO:0060255,GO:0060548,GO:0061458,GO:0065007,GO:0065008,GO:0065009,GO:0070013,GO:0070647,GO:0071704,GO:0071840,GO:0080090,GO:0090066,GO:0097190,GO:0097193,GO:1901575,GO:1901987,GO:1901988,GO:1901990,GO:1901991 | Cullin 1 |
| comp219686_c0_seq2.p1 | GO:0002119,GO:0002164,GO:0003008,GO:0005575,GO:0005623,GO:0007154,GO:0007267,GO:0007268,GO:0007275,GO:0008150,GO:0009790,GO:0009791,GO:0009792,GO:0009987,GO:0016020,GO:0019226,GO:0023052,GO:0032501,GO:0032502,GO:0035637,GO:0040007,GO:0044699,GO:0044700,GO:0044707,GO:0044763,GO:0044767,GO:0048856,GO:0050877 | neural proliferation, differentiation and control, 1 |
| comp215198_c7_seq16.p1 | GO:0005575,GO:0005622,GO:0005623,GO:0005634,GO:0043226,GO:0043227,GO:0043229,GO:0043231,GO:0044424,GO:0044464 | SIX homeobox 1 |
| comp213213_c0_seq1.p1 | | Inherit from NOG: Viral A-type inclusion protein |

| | | |
|---|---|---|
| comp220037_c1_seq2.p1 | GO:0000003,GO:0000022,GO:0000090,GO:0000092,GO:0000226,GO:0000278,GO:0000279,GO:0000280,GO:0000910,GO:0001654,GO:0001700,GO:0001736,GO:0001737,GO:0001738,GO:0001745,GO:0001932,GO:0002009,GO:0002165,GO:0003006,GO:0003674,GO:0003824,GO:0004672,GO:0004674,GO:0005488,GO:0005515,GO:0006464,GO:0006468,GO:0006793,GO:0006796,GO:0006810,GO:0006996,GO:0007010,GO:0007017,GO:0007049,GO:0007051,GO:0007052,GO:0007067,GO:0007097,GO:0007154,GO:0007163,GO:0007164,GO:0007165,GO:0007264,GO:0007265,GO:0007266,GO:0007275,GO:0007276,GO:0007281,GO:0007292,GO:0007300,GO:0007302,GO:0007308,GO:0007309,GO:0007389,GO:0007391,GO:0007399,GO:0007423,GO:0007444,GO:0007472,GO:0007476,GO:0007552,GO:0007560,GO:0008104,GO:0008150,GO:0008152,GO:0008289,GO:0008335,GO:0008360,GO:0008544,GO:0009653,GO:0009790,GO:0009791,GO:0009792,GO:0009798,GO:0009886,GO:0009887,GO:0009888,GO:0009913,GO:0009966,GO:0009987,GO:0009994,GO:0010646,GO:0010769,GO:0010975,GO:0016043,GO:0016301,GO:0016310,GO:0016318,GO:0016331,GO:0016740,GO:0016772,GO:0016773,GO:0019220,GO:0019222,GO:0019538,GO:0019899,GO:0019953,GO:0019992,GO:0021700,GO:0022008,GO:0022402,GO:0022403,GO:0022412,GO:0022414,GO:0022603,GO:0022604,GO:0023051,GO:0023052,GO:0030029,GO:0030030,GO:0030036,GO:0030111,GO:0030154,GO:0030182,GO:0030855,GO:0031032,GO:0031323,GO:0031344,GO:0031399,GO:0032065,GO:0032231,GO:0032268,GO:0032501,GO:0032502,GO:0032504,GO:0032507,GO:0032879,GO:0032880,GO:0032956,GO:0032970,GO:0033036,GO:0033043,GO:0034613,GO:0035107,GO:0035114,GO:0035120,GO:0035190,GO:0035191,GO:0035220,GO:0035315,GO:0035316,GO:0035317,GO:0035556,GO:0036211,GO:0040008,GO:0040023,GO:0042067,GO:0042325,GO:0043170,GO:0043412,GO:0043588,GO:0043933,GO:0044087,GO:0044237,GO:0044238,GO:0044260,GO:0044267,GO:0044699,GO:0044700,GO:0044702,GO:0044707,GO:0044763,GO:0044765,GO:0044767,GO:0045184,GO:0045185,GO:0045595,GO:0045664,GO:0048285,GO:0048468,GO:0048469,GO:0048477,GO:0048513,GO:0048519,GO:0048563,GO:0048569,GO:0048583,GO:0048592,GO:0048598,GO:0048599,GO:0048609,GO:0048610,GO:0048638,GO:0048699,GO:0048707,GO:0048729,GO:0048731,GO:0048736,GO:0048737,GO:0048749,GO:0048856,GO:0048869,GO:0050767,GO:0050770,GO:0050789,GO:0050793,GO:0050794,GO:0050896,GO:0051020,GO:0051049,GO:0051051,GO:0051128,GO:0051174,GO:0051179,GO:0051220,GO:0051223,GO:0051224,GO:0051231,GO:0051234,GO:0051235,GO:0051239,GO:0051246,GO:0051301,GO:0051322,GO:0051493,GO:0051640,GO:0051641,GO:0051647,GO:0051649,GO:0051651,GO:0051656,GO:0051716,GO:0051960,GO:0060255,GO:0060284,GO:0060429,GO:0065007,GO:0065008,GO:0070201,GO:0070727,GO:0071704,GO:0071822,GO:0071840,GO:0080090,GO:2000026,GO:2000040,GO:2000050,GO:2000095 | Rho Binding |

| | | |
|---|---|---|
| comp220037_c1_seq2.p2 | GO:0000003,GO:0000022,GO:0000090,GO:0000092,GO:0000226,GO:0000278,GO:0000279,GO:0000280,GO:0000910,GO:0001654,GO:0001700,GO:0001736,GO:0001737,GO:0001738,GO:0001745,GO:0001932,GO:0002009,GO:0002165,GO:0003006,GO:0003674,GO:0003824,GO:0004672,GO:0004674,GO:0005488,GO:0005515,GO:0006464,GO:0006468,GO:0006793,GO:0006796,GO:0006810,GO:0006996,GO:0007010,GO:0007017,GO:0007049,GO:0007051,GO:0007052,GO:0007067,GO:0007097,GO:0007154,GO:0007163,GO:0007164,GO:0007165,GO:0007264,GO:0007265,GO:0007266,GO:0007275,GO:0007276,GO:0007281,GO:0007292,GO:0007300,GO:0007302,GO:0007308,GO:0007309,GO:0007389,GO:0007391,GO:0007399,GO:0007423,GO:0007444,GO:0007472,GO:0007476,GO:0007552,GO:0007560,GO:0008104,GO:0008150,GO:0008152,GO:0008289,GO:0008335,GO:0008360,GO:0008544,GO:0009653,GO:0009790,GO:0009791,GO:0009792,GO:0009798,GO:0009886,GO:0009887,GO:0009888,GO:0009913,GO:0009966,GO:0009987,GO:0009994,GO:0010646,GO:0010769,GO:0010975,GO:0016043,GO:0016301,GO:0016310,GO:0016318,GO:0016331,GO:0016740,GO:0016772,GO:0016773,GO:0019220,GO:0019222,GO:0019538,GO:0019899,GO:0019953,GO:0019992,GO:0021700,GO:0022008,GO:0022402,GO:0022403,GO:0022412,GO:0022414,GO:0022603,GO:0022604,GO:0023051,GO:0023052,GO:0030029,GO:0030030,GO:0030036,GO:0030111,GO:0030154,GO:0030182,GO:0030855,GO:0031032,GO:0031323,GO:0031344,GO:0031399,GO:0032065,GO:0032231,GO:0032268,GO:0032501,GO:0032502,GO:0032504,GO:0032507,GO:0032879,GO:0032880,GO:0032956,GO:0032970,GO:0033036,GO:0033043,GO:0034613,GO:0035107,GO:0035114,GO:0035120,GO:0035190,GO:0035191,GO:0035220,GO:0035315,GO:0035316,GO:0035317,GO:0035556,GO:0036211,GO:0040008,GO:0040023,GO:0042067,GO:0042325,GO:0043170,GO:0043412,GO:0043588,GO:0043933,GO:0044087,GO:0044237,GO:0044238,GO:0044260,GO:0044267,GO:0044699,GO:0044700,GO:0044702,GO:0044707,GO:0044763,GO:0044765,GO:0044767,GO:0045184,GO:0045185,GO:0045595,GO:0045664,GO:0048285,GO:0048468,GO:0048469,GO:0048477,GO:0048513,GO:0048519,GO:0048563,GO:0048569,GO:0048583,GO:0048592,GO:0048598,GO:0048599,GO:0048609,GO:0048610,GO:0048638,GO:0048699,GO:0048707,GO:0048729,GO:0048731,GO:0048736,GO:0048737,GO:0048749,GO:0048856,GO:0048869,GO:0050767,GO:0050770,GO:0050789,GO:0050793,GO:0050794,GO:0050896,GO:0051020,GO:0051049,GO:0051051,GO:0051128,GO:0051174,GO:0051179,GO:0051220,GO:0051223,GO:0051224,GO:0051231,GO:0051234,GO:0051235,GO:0051239,GO:0051246,GO:0051301,GO:0051322,GO:0051493,GO:0051640,GO:0051641,GO:0051647,GO:0051649,GO:0051651,GO:0051656,GO:0051716,GO:0051960,GO:0060255,GO:0060284,GO:0060429,GO:0065007,GO:0065008,GO:0070201,GO:0070727,GO:0071704,GO:0071822,GO:0071840,GO:0080090,GO:2000026,GO:2000040,GO:2000050,GO:2000095 | Rho Binding |
| comp216493_c6_seq4.p1 | GO:0001504,GO:0001505,GO:0003008,GO:0003333,GO:0003674,GO:0005215,GO:0005277,GO:0005283,GO:0005294,GO:0005326,GO:0005328,GO:0005342,GO:0005343,GO:0005416,GO:0005488,GO:0005515,GO:0005575,GO:0005623,GO:0005886,GO:0005887,GO:0006810,GO:0006811,GO:0006812,GO:0006814,GO:0006820,GO:0006836,GO:0006865,GO:0007154,GO:0007267,GO:0007268,GO:0007274,GO:0008150,GO:0008324,GO:0008509,GO:0008514,GO:0009987,GO:0015075,GO:0015077,GO:0015081,GO:0015171,GO:0015175,GO:0015179,GO:0015187,GO:0015193,GO:0015291,GO:0015293,GO:0015294,GO:0015370,GO:0015375,GO:0015672,GO:0015711,GO:0015804,GO:0015807,GO:0015816,GO:0015824,GO:0015849,GO:0015850,GO:0015870,GO:0015871,GO:0016020,GO:0016021,GO:0019226,GO:0019904,GO:0022804,GO:0022857,GO:0022890,GO:0022891,GO:0022892,GO:0023052,GO:0030001,GO:0030165,GO:0031224,GO:0031226,GO:0032501,GO:0034220,GO:0035524,GO:0035637,GO:0035725,GO:0040011,GO:0040012,GO:0040017,GO:0044425,GO:0044456,GO:0044459,GO:0044464,GO:0044699,GO:0044700,GO:0044707,GO:0044763,GO:0044765,GO:0045202,GO:0045211,GO:0046873,GO:0046942,GO:0046943,GO:0048518,GO:0050789,GO:0050877,GO:0051179,GO:0051234,GO:0055085,GO:0065007,GO:0065008,GO:0071702,GO:0071705,GO:0071944,GO:0097060,GO:1901374,GO:1901375 | solute carrier family 6 (neurotransmitter transporter) |

| | | |
|---|---|---|
| comp203943_c0_seq2.p1 | GO:0001775,GO:0001932,GO:0001934,GO:0002119,GO:0002164,GO:0003674,GO:0003756,GO:0003824,GO:0005488,GO:0005509,GO:0005575,GO:0005622,GO:0005623,GO:0005737,GO:0005783,GO:0005788,GO:0005789,GO:0005793,GO:0005811,GO:0005886,GO:0006457,GO:0006950,GO:0006984,GO:0006986,GO:0006987,GO:0007154,GO:0007155,GO:0007165,GO:0007275,GO:0007596,GO:0007599,GO:0008150,GO:0008152,GO:0009611,GO:0009791,GO:0009893,GO:0009987,GO:0010033,GO:0010562,GO:0010604,GO:0012505,GO:0016020,GO:0016023,GO:0016337,GO:0016853,GO:0016860,GO:0016862,GO:0016864,GO:0019219,GO:0019220,GO:0019222,GO:0019538,GO:0022610,GO:0023052,GO:0030100,GO:0030168,GO:0030968,GO:0031090,GO:0031323,GO:0031325,GO:0031399,GO:0031401,GO:0031410,GO:0031974,GO:0031982,GO:0031988,GO:0032069,GO:0032075,GO:0032268,GO:0032270,GO:0032501,GO:0032502,GO:0032879,GO:0033554,GO:0033674,GO:0034109,GO:0034620,GO:0034976,GO:0035966,GO:0035967,GO:0040007,GO:0040011,GO:0042060,GO:0042175,GO:0042221,GO:0042325,GO:0042327,GO:0042470,GO:0043085,GO:0043167,GO:0043169,GO:0043170,GO:0043226,GO:0043227,GO:0043229,GO:0043231,GO:0043233,GO:0043549,GO:0044093,GO:0044237,GO:0044238,GO:0044260,GO:0044267,GO:0044422,GO:0044424,GO:0044425,GO:0044432,GO:0044444,GO:0044446,GO:0044464,GO:0044699,GO:0044700,GO:0044707,GO:0044763,GO:0045807,GO:0045859,GO:0045860,GO:0045937,GO:0046872,GO:0048518,GO:0048522,GO:0048770,GO:0050764,GO:0050766,GO:0050789,GO:0050790,GO:0050794,GO:0050817,GO:0050878,GO:0050896,GO:0051049,GO:0051050,GO:0051128,GO:0051130,GO:0051171,GO:0051174,GO:0051246,GO:0051247,GO:0051336,GO:0051338,GO:0051345,GO:0051347,GO:0051716,GO:0060255,GO:0060627,GO:0065007,GO:0065008,GO:0065009,GO:0070013,GO:0070527,GO:0070887,GO:0071310,GO:0071704,GO:0071944,GO:0080090,GO:2000425,GO:2000427 | Protein disulfide isomerase family A, member 6 |
| comp218214_c0_seq7.p1 | GO:0000003,GO:0001725,GO:0001932,GO:0001934,GO:0003674,GO:0003824,GO:0004672,GO:0004713,GO:0004714,GO:0004871,GO:0004872,GO:0004888,GO:0005011,GO:0005575,GO:0005622,GO:0005623,GO:0005856,GO:0005886,GO:0005887,GO:0006464,GO:0006468,GO:0006793,GO:0006796,GO:0006928,GO:0006950,GO:0006952,GO:0007154,GO:0007165,GO:0007166,GO:0007338,GO:0008150,GO:0008152,GO:0008284,GO:0009566,GO:0009607,GO:0009615,GO:0009893,GO:0009966,GO:0009967,GO:0009987,GO:0010033,GO:0010562,GO:0010604,GO:0010627,GO:0010646,GO:0010647,GO:0010740,GO:0015629,GO:0016020,GO:0016021,GO:0016301,GO:0016310,GO:0016740,GO:0016772,GO:0016773,GO:0019199,GO:0019220,GO:0019221,GO:0019222,GO:0019538,GO:0019953,GO:0022414,GO:0023051,GO:0023052,GO:0023056,GO:0031224,GO:0031226,GO:0031323,GO:0031325,GO:0031399,GO:0031401,GO:0032268,GO:0032270,GO:0032432,GO:0033674,GO:0034097,GO:0036005,GO:0036006,GO:0036211,GO:0038023,GO:0038145,GO:0042127,GO:0042221,GO:0042325,GO:0042327,GO:0042641,GO:0043085,GO:0043170,GO:0043226,GO:0043228,GO:0043229,GO:0043232,GO:0043405,GO:0043406,GO:0043408,GO:0043410,GO:0043412,GO:0043549,GO:0044093,GO:0044237,GO:0044238,GO:0044260,GO:0044267,GO:0044422,GO:0044424,GO:0044425,GO:0044430,GO:0044446,GO:0044459,GO:0044464,GO:0044699,GO:0044700,GO:0044702,GO:0044763,GO:0045859,GO:0045860,GO:0045937,GO:0048518,GO:0048522,GO:0048583,GO:0048584,GO:0050789,GO:0050790,GO:0050794,GO:0050896,GO:0051174,GO:0051246,GO:0051247,GO:0051338,GO:0051347,GO:0051704,GO:0051707,GO:0051716,GO:0051896,GO:0051897,GO:0060089,GO:0060255,GO:0065007,GO:0065009,GO:0070887,GO:0071310,GO:0071345,GO:0071704,GO:0071900,GO:0071902,GO:0071944,GO:0080090 | macrophage stimulating 1 receptor (c-met-related tyrosine kinase) |
| comp221153_c0_seq1.p1 | GO:0003674,GO:0005198,GO:0005200 | IG |
| comp166582_c0_seq1.p1 | | ZnF_C2H2 |
| comp220511_c0_seq15.p1 | | Inherit from NOG: pathogenesis |

| | | |
|---|---|---|
| comp220511_c0_seq15.p2 | GO:0001501,GO:0001502,GO:0001503,GO:0001894,GO:0001958,GO:0002062,GO:0003007,GO:0005575,GO:0005576,GO:0005578,GO:0005581,GO:0005583,GO:0005585,GO:0005604,GO:0005615,GO:0005622,GO:0005623,GO:0005737,GO:0006029,GO:0006950,GO:0007155,GO:0007275,GO:0007399,GO:0007417,GO:0007423,GO:0007507,GO:0008150,GO:0008152,GO:0009100,GO:0009611,GO:0009653,GO:0009719,GO:0009790,GO:0009887,GO:0009888,GO:0009987,GO:0010033,GO:0010468,GO:0010941,GO:0016043,GO:0016337,GO:0019222,GO:0019538,GO:0022610,GO:0030154,GO:0030198,GO:0030199,GO:0030278,GO:0030903,GO:0031012,GO:0031099,GO:0031101,GO:0032501,GO:0032502,GO:0033333,GO:0033339,GO:0035107,GO:0035108,GO:0036075,GO:0040007,GO:0042060,GO:0042221,GO:0042246,GO:0042471,GO:0042472,GO:0042592,GO:0042981,GO:0043062,GO:0043066,GO:0043067,GO:0043069,GO:0043170,GO:0043583,GO:0044237,GO:0044238,GO:0044260,GO:0044420,GO:0044421,GO:0044424,GO:0044464,GO:0044699,GO:0044707,GO:0044763,GO:0044767,GO:0048513,GO:0048519,GO:0048523,GO:0048562,GO:0048568,GO:0048589,GO:0048598,GO:0048646,GO:0048705,GO:0048731,GO:0048736,GO:0048839,GO:0048856,GO:0048869,GO:0048871,GO:0050789,GO:0050794,GO:0050896,GO:0051216,GO:0051239,GO:0051716,GO:0060021,GO:0060173,GO:0060174,GO:0060249,GO:0060255,GO:0060348,GO:0060349,GO:0060350,GO:0060351,GO:0060548,GO:0061448,GO:0065007,GO:0065008,GO:0070848,GO:0070887,GO:0071310,GO:0071363,GO:0071495,GO:0071599,GO:0071704,GO:0071772,GO:0071773,GO:0071840,GO:0072358,GO:0072359,GO:1901135 | Collagen, type I, alpha 1 |
| comp215270_c0_seq3.p1 | GO:0000139,GO:0001558,GO:0001701,GO:0001775,GO:0002274,GO:0002376,GO:0002682,GO:0002684,GO:0002685,GO:0002687,GO:0002688,GO:0002690,GO:0002761,GO:0003674,GO:0003824,GO:0004175,GO:0004222,GO:0005102,GO:0005178,GO:0005488,GO:0005515,GO:0005575,GO:0005622,GO:0005623,GO:0005634,GO:0005737,GO:0005794,GO:0005798,GO:0005802,GO:0005856,GO:0005886,GO:0006464,GO:0006468,GO:0006508,GO:0006509,GO:0006793,GO:0006796,GO:0006810,GO:0006913,GO:0007154,GO:0007162,GO:0007165,GO:0007166,GO:0007167,GO:0007169,GO:0007173,GO:0007219,GO:0007220,GO:0007229,GO:0007267,GO:0007275,GO:0008150,GO:0008152,GO:0008233,GO:0008237,GO:0008284,GO:0009790,GO:0009792,GO:0009986,GO:0009987,GO:0010033,GO:0010819,GO:0010820,GO:0012505,GO:0014069,GO:0016020,GO:0016021,GO:0016023,GO:0016310,GO:0016787,GO:0019538,GO:0019899,GO:0019900,GO:0019901,GO:0019904,GO:0023052,GO:0030155,GO:0030307,GO:0030334,GO:0030335,GO:0031090,GO:0031224,GO:0031410,GO:0031982,GO:0031988,GO:0032101,GO:0032103,GO:0032403,GO:0032501,GO:0032502,GO:0032879,GO:0033619,GO:0034097,GO:0034612,GO:0036211,GO:0038127,GO:0040008,GO:0040012,GO:0040017,GO:0042117,GO:0042127,GO:0042169,GO:0042221,GO:0042802,GO:0042803,GO:0042995,GO:0043009,GO:0043170,GO:0043226,GO:0043227,GO:0043228,GO:0043229,GO:0043231,GO:0043232,GO:0043412,GO:0044237,GO:0044238,GO:0044260,GO:0044267,GO:0044422,GO:0044424,GO:0044425,GO:0044430,GO:0044431,GO:0044444,GO:0044446,GO:0044456,GO:0044463,GO:0044464,GO:0044699,GO:0044700,GO:0044707,GO:0044763,GO:0044767,GO:0045202,GO:0045321,GO:0045595,GO:0045637,GO:0045670,GO:0045927,GO:0046907,GO:0046983,GO:0048518,GO:0048519,GO:0048520,GO:0048522,GO:0048523,GO:0048583,GO:0048584,GO:0048856,GO:0050789,GO:0050793,GO:0050794,GO:0050795,GO:0050896,GO:0050920,GO:0050921,GO:0051088,GO:0051089,GO:0051128,GO:0051169,GO:0051179,GO:0051234,GO:0051239,GO:0051270,GO:0051272,GO:0051641,GO:0051649,GO:0051716,GO:0065007,GO:0070011,GO:0071704,GO:0071944,GO:1901623,GO:1902105,GO:2000026,GO:2000145,GO:2000147,GO:2000401,GO:2000403,GO:2000404,GO:2000406 | ADAM metallopeptidase domain 10 |
| comp202301_c0_seq1.p1 | GO:0005575,GO:0005622,GO:0005623,GO:0005737,GO:0005777,GO:0005778,GO:0005779,GO:0016020,GO:0016021,GO:0031090,GO:0031224,GO:0031231,GO:0031300,GO:0031301,GO:0031903,GO:0042579,GO:0043226,GO:0043227,GO:0043229,GO:0043231,GO:0044422,GO:0044424,GO:0044425,GO:0044438,GO:0044439,GO:0044444,GO:0044446,GO:0044464 | Mpv17 / PMP22 family |

| | | |
|---|---|---|
| comp215575_c1_seq1.p1 | GO:0005575,GO:0005622,GO:0005623,GO:0005815,GO:0005856,GO:0005929,GO:0005932,GO:0007275,GO:0007368,GO:0007389,GO:0008150,GO:0009799,GO:0009855,GO:0015630,GO:0016020,GO:0016021,GO:0031224,GO:0031514,GO:0032501,GO:0032502,GO:0036064,GO:0042995,GO:0043226,GO:0043227,GO:0043228,GO:0043229,GO:0043231,GO:0043232,GO:0044422,GO:0044424,GO:0044425,GO:0044430,GO:0044441,GO:0044446,GO:0044463,GO:0044464,GO:0044699,GO:0044707 | determination of left/right symmetry |
| comp215575_c1_seq1.p2 | GO:0005575,GO:0005622,GO:0005623,GO:0005815,GO:0005856,GO:0005929,GO:0005932,GO:0007275,GO:0007368,GO:0007389,GO:0008150,GO:0009799,GO:0009855,GO:0015630,GO:0016020,GO:0016021,GO:0031224,GO:0031514,GO:0032501,GO:0032502,GO:0036064,GO:0042995,GO:0043226,GO:0043227,GO:0043228,GO:0043229,GO:0043231,GO:0043232,GO:0044422,GO:0044424,GO:0044425,GO:0044430,GO:0044441,GO:0044446,GO:0044463,GO:0044464,GO:0044699,GO:0044707 | Inherit from spriNOG: determination of left/right symmetry |
| comp219986_c3_seq1.p1 | | pumilio homolog 1 (Drosophila) |
| comp216680_c0_seq2.p1 | GO:0000003,GO:0000902,GO:0000904,GO:0003674,GO:0005488,GO:0005515,GO:0005575,GO:0005623,GO:0005886,GO:0006928,GO:0006935,GO:0007049,GO:0007126,GO:0007275,GO:0007399,GO:0007409,GO:0007411,GO:0008150,GO:0009605,GO:0009653,GO:0009987,GO:0010171,GO:0016020,GO:0016043,GO:0016328,GO:0016358,GO:0022008,GO:0022402,GO:0030030,GO:0030154,GO:0030182,GO:0031175,GO:0032501,GO:0032502,GO:0032989,GO:0032990,GO:0033563,GO:0040008,GO:0040009,GO:0040010,GO:0040011,GO:0040012,GO:0040014,GO:0040017,GO:0040018,GO:0042221,GO:0042330,GO:0044425,GO:0044459,GO:0044464,GO:0044699,GO:0044707,GO:0044763,GO:0044767,GO:0045927,GO:0048468,GO:0048518,GO:0048610,GO:0048666,GO:0048667,GO:0048699,GO:0048731,GO:0048812,GO:0048856,GO:0048858,GO:0048869,GO:0050789,GO:0050839,GO:0050896,GO:0051239,GO:0051240,GO:0051321,GO:0061564,GO:0065007,GO:0071840,GO:0071944,GO:0097485 | Ankyrin Repeat Protein |
| comp221093_c1_seq1.p1 | GO:0001505,GO:0001540,GO:0001654,GO:0003001,GO:0003008,GO:0003674,GO:0005488,GO:0005515,GO:0005575,GO:0005622,GO:0005623,GO:0005737,GO:0006810,GO:0006836,GO:0006887,GO:0006903,GO:0006904,GO:0007154,GO:0007163,GO:0007267,GO:0007268,GO:0007269,GO:0007275,GO:0007423,GO:0008150,GO:0009987,GO:0016079,GO:0016080,GO:0016081,GO:0016192,GO:0019226,GO:0022406,GO:0023052,GO:0023061,GO:0032501,GO:0032502,GO:0032940,GO:0033218,GO:0035637,GO:0042277,GO:0044424,GO:0044464,GO:0044699,GO:0044700,GO:0044707,GO:0044763,GO:0044765,GO:0044767,GO:0046903,GO:0048278,GO:0048489,GO:0048513,GO:0048731,GO:0048749,GO:0048856,GO:0050839,GO:0050877,GO:0051179,GO:0051234,GO:0051640,GO:0051641,GO:0051648,GO:0051649,GO:0051650,GO:0051656,GO:0065007,GO:0065008,GO:0097479,GO:0097480 | binding |
| comp221093_c1_seq1.p2 | GO:0001505,GO:0001540,GO:0001654,GO:0003001,GO:0003008,GO:0003674,GO:0005488,GO:0005515,GO:0005575,GO:0005622,GO:0005623,GO:0005737,GO:0006810,GO:0006836,GO:0006887,GO:0006903,GO:0006904,GO:0007154,GO:0007163,GO:0007267,GO:0007268,GO:0007269,GO:0007275,GO:0007423,GO:0008150,GO:0009987,GO:0016079,GO:0016080,GO:0016081,GO:0016192,GO:0019226,GO:0022406,GO:0023052,GO:0023061,GO:0032501,GO:0032502,GO:0032940,GO:0033218,GO:0035637,GO:0042277,GO:0044424,GO:0044464,GO:0044699,GO:0044700,GO:0044707,GO:0044763,GO:0044765,GO:0044767,GO:0046903,GO:0048278,GO:0048489,GO:0048513,GO:0048731,GO:0048749,GO:0048856,GO:0050839,GO:0050877,GO:0051179,GO:0051234,GO:0051640,GO:0051641,GO:0051648,GO:0051649,GO:0051650,GO:0051656,GO:0065007,GO:0065008,GO:0097479,GO:0097480 | binding |

| | | |
|---|---|---|
| comp213223_c0_seq30.p1 | GO:0003674,GO:0003824,GO:0003887,GO:0005575,GO:0005622,GO:0005623,GO:0005634,GO:0005654,GO:0005694,GO:0006139,GO:0006259,GO:0006260,GO:0006261,GO:0006281,GO:0006301,GO:0006725,GO:0006807,GO:0006950,GO:0006974,GO:0008150,GO:0008152,GO:0009058,GO:0009059,GO:0009987,GO:0016035,GO:0016740,GO:0016772,GO:0016779,GO:0018130,GO:0019438,GO:0019985,GO:0031974,GO:0031981,GO:0032991,GO:0033554,GO:0034061,GO:0034641,GO:0034645,GO:0034654,GO:0042575,GO:0043170,GO:0043226,GO:0043227,GO:0043228,GO:0043229,GO:0043231,GO:0043232,GO:0043233,GO:0043234,GO:0044237,GO:0044238,GO:0044249,GO:0044260,GO:0044271,GO:0044422,GO:0044424,GO:0044428,GO:0044446,GO:0044464,GO:0044699,GO:0044763,GO:0046483,GO:0050896,GO:0051716,GO:0070013,GO:0071704,GO:0071897,GO:0090304,GO:1901360,GO:1901362,GO:1901576 | REV3-like, polymerase (DNA directed), zeta, catalytic subunit |
| comp213605_c1_seq2.p1 | GO:0005575,GO:0005622,GO:0005623,GO:0005634,GO:0005737,GO:0043226,GO:0043227,GO:0043229,GO:0043231,GO:0044424,GO:0044464 | RWD domain containing 3 |
| comp215971_c0_seq1.p1 | | Phosphatidylinositol-specific phospholipase C, X domain containing 3 |
| comp214790_c0_seq2.p1 | GO:0003674,GO:0003676,GO:0003677,GO:0005488,GO:0005575,GO:0005622,GO:0005623,GO:0005634,GO:0006355,GO:0008150,GO:0008270,GO:0009889,GO:0010468,GO:0010556,GO:0019219,GO:0019222,GO:0031323,GO:0031326,GO:0043167,GO:0043169,GO:0043226,GO:0043227,GO:0043229,GO:0043231,GO:0044424,GO:0044464,GO:0046872,GO:0046914,GO:0050789,GO:0050794,GO:0051171,GO:0051252,GO:0060255,GO:0065007,GO:0080090,GO:0097159,GO:1901363,GO:2000112,GO:2001141 | zinc finger protein 184 |

| | | |
|---|---|---|
| comp212446_c0_seq7.p1 | GO:0000139,GO:0000975,GO:0001067,GO:0001071,GO:0001726,GO:0001919,GO:0001921,GO:0002165,GO:0003674,GO:0003676,GO:0003677,GO:0003700,GO:0005083,GO:0005096,GO:0005099,GO:0005100,GO:0005488,GO:0005515,GO:0005543,GO:0005547,GO:0005575,GO:0005622,GO:0005623,GO:0005737,GO:0005794,GO:0005795,GO:0005802,GO:0005829,GO:0005856,GO:0005886,GO:0006140,GO:0006355,GO:0006605,GO:0006606,GO:0006810,GO:0006886,GO:0006913,GO:0006996,GO:0007010,GO:0007015,GO:0007049,GO:0007154,GO:0007165,GO:0007166,GO:0007167,GO:0007178,GO:0007179,GO:0007184,GO:0007264,GO:0007275,GO:0007444,GO:0007478,GO:0007480,GO:0007552,GO:0007560,GO:0008047,GO:0008060,GO:0008104,GO:0008134,GO:0008150,GO:0008289,GO:0008360,GO:0009118,GO:0009653,GO:0009719,GO:0009790,GO:0009791,GO:0009792,GO:0009886,GO:0009887,GO:0009889,GO:0009891,GO:0009893,GO:0009894,GO:0009966,GO:0009968,GO:0009987,GO:0010033,GO:0010468,GO:0010556,GO:0010557,GO:0010604,GO:0010628,GO:0010639,GO:0010646,GO:0010648,GO:0012505,GO:0015031,GO:0016020,GO:0016043,GO:0016192,GO:0016482,GO:0017038,GO:0019219,GO:0019220,GO:0019222,GO:0022402,GO:0022603,GO:0022604,GO:0023051,GO:0023052,GO:0023056,GO:0023057,GO:0030027,GO:0030029,GO:0030036,GO:0030037,GO:0030234,GO:0030334,GO:0030336,GO:0030695,GO:0030811,GO:0031090,GO:0031252,GO:0031323,GO:0031325,GO:0031326,GO:0031328,GO:0031329,GO:0031344,GO:0031346,GO:0031410,GO:0031532,GO:0031982,GO:0031984,GO:0031985,GO:0032231,GO:0032232,GO:0032318,GO:0032319,GO:0032320,GO:0032321,GO:0032489,GO:0032501,GO:0032502,GO:0032580,GO:0032879,GO:0032956,GO:0032970,GO:0033036,GO:0033043,GO:0033121,GO:0033124,GO:0033365,GO:0034504,GO:0034613,GO:0035020,GO:0035021,GO:0035023,GO:0035024,GO:0035091,GO:0035107,GO:0035108,GO:0035114,GO:0035120,GO:0035127,GO:0035218,GO:0035556,GO:0040012,GO:0040013,GO:0042221,GO:0042995,GO:0043085,GO:0043087,GO:0043088,GO:0043089,GO:0043167,GO:0043168,GO:0043226,GO:0043227,GO:0043228,GO:0043229,GO:0043231,GO:0043232,GO:0043325,GO:0043547,GO:0043933,GO:0044087,GO:0044093,GO:0044212,GO:0044422,GO:0044424,GO:0044431,GO:0044444,GO:0044446,GO:0044464,GO:0044699,GO:0044700,GO:0044707,GO:0044744,GO:0044763,GO:0044765,GO:0044767,GO:0045184,GO:0045893,GO:0045935,GO:0046332,GO:0046578,GO:0046580,GO:0046907,GO:0048513,GO:0048518,GO:0048519,GO:0048522,GO:0048523,GO:0048563,GO:0048569,GO:0048583,GO:0048585,GO:0048707,GO:0048731,GO:0048736,GO:0048737,GO:0048856,GO:0050789,GO:0050790,GO:0050793,GO:0050794,GO:0050896,GO:0051056,GO:0051058,GO:0051128,GO:0051129,GO:0051130,GO:0051169,GO:0051170,GO:0051171,GO:0051173,GO:0051174,GO:0051179,GO:0051234,GO:0051252,GO:0051254,GO:0051270,GO:0051271,GO:0051336,GO:0051345,GO:0051489,GO:0051491,GO:0051492,GO:0051493,GO:0051494,GO:0051497,GO:0051641,GO:0051649,GO:0051716,GO:0060173,GO:0060255,GO:0060395,GO:0060491,GO:0060589,GO:0065007,GO:0065008,GO:0065009,GO:0070412,GO:0070727,GO:0070848,GO:0070887,GO:0071310,GO:0071363,GO:0071495,GO:0071559,GO:0071560,GO:0071702,GO:0071822,GO:0071840,GO:0071944,GO:0080090,GO:0090527,GO:0097159,GO:1900542,GO:1901363,GO:1901981,GO:2000112,GO:2000145,GO:2000146,GO:2001141 | ArfGAP with RhoGAP domain, ankyrin repeat and PH domain |

| | | |
|---|---|---|
| comp212446_c0_seq7.p2 | GO:0000139,GO:0000975,GO:0001067,GO:0001071,GO:0001919,GO:0001921,GO:0002165,GO:0003674,GO:0003676,GO:0003677,GO:0003700,GO:0005083,GO:0005096,GO:0005099,GO:0005100,GO:0005488,GO:0005515,GO:0005543,GO:0005547,GO:0005575,GO:0005622,GO:0005623,GO:0005737,GO:0005794,GO:0005795,GO:0005802,GO:0005829,GO:0005886,GO:0006140,GO:0006355,GO:0006605,GO:0006606,GO:0006810,GO:0006886,GO:0006913,GO:0006996,GO:0007010,GO:0007015,GO:0007049,GO:0007154,GO:0007165,GO:0007166,GO:0007167,GO:0007178,GO:0007179,GO:0007184,GO:0007264,GO:0007275,GO:0007444,GO:0007478,GO:0007480,GO:0007552,GO:0007560,GO:0008047,GO:0008060,GO:0008104,GO:0008134,GO:0008150,GO:0008289,GO:0008360,GO:0009118,GO:0009653,GO:0009719,GO:0009790,GO:0009791,GO:0009792,GO:0009886,GO:0009887,GO:0009889,GO:0009891,GO:0009893,GO:0009894,GO:0009966,GO:0009987,GO:0010033,GO:0010468,GO:0010556,GO:0010557,GO:0010604,GO:0010628,GO:0010639,GO:0010646,GO:0012505,GO:0015031,GO:0016020,GO:0016043,GO:0016482,GO:0017038,GO:0019219,GO:0019220,GO:0019222,GO:0022402,GO:0022603,GO:0022604,GO:0023051,GO:0023052,GO:0023056,GO:0030029,GO:0030036,GO:0030037,GO:0030234,GO:0030695,GO:0030811,GO:0031090,GO:0031323,GO:0031325,GO:0031326,GO:0031328,GO:0031329,GO:0031344,GO:0031346,GO:0031410,GO:0031532,GO:0031982,GO:0031984,GO:0031985,GO:0032231,GO:0032232,GO:0032318,GO:0032319,GO:0032320,GO:0032321,GO:0032489,GO:0032501,GO:0032502,GO:0032580,GO:0032879,GO:0032956,GO:0032970,GO:0033036,GO:0033043,GO:0033121,GO:0033124,GO:0033365,GO:0034504,GO:0034613,GO:0035023,GO:0035091,GO:0035107,GO:0035108,GO:0035114,GO:0035120,GO:0035127,GO:0035218,GO:0035556,GO:0042221,GO:0043085,GO:0043087,GO:0043088,GO:0043089,GO:0043167,GO:0043168,GO:0043226,GO:0043227,GO:0043229,GO:0043231,GO:0043547,GO:0043933,GO:0044087,GO:0044093,GO:0044212,GO:0044422,GO:0044424,GO:0044431,GO:0044444,GO:0044446,GO:0044464,GO:0044699,GO:0044700,GO:0044707,GO:0044744,GO:0044763,GO:0044765,GO:0044767,GO:0045184,GO:0045893,GO:0045935,GO:0046332,GO:0046578,GO:0046907,GO:0048513,GO:0048518,GO:0048519,GO:0048522,GO:0048523,GO:0048563,GO:0048569,GO:0048583,GO:0048707,GO:0048731,GO:0048736,GO:0048737,GO:0048856,GO:0050789,GO:0050790,GO:0050793,GO:0050794,GO:0050896,GO:0051056,GO:0051128,GO:0051129,GO:0051130,GO:0051169,GO:0051170,GO:0051171,GO:0051173,GO:0051174,GO:0051179,GO:0051234,GO:0051252,GO:0051254,GO:0051270,GO:0051336,GO:0051345,GO:0051489,GO:0051491,GO:0051492,GO:0051493,GO:0051494,GO:0051497,GO:0051641,GO:0051649,GO:0051716,GO:0060173,GO:0060255,GO:0060395,GO:0060491,GO:0060589,GO:0065007,GO:0065008,GO:0065009,GO:0070412,GO:0070727,GO:0070848,GO:0070887,GO:0071310,GO:0071363,GO:0071495,GO:0071559,GO:0071560,GO:0071702,GO:0071822,GO:0071840,GO:0071944,GO:0080090,GO:0090527,GO:0097159,GO:1900542,GO:1901363,GO:1901981,GO:2000112,GO:2001141 | ArfGAP with RhoGAP domain, ankyrin repeat and PH domain |

| | | |
|---|---|---|
| comp217686_c2_seq8.p1 | GO:0000151,GO:0000209,GO:0001775,GO:0002200,GO:0002204,GO:0002208,GO:0002250,GO:0002252,GO:0002263,GO:0002285,GO:0002312,GO:0002366,GO:0002376,GO:0002377,GO:0002381,GO:0002440,GO:0002443,GO:0002449,GO:0002460,GO:0002520,GO:0002562,GO:0003674,GO:0003682,GO:0003824,GO:0004842,GO:0005488,GO:0005515,GO:0005575,GO:0005622,GO:0005623,GO:0005634,GO:0005694,GO:0006139,GO:0006259,GO:0006281,GO:0006282,GO:0006302,GO:0006310,GO:0006325,GO:0006417,GO:0006448,GO:0006464,GO:0006508,GO:0006511,GO:0006513,GO:0006725,GO:0006807,GO:0006950,GO:0006955,GO:0006974,GO:0006996,GO:0007275,GO:0008150,GO:0008152,GO:0009056,GO:0009057,GO:0009314,GO:0009628,GO:0009889,GO:0009890,GO:0009892,GO:0009893,GO:0009987,GO:0010212,GO:0010390,GO:0010468,GO:0010556,GO:0010558,GO:0010604,GO:0010605,GO:0010608,GO:0016043,GO:0016064,GO:0016444,GO:0016445,GO:0016447,GO:0016567,GO:0016568,GO:0016569,GO:0016570,GO:0016574,GO:0016874,GO:0016879,GO:0016881,GO:0017148,GO:0019219,GO:0019222,GO:0019538,GO:0019724,GO:0019787,GO:0019941,GO:0030163,GO:0031323,GO:0031324,GO:0031325,GO:0031326,GO:0031327,GO:0031491,GO:0031593,GO:0032182,GO:0032268,GO:0032269,GO:0032446,GO:0032501,GO:0032502,GO:0032991,GO:0033522,GO:0033554,GO:0034641,GO:0035518,GO:0035861,GO:0036211,GO:0036297,GO:0036351,GO:0036352,GO:0042113,GO:0042393,GO:0043130,GO:0043170,GO:0043226,GO:0043227,GO:0043228,GO:0043229,GO:0043231,GO:0043232,GO:0043234,GO:0043412,GO:0043632,GO:0044237,GO:0044238,GO:0044248,GO:0044257,GO:0044260,GO:0044265,GO:0044267,GO:0044422,GO:0044424,GO:0044427,GO:0044446,GO:0044464,GO:0044699,GO:0044707,GO:0044763,GO:0045190,GO:0045321,GO:0045739,GO:0045900,GO:0045935,GO:0046483,GO:0046649,GO:0048518,GO:0048519,GO:0048522,GO:0048523,GO:0048583,GO:0048584,GO:0048731,GO:0048856,GO:0050789,GO:0050794,GO:0050896,GO:0051052,GO:0051054,GO:0051171,GO:0051173,GO:0051246,GO:0051248,GO:0051276,GO:0051603,GO:0051716,GO:0060255,GO:0065007,GO:0070530,GO:0070534,GO:0070535,GO:0070647,GO:0071704,GO:0071840,GO:0080090,GO:0080134,GO:0080135,GO:0090304,GO:1901360,GO:1901575,GO:2000112,GO:2000113,GO:2001020,GO:2001022 | ring finger protein 168, E3 ubiquitin protein ligase |
| comp216612_c0_seq5.p1 | GO:0005575,GO:0005622,GO:0005623,GO:0005737,GO:0005813,GO:0005815,GO:0005856,GO:0015630,GO:0043226,GO:0043228,GO:0043229,GO:0043232,GO:0044422,GO:0044424,GO:0044430,GO:0044446,GO:0044464 | kiaa0753 |
| comp212056_c0_seq4.p1 | GO:0005575,GO:0005622,GO:0005623,GO:0005737,GO:0044424,GO:0044464 | fem-1 homolog c (C. elegans) |
| comp213316_c2_seq1.p1 | | Zinc finger protein |

| | | |
|---|---|---|
| comp221267_c2_seq2.p1 | GO:0000003,GO:0000122,GO:0000151,GO:0000209,GO:0001558,GO:0001700,GO:0003006,GO:0003674,GO:0003824,GO:0004842,GO:0005488,GO:0005515,GO:0005575,GO:0005622,GO:0005623,GO:0005634,GO:0005654,GO:0005737,GO:0005829,GO:0005886,GO:0006139,GO:0006351,GO:0006352,GO:0006355,GO:0006357,GO:0006366,GO:0006367,GO:0006464,GO:0006508,GO:0006511,GO:0006611,GO:0006725,GO:0006807,GO:0006810,GO:0006886,GO:0006913,GO:0007154,GO:0007165,GO:0007166,GO:0007167,GO:0007178,GO:0007179,GO:0007275,GO:0007276,GO:0007292,GO:0007293,GO:0007398,GO:0007399,GO:0007442,GO:0008101,GO:0008104,GO:0008150,GO:0008152,GO:0008361,GO:0009056,GO:0009057,GO:0009058,GO:0009059,GO:0009653,GO:0009719,GO:0009790,GO:0009792,GO:0009888,GO:0009889,GO:0009890,GO:0009892,GO:0009893,GO:0009966,GO:0009968,GO:0009987,GO:0010033,GO:0010467,GO:0010468,GO:0010498,GO:0010556,GO:0010558,GO:0010604,GO:0010605,GO:0010629,GO:0010646,GO:0010648,GO:0010769,GO:0010975,GO:0015031,GO:0016020,GO:0016043,GO:0016070,GO:0016482,GO:0016567,GO:0016874,GO:0016879,GO:0016881,GO:0017015,GO:0018130,GO:0019219,GO:0019222,GO:0019438,GO:0019538,GO:0019787,GO:0019941,GO:0019953,GO:0022008,GO:0022414,GO:0022603,GO:0022604,GO:0023051,GO:0023052,GO:0023057,GO:0030154,GO:0030163,GO:0030278,GO:0030279,GO:0030509,GO:0030510,GO:0030512,GO:0030514,GO:0030516,GO:0030579,GO:0031323,GO:0031324,GO:0031325,GO:0031326,GO:0031327,GO:0031344,GO:0031396,GO:0031398,GO:0031399,GO:0031401,GO:0031974,GO:0031981,GO:0032268,GO:0032270,GO:0032446,GO:0032501,GO:0032502,GO:0032504,GO:0032535,GO:0032774,GO:0032801,GO:0032991,GO:0033036,GO:0034394,GO:0034613,GO:0034641,GO:0034645,GO:0034654,GO:0036211,GO:0040008,GO:0042221,GO:0042787,GO:0043025,GO:0043112,GO:0043161,GO:0043170,GO:0043226,GO:0043227,GO:0043229,GO:0043231,GO:0043233,GO:0043234,GO:0043412,GO:0043632,GO:0044237,GO:0044238,GO:0044248,GO:0044249,GO:0044257,GO:0044260,GO:0044265,GO:0044267,GO:0044271,GO:0044297,GO:0044422,GO:0044424,GO:0044425,GO:0044428,GO:0044444,GO:0044446,GO:0044464,GO:0044699,GO:0044700,GO:0044702,GO:0044707,GO:0044763,GO:0044765,GO:0044767,GO:0045121,GO:0045184,GO:0045595,GO:0045664,GO:0045892,GO:0045934,GO:0046483,GO:0046907,GO:0048477,GO:0048518,GO:0048519,GO:0048522,GO:0048523,GO:0048546,GO:0048565,GO:0048583,GO:0048585,GO:0048598,GO:0048609,GO:0048619,GO:0048638,GO:0048646,GO:0048699,GO:0048731,GO:0048856,GO:0048869,GO:0050767,GO:0050770,GO:0050789,GO:0050793,GO:0050794,GO:0050896,GO:0051128,GO:0051168,GO:0051169,GO:0051171,GO:0051172,GO:0051179,GO:0051234,GO:0051239,GO:0051241,GO:0051246,GO:0051247,GO:0051252,GO:0051253,GO:0051603,GO:0051641,GO:0051649,GO:0051716,GO:0051960,GO:0055123,GO:0060255,GO:0060284,GO:0061387,GO:0061525,GO:0065007,GO:0065008,GO:0070013,GO:0070647,GO:0070727,GO:0070848,GO:0070887,GO:0071310,GO:0071363,GO:0071495,GO:0071559,GO:0071560,GO:0071702,GO:0071704,GO:0071840,GO:0071944,GO:0080090,GO:0090066,GO:0090092,GO:0090101,GO:0090287,GO:0090288,GO:0090304,GO:0097458,GO:1901360,GO:1901362,GO:1901575,GO:1901576,GO:2000026,GO:2000112,GO:2000113,GO:2001141 | SMAD specific E3 ubiquitin protein ligase |
| comp221273_c0_seq1.p1 | | low density lipoprotein receptor-related protein 2 |

| | | |
|---|---|---|
| comp221273_c0_seq1.p2 | GO:0000003,GO:0002165,GO:0003674,GO:0003682,GO:0003824,GO:0004386,GO:0005488,GO:0005575,GO:0005622,GO:0005623,GO:0005634,GO:0005694,GO:0005700,GO:0005703,GO:0005705,GO:0006325,GO:0006338,GO:0006996,GO:0006997,GO:0007275,GO:0007276,GO:0007292,GO:0007338,GO:0007444,GO:0007472,GO:0007476,GO:0007552,GO:0007560,GO:0008150,GO:0008152,GO:0009566,GO:0009653,GO:0009791,GO:0009886,GO:0009887,GO:0009987,GO:0016043,GO:0016462,GO:0016568,GO:0016787,GO:0016817,GO:0016818,GO:0017111,GO:0019953,GO:0022414,GO:0022607,GO:0032501,GO:0032502,GO:0032504,GO:0034728,GO:0035039,GO:0035041,GO:0035042,GO:0035107,GO:0035114,GO:0035120,GO:0035220,GO:0043044,GO:0043226,GO:0043227,GO:0043228,GO:0043229,GO:0043231,GO:0043232,GO:0043486,GO:0043933,GO:0044085,GO:0044422,GO:0044424,GO:0044427,GO:0044446,GO:0044464,GO:0044699,GO:0044702,GO:0044707,GO:0044763,GO:0044767,GO:0048477,GO:0048513,GO:0048563,GO:0048569,GO:0048609,GO:0048610,GO:0048707,GO:0048731,GO:0048736,GO:0048737,GO:0048856,GO:0051276,GO:0070925,GO:0071824,GO:0071840 | ATP-dependent chromatin-remodeling factor which functions as substrate recognition component of the transcription regulatory histone acetylation (HAT) complex SAGA. Regulates polymerase II transcription. Also required for efficient transcription by RNA polymerase I, and more specifically the polymerase I transcription termination step. Regulates negatively DNA replication. Not only involved in transcription-related chromatin remodeling, but also required to maintain a specific chromatin configuration across the genome |
| comp218822_c1_seq4.p1 | | B-box zinc finger |
| comp219642_c2_seq3.p1 | GO:0005575,GO:0005622,GO:0005623,GO:0005737,GO:0044424,GO:0044464,GO:0045202 | phosphatase and actin regulator 1 |
| comp214192_c0_seq2.p1 | GO:0005575,GO:0005622,GO:0005623,GO:0005737,GO:0005739,GO:0005740,GO:0005743,GO:0006996,GO:0007005,GO:0008150,GO:0009987,GO:0016020,GO:0016021,GO:0016043,GO:0019866,GO:0031090,GO:0031224,GO:0031300,GO:0031301,GO:0031304,GO:0031305,GO:0031966,GO:0031967,GO:0031975,GO:0032592,GO:0043226,GO:0043227,GO:0043229,GO:0043231,GO:0044422,GO:0044424,GO:0044425,GO:0044429,GO:0044444,GO:0044446,GO:0044455,GO:0044464,GO:0044699,GO:0044763,GO:0071840 | Transmembrane protein 11 |
| comp220190_c0_seq1.p1 | | Inherit from euNOG: Glutamine and serine rich 1 |

| | | |
|---|---|---|
| comp220818_c1_seq23.p1 | GO:0001666,GO:0001775,GO:0002791,GO:0003674,GO:0004871,GO:0004872,GO:0005215,GO:0005216,GO:0005217,GO:0005218,GO:0005220,GO:0005261,GO:0005262,GO:0005488,GO:0005543,GO:0005575,GO:0005622,GO:0005623,GO:0005634,GO:0005635,GO:0005637,GO:0005730,GO:0005737,GO:0005783,GO:0005789,GO:0005856,GO:0005955,GO:0006091,GO:0006112,GO:0006810,GO:0006811,GO:0006812,GO:0006816,GO:0006873,GO:0006874,GO:0006875,GO:0006915,GO:0006950,GO:0007154,GO:0007165,GO:0007166,GO:0007167,GO:0007169,GO:0007173,GO:0007202,GO:0007204,GO:0007275,GO:0007596,GO:0007599,GO:0008150,GO:0008152,GO:0008219,GO:0008287,GO:0008289,GO:0008324,GO:0008543,GO:0009611,GO:0009628,GO:0009719,GO:0009791,GO:0009987,GO:0010033,GO:0010517,GO:0010518,GO:0010646,GO:0010863,GO:0012501,GO:0012505,GO:0012506,GO:0014069,GO:0015075,GO:0015085,GO:0015267,GO:0015276,GO:0015278,GO:0015980,GO:0016020,GO:0016023,GO:0016265,GO:0016482,GO:0016528,GO:0016529,GO:0019222,GO:0019725,GO:0019866,GO:0019932,GO:0022803,GO:0022834,GO:0022836,GO:0022838,GO:0022839,GO:0022857,GO:0022890,GO:0022891,GO:0022892,GO:0023051,GO:0023052,GO:0030001,GO:0030003,GO:0030133,GO:0030141,GO:0030168,GO:0030658,GO:0030659,GO:0030667,GO:0031088,GO:0031090,GO:0031094,GO:0031095,GO:0031410,GO:0031965,GO:0031967,GO:0031974,GO:0031975,GO:0031981,GO:0031982,GO:0031988,GO:0032469,GO:0032501,GO:0032502,GO:0032844,GO:0032879,GO:0032991,GO:0033554,GO:0034220,GO:0034976,GO:0035091,GO:0035556,GO:0036293,GO:0038023,GO:0038127,GO:0038179,GO:0042060,GO:0042175,GO:0042221,GO:0042592,GO:0042827,GO:0042995,GO:0043085,GO:0043167,GO:0043168,GO:0043226,GO:0043227,GO:0043228,GO:0043229,GO:0043231,GO:0043232,GO:0043233,GO:0043234,GO:0044093,GO:0044237,GO:0044281,GO:0044344,GO:0044422,GO:0044424,GO:0044425,GO:0044428,GO:0044430,GO:0044432,GO:0044433,GO:0044444,GO:0044446,GO:0044456,GO:0044463,GO:0044464,GO:0044699,GO:0044700,GO:0044707,GO:0044710,GO:0044763,GO:0044765,GO:0045202,GO:0046873,GO:0046883,GO:0046907,GO:0048011,GO:0048016,GO:0048519,GO:0048878,GO:0050789,GO:0050790,GO:0050794,GO:0050796,GO:0050801,GO:0050817,GO:0050878,GO:0050879,GO:0050881,GO:0050882,GO:0050896,GO:0051046,GO:0051049,GO:0051179,GO:0051209,GO:0051234,GO:0051282,GO:0051283,GO:0051336,GO:0051345,GO:0051480,GO:0051641,GO:0051649,GO:0051716,GO:0055065,GO:0055074,GO:0055080,GO:0055082,GO:0055085,GO:0055114,GO:0060089,GO:0060191,GO:0060193,GO:0060341,GO:0060401,GO:0060402,GO:0065007,GO:0065008,GO:0065009,GO:0070013,GO:0070059,GO:0070482,GO:0070588,GO:0070838,GO:0070848,GO:0070887,GO:0071310,GO:0071363,GO:0071495,GO:0071774,GO:0072503,GO:0072507,GO:0072509,GO:0072511,GO:0090087,GO:0090276,GO:0097190,GO:0097193,GO:1900274,GO:2000021 | inositol 1,4,5-trisphosphate receptor type |
| comp219151_c0_seq5.p1 | | Inherit from opiNOG: Pfam:DDE |
| comp219891_c3_seq8.p1 | GO:0001701,GO:0003674,GO:0003824,GO:0004721,GO:0004725,GO:0004871,GO:0004872,GO:0004888,GO:0005001,GO:0005488,GO:0005515,GO:0005575,GO:0005622,GO:0005623,GO:0005634,GO:0005737,GO:0005886,GO:0006464,GO:0006470,GO:0006793,GO:0006796,GO:0007154,GO:0007165,GO:0007275,GO:0007399,GO:0008150,GO:0008152,GO:0009790,GO:0009792,GO:0009986,GO:0009987,GO:0016020,GO:0016311,GO:0016787,GO:0016788,GO:0016791,GO:0019198,GO:0019538,GO:0019899,GO:0019900,GO:0019901,GO:0022008,GO:0023052,GO:0030154,GO:0030182,GO:0032501,GO:0032502,GO:0035335,GO:0036211,GO:0038023,GO:0042578,GO:0043009,GO:0043170,GO:0043226,GO:0043227,GO:0043229,GO:0043231,GO:0043412,GO:0044237,GO:0044238,GO:0044260,GO:0044267,GO:0044424,GO:0044444,GO:0044464,GO:0044699,GO:0044700,GO:0044707,GO:0044763,GO:0044767,GO:0048471,GO:0048699,GO:0048731,GO:0048856,GO:0048869,GO:0050789,GO:0050794,GO:0050896,GO:0051716,GO:0060089,GO:0065007,GO:0071704,GO:0071944 | Protein tyrosine phosphatase, receptor type, R |

| | | |
|---|---|---|
| comp219061_c2_seq1.p1 | GO:0000221,GO:0002165,GO:0003674,GO:0003824,GO:0005215,GO:0005575,GO:0005622,GO:0005623,GO:0005737,GO:0005773,GO:0005774,GO:0005886,GO:0006810,GO:0006811,GO:0006812,GO:0006818,GO:0007275,GO:0007556,GO:0007557,GO:0008150,GO:0008152,GO:0009791,GO:0009889,GO:0009987,GO:0015672,GO:0015988,GO:0015991,GO:0015992,GO:0016020,GO:0016469,GO:0016471,GO:0016787,GO:0016817,GO:0016820,GO:0019216,GO:0019222,GO:0019747,GO:0022857,GO:0031090,GO:0031323,GO:0031326,GO:0032350,GO:0032501,GO:0032502,GO:0032991,GO:0033176,GO:0033178,GO:0033180,GO:0033181,GO:0034220,GO:0043226,GO:0043227,GO:0043229,GO:0043231,GO:0043234,GO:0043455,GO:0044422,GO:0044424,GO:0044425,GO:0044437,GO:0044444,GO:0044446,GO:0044459,GO:0044464,GO:0044699,GO:0044707,GO:0044763,GO:0044765,GO:0046885,GO:0046890,GO:0050789,GO:0050794,GO:0051179,GO:0051234,GO:0055085,GO:0065007,GO:0071944,GO:0080090 | subunit C |
| comp216232_c0_seq1.p2 | GO:0002376,GO:0002520,GO:0003674,GO:0003824,GO:0005575,GO:0005622,GO:0005623,GO:0005634,GO:0005737,GO:0005829,GO:0006576,GO:0006595,GO:0006596,GO:0006598,GO:0006805,GO:0006807,GO:0007275,GO:0007568,GO:0008150,GO:0008152,GO:0008215,GO:0008340,GO:0009056,GO:0009058,GO:0009308,GO:0009309,GO:0009310,GO:0009410,GO:0009987,GO:0010259,GO:0016491,GO:0016645,GO:0016647,GO:0030097,GO:0032501,GO:0032502,GO:0034641,GO:0042221,GO:0042401,GO:0042402,GO:0043226,GO:0043227,GO:0043229,GO:0043231,GO:0044106,GO:0044237,GO:0044249,GO:0044271,GO:0044281,GO:0044424,GO:0044444,GO:0044464,GO:0044699,GO:0044707,GO:0044710,GO:0044763,GO:0044767,GO:0046208,GO:0046592,GO:0048513,GO:0048534,GO:0048731,GO:0048856,GO:0050896,GO:0051716,GO:0055114,GO:0070887,GO:0071466,GO:0071704,GO:1901564,GO:1901565,GO:1901566,GO:1901575,GO:1901576 | Spermine oxidase |
| comp209529_c0_seq1.p1 | | IQ motif containing G |
| comp218920_c2_seq1.p1 | | dnaJ homolog subfamily B member |

| | | |
|---|---|---|
| comp219279_c0_seq2.p1 | GO:0000122,GO:0000139,GO:0000902,GO:0000975,GO:0000976,GO:0000977,GO:0000978,GO:0000981,GO:0000982,GO:0000987,GO:0001012,GO:0001067,GO:0001071,GO:0001077,GO:0001159,GO:0001228,GO:0003674,GO:0003676,GO:0003677,GO:0003700,GO:0005488,GO:0005575,GO:0005622,GO:0005623,GO:0005634,GO:0005635,GO:0005737,GO:0005783,GO:0005789,GO:0005794,GO:0006082,GO:0006139,GO:0006351,GO:0006355,GO:0006357,GO:0006366,GO:0006629,GO:0006631,GO:0006633,GO:0006725,GO:0006807,GO:0006810,GO:0006897,GO:0006909,GO:0006950,GO:0006984,GO:0006991,GO:0007154,GO:0007165,GO:0008150,GO:0008152,GO:0008610,GO:0009058,GO:0009059,GO:0009653,GO:0009719,GO:0009889,GO:0009890,GO:0009891,GO:0009892,GO:0009893,GO:0009987,GO:0010033,GO:0010467,GO:0010468,GO:0010556,GO:0010557,GO:0010558,GO:0010565,GO:0010604,GO:0010605,GO:0010628,GO:0010629,GO:0010874,GO:0010883,GO:0010884,GO:0010885,GO:0010886,GO:0012505,GO:0012506,GO:0012507,GO:0016020,GO:0016021,GO:0016023,GO:0016043,GO:0016053,GO:0016070,GO:0016192,GO:0018130,GO:0019216,GO:0019217,GO:0019219,GO:0019222,GO:0019438,GO:0019752,GO:0023052,GO:0030133,GO:0030134,GO:0030135,GO:0030176,GO:0030658,GO:0030659,GO:0030662,GO:0031090,GO:0031224,GO:0031227,GO:0031300,GO:0031301,GO:0031323,GO:0031324,GO:0031325,GO:0031326,GO:0031327,GO:0031328,GO:0031410,GO:0031965,GO:0031967,GO:0031975,GO:0031982,GO:0031988,GO:0032368,GO:0032369,GO:0032371,GO:0032372,GO:0032374,GO:0032375,GO:0032502,GO:0032774,GO:0032787,GO:0032879,GO:0032933,GO:0032937,GO:0032989,GO:0032991,GO:0033554,GO:0034405,GO:0034616,GO:0034641,GO:0034645,GO:0034654,GO:0042175,GO:0042221,GO:0042304,GO:0042592,GO:0043170,GO:0043226,GO:0043227,GO:0043229,GO:0043231,GO:0043234,GO:0043436,GO:0043565,GO:0044212,GO:0044237,GO:0044238,GO:0044249,GO:0044255,GO:0044260,GO:0044271,GO:0044281,GO:0044283,GO:0044422,GO:0044424,GO:0044425,GO:0044428,GO:0044431,GO:0044432,GO:0044433,GO:0044444,GO:0044446,GO:0044464,GO:0044699,GO:0044700,GO:0044710,GO:0044711,GO:0044763,GO:0044765,GO:0044767,GO:0045892,GO:0045893,GO:0045934,GO:0045935,GO:0045944,GO:0046394,GO:0046483,GO:0046890,GO:0048518,GO:0048519,GO:0048522,GO:0048523,GO:0048856,GO:0048869,GO:0048878,GO:0050789,GO:0050794,GO:0050801,GO:0050896,GO:0051049,GO:0051051,GO:0051171,GO:0051172,GO:0051173,GO:0051179,GO:0051234,GO:0051252,GO:0051253,GO:0051254,GO:0051716,GO:0055081,GO:0055088,GO:0055089,GO:0055094,GO:0055098,GO:0060255,GO:0065007,GO:0065008,GO:0070888,GO:0071498,GO:0071499,GO:0071501,GO:0071704,GO:0071840,GO:0072330,GO:0072367,GO:0072368,GO:0080090,GO:0090304,GO:0090370,GO:0097159,GO:1901360,GO:1901362,GO:1901363,GO:1901576,GO:2000112,GO:2000113,GO:2001141 | sterol regulatory element binding transcription factor |

| | | |
|---|---|---|
| comp219279_c0_seq2.p2 | GO:0000122,GO:0000139,GO:0000902,GO:0000975,GO:0000976,GO:0000977,GO:0000978,GO:0000981,GO:0000982,GO:0000987,GO:0001012,GO:0001067,GO:0001071,GO:0001077,GO:0001159,GO:0001228,GO:0003674,GO:0003676,GO:0003677,GO:0003700,GO:0005488,GO:0005575,GO:0005622,GO:0005623,GO:0005634,GO:0005635,GO:0005737,GO:0005783,GO:0005789,GO:0005794,GO:0006082,GO:0006139,GO:0006351,GO:0006355,GO:0006357,GO:0006366,GO:0006629,GO:0006631,GO:0006633,GO:0006725,GO:0006807,GO:0006810,GO:0006897,GO:0006909,GO:0006950,GO:0006984,GO:0006991,GO:0007154,GO:0007165,GO:0008150,GO:0008152,GO:0008610,GO:0009058,GO:0009059,GO:0009653,GO:0009719,GO:0009889,GO:0009890,GO:0009891,GO:0009892,GO:0009893,GO:0009987,GO:0010033,GO:0010467,GO:0010468,GO:0010556,GO:0010557,GO:0010558,GO:0010565,GO:0010604,GO:0010605,GO:0010628,GO:0010629,GO:0010874,GO:0010883,GO:0010884,GO:0010885,GO:0010886,GO:0012505,GO:0012506,GO:0012507,GO:0016020,GO:0016021,GO:0016023,GO:0016043,GO:0016053,GO:0016070,GO:0016192,GO:0018130,GO:0019216,GO:0019217,GO:0019219,GO:0019222,GO:0019438,GO:0019752,GO:0023052,GO:0030133,GO:0030134,GO:0030135,GO:0030176,GO:0030658,GO:0030659,GO:0030662,GO:0031090,GO:0031224,GO:0031227,GO:0031300,GO:0031301,GO:0031323,GO:0031324,GO:0031325,GO:0031326,GO:0031327,GO:0031328,GO:0031410,GO:0031965,GO:0031967,GO:0031975,GO:0031982,GO:0031988,GO:0032368,GO:0032369,GO:0032371,GO:0032372,GO:0032374,GO:0032375,GO:0032502,GO:0032774,GO:0032787,GO:0032879,GO:0032933,GO:0032937,GO:0032989,GO:0032991,GO:0033554,GO:0034405,GO:0034616,GO:0034641,GO:0034645,GO:0034654,GO:0042175,GO:0042221,GO:0042304,GO:0042592,GO:0043170,GO:0043226,GO:0043227,GO:0043229,GO:0043231,GO:0043234,GO:0043436,GO:0043565,GO:0044212,GO:0044237,GO:0044238,GO:0044249,GO:0044255,GO:0044260,GO:0044271,GO:0044281,GO:0044283,GO:0044422,GO:0044424,GO:0044425,GO:0044428,GO:0044431,GO:0044432,GO:0044433,GO:0044444,GO:0044446,GO:0044464,GO:0044699,GO:0044700,GO:0044710,GO:0044711,GO:0044763,GO:0044765,GO:0044767,GO:0045892,GO:0045893,GO:0045934,GO:0045935,GO:0045944,GO:0046394,GO:0046483,GO:0046890,GO:0048518,GO:0048519,GO:0048522,GO:0048523,GO:0048856,GO:0048869,GO:0048878,GO:0050789,GO:0050794,GO:0050801,GO:0050896,GO:0051049,GO:0051051,GO:0051171,GO:0051172,GO:0051173,GO:0051179,GO:0051234,GO:0051252,GO:0051253,GO:0051254,GO:0051716,GO:0055081,GO:0055088,GO:0055089,GO:0055094,GO:0055098,GO:0060255,GO:0065007,GO:0065008,GO:0070888,GO:0071498,GO:0071499,GO:0071501,GO:0071704,GO:0071840,GO:0072330,GO:0072367,GO:0072368,GO:0080090,GO:0090304,GO:0090370,GO:0097159,GO:1901360,GO:1901362,GO:1901363,GO:1901576,GO:2000112,GO:2000113,GO:2001141 | sterol regulatory element binding transcription factor |
| comp217039_c1_seq8.p1 | GO:0000127,GO:0005575,GO:0005622,GO:0005623,GO:0005634,GO:0005654,GO:0005667,GO:0006139,GO:0006351,GO:0006383,GO:0006725,GO:0006807,GO:0007275,GO:0007517,GO:0007519,GO:0008150,GO:0008152,GO:0009058,GO:0009059,GO:0009303,GO:0009304,GO:0009888,GO:0009987,GO:0010467,GO:0014706,GO:0016070,GO:0018130,GO:0019438,GO:0030154,GO:0031974,GO:0031981,GO:0032501,GO:0032502,GO:0032774,GO:0032991,GO:0034641,GO:0034645,GO:0034654,GO:0035914,GO:0042692,GO:0042791,GO:0042797,GO:0043170,GO:0043226,GO:0043227,GO:0043229,GO:0043231,GO:0043233,GO:0043234,GO:0044237,GO:0044238,GO:0044249,GO:0044260,GO:0044271,GO:0044422,GO:0044424,GO:0044428,GO:0044446,GO:0044451,GO:0044464,GO:0044699,GO:0044707,GO:0044763,GO:0044767,GO:0046483,GO:0048513,GO:0048731,GO:0048856,GO:0048869,GO:0051146,GO:0060537,GO:0060538,GO:0061061,GO:0070013,GO:0071704,GO:0090304,GO:1901360,GO:1901362,GO:1901576 | general transcription factor IIIC, polypeptide 5 |
| comp217039_c1_seq8.p2 | GO:0003674,GO:0005488,GO:0005515 | methyltransferase like 13 |
| comp211982_c0_seq11.p1 | | |
| comp217303_c1_seq7.p1 | | Zinc finger protein 192 |
| comp201502_c0_seq4.p1 | GO:0005575,GO:0005622,GO:0005623,GO:0005634,GO:0043226,GO:0043227,GO:0043229,GO:0043231,GO:0044424,GO:0044464 | Zinc finger protein |

| | | |
|---|---|---|
| comp219878_c0_seq6.p1 | GO:0000902,GO:0000904,GO:0005575,GO:0005622,GO:0005623,GO:0005634,GO:0005737,GO:0005829,GO:0006928,GO:0006935,GO:0007154,GO:0007165,GO:0007264,GO:0007275,GO:0007399,GO:0007409,GO:0007411,GO:0008150,GO:0009605,GO:0009653,GO:0009966,GO:0009987,GO:0010646,GO:0016043,GO:0022008,GO:0023051,GO:0023052,GO:0030030,GO:0030154,GO:0030182,GO:0031175,GO:0032501,GO:0032502,GO:0032989,GO:0032990,GO:0035556,GO:0040011,GO:0042221,GO:0042330,GO:0043226,GO:0043227,GO:0043229,GO:0043231,GO:0044424,GO:0044444,GO:0044464,GO:0044699,GO:0044700,GO:0044707,GO:0044763,GO:0044767,GO:0048468,GO:0048583,GO:0048666,GO:0048667,GO:0048699,GO:0048731,GO:0048812,GO:0048856,GO:0048858,GO:0048869,GO:0050789,GO:0050794,GO:0050896,GO:0051056,GO:0051716,GO:0061564,GO:0065007,GO:0071840,GO:0097485 | Rho GTPase activating protein 39 |
| comp218906_c0_seq1.p1 | | SOCS box |
| comp218906_c0_seq1.p2 | GO:0003008,GO:0005575,GO:0005622,GO:0005623,GO:0005737,GO:0007600,GO:0007605,GO:0008150,GO:0032501,GO:0044424,GO:0044464,GO:0044699,GO:0044707,GO:0050877,GO:0050954 | zinc finger, MYND-type containing 10 |
| comp203864_c0_seq4.p1 | GO:0000313,GO:0000315,GO:0003674,GO:0003735,GO:0005198,GO:0005575,GO:0005622,GO:0005623,GO:0005737,GO:0005739,GO:0005759,GO:0005761,GO:0005762,GO:0005840,GO:0006412,GO:0008150,GO:0008152,GO:0009058,GO:0009059,GO:0009987,GO:0010467,GO:0015934,GO:0019538,GO:0030529,GO:0031974,GO:0032991,GO:0034645,GO:0043170,GO:0043226,GO:0043227,GO:0043228,GO:0043229,GO:0043231,GO:0043232,GO:0043233,GO:0044237,GO:0044238,GO:0044249,GO:0044260,GO:0044267,GO:0044391,GO:0044422,GO:0044424,GO:0044429,GO:0044444,GO:0044446,GO:0044464,GO:0070013,GO:0071704,GO:1901576 | Ribosomal protein L17 |
| comp208283_c2_seq1.p1 | GO:0000003,GO:0002119,GO:0002164,GO:0003333,GO:0003674,GO:0005215,GO:0005216,GO:0005261,GO:0005342,GO:0005427,GO:0005575,GO:0005623,GO:0005886,GO:0005887,GO:0006629,GO:0006810,GO:0006811,GO:0006812,GO:0006818,GO:0006820,GO:0006857,GO:0006865,GO:0006885,GO:0006950,GO:0006979,GO:0007049,GO:0007059,GO:0007126,GO:0007275,GO:0007568,GO:0007586,GO:0008150,GO:0008152,GO:0008324,GO:0008340,GO:0008509,GO:0008514,GO:0009266,GO:0009408,GO:0009628,GO:0009791,GO:0009987,GO:0010259,GO:0010876,GO:0015075,GO:0015171,GO:0015175,GO:0015197,GO:0015198,GO:0015252,GO:0015267,GO:0015291,GO:0015293,GO:0015294,GO:0015295,GO:0015322,GO:0015333,GO:0015672,GO:0015711,GO:0015804,GO:0015833,GO:0015849,GO:0015992,GO:0016020,GO:0016021,GO:0016247,GO:0016248,GO:0016324,GO:0019915,GO:0022402,GO:0022803,GO:0022804,GO:0022838,GO:0022857,GO:0022891,GO:0022892,GO:0022897,GO:0031224,GO:0031226,GO:0032368,GO:0032369,GO:0032501,GO:0032502,GO:0032879,GO:0032890,GO:0032891,GO:0033036,GO:0034220,GO:0035672,GO:0035673,GO:0040007,GO:0040008,GO:0040009,GO:0040010,GO:0040014,GO:0040018,GO:0042592,GO:0042886,GO:0042887,GO:0042936,GO:0042938,GO:0043269,GO:0043271,GO:0044070,GO:0044238,GO:0044425,GO:0044459,GO:0044464,GO:0044699,GO:0044707,GO:0044710,GO:0044763,GO:0044765,GO:0044767,GO:0045132,GO:0045177,GO:0045927,GO:0046942,GO:0046943,GO:0048518,GO:0048519,GO:0048610,GO:0048878,GO:0050789,GO:0050801,GO:0050896,GO:0051049,GO:0051051,GO:0051179,GO:0051234,GO:0051239,GO:0051240,GO:0051321,GO:0051952,GO:0051953,GO:0051955,GO:0051956,GO:0055067,GO:0055080,GO:0055085,GO:0065007,GO:0065008,GO:0071702,GO:0071704,GO:0071705,GO:0071944,GO:2000191,GO:2000192 | solute carrier family 15 |

| | | |
|---|---|---|
| comp88296_c0_seq1.p1 | GO:0003674,GO:0003824,GO:0003954,GO:0005575,GO:0005622,GO:0005623,GO:0005737,GO:0005739,GO:0005740,GO:0005743,GO:0005746,GO:0005747,GO:0005759,GO:0005811,GO:0006091,GO:0006119,GO:0006120,GO:0006793,GO:0006796,GO:0008137,GO:0008150,GO:0008152,GO:0009987,GO:0015980,GO:0016020,GO:0016310,GO:0016491,GO:0016651,GO:0016655,GO:0019866,GO:0022900,GO:0022904,GO:0030964,GO:0031090,GO:0031966,GO:0031967,GO:0031974,GO:0031975,GO:0032991,GO:0042773,GO:0042775,GO:0043226,GO:0043227,GO:0043229,GO:0043231,GO:0043233,GO:0043234,GO:0044237,GO:0044422,GO:0044424,GO:0044425,GO:0044429,GO:0044444,GO:0044446,GO:0044455,GO:0044464,GO:0044710,GO:0045271,GO:0045333,GO:0050136,GO:0055114,GO:0070013,GO:0070469 | NADH-ubiquinone oxidoreductase 42 kda subunit |
| comp201929_c0_seq1.p1 | | tripartite motif containing |
| comp197644_c0_seq1.p1 | GO:0005575,GO:0005576,GO:0005578,GO:0007155,GO:0008150,GO:0009987,GO:0022610,GO:0031012,GO:0044421,GO:0044699,GO:0044763 | EGF_like |
| comp213929_c0_seq2.p1 | GO:0001071,GO:0001101,GO:0001654,GO:0001666,GO:0002165,GO:0002237,GO:0003008,GO:0003674,GO:0003676,GO:0003677,GO:0003700,GO:0005488,GO:0005575,GO:0005622,GO:0005623,GO:0005634,GO:0005654,GO:0005667,GO:0005737,GO:0006091,GO:0006355,GO:0006357,GO:0006950,GO:0006996,GO:0007005,GO:0007154,GO:0007267,GO:0007268,GO:0007275,GO:0007399,GO:0007417,GO:0007423,GO:0007444,GO:0007517,GO:0007552,GO:0007560,GO:0007584,GO:0008150,GO:0008152,GO:0008582,GO:0009266,GO:0009409,GO:0009605,GO:0009607,GO:0009617,GO:0009628,GO:0009653,GO:0009719,GO:0009725,GO:0009791,GO:0009886,GO:0009887,GO:0009889,GO:0009966,GO:0009967,GO:0009987,GO:0009991,GO:0010033,GO:0010243,GO:0010468,GO:0010556,GO:0010646,GO:0010647,GO:0014070,GO:0016043,GO:0016202,GO:0019219,GO:0019222,GO:0019226,GO:0023051,GO:0023052,GO:0023056,GO:0030111,GO:0030177,GO:0031099,GO:0031100,GO:0031323,GO:0031326,GO:0031667,GO:0031974,GO:0031981,GO:0032355,GO:0032496,GO:0032501,GO:0032502,GO:0032991,GO:0033273,GO:0033993,GO:0035637,GO:0036293,GO:0040008,GO:0042221,GO:0043010,GO:0043200,GO:0043226,GO:0043227,GO:0043229,GO:0043231,GO:0043233,GO:0043234,GO:0043627,GO:0044087,GO:0044237,GO:0044422,GO:0044424,GO:0044428,GO:0044446,GO:0044451,GO:0044464,GO:0044699,GO:0044700,GO:0044707,GO:0044763,GO:0044767,GO:0045595,GO:0045886,GO:0045926,GO:0048513,GO:0048518,GO:0048519,GO:0048522,GO:0048523,GO:0048545,GO:0048563,GO:0048569,GO:0048583,GO:0048584,GO:0048634,GO:0048638,GO:0048640,GO:0048641,GO:0048707,GO:0048731,GO:0048742,GO:0048856,GO:0050789,GO:0050793,GO:0050794,GO:0050803,GO:0050807,GO:0050877,GO:0050896,GO:0051093,GO:0051128,GO:0051129,GO:0051147,GO:0051153,GO:0051171,GO:0051239,GO:0051252,GO:0051593,GO:0051602,GO:0051704,GO:0051707,GO:0051960,GO:0051961,GO:0051963,GO:0051964,GO:0060041,GO:0060255,GO:0060284,GO:0061061,GO:0065007,GO:0065008,GO:0070013,GO:0070482,GO:0071840,GO:0080090,GO:0097159,GO:0097305,GO:1901363,GO:1901698,GO:1901700,GO:1901861,GO:2000026,GO:2000112,GO:2001141 | nuclear respiratory factor 1 |

| | | |
|---|---|---|
| comp206822_c0_seq1.p1 | GO:0000187,GO:0001932,GO:0001934,GO:0002218,GO:0002221,GO:0002224,GO:0002253,GO:0002376,GO:0002682,GO:0002684,GO:0002755,GO:0002757,GO:0002758,GO:0002764,GO:0005575,GO:0005622,GO:0005623,GO:0005634,GO:0007154,GO:0007165,GO:0008150,GO:0009893,GO:0009966,GO:0009967,GO:0009987,GO:0010562,GO:0010604,GO:0010627,GO:0010646,GO:0010647,GO:0010740,GO:0019220,GO:0019222,GO:0023051,GO:0023052,GO:0023056,GO:0031323,GO:0031325,GO:0031347,GO:0031349,GO:0031399,GO:0031401,GO:0032147,GO:0032268,GO:0032270,GO:0033674,GO:0034134,GO:0034138,GO:0034142,GO:0034146,GO:0034154,GO:0035419,GO:0035681,GO:0035682,GO:0042325,GO:0042327,GO:0043085,GO:0043226,GO:0043227,GO:0043229,GO:0043231,GO:0043405,GO:0043406,GO:0043408,GO:0043410,GO:0043549,GO:0044093,GO:0044424,GO:0044464,GO:0044699,GO:0044700,GO:0044763,GO:0045088,GO:0045089,GO:0045859,GO:0045860,GO:0045937,GO:0048518,GO:0048522,GO:0048583,GO:0048584,GO:0050776,GO:0050778,GO:0050789,GO:0050790,GO:0050794,GO:0050896,GO:0051174,GO:0051246,GO:0051247,GO:0051338,GO:0051347,GO:0051716,GO:0060255,GO:0065007,GO:0065009,GO:0071900,GO:0071902,GO:0080090,GO:0080134 | Jun-like transcription factor |
| comp83104_c0_seq2.p1 | GO:0003674,GO:0003676,GO:0003723,GO:0003746,GO:0004871,GO:0005085,GO:0005488,GO:0005575,GO:0005622,GO:0005623,GO:0005634,GO:0005737,GO:0005829,GO:0005853,GO:0005856,GO:0005875,GO:0006412,GO:0006414,GO:0007154,GO:0007165,GO:0008135,GO:0008150,GO:0008152,GO:0009058,GO:0009059,GO:0009966,GO:0009967,GO:0009987,GO:0010467,GO:0010627,GO:0010646,GO:0010647,GO:0010740,GO:0010876,GO:0015630,GO:0019222,GO:0019538,GO:0019915,GO:0023051,GO:0023052,GO:0023056,GO:0030234,GO:0030695,GO:0032991,GO:0033036,GO:0034645,GO:0043122,GO:0043123,GO:0043170,GO:0043226,GO:0043227,GO:0043228,GO:0043229,GO:0043231,GO:0043232,GO:0043234,GO:0044237,GO:0044238,GO:0044249,GO:0044260,GO:0044267,GO:0044422,GO:0044424,GO:0044430,GO:0044444,GO:0044446,GO:0044464,GO:0044699,GO:0044700,GO:0044763,GO:0048518,GO:0048522,GO:0048583,GO:0048584,GO:0050789,GO:0050790,GO:0050794,GO:0050896,GO:0051179,GO:0051716,GO:0060089,GO:0060589,GO:0065007,GO:0065009,GO:0071704,GO:0097159,GO:1901363,GO:1901576 | Eukaryotic translation elongation factor 1 delta (Guanine nucleotide exchange protein) |
| comp213133_c0_seq1.p1 | | UDP-Gal betaGal beta 1,3-galactosyltransferase polypeptide 6 |
| comp219051_c1_seq3.p1 | GO:0000932,GO:0001700,GO:0003674,GO:0005488,GO:0005515,GO:0005575,GO:0005622,GO:0005623,GO:0005737,GO:0006139,GO:0006401,GO:0006402,GO:0006725,GO:0006807,GO:0007275,GO:0008150,GO:0008152,GO:0009056,GO:0009057,GO:0009790,GO:0009792,GO:0009892,GO:0009987,GO:0010468,GO:0010605,GO:0010608,GO:0010629,GO:0016070,GO:0016071,GO:0016441,GO:0016458,GO:0019222,GO:0019439,GO:0030529,GO:0031047,GO:0032501,GO:0032502,GO:0032879,GO:0032880,GO:0032991,GO:0034641,GO:0034655,GO:0035194,GO:0035195,GO:0035770,GO:0040029,GO:0043170,GO:0043226,GO:0043228,GO:0043229,GO:0043232,GO:0044237,GO:0044238,GO:0044248,GO:0044260,GO:0044265,GO:0044270,GO:0044424,GO:0044444,GO:0044464,GO:0044699,GO:0044707,GO:0044763,GO:0044767,GO:0046483,GO:0046700,GO:0048519,GO:0048856,GO:0050789,GO:0060255,GO:0065007,GO:0071704,GO:0090304,GO:1901360,GO:1901361,GO:1901575 | RNA recognition motif. (a.k.a. RRM, RBD, or RNP domain) |
| comp209795_c0_seq1.p1 | GO:0005575,GO:0005622,GO:0005623,GO:0005737,GO:0044424,GO:0044464 | Calmodulin-lysine N-methyltransferase |
| comp218497_c1_seq71.p1 | | non-SMC condensin II complex, subunit G2 |
| comp218497_c1_seq71.p2 | GO:0005575,GO:0005622,GO:0005623,GO:0005634,GO:0043226,GO:0043227,GO:0043229,GO:0043231,GO:0044424,GO:0044464 | ElaC homolog 1 (E. coli) |
| comp218497_c1_seq71.p3 | GO:0005575,GO:0005622,GO:0005623,GO:0005634,GO:0043226,GO:0043227,GO:0043229,GO:0043231,GO:0044424,GO:0044464 | ElaC homolog 1 (E. coli) |

| | | |
|---|---|---|
| comp219960_c1_seq3.p1 | GO:0007275,GO:0007423,GO:0007444,GO:0008150,GO:0022416,GO:0032501,GO:0032502,GO:0035220,GO:0044699,GO:0044707,GO:0044767,GO:0048513,GO:0048731,GO:0048856 | ZnF_C2H2 |
| comp219960_c1_seq3.p4 | GO:0000902,GO:0001700,GO:0002009,GO:0003674,GO:0005215,GO:0005575,GO:0005622,GO:0005623,GO:0005737,GO:0005783,GO:0005784,GO:0005789,GO:0005791,GO:0006605,GO:0006612,GO:0006613,GO:0006614,GO:0006616,GO:0006810,GO:0006886,GO:0007275,GO:0007391,GO:0007399,GO:0008104,GO:0008150,GO:0008219,GO:0008258,GO:0008565,GO:0009653,GO:0009790,GO:0009792,GO:0009888,GO:0009892,GO:0009894,GO:0009895,GO:0009987,GO:0010506,GO:0010507,GO:0010941,GO:0010942,GO:0012505,GO:0015031,GO:0016020,GO:0016043,GO:0016265,GO:0016331,GO:0016482,GO:0019222,GO:0022008,GO:0022892,GO:0030030,GO:0030154,GO:0030182,GO:0030867,GO:0031090,GO:0031175,GO:0031323,GO:0031324,GO:0031329,GO:0031330,GO:0032501,GO:0032502,GO:0032989,GO:0032990,GO:0032991,GO:0033036,GO:0033365,GO:0034613,GO:0042175,GO:0042335,GO:0043226,GO:0043227,GO:0043229,GO:0043231,GO:0043234,GO:0044422,GO:0044424,GO:0044425,GO:0044432,GO:0044444,GO:0044446,GO:0044464,GO:0044699,GO:0044707,GO:0044763,GO:0044765,GO:0044767,GO:0045047,GO:0045169,GO:0045184,GO:0046907,GO:0048468,GO:0048518,GO:0048519,GO:0048522,GO:0048523,GO:0048598,GO:0048666,GO:0048699,GO:0048729,GO:0048731,GO:0048812,GO:0048856,GO:0048858,GO:0048869,GO:0050789,GO:0050794,GO:0051179,GO:0051234,GO:0051641,GO:0051649,GO:0055085,GO:0060429,GO:0065002,GO:0065007,GO:0070727,GO:0070972,GO:0071256,GO:0071702,GO:0071806,GO:0071840,GO:0072594,GO:0072599 | Plug domain of Sec61p |
| comp216070_c0_seq3.p1 | GO:0005575,GO:0016020 | Transmembrane protein 248 |
| comp218060_c1_seq15.p1 | GO:0000165,GO:0000186,GO:0000902,GO:0000904,GO:0001932,GO:0001934,GO:0002119,GO:0002164,GO:0003008,GO:0003674,GO:0003824,GO:0004672,GO:0004674,GO:0004702,GO:0004709,GO:0004871,GO:0005057,GO:0005488,GO:0005515,GO:0005575,GO:0005622,GO:0005623,GO:0005737,GO:0005829,GO:0005886,GO:0006325,GO:0006355,GO:0006464,GO:0006468,GO:0006793,GO:0006796,GO:0006950,GO:0006996,GO:0007030,GO:0007154,GO:0007165,GO:0007243,GO:0007254,GO:0007267,GO:0007268,GO:0007275,GO:0007399,GO:0007409,GO:0008022,GO:0008150,GO:0008152,GO:0009611,GO:0009653,GO:0009791,GO:0009889,GO:0009893,GO:0009966,GO:0009987,GO:0010468,GO:0010556,GO:0010562,GO:0010604,GO:0010627,GO:0010646,GO:0016020,GO:0016043,GO:0016049,GO:0016301,GO:0016310,GO:0016568,GO:0016569,GO:0016570,GO:0016572,GO:0016740,GO:0016772,GO:0016773,GO:0018105,GO:0018107,GO:0018193,GO:0018209,GO:0018210,GO:0019219,GO:0019220,GO:0019222,GO:0019226,GO:0019538,GO:0019899,GO:0019900,GO:0019901,GO:0022008,GO:0023014,GO:0023051,GO:0023052,GO:0030030,GO:0030154,GO:0030182,GO:0030424,GO:0030426,GO:0030427,GO:0031098,GO:0031099,GO:0031102,GO:0031103,GO:0031175,GO:0031323,GO:0031325,GO:0031326,GO:0031399,GO:0031401,GO:0032147,GO:0032268,GO:0032270,GO:0032501,GO:0032502,GO:0032989,GO:0032990,GO:0033267,GO:0033554,GO:0033674,GO:0035556,GO:0035637,GO:0036211,GO:0040007,GO:0042325,GO:0042327,GO:0042802,GO:0042803,GO:0042995,GO:0043005,GO:0043025,GO:0043085,GO:0043170,GO:0043408,GO:0043412,GO:0043549,GO:0044093,GO:0044237,GO:0044238,GO:0044260,GO:0044267,GO:0044297,GO:0044424,GO:0044444,GO:0044463,GO:0044464,GO:0044699,GO:0044700,GO:0044707,GO:0044763,GO:0044767,GO:0045202,GO:0045859,GO:0045860,GO:0045937,GO:0046777,GO:0046983,GO:0048468,GO:0048518,GO:0048522,GO:0048583,GO:0048588,GO:0048589,GO:0048666,GO:0048667,GO:0048668,GO:0048674,GO:0048678,GO:0048682,GO:0048699,GO:0048731,GO:0048812,GO:0048856,GO:0048858,GO:0048869,GO:0050789,GO:0050790,GO:0050794,GO:0050803,GO:0050807,GO:0050877,GO:0050896,GO:0051090,GO:0051091,GO:0051092,GO:0051128,GO:0051171,GO:0051174,GO:0051246,GO:0051247,GO:0051252,GO:0051276,GO:0051338,GO:0051347,GO:0051403,GO:0051716,GO:0060089,GO:0060255,GO:0060560,GO:0061564,GO:0065007,GO:0065008,GO:0065009,GO:0071704,GO:0071840,GO:0071944,GO:0080090,GO:0097458,GO:2000112,GO:2001141 | Mitogen-activated protein kinase kinase kinase |

| | | |
|---|---|---|
| comp218060_c1_seq15.p2 | GO:0005575,GO:0005623,GO:0030424,GO:0030426,GO:0030427,GO:0042995,GO:0043005,GO:0044463,GO:0044464,GO:0097458 | |
| comp214752_c1_seq1.p1 | GO:0000003,GO:0005575,GO:0005622,GO:0005623,GO:0005737,GO:0005777,GO:0007276,GO:0007283,GO:0008150,GO:0019953,GO:0022414,GO:0032501,GO:0032504,GO:0042579,GO:0043226,GO:0043227,GO:0043229,GO:0043231,GO:0044424,GO:0044444,GO:0044464,GO:0044699,GO:0044702,GO:0048232,GO:0048609 | Coiled-coil domain containing 33 |
| comp221250_c0_seq1.p1 | | aquarius homolog (mouse) |
| comp215510_c0_seq1.p1 | | Ring finger protein 219 |
| comp215838_c1_seq2.p1 | | Inherit from KOG: muscleblind-like splicing regulator |
| comp203997_c1_seq5.p1 | GO:0000118,GO:0000228,GO:0000785,GO:0000790,GO:0005575,GO:0005622,GO:0005623,GO:0005634,GO:0005654,GO:0005694,GO:0006810,GO:0006897,GO:0006909,GO:0008150,GO:0016192,GO:0031974,GO:0031981,GO:0032991,GO:0043226,GO:0043227,GO:0043228,GO:0043229,GO:0043231,GO:0043232,GO:0043233,GO:0043234,GO:0044422,GO:0044424,GO:0044427,GO:0044428,GO:0044446,GO:0044451,GO:0044454,GO:0044464,GO:0044699,GO:0044765,GO:0051179,GO:0051234,GO:0070013,GO:0070822 | RBB1NT (NUC162) domain |
| comp208911_c0_seq2.p1 | | KR domain |
| comp220955_c1_seq8.p1 | GO:0000003,GO:0000166,GO:0000287,GO:0001817,GO:0001882,GO:0001883,GO:0003006,GO:0003674,GO:0003824,GO:0004672,GO:0004674,GO:0005488,GO:0005515,GO:0005524,GO:0005575,GO:0005622,GO:0005623,GO:0005737,GO:0005856,GO:0005886,GO:0006464,GO:0006468,GO:0006793,GO:0006796,GO:0007276,GO:0007283,GO:0008017,GO:0008092,GO:0008150,GO:0008152,GO:0009889,GO:0009987,GO:0010556,GO:0015630,GO:0015631,GO:0016020,GO:0016301,GO:0016310,GO:0016740,GO:0016772,GO:0016773,GO:0017076,GO:0019222,GO:0019538,GO:0019953,GO:0022412,GO:0022414,GO:0030054,GO:0030154,GO:0030554,GO:0031323,GO:0031326,GO:0032501,GO:0032502,GO:0032504,GO:0032549,GO:0032550,GO:0032553,GO:0032555,GO:0032559,GO:0032655,GO:0035639,GO:0036094,GO:0036211,GO:0042035,GO:0043167,GO:0043168,GO:0043169,GO:0043170,GO:0043226,GO:0043228,GO:0043229,GO:0043232,GO:0043412,GO:0044237,GO:0044238,GO:0044260,GO:0044267,GO:0044424,GO:0044464,GO:0044699,GO:0044702,GO:0044763,GO:0045075,GO:0046872,GO:0048232,GO:0048515,GO:0048609,GO:0048610,GO:0048869,GO:0050789,GO:0050794,GO:0051239,GO:0051246,GO:0060255,GO:0065007,GO:0071704,GO:0071944,GO:0080090,GO:0097159,GO:1901265,GO:1901363 | Microtubule associated serine threonine kinase 2 |
| comp213139_c1_seq1.p1 | | RNA-directed DNA polymerase from mobile element jockey-like |
| comp213139_c1_seq1.p2 | | Inherit from KOG: Retrotransposon protein |
| comp213139_c1_seq1.p3 | GO:0003674,GO:0003824,GO:0004175,GO:0004252,GO:0006508,GO:0008150,GO:0008152,GO:0008233,GO:0008236,GO:0016787,GO:0017171,GO:0019538,GO:0043170,GO:0044238,GO:0070011,GO:0071704 | Transmembrane protease serine |
| comp220446_c0_seq2.p1 | GO:0007275,GO:0008150,GO:0009653,GO:0032501,GO:0032502,GO:0033333,GO:0033334,GO:0035107,GO:0044699,GO:0044707,GO:0044767,GO:0048736,GO:0048856 | Collagen type II, alpha 1 |
| comp211842_c0_seq1.p1 | GO:0005575,GO:0005622,GO:0005623,GO:0005634,GO:0043226,GO:0043227,GO:0043229,GO:0043231,GO:0044424,GO:0044464 | BRF2, subunit of RNA polymerase III transcription initiation factor, BRF1-like |

| | | |
|---|---|---|
| comp211842_c0_seq1.p2 | GO:0005575,GO:0005622,GO:0005623,GO:0005634,GO:0005654,GO:0005681,GO:0005689,GO:0006139,GO:0006396,GO:0006725,GO:0006807,GO:0008150,GO:0008152,GO:0008380,GO:0009987,GO:0010467,GO:0016070,GO:0030529,GO:0031974,GO:0031981,GO:0032991,GO:0034641,GO:0043170,GO:0043226,GO:0043227,GO:0043229,GO:0043231,GO:0043233,GO:0044237,GO:0044238,GO:0044260,GO:0044422,GO:0044424,GO:0044428,GO:0044446,GO:0044464,GO:0046483,GO:0070013,GO:0071704,GO:0090304,GO:1901360 | Zinc finger CCHC-type and RNA binding motif 1 |
| comp215357_c1_seq5.p1 | | leucine rich repeat containing 63 |
| comp215357_c1_seq5.p2 | | tetraspanin 18 |
| comp194919_c2_seq1.p1 | | CASK interacting protein 2 |
| comp215141_c0_seq1.p1 | | Laminin G domain |

| | |
|---|---|
| ***Titin-like isoform X1*** | Key component in the assembly and functioning of striated muscles and muscle tendons. |
| ***Annetocin receptor*** | Receptor for annetocin. Activation by annetocin may induce egg-laying behavior |
| ***ATP-dependent RNA helicase* DHX58 *isoform X2*** | Involved in the innate immune response to various RNA viruses and some DNA viruses |
| ***Serine/threonine-protein kinase* TBK1** | Plays an essential role in regulating inflammatory responses to foreign agents |
| ***Sarcoplasmic/endoplasmic reticulum calcium ATPase 1*** | Key regulator of striated muscle performance |
| ***Sal-like protein 1*** | Transcriptional repressor involved in organogenesis. Necessary for maintaining stem cell pluripotency and proliferation |
| ***14-3-3 protein zeta-like*** | Involved in a wide array of cellular processes including the cell cycle, apoptosis, signal transduction, cytoskeletal rearrangements, neurogenesis, metabolism and the regulation of gene transcription |
| ***Coiled-coil and C2 domain-containing protein 1-like protein*** | Causes negative regulation of Notch during vein, eye, and bristle development in *Drosophila* . Performs essential function in controlling functional maturation of synapses in humans. |
| ***Mitogen-activated protein kinase kinase kinase 12* (MAP3K12)** | Involved in cellular responses to extracellular stimuli such as physical stress. Could aid cell adaptation to osmolarity changes in the environment. Regulates repair of UV-induced DNA damage. Promotes the expansion of myogenic precursor cells during muscle growth and regeneration |
| ***Homeodomain-interacting protein kinase 2*** | Inhibits cell growth and promotes apoptosis. Involved in the response to hypoxia. Promotes angiogenesis, and is involved in erythroid differentiation |
| ***Manganese superoxide dismutase*** | Destroys superoxide anion radicals which are normally produced within the cells and which are toxic to biological systems |
| ***Ubiquitin carboxyl-terminal hydrolase 46-like*** | Deubiquitinating enzyme that plays a role in behavior |
| ***Zinc finger homeobox protein 4*** | May play a role in neural and muscle differentiation |
| ***Putative transcription factor capicua*** | Transcriptional repressor which plays a role in development of the central nervous system |
| ***Hypoxia-inducible factor 1-alpha isoform X1*** | Master transcriptional regulator of the adaptive response to hypoxia: activates over 40 genes whose protein products increase oxygen delivery or facilitate metabolic adaptation to hypoxia. Involved in the axonal distribution and transport of mitochondria in neurons during hypoxia. |

| Contig | SNP position | ORF start | ORF end |
|---|---|---|---|
| COMP1473204_C0_SEQ1 | 46 | 221 | 781 |
| COMP148439_C0_SEQ1 | 978 | 583 | 960 |
| COMP174335_C1_SEQ1 | 150 | 550 | 152 |
| | | 167 | 517 |
| COMP175699_C0_SEQ2 | 6484 | 5164 | 5562 |
| COMP186564_C0_SEQ1 | 206 | 273 | 599 |
| COMP188521_C0_SEQ1 | 233 | 517 | 987 |
| COMP191988_C0_SEQ1 | 609 | 390 | 61 |
| COMP198175_C0_SEQ1 | 1083 | 2 | 940 |
| | | 1690 | 1379 |
| COMP198260_C0_SEQ4 | 199 | 548 | 1633 |
| COMP198260_C0_SEQ4 | 233 | 1330 | 776 |
| COMP198525_C0_SEQ2 | 579 | 1 | 525 |
| COMP199566_C1_SEQ1 | 101 | 107 | 451 |
| COMP200529_C0_SEQ2 | 1145 | 1781 | 1182 |
| COMP201466_C0_SEQ1 | 2155 | 1484 | 3 |
| | | 2724 | 1870 |
| COMP201482_C0_SEQ1 | 536 | 1139 | 591 |
| | | 550 | 855 |
| COMP201938_C0_SEQ1 | 1502 | 1576 | 2019 |
| COMP202301_C0_SEQ1 | 334 | 1131 | 394 |
| COMP202572_C0_SEQ1 | 277 | 1083 | 421 |
| COMP203463_C0_SEQ1 | 780 | 1740 | 1183 |
| COMP203597_C2_SEQ5 | 3777 | 3516 | 2035 |
| | | 3184 | 3525 |
| COMP203997_C1_SEQ5 | 2233 | 2016 | 1387 |
| | | 2844 | 2425 |
| COMP205425_C0_SEQ1 | 514 | 996 | 1 |
| COMP205553_C0_SEQ2 | 3950 | 1853 | 3112 |
| COMP205932_C1_SEQ2 | 1606 | 556 | 1311 |
| COMP206877_C0_SEQ1 | 226 | 258 | 1067 |
| COMP207023_C0_SEQ7 | 2155 | 2 | 1951 |
| COMP207023_C0_SEQ7 | 2249 | | |
| COMP207425_C0_SEQ4 | 1254 | 402 | 1061 |
| COMP207765_C1_SEQ3 | 1347 | 547 | 1251 |
| COMP207929_C0_SEQ4 | 126 | 2098 | 437 |
| COMP208251_C1_SEQ2 | 3469 | 3235 | 2735 |
| COMP208283_C2_SEQ1 | 1394 | 1 | 972 |
| COMP208585_C0_SEQ1 | 1759 | 2814 | 1840 |
| COMP208701_C0_SEQ2 | 43 | 213 | 3356 |
| COMP208775_C0_SEQ5 | 486 | 1804 | 665 |
| COMP208817_C2_SEQ1 | 893 | 68 | 889 |
| COMP209795_C0_SEQ1 | 200 | 743 | 216 |
| COMP209845_C0_SEQ2 | 1000 | 148 | 798 |
| COMP210439_C0_SEQ1 | 2270 | 2817 | 2275 |
| COMP211143_C0_SEQ1 | 561 | 158 | 469 |
| COMP211989_C2_SEQ59 | 1085 | 1 | 918 |

| | | | |
|---|---|---|---|
| COMP212759_C0_SEQ2 | 1376 | 2039 | 1701 |
| COMP213139_C1_SEQ1 | 2754 | 2796 | 3302 |
| COMP213195_C0_SEQ4 | 1472 | 1250 | 897 |
| COMP213538_C1_SEQ3 | 90 | 544 | 1746 |
| COMP213838_C0_SEQ1 | 1417 | 328 | 1218 |
| | | 1164 | 799 |
| COMP213922_C0_SEQ1 | 2829 | 513 | 2600 |
| COMP213961_C2_SEQ2 | 145 | 1264 | 188 |
| COMP214282_C0_SEQ2 | 1418 | 3 | 1274 |
| | | 1257 | 697 |
| COMP214398_C0_SEQ1 | 3178 | 2791 | 3156 |
| COMP215008_C0_SEQ6 | 790 | 1135 | 812 |
| COMP215198_C7_SEQ16 | 1922 | 1815 | 613 |
| | | 1352 | 1714 |
| COMP215526_C1_SEQ1 | 388 | 1496 | 909 |
| COMP216281_C0_SEQ13 | 3057 | 3324 | 3686 |
| COMP216493_C6_SEQ4 | 659 | 2278 | 680 |
| COMP216680_C0_SEQ2 | 684 | 2374 | 3327 |
| COMP217018_C3_SEQ6 | 620 | 4159 | 842 |
| COMP217150_C1_SEQ1 | 1130 | 2 | 895 |
| COMP217229_C4_SEQ38 | 3235 | 582 | 2972 |
| | | 2863 | 3165 |
| COMP217348_C0_SEQ9 | 2403 | 4343 | 2715 |
| COMP217615_C0_SEQ4 | 783 | 2 | 772 |
| COMP217672_C1_SEQ3 | 1210 | 4079 | 1311 |
| COMP217871_C0_SEQ5 | 1847 | 151 | 1716 |
| | | 1837 | 1517 |
| COMP218083_C0_SEQ2 | 1498 | 972 | 1334 |
| COMP218083_C0_SEQ2 | 1566 | | |
| COMP218414_C0_SEQ16 | 1744 | 176 | 1675 |
| COMP218711_C0_SEQ1 | 1425 | 2 | 1309 |
| COMP219124_C1_SEQ11 | 1380 | 2568 | 2245 |
| COMP219249_C0_SEQ5 | 1185 | 1524 | 1901 |
| COMP219472_C0_SEQ6 | 152 | 2853 | 736 |
| COMP219559_C2_SEQ7 | 3250 | 2500 | 2850 |
| COMP219642_C2_SEQ3 | 551 | 558 | 2186 |
| COMP219686_C0_SEQ2 | 2010 | 2669 | 2343 |
| COMP219846_C0_SEQ3 | 6386 | 4859 | 6319 |
| COMP219875_C0_SEQ3 | 3810 | 2 | 3337 |
| COMP220533_C0_SEQ1 | 3839 | 196 | 3525 |
| COMP220611_C0_SEQ19 | 2977 | 3881 | 3219 |
| COMP221093_C1_SEQ1 | 5527 | 5483 | 2316 |
| COMP221153_C0_SEQ1 | 186 | 397 | 4203 |
| COMP221163_C0_SEQ8 | 8637 | 7579 | 7986 |
| | | 7677 | 8063 |
| COMP254358_C0_SEQ1 | 651 | 2 | 421 |
| COMP618580_C0_SEQ1 | 1035 | 1298 | 3 |
| COMP83104_C0_SEQ2 | 576 | 67 | 981 |
